# Evolution of the *Cdk4/6*–*Cdkn2* system in invertebrates

**DOI:** 10.1101/2024.05.26.595977

**Authors:** Shiori Yuki, Shunsuke Sasaki, Yuta Yamamoto, Fumika Murakami, Kazumi Sakata, Isato Araki

## Abstract

The cell cycle is driven by cyclin-dependent kinases (Cdks). The decision whether the cell cycle proceeds is made during G1 phase, when Cdk4/6 functions. Cyclin-dependent kinase inhibitor 2 (Cdkn2) is a specific inhibitor of Cdk4/6, and their interaction depends on D84 in Cdkn2 and R24/31 in Cdk4/6. This knowledge is based mainly on studies in mammalian cells. Here, we comprehensively analyzed *Cdk4/6* and *Cdkn2* in invertebrates and found that *Cdk4/6* was present in most of the investigated phyla, but the distribution of *Cdkn2* was rather uneven among and within the phyla. The positive charge of R24/R31 in Cdk4/6 was conserved in all analyzed species in phyla with Cdkn2. The presence of *Cdkn2* and the conservation of the positive charge were statistically correlated. We also found that *Cdkn2* has been tightly linked to *Fas associated factor 1* (*Faf1*) during evolution. We discuss potential interactions between Cdkn2 and Cdk4/6 in evolution and the possible cause of the strong conservation of the microsynteny.

## Introduction

Regulation of the cell cycle is closely associated with ontogeny, tissue regeneration, cell senescence, pathogenesis, especially carcinogenesis, and evolution. Cyclin-dependent kinases (Cdks) and their regulatory subunits, cyclins, promote cell cycle, while Cdk inhibitors (Cdkns) repress the Cdk function (reviewed in Weinberg, 2014). In mammalian cells, Cdk4 or Cdk6, and cyclin Ds drive the cell cycle in G1 phase, when cell proliferation is sensitive to extracellular signals. Fish have Cdk21, the third member of the Cdk4/6 subgroup of the Cdk family (Webster et al., 2018). Mammals have three types of structurally different Cdkns, Cdkn1–Cdkn3; the former includes Cdkn1a–Cdkn1c (aka p21^Cip1/Waf1^, p27^Kip1^, and p57^Kip2^, respectively; reviewed in Muühleder *et al*., 2021). Cdkn1s preferentially inhibit Cdk1 and Cdk2 but also Cdk4/6.

Cdk inhibitor 2s (Cdkn2s) are specific for Cdk4/6. Mammals have Cdkn2a–Cdkn2d (aka p16^Ink4a^, p15^Ink4b^, p18^Ink4c^, and p18^Ink4d^, respectively); Cdkn2a and Cdkn2b have four ankyrin repeats, and Cdkn2c and Cdkn2d have five (Baumgartner et al., 1998). Cdkn1s and Cdkn2s are known as tumor suppressors (El-Deiry et al., 1993; Jen *et al*., 1994; Kamb et al., 1994; Chen et al., 1996; Zhang et al., 1997; Husemann et al., 1999; Gluick et al., 2013). In mammalian cells, Cdkn2s accumulate during cell senescence (Serrano et al., 1997; Kita et al., 2022; Liu et al., 2023). Electrostatic interaction between R24/R31 in Cdk4/6 and D84 in Cdkn2s is important for the formation of the Cdk4/6–Cdkn2 complex, which causes a structural change preventing cyclin and ATP binding to Cdk4/6 and its following activation (Brotherton et al., 1998; Russo et al., 1998; reviewed in Baker & Reddy 2013)*. Mutations R24C and R24H in CDK4 cause familial melanoma, suggesting that strong positive charge (R or possibly K, not H) at the residue is necessary for the complex formation (Wölfel et al., 1995; Zuo et al., 1996; Soufir et al., 1998; Tsao et al., 1998; Molven et al., 2005; Veinalde et al., 2013). This interaction mechanism is used in a cell immortalization protocol (Shiomi et al., 2011).

Phosphatase Cdkn3 (aka Cdk2-associated dual specificity phosphatase, Kap or Cip2) specifically inhibits Cdk2 (reviewed in Patterson et al., 2009). Unlike Cdkn1s and Cdkn2s, which act as tumor suppressors, Cdkn3 has been suggested to act as an oncogene (Liu et al., 2019). No direct interaction between Cdk4/6 and Cdkn3 has been reported.

Unlike the molecular mechanisms of the cell cycle in mammals, those in invertebrates (except *Caenorhabditis elegans* and *Drosophila melanogaster*) are poorly studied (reviewed in Kipreos & van den Heuvel, 2019). These two species have no Cdkn2, and Cdkn1 is thought to subrogate the Cdk4/6 function in G1 phase (Kipreos & van den Heuvel, 2019). On the other hand, *Cdkn1* compensates for the lack of *Cdkn2* in the embryos of *Ciona* (Urochordata; Kobayashi et al., 2022). Planarians lack Cdkn1s, and Cdh1 is important for exiting the cell cycle (Sato et al., 2022).

A few pioneering small-scale genome studies reported the existence of *Ckd4/6* genes in invertebrates, namely Chordata (*Ciona intestinalis* and *Branchiostoma floridae)*, Echinodermata (*Strongylocentrotus purpuratus*), Arthropoda (*Drosophila melanogaster*), Platyhelminthes (*Schmidtea mediterranea*), Cnidaria (*Nematostella vectensis* and *Hydra vulgaris*), and Placozoa (*Trichoplax adhaerens*) (Cao et al., 2014; Espinal-Centeno et al., 2020). No *Ckd4/6* gene was found in Porifera (*Amphimedon queenslandica*) nor *Cdkn2* gene in Platyhelminthes (*S. mediterranea*) or Cnidaria (*H. vulgaris*) (Cao et al., 2014; Espinal-Centeno et al., 2020). Considering the diversity of invertebrates, more systematic and comprehensive investigation of the regulatory mechanisms of the cell cycle is needed.

Here, we investigated the presence of *Cdkn2* and *Cdk4/6* genes in invertebrates on a large scale (61 Cdkn2 sequences from 61 invertebrate species, and 1152 Cdk4/6 sequences from 1172 invertebrate species).

## Results

### Presence of Cdkn2 in invertebrates

First, we systematically searched for *Cdkn2* in the genomes of all 24 invertebrate phyla whose genome data are available (Figure 1). During the initial keyword and blastp searches and the following more thorough tblastn searches in whole-genome shotgun contig databases, we detected *Cdkn2* only in seven phyla (Figures 1, S1, Tables 1, S1, S2). Among invertebrate Deuterostomia, we detected *Cdkn2* in Cephalochordata, Hemichordata, and Echinodermata, but not in Urochordata or Xenacoelomorpha. Among Lophotrochozoa, we detected *Cdkn2* only in Mollusca, Annelida, and Brachiopoda/Phoronida; among Mollusca, we detected *Cdkn2* in Gastropoda (except Heterobranchia) and Cephalopoda (except Coleoidea), but not in Bivalvia. Among Ecdysozoa, we found *Cdkn2* only in Onychophora. We did not find it in the basal invertebrates (non-Bilateria).

**Figure 1.**
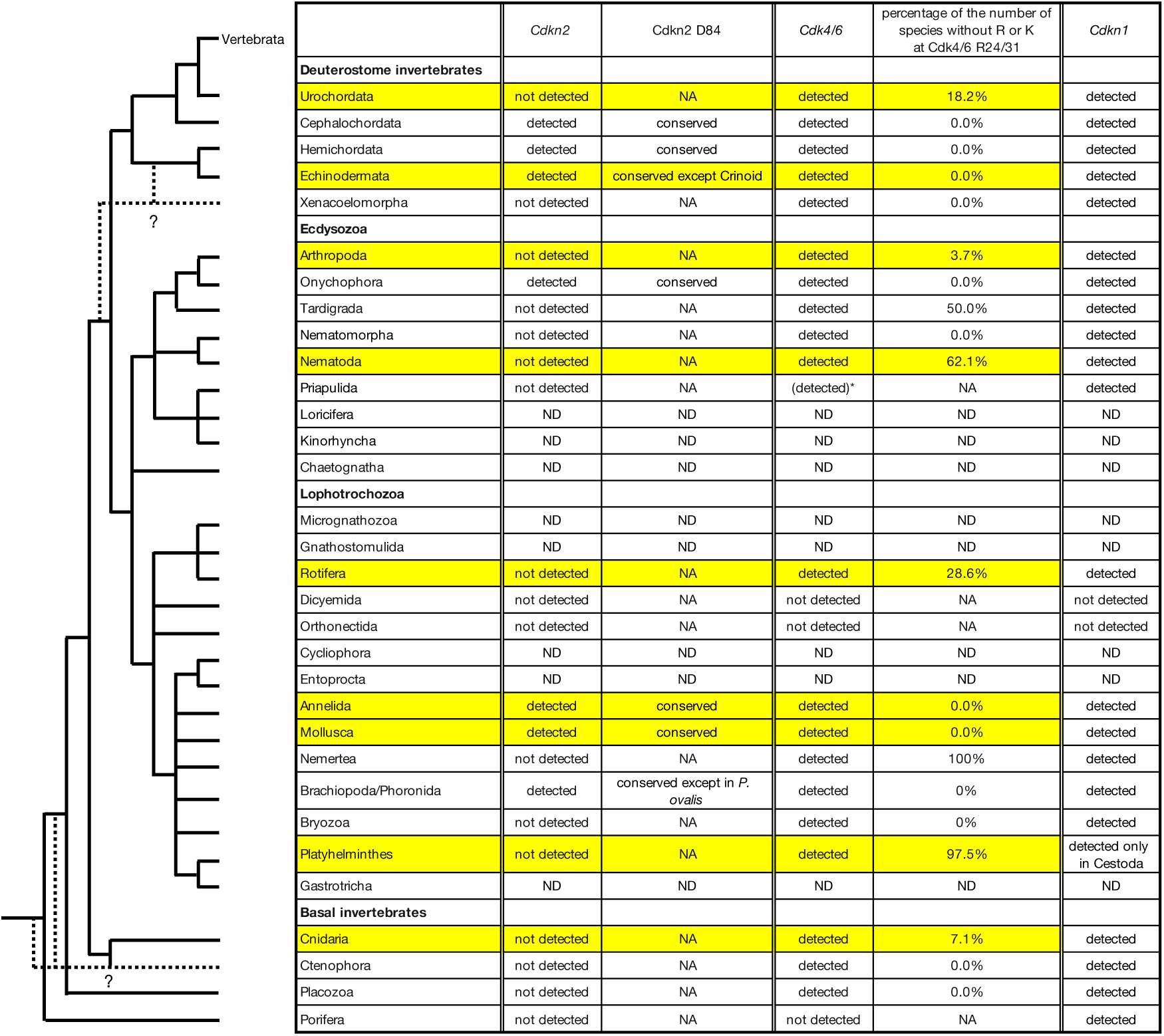
Distribution of *Cdkn2*, *Cdk4/6* and *Cdkn1* and conservation of the key charged amino acids in all invertebrate phyla. The phylogenetic tree was modified from Telford et al. (2015). The rows for phyla in which at least five species were analyzed are in yellow. * We did not include the Priapulida candidate sequence (XP_014677898.1) for the phylogenetic analysis because of the lack of a part of the PKD due to the gaps of the contig (see the main text). ND, not determined because of the lack of genome data. NA, not applicable.

**Table 1.** Detailed distribution of *Cdkn2* and *Cdk4/6* in the subtaxons of the seven phyla where *Cdkn2* was detected.

| Phyla/subtaxons | Total number of analyzed species | The number of species where *Cdk4/6* was detected | The number of species where *Cdkn2* was detected |
| --- | --- | --- | --- |
| **Cephalochordata** | 4 | 3 | 4 |
| **Hemichordata** | 2 | 1 | 2 |
| **Echinodermata** | 21 | 21 | 21 |
| Echinozoa (incl. Holothuroidea) | 10 | 7 | 10 |
| Asterozoa (incl. Ophiuroidea) | 9 | 6 | 9 |
| Pelmatozoa | 2 | 1 | 2 |
| **Onychophora** | 2 | 1 | 2 |
| **Annelida** | 9 | 7 | 6 |
| Polychaeta (incl. Pogonophora) | 7 | 6 | 5 |
| Sipuncula | 1 | 0 | 1 |
| Clitellata | 1 | 1 | 0 |
| **Mollusca** | 62 | 54 | 22 |
| Bivalvia | 29 | 29 | 0 |
| Gastropoda | 30 | 23 | 21 |
| Caenogastropoda | 11 | 5 | 10 |
| Heterobranchia | 6 | 6 | 0 |
| Neomphalina | 2 | 1 | 1 |
| Patellogastropoda | 4 | 4 | 3 |
| Vetigastropoda | 7 | 7 | 7 |
| Cephalopoda | 2 | 2 | 1 |
| Coleoidea | 2 | 2 | 0 |
| Nautiloidea | 1 | 1 | 1 |
| **Brachiopoda/Phoronida** | 4 | 2 | 4 |

During tblastn searches, we noticed sporadic “weakly positive” sequences in phyla where our searches mostly suggested the lack *Cdkn2* (Table S2b). Some of them apparently originated from contaminating DNA of other species. We used three additional criteria to clarify whether the “weakly positive” sequences other than obvious contaminating DNA might be *bona fide Cdkn2*. In all Cdkn2 candidate sequences analyzed in this study, the exon junction between Q50 and V51 was conserved (Table S1). We also determined the amino acid residues conserved among invertebrate Cdkn2s (Figure S1c). The “weakly positive” sequences failed both criteria. In the phylogenetic analysis of the Cdkn2 candidates and the human homologs of other ankyrin repeat–containing sequences with human CDKN1A as an outgroup (Tables S1 and S2), the Cdkn2 candidates formed one clade (Figure S1d; Table S3). On the basis of these data, we concluded that the “weakly positive” sequences are not *bona fide* Cdkn2 (for details, see notes in Table S2b).

Some invertebrate sequences were potentially misannotated as Cdkn2 in the NCBI Protein database (Table S4). Some of them appeared to be Cdkn1, and others were also not *bona fide* Cdkn2 but other ankyrin repeat–containing proteins on the basis of the criteria mentioned above (for details, see notes in Table S4).

We found at least one *Cdkn1* gene in all invertebrate phyla except Dicyemida and Orthonectida (Figure 1; Table S5). Among Platyhelminthes, we found *Cdkn1* in Cestoda, but not in Rhabditophora or Trematoda.

### Analysis of residues corresponding to D84 in Cdkn2

In vertebrates (reference), the residue corresponding to Cdkn2 D84 was conserved in almost all Cdkn2s (1852/1853, 99.9%; Figure S2; Table S6). The negative charge was conserved in most of the detected invertebrate *Cdkn2* (57/60, 95.0%; Figures 2, S1b). This residue was substituted in two Crinoid species (D84Q in *Anneissia japonica* and D84N in *Nesometra* (*Dorometra*) *sesokonis*) from Echinodermata and in a Brachiopoda/Phoronida species (D84N in *Phoronis ovalis*), resulting in a loss of the negative charge (Figures 2, S1b).

**Figure 2.**
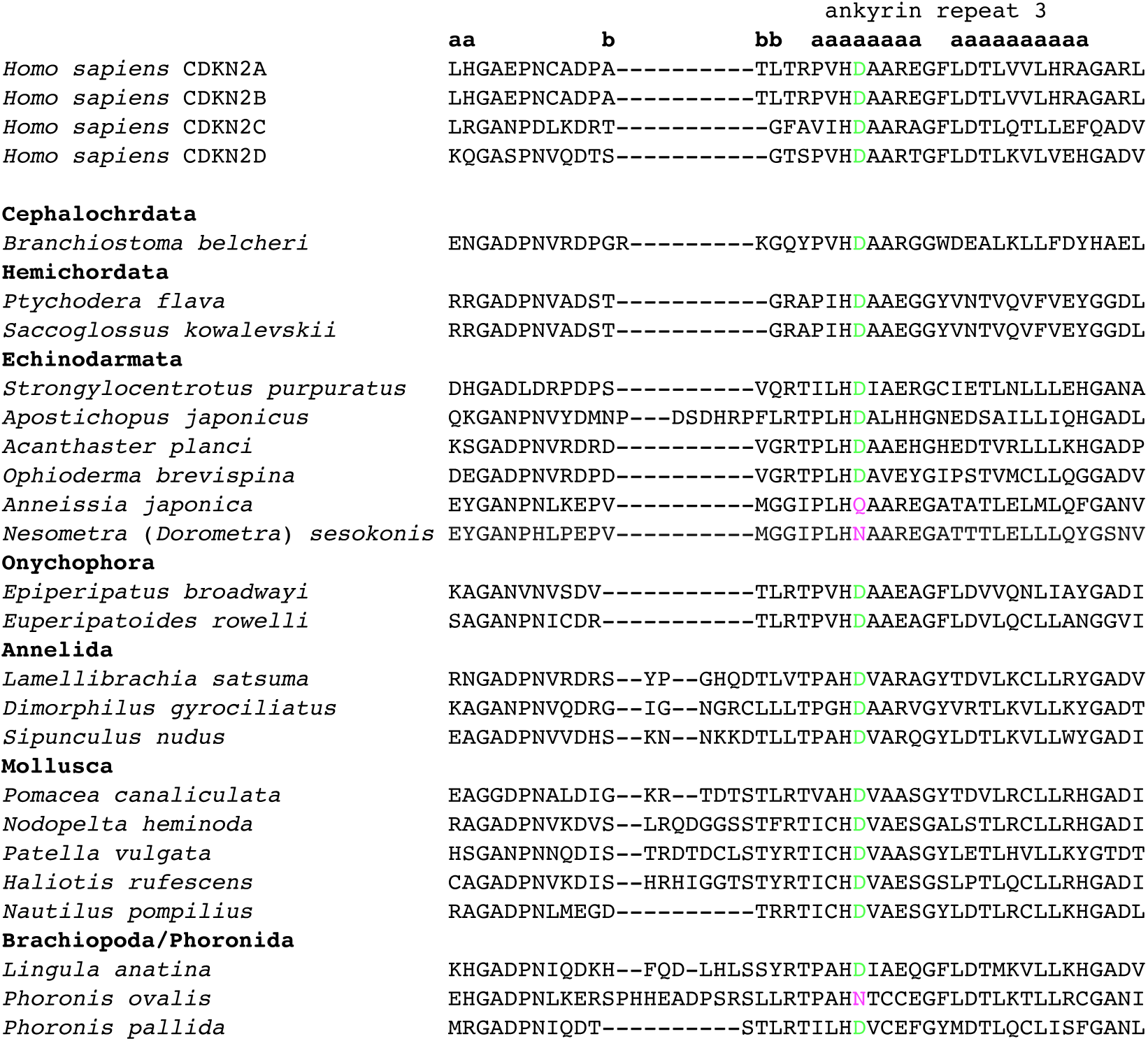
Amino acid alignment around the residue corresponding to Cdkn2 D84 in representative invertebrate species from the seven phyla where *Cdkn2* was detected. The residues corresponding to Cdkn2 D84 are in green if negatively charged or in magenta if not. Human CDKN2s are aligned as references. **a** and **b** represent the amino acids of α-helices and a β-turn in CDKN2A, respectively (Baumgartner et al., 1998).

### Microsynteny around Cdkn2 and Faf1 loci

We analyzed the microsynteny around the *Cdkn2* locus in invertebrates and found that all analyzed invertebrate *Cdkn2* genes were linked to *Fas associated factor 1* (*Faf1*), similar to vertebrate *Cdkn2c* (Figure 3, Tables S1, S7). Cephalochordate *Cdkn2* genes were linked to *methylthioadenosine phosphorylase* (*Mtap*), similar to vertebrate *Cdkn2a*/*b* (Figure 3).

**Figure 3.**
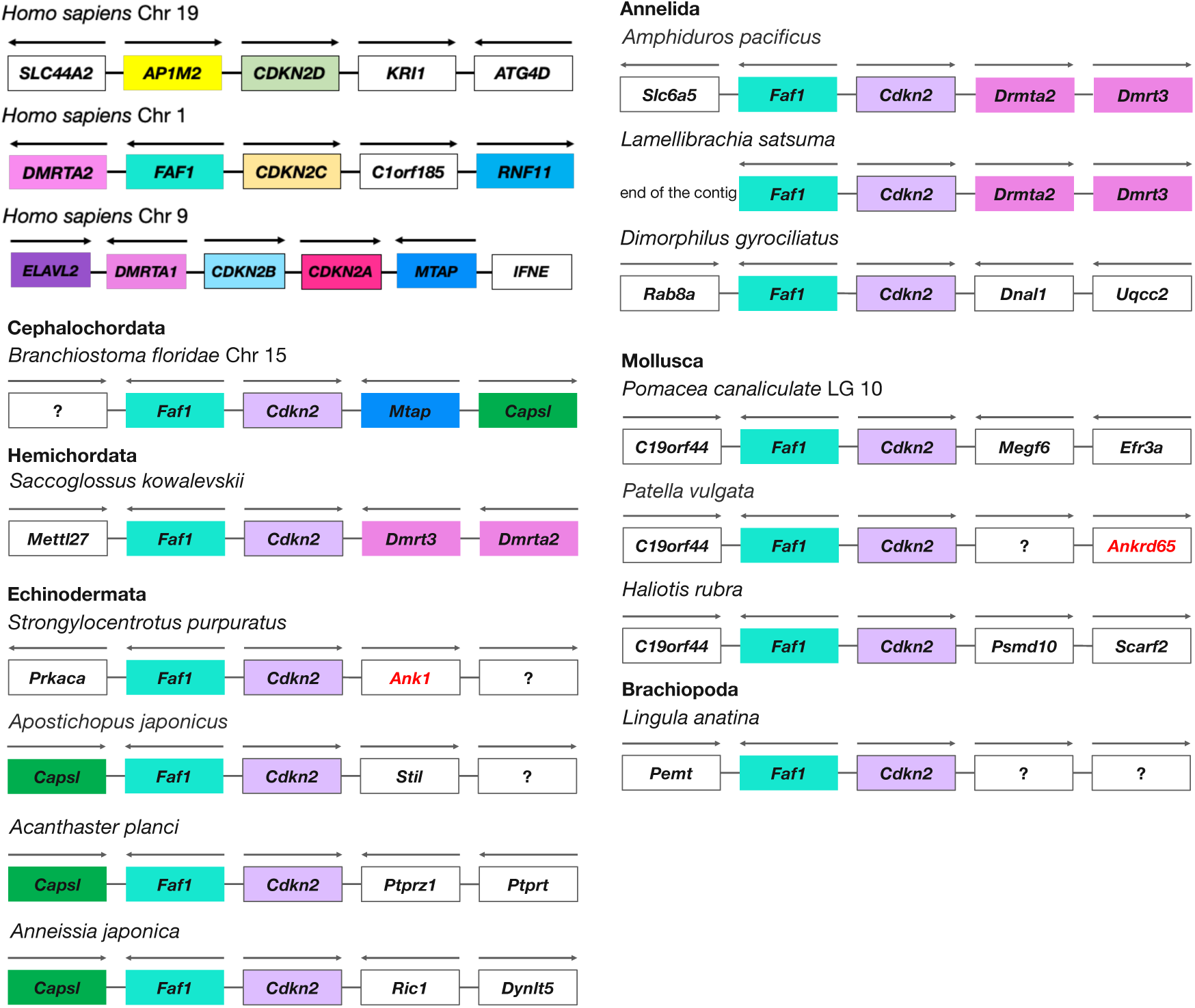
The microsynteny analysis revealed that *Cdkn2* and *Faf1* are tightly linked during evolution. Arrows show the direction of transcription. The chromosome (Chr) number or the linkage group (LG) is shown when available. Microsynteny around human *CDKN2* loci is shown as a reference. Genes encoding ankyrin repeat–containing proteins other than Cdkn2 are in red.

To obtain clues on the origin and loss of *Cdkn2*, we analyzed microsynteny around *Faf1* in the species without *Cdkn2*. We could not detect any evolutionary conservation of linkages comparable to that between *Cdkn2* and *Faf1*. No “weakly positive” sequences mentioned above was not linked with *Faf1*, which is another evidence that they are not *bona fide Cdkn2*. The linkage between *Faf1* and *KN Motif and Ankyrin Repeat Domains 1 (Kank1*) was conserved among some species in Anthozoa of Cnidaria (Figure 4, Table S8).

**Figure 4.**
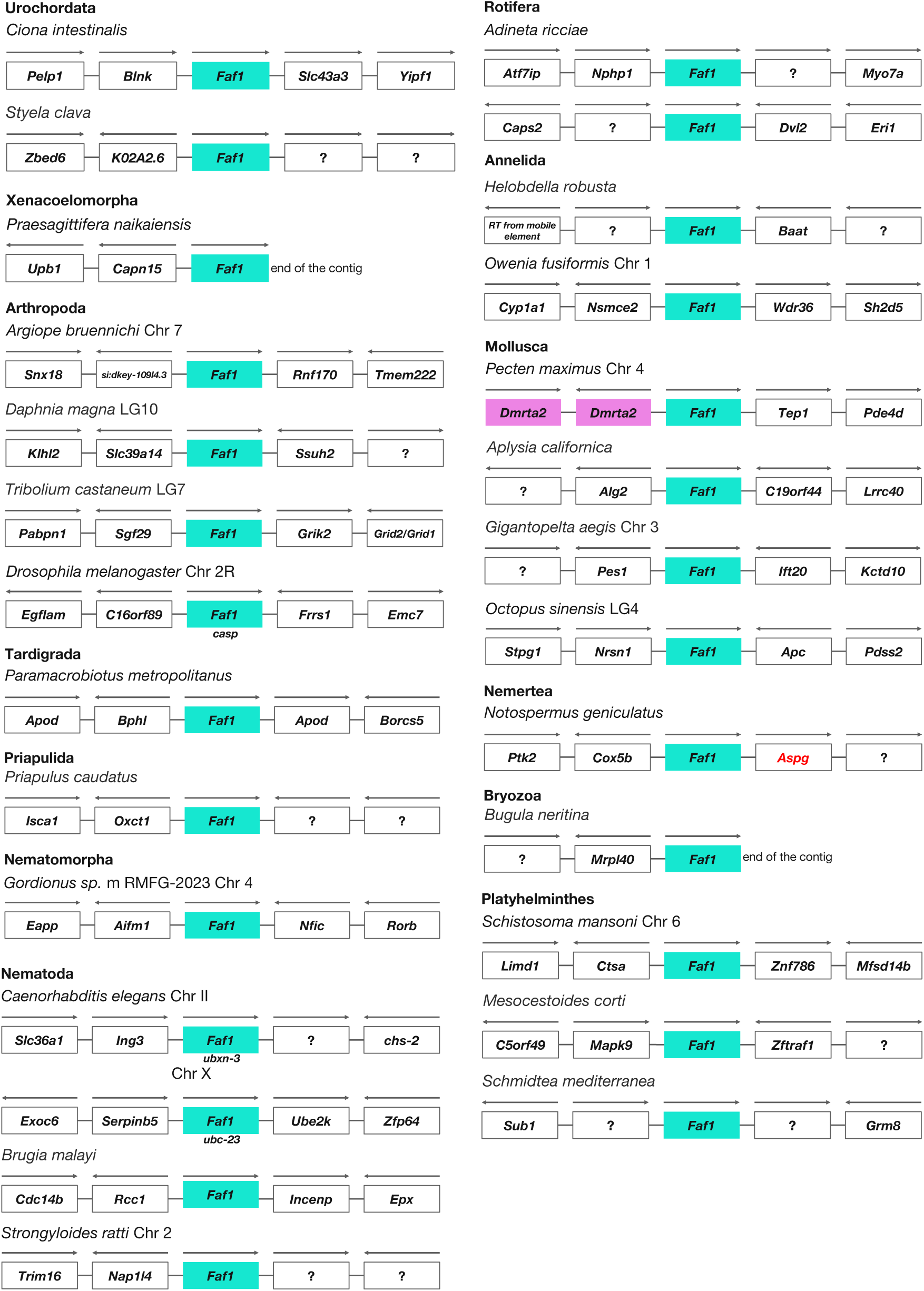
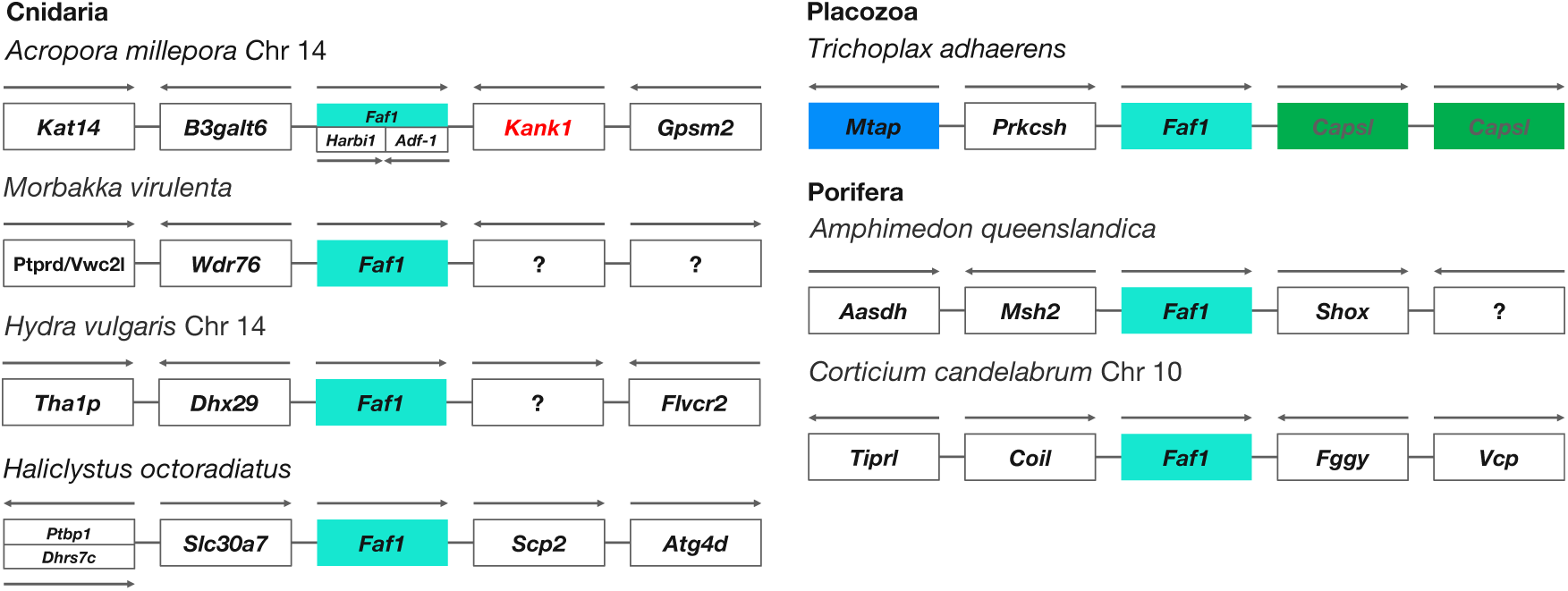
Microsynteny around the *Faf1* locus in representative invertebrates without *Cdkn2*. Arrows show the direction of transcription. The chromosome (Chr) number or the linkage group (LG) is shown when available. A gene encoding an ankyrin repeat– containing protein other than Cdkn2 is in red.

### Presence of Cdk4/6 in invertebrates

Our systematic search for *Cdk4/6* in invertebrate genome databases with gene names and blastp/tblast revealed 1296 candidate sequences from 1265 species among 21 phyla (Table S9). We could not detect Cdk4/6 candidate sequences in Porifera (Cao et al., 2014), Dicyemida and Orthonectida (Tables 1, S9). To confirm the identity of the sequences as Cdk4/6, we performed a phylogenetic analysis. Because of the various lengths of the 1296 sequences, we selected 1172 sequences, from 1152 species among 20 phyla, which contain the protein kinase domain (PKD) without apparent deletions for the analysis to obtain a reliable result. The phylum excluded by this selection was Priapulida. Although the candidate sequence (XP_014677898.1) is annotated as a Cdk6-like and the blastp analysis supported it, we did not include the phylogenetic analysis because a part of the PKD was missing due to gaps of the contigs. The result with the PKD of the 1172 sequences as well as the ones of human representative Cdk members (CDK1, 4, 5, 6, 7, 8, 9, 11B, and 20; Malumbres, 2014) showed that all the Cdk4/6 candidate sequences formed one clade (Figure S3, Table S10). We used the 1172 sequences for further analyses.

### Analysis of the residues corresponding to R24/R31 in Cdk4/6

In vertebrate Cdk4/6/21 from 572 species, the strong positive charge of the residues corresponding to R24/R31 was conserved (99.7%; 1248/1252; Figure S5, Table S11). In invertebrate the charge was conserved in 1048 sequences (89.4%), including 198 sequences (16.9%) in which the R was substituted with K (Tables 2, S9). The strong positive charge was lost in 124 sequences (10.6%); among invertebrate Deuterostomia, two *Ciona* species in Urochordata lacked the positive charge (2/31, 6.5%) (Figures 5, S4a). In contrast, in *Corella inflata* and *Ascidia aspersa*, which belong to the same order (Phlebobranchia) as *Ciona*, the R was substituted with K, thus, the charge was conserved (Shito & Hotta, personal communication). The strong positive charge was lost in 37.7% (43/114) of the sequences investigated in Lophotrochozoa (Table 2, Figure S4b, c), in 7.7% (75/969) in Ecdysozoa (Table 2, Figure S4d, e) and in 6.9% (4/58) in basal invertebrates (Table 2, Figure S4f). The corresponding residue was mostly T in 97.5% (39/40) of Platyhelminthes species (Lophotrochozoa) and in 62.1% (41/66) of the Nematoda species (Ecdysozoa). It is interesting that many species in the evolutionarily distant Platyhelminthes and Nematoda, both of which contain many parasites, had T instead of R24/R31, although the importance is unknown.

**Figure 5.**
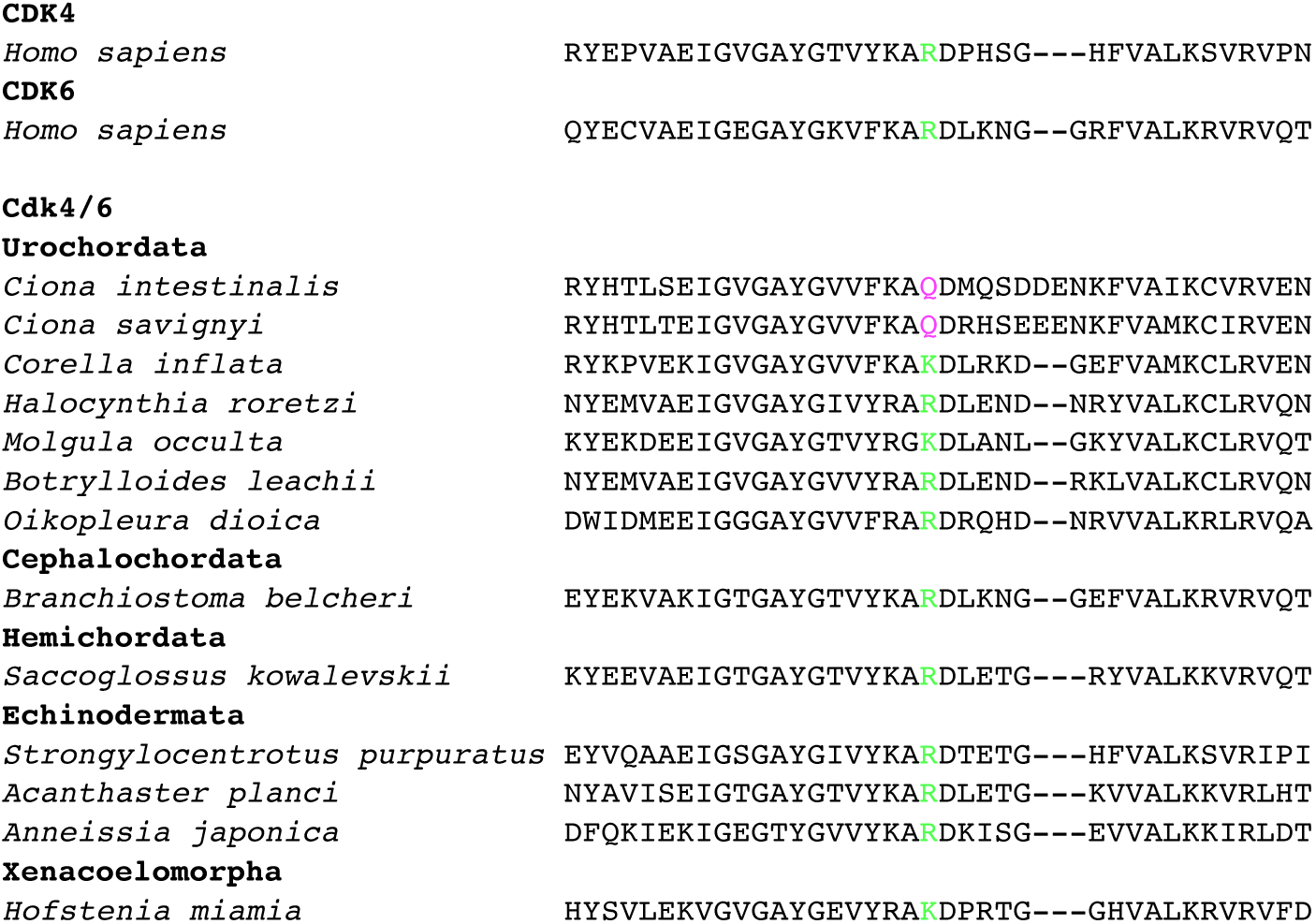
Amino acid alignment around the residue corresponding to Cdk4/6 R24/31 in representative invertebrate Deuterostomia species. The residues corresponding to Cdk4/6 R24/31 are green if positively charged or in magenta if not. Human CDK4 and CDK6 are aligned as references.

**Table 2.**
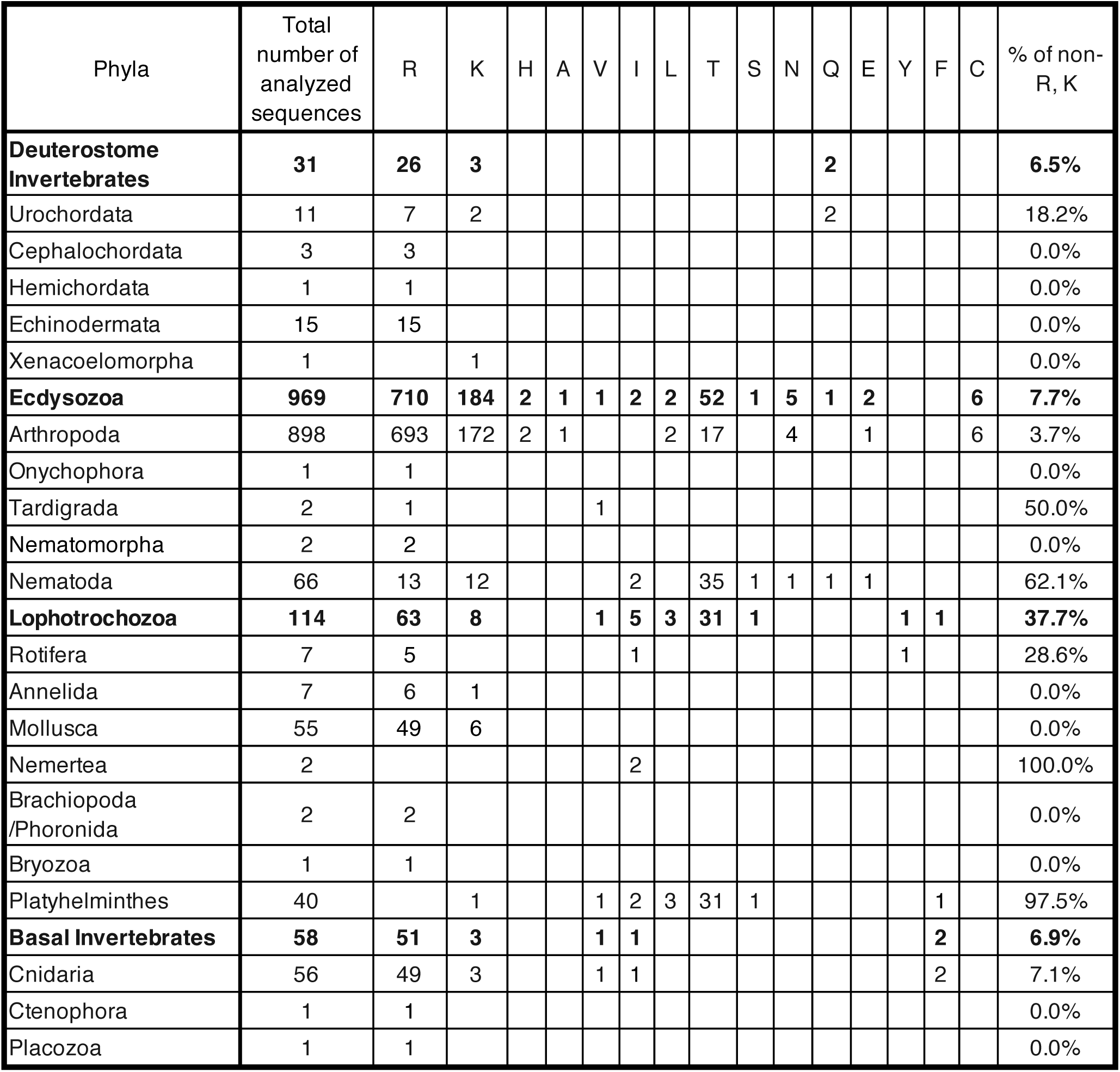
Number of species with each type of substituted amino acids for CDK4/6 R24/R31.

### Relationships between the presence of Cdkn2 and the positive charge of the residue corresponding to R24/R31 in Cdk4/6 among phyla

While species in most of animal phyla have *Cdk4/6*, the extent of the evolutionary conservation of the residue corresponding to R24/R31 greatly differed among phyla. We speculated that the presence of *Cdkn2* might affect the extent, we compared them in each phylum (Figure 1). In all analyzed species in phyla with *Cdkn2* gene(s), the strong positive charge of the residue corresponding to the R24/R31 of Cdk4/6 homologs was conserved, even in the Crinoid *A. japonica* where the corresponding negative charge of D84 in Cdkn2 was lost. In phyla without *Cdkn2*, high diversity was observed. The strong positive charge was lacking in a few species (Cnidaria, 7.1%; Arthropoda, 3.7%; and Urochordata, 18.2%), many species (Nematoda, 62.1%), or almost all species (Platyhelminthes, 97.5%).

Finally, we analyzed the statistical significance of correlation between the presence of *Cdkn2* and the conservation of a strong positive charge corresponding to R24/R31 in Cdk4/6. We analyzed nine phyla (Figure S6) in which at least five species were included in this study (Figure 1). The Spearman’s rank coefficient was 0.816 (*P* = 0.00735), suggesting that the presence of *Cdkn2* and the conservation of the positive charge were strongly correlated with statistical significance.

## Discussion

In this study, we comprehensively investigated the presence of *Cdkn2* and *Cdk4/6* in invertebrate genomes and the evolutionary conservation of the key charged residues; we also investigated the *Cdkn2* synteny. The distribution of *Cdkn2* was uneven among and within invertebrate phyla while most of the phyla contained *Cdk4/6* (Table 1). The lack of *Cdk4/6* in Dicyemida and Orthonectida may be caused by the genome size shrinkage (Lu et al., 2019; Slyusarev et al., 2020).

### Evolution of Cdkn2 gene

On the basis of the data on the *Cdkn2* presence in invertebrates, we propose a scenario of *Cdkn2* evolution (Figure 6a). The ancestral *Cdkn2* gene first originated from an ankyrin repeat–encoding gene during the early evolution of Bilateria. During the diversification of Bilateria, this gene has been lost in some phyla, although to a different extent among Ecdysozoa, Lophotrochozoa, and Deuterostomia.

**Figure 6.**
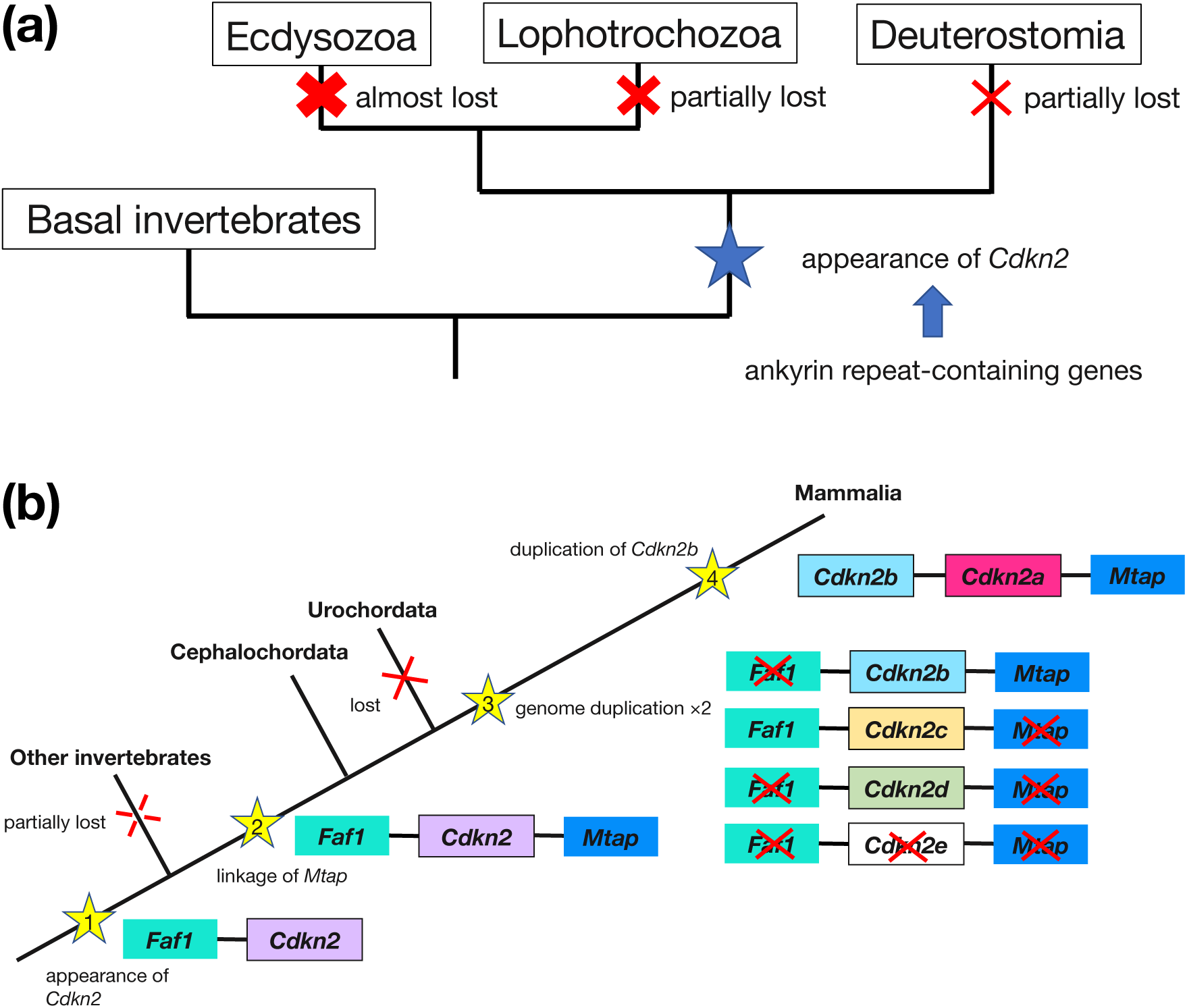
Hypothesis on the evolution of the *Cdkn2* locus. (a) Loss of *Cdkn2* occurred multiple times. (b) Evolution of microsynteny at the *Cdkn2* locus. *Cdkn2e* is a hypothetical transient gene.

The uneven distribution could indicate that the loss of *Cdkn2* might be neutral or non-harmful in some cases, and its presence might be beneficial for survival or reproduction in other cases. Because Cdkn2s regulate the cell cycle and are known as tumor suppressors as well as cell senescence markers in mammals, their beneficial role in invertebrates might be related to cell proliferation, cell differentiation, cell aging, or prevention of cancer. However, it should be noted that even in deuterostome invertebrates, the mechanisms of cell aging seem to differ from those in mammals (reviewed in Amir et al., 2020). Species in the invertebrate phyla without *Cdkn2*, like those in Platyhelminthes, Nemertea, Bryozoa, and Urochordata as well as non-Bilateria, are known for their extreme regenerative ability. However, species in the invertebrate phyla with *Cdkn2*, like those in Phoronida, Annelida, and Echinodermata, also have this ability (reviewed in Bely & Nyberg, 2010). Thus, *Cdkn2* is not another example of genes that are highly linked to extreme regeneration ability, such as the JmjC domain– encoding gene (Cao et al., 2019). It seems that longevity may not be related to the presence of *Cdkn2* in invertebrates. Even if we exclude non-Bilateria, which include species with extreme longevity or theoretical immortality (Brien, 1953; Piraino et al., 1996; Jochum et al., 2012), both a sea urchin, *Mesocentrotus* (*Strongylocentrotus*) *franciscanus*, which has a *Cdkn2* gene (Figure S1), and a clam, *Arctica islandica* (Bivalvia), without it (Table 1), are known for to their longevity, although the nuclear genome of *A. islandica* has not yet been published (Ebert & Southon, 2003; Scho□ne et al., 2005). Thus, we have currently no compelling clear-cut hypothesis to explain the uneven distribution of *Cdkn2*.

### Strong evolutionary conservation of the microsynteny in the Cdkn2–Faf1 locus

Our synteny analysis revealed a strong linkage between *Cdkn2* and *Faf1*. On the basis of these data, we propose a scenario of the evolution of the linkage around the *Cdkn2* locus in invertebrates (Figure 6b). When the ancestral *Cdkn2* gene first appeared, it was linked to *Faf1*. In an ancestral chordate, the *Cdkn2* gene became linked also to *Mtap*. During the double duplication of the genome in an ancestral vertebrate, a *Cdkn2* copy (*Cdkn2c*) remained linked to *Faf1*, and another one (*Cdkn2b*) remained linked to *Mtap*. One of the four *Cdkn2* copies became *Cdkn2d*, and the last one, “*Cdkn2e*,” was lost. Later, the duplication of *Cdkn2b* produced *Cdkn2a*. Human *Faf1*, *Mtap*, and *Cdkn2* are tumor suppressors, and the genetic linkage between *Cdkn2* and *Faf1* can be traced back to the *Cdkn2* origin (Jen et al., 1994; Kamb et al., 1994; Husemann et al., 1999; Christopher et al., 2002; Menges et al., 2009; Gluick et al., 2013). Since *Faf1* is linked with *Kank1* encoding another ankyrin repeat–containing protein in Anthozoa of Cnidaria, *Cdkn2* in Bilateria and *Kank1* in Anthozoa might share the origin in ancient metazoans.

Why has the microsynteny in the *Cdkn2–Faf1* locus been conserved during bilaterian evolution? Microsynteny in a locus with multiple genes may be conserved by *cis*-regulatory constraints between the genes and/or the functional interdependency of the gene products (Imai et al., 2012; Irimia et al., 2012; Simakov & Kawashima, 2017; Cai et al., 2010). Although both *Cdkn2*s and *Faf1* are tumor suppressor genes in mammals, they function in different cellular systems (Cdkn2s in the cell cycle and Faf1 in apoptosis), and their physical interaction has not been reported. Thus, *cis*-regulatory constraints between them might maintain the microsynteny. In this context, it is interesting that the expression of a lncRNA gene, which is on the immediate opposite side of the *C1orf185* gene from *Cdkn2c*, can affect *Faf1* expression in humans, although the linkage between *C1orf185* and the *Cdkn2c*–*Faf1* locus is not evolutionarily conserved even in vertebrates (Wan et al., 2014; Araki, unpublished data). A conservation of microsynteny might also reflect a hidden physical interaction (Moretti et al., 2017).

### Potential interactions between Cdkn2 and Cdk4/6 evolution

Our comparison between the presence of *Cdkn2* and the differential conservation of the strong positive charge at the residue corresponding to R24/R31 in Cdk4/6 among phyla suggests potential interactions between *Cdkn2* and *Cdk4/6* evolution. All species in which we detected both *Cdkn2* and *Cdk4/6* in Annelida, Mollusca, and Echinodermata retain the charge, while all investigated species in Platyhelminthes and Nematoda lack *Cdkn2*, and a high percentage of them also lack the positive charge, implying that the presence of *Cdkn2* might constrain the residue corresponding to R24/R31 as the interface from a substitution or loss when the presence of *Cdkn2* is beneficial, or its loss is harmful to the organism.

On the other hand, most of the species in Cnidaria, Arthropoda, and Urochordata retain the charge but lack *Cdkn2*. The Crinoid *A. japonica* lacks the negative charge in Cdkn2 at the residue corresponding to D84, but retains R24/R31 in Cdk4/6s. In this species, the Cdk4/6 sequence might lose the positive charge in the future. Alternatively, this positive charge in the Cdk4/6 homologs might be conserved to maintain the three-dimensional structure and/or interactions with proteins other than Cdkn2.

### Cell cycle regulation without Cdkn2

What does substitute Cdkn2 in species without it? In mammalian cells, Cdkn1 preferentially inhibits Cdk1/2 but can also inhibit Cdk4/6, although the structure of the Cdk4/6–Cdkn1 complex has not yet been solved by X-ray crystallography (reviewed in Wood & Endicott, 2018, and Muühleder et al., 2021). It is easy to speculate that Cdkn1 substitutes for Cdkn2 in the species without *Cdkn2*, as reported for some invertebrates including *Ciona*, *D. melanogaster* and *C. elegans* (Kobayashi et al., 2022; reviewed in Kipreos & van den Heuvel, 2019). However, caution is needed in extrapolating this mechanism to other urochordates or species lacking *Cdkn2* in other phyla, because *Ciona* is the only urochordate investigated that lacks the positive charge at the residue corresponding to R24/R31 in Cdk4/6. Since the extent of the evolutionary conservation of the strong positive charge greatly differs among phyla lacking *Cdkn2*, the conserved positive charge in Cdk4/6 of Cnidaria, Arthropoda, and Urochordata might not be involved in the interaction with Cdkn1 and might be necessary for the interactions with other proteins.

### Conclusion

We believe that the results in this study not only widen our knowledge on the mechanisms of cell cycle regulation and their evolution in metazoans, but also provide fundamentals to design a cell immortalization protocol in invertebrates for basic sciences as well as drug discovery and clinical research.

## Experimental procedures

### Taxonomy

We used the taxonomy of invertebrates by Telford et al. (2015) for phyla, and the one in the NCBI Taxonomy database for subclassification. The evolutionary position of Xenacoelomorpha is controversial (Cannon et al., 2016; Philippe et al., 2019); we tentatively categorized it as a diverged deuterostome.

### Database searches

To analyze the presence of *Cdkn2* and *Cdk4/6* genes and to obtain the corresponding amino acid sequences, we used keyword and blastp searches in the NCBI Protein database (https://www.ncbi.nlm.nih.gov/protein/), Ensembl Metazoa (http://metazoa.ensembl.org/index.html), Ensembl Rapid Release (https://rapid.ensembl.org/index.html), Ghost Database: *Ciona intestinalis* genomic and cDNA resources (http://ghost.zool.kyoto-u.ac.jp/SearchGenomekh.html), Aniseed - Ascidian Network for In Situ Expression and Embryological Data (https://aniseed.fr/), PlanMine (https://planmine.mpinat.mpg.de/planmine/begin.do), and OIST Marine Genomics Unit Genome Browser (https://marinegenomics.oist.jp/gallery) (Sayers et al., 2024; Yates et al., 2022; Satou et al., 2005; Dardaillon et al., 2020; Rozanski et al., 2019; Shinzato et al., 2020, 2021; Yoshioka et al., 2022; Khalturin et al., 2019; Luo et al., 2018; Takeuchi et al., 2022; Arimoto et al., 2019; Simakov et al., 2015). We also performed tblastn searches in the NCBI Whole-genome shotgun contigs database. To confirm the identity of the sequences obtained, we used blastp searches against vertebrate sequences. We also used vertebrate *Cdkn2* and *Cdk4/6/21* sequences as references and noticed that some vertebrate genes were misannotated in the NCBI databases. For example, a platypus Cdkn2 (XP_028910528.1) is annotated as “cyclin-dependent kinase 4 inhibitor B-like”, but the microsynteny suggested that it is Cdkn2a; thus, we treated it as Cdkn2a, not Cdkn2b. As the paralogs Cdkn2a–Cdkn2d are specific to vertebrates, such misannotations do not affect this study.

In this analysis, we included 61 Cdkn2 sequences from 61 invertebrate species, and 1172 Cdk4/6 sequences from 1152 invertebrate species. As references, we used 1853 Cdkn2 sequences and 1252 Cdk4/6/21 sequences from 572 vertebrates. The database IDs of the analyzed sequences are listed in Tables S1 (invertebrate Cdkn2), S6 (vertebrate Cdkn2), S9 (invertebrate Cdk4/6), S11 (vertebrate Cdk4/6/21).

### Multiple alignment of protein sequences

We used NCBI COBALT (Constraint-based Multiple Alignment Tool) to align protein sequences (Papadopoulos & Agarwala, 2007). Questionable alignments were corrected manually. We used WebLogo (https://weblogo.berkeley.edu/logo.cgi) to create a consensus sequence logo, and MOTIF Search (https://www.genome.jp/tools/motif/) to identify protein motives.

### Phylogenetic tree construction

The phylogenetic trees (Figures S1d and S4) were constructed with NGPhylogeny.fr (https://ngphylogeny.fr/) (Edgar, 2004; Capella-Gutiérrez et al., 2009; Junier & Zdobnov, 2010; Katoh & Standley, 2013; Goloboff & Catalano, 2016; Lemoine et al., 2019). To construct the *Cdkn2* phylogenetic tree, we used Cdkn2 sequences without apparent truncation at the N– or C– terminal (47 of a total 61 invertebrate sequences [77.0%] and 4 human sequences; the 47 sequences were from all the phyla that we judged *Cdkn2*-positive), sequences for 73 human ankyrin repeat–containing proteins and human CDKN1A sequence, and the following programs with default settings: MAFFT, trimAI, TNT, Newick Display. To construct the Cdk4/6 phylogenetic tree, we used the PKDs (PF00069) without apparent deletions (1172 out of total 1296 (90.4%) invertebrate sequences and 2 human sequences) and 7 human sequences of other representative Cdk family members, and the following programs with default settings: MUSCLE, TNT, Newick Display.

### Synteny analysis

We analyzed protein–coding genes only in the synteny analysis. We used the NCBI gene database for the sequences with an NCBI gene ID (Brown et al., 2015). For long genomic sequences without ID, we analyzed the annotation, if available, or used a blastp search against the flanking sequences. If the surrounding genes have been annotated, we used blastp against vertebrates to confirm the identities. We checked GeneCards (https://www.genecards.org) for the abbreviated gene names.

### Statistical analysis

We used the statistical software EZR to determine Spearman’s rank correlation coefficient and create the scatter plot (Kanda, 2013). To determine the Spearman’s rank correlation coefficient, we used the percentage of the number of species with both Cdkn2 and Cdk4/6 to that with Cdk4/6, and the percentage of the number of species with the conserved strong positive charge at Cdk4/6 R24/31.

## Acknowledgements

We thank Drs. Yutaka Satou (Kyoto Univ) and Takeshi Kawashima (Inuyama Bio) for critically reading the manuscript, and Takuya Minokawa (Tohoku Univ), Kazuya Kobayashi (Hirosaki Univ), Hiroshi Wada (Tsukuba Univ), and Koji Tamura (Tohoku Univ) for helpful discussion. We also thank Drs. Takumi Shito and Kohji Hotta (Keio Univ) for sharing their unpublished data. This research was financially supported by Iwate University.

## Supplementary materials

**Table S1** The list of the analyzed invertebrate *Cdkn2*s.

**Table S2** Results of blast searches for invertebrate *Cdkn2*. (a) blastp on refseq_protein, (b) tblastn on wgs database.

**Table S3** The list of the invertebrate Cdkn2 candidates and the human homologs of other ankyrin repeat–containing sequences as well as human CDKN1A as an outgroup for the phylogenetic analysis in FASTA format.

**Table S4** The list of invertebrate genes possibly misannotated as *Cdkn2*.

**Table S5** The list of the identified *Cdkn1*s (a) in invertebrates other than Arthropoda and (b) in Arthropoda.

**Table S6** The list of the analyzed vertebrate *Cdkn2*s.

**Table S7** Microsynteny around the *Cdkn2* locus in invertebrates.

**Table S8** Microsynteny around the *Faf1* locus in (a) invertebrates other than Arthropoda and (b) Arthropoda.

**Table S9** The list of the analyzed invertebrate *Cdk4/6*s. (a) deuterostome invertebrates, (b) Platyhelminthes, (c) other Lophotrochozoa, (d) Nematoda, (e) other Ecdysozoa, and (f) basal invertebrates.

**Table S10** The list of the analyzed vertebrate *Cdk4/6*s.

**Table S11** The list of the invertebrate Cdk4/6 candidate sequences and human homolog sequences of other representative Cdk members in FASTA format.

**Figure S1** Amino acid alignment of invertebrate Cdkn2s. (a) N-terminal half of Cdkn2. (b) C-terminal half. The residue corresponding to D84 is in green if negatively charged, or in magenta if not. (c) Consensus residues in invertebrate Cdkn2. **`a`** and **`b`** represent amino acids of α-helices and β-turns in *Homo sapiens* CDKN2s, respectively (Baumgartner et al., 1998). (d) Phylogenetic analysis of invertebrate Cdkn2 candidate sequences.

**Figure S2** Cdkn2 amino acid alignment around the residue corresponding to D84 in vertebrates. Cdkn2a in (a) mammals and (b) non-mammalian vertebrates; Cdkn2b in (c) mammals, (d) non-mammalian/non-fish vertebrates, and (e) fish; Cdkn2c in (f) mammals, (g) non-mammalian/non-fish vertebrates, and (h) fish; Cdkn2d in (i) mammals and (j) non-mammalian vertebrates. The residue corresponding to Cdkn2 D84 is in green if negatively charged or in magenta, if not.

**Figure S3** Phylogenetic analysis of invertebrate Cdk4/6 candidate sequences and human representative Cdk members (CDK1, 4, 5, 6, 7, 8, 9, 11B, and 20).

**Figure S4** Cdk4/6 amino acid alignment around the residue corresponding to R24/R31 in invertebrates. (a) deuterostome invertebrates, (b) Nematoda, (c) other Ecdysozoa, (d) Platyhelminthes, (e) other Lophotrochozoa, and (f) basal invertebrates. The residue corresponding to R24/31 in Cdk4/6 is in green if positively charged and in magenta, if not.

**Figure S5** Cdk4/6/21 amino acid alignment around the residue corresponding to R24/R31 in vertebrates. Cdk4 in (a) mammals, (b) non-mammalian/non-fish vertebrates, and (c) fish; Cdk6 in (d) mammals, (e) non-mammalian/non-fish vertebrates, and (f) fish; Cdk21 in (g) fish. The residue corresponding to Cdk4/6 R24/31 is in green if the strong positive charge is conserved and in magenta, if not.

**Figure S6** Determination of the Spearman’s rank correlation coefficient. (a) The data used and (b) the scatter plot.

## Footnotes

* The numbering of the amino acids is based on human proteins. D84 in human CDKN2A corresponds to D86 in CDKN2B, D76 in CDKN2C, and D80 in CDKN2D. For simplicity, we use D84 for D84/D86/D76/D80 in this paper.

## Notes

### Competing Interest Statement

The authors have declared no competing interest.

### Summary of Updates

Added the results of phylogenetic analyses for Cdkn2 and Cdk4/6 candidates; Thoroughly improved English in the manuscript; Figures, Tables and Supplemental files updated.

