## Supplementary Tables S1-S11 and Figures S1-S6 for "Evolution of the *Cdk4/6*–*Cdkn2* system in invertebrates": Figure S1.pdf

```
Branchiostoma lanceolatum          MDGGGDTYLKGHAGDFRRRRSAWL------------------REGRATNKRRFLRDRLSGLP-----G-KDWCQLVPTTDDIQVEIIALLIELVFW---------PESELCGYNKVGSGDES-------SDGALCGYCE 99
Patiria miniata  (X1)              MLLP-----------------------------RKFCR----R-------------RTHGGW-R- 17
Acanthaster planci (X1)            MDISHVIMPFG-----------------------SPARSDFTRARDRR-------------DSSLSWWE- 33
Acanthaster planci (X2)            MDISHVIMPFG-----------------------SPARSDFTRARDRR-------------DSSLSWWE- 33
Dimorphilus gyrociliatus           MSSRSFRI---PREGAFHPVQRRIPEATPLCPLQS 32
Marisa cornuarietis                MIGGSAHGCVSCPVVVKPWEPRSTPLTQRSSHFQRSSSIHPLSDEDVRSKQDRSSAFRRIPFQNVTDDSKPKIRRRSNGEKDPKAGTDIGQQDFVSLIYRGHDTSQQQDETKMTV-LHPFEEINATIDDATIRTKHDKRVLHHHGLHHSSFAVHSSLDAGSTREDNRE-IMQMPLSHYVS 178
Pomacea canaliculata               MI-GSAHGCVSCPVEVKPWEPRSTPLTQRSSHFQRSASIHPLSDEDESSKQDQSSAFQRIPIQSVTDDSKLKIRRQSNGEKDPKAGTDIGQQDFVSPIYHGPDTSQQQGEAQMTVLMKPVEEIRSTINDATIRTKHYKSVLHHHGLHHSSFAVHSSLDEGSTREDLRE-IIKIPLSHYVT 178
Pomacea maculata                   MI-GSAHGCVSCPVEVKPWEPRSTPLTQRSSHFQRSASIHPLSDEDERSKQDQSSAFQRIPIQSVTDDSKLKIRRRSNGEKDPKAGTDIGQQDFVSPIYHGPDTSRQQGEAQMTVLMKPVEEIKATINDATIRTKHYKSVLHHHGLKHSSFAVHSSLDEGSTREDLRE-IIKIPLSHYVT 178
Alviniconcha marisindica           MQDPF---QCLTVDA-----------------RREAAVQSHQSA-AV-LNAH---HNP--CGART 38
Conus tribblei                     MKGPQK-KKCAISDGPAGDSLLKKQLGK----KGEPAVPNSPPT-SFDTTPK---VLPPGTSASD 56


(Cont'd)
                                   aaaaaaaaaa  aaaaaaaaa    a        bb aaaa
CDKN2A
Homo sapiens                       MEPAAGSSMEPSADWLATAARGRVEEVRALLEA----GALPNAPNSYGRRPIQ 50
CDKN2B
Homo sapiens                       MR----EENKGMPSGGGSDEGLASAAARGLVEKVRQLLEA----GADPNGVNRFGRRAIQ 52
                                   aaaaaaaaaa  aaaaaaaaa              bb  aaa
CDKN2C
Homo sapiens                       MAEPWGNELASAAARGDLEQLTSLLQN----NVNVNAQNGFGRTALQ 43
                                   aaaaaaaaaa  aaaaaaaaa              bb  aaa
CDKN2D
Homo sapiens                       MLLEEVRAGDRLSGAAARGDVQEVRRLLHR---ELVHPDALNRFGKTALQ 47
Cdkn2
Cephalochordate
Branchiostoma belcheri             MAHNGAAGEAGTSSPEGAAGGQAV---QE-------QAVGFHAIN--DGNDLTTAAAGDLVEVQRLLEV----GVDVNAQNQGGRTAIQ 74
Branchiostoma floridae             MAHNGAAREDGTSFPEGAAGGQAA---QE-------QAAGFHAINVHDGNDLTTAAARGDLEEVRRLLAA----GVDVNARNQHGMTAIQ 76
Branchiostoma lanceolatum          VAELQVKLGAIICCIDCV---RSPA--AGLAIQVDASH--------------RAHSKTP--GVGERREDPVRGTGSPRRGMAHNGAAGEAGTSSPEGAAGGQAV---QA-------PAAGFHAIN--DGNDLTTAAARGDLDEVRRLLEV----GVDVNAQNQHGMTAIQ 232
Asymmetron lucayanum               MAHNGA----EASSPGGAAGGQAG---EE-------QAAGFQPIS--DGNDLTTAAAKGDMDEVRRLLQA----GVEVNARNQLGQTAIQ 70
Hemichordate
Ptychodera flava                   MLLEKGVSIELTS---FF-------VLPAMNNDGGHVQMSLAEAAARGDDGTVRDLLAQ----GCDVHERNRYNRTPIQ 65
Saccoglossus kowalevskii           MVAGKIR---EFKDVCL--ITVMANDVQEESELTLIEAAAKGLEDQVRRLLEL----GSDVHRRNIFGRTALQ 64
Echinoderm
Strongylocentrotus purpuratus (X1~2)   MAEQETIDAMTSAAAKADDAALVKILDG--APPGIVNRPNIYDRTAIQ 46
Mesocentrotus franciscanus         MADQETIDAMTTAAVNADDATLERILDG--APPGTVNRPNTDGYTAIQ 46
Hemicentrotus pulcherrimus         MADQETINEMTSAAANADDATLVRILDG--APPGTVNRPNKYNRTAIQ 46
Paracentrotus lividus              MADDQETINAMTTAAAKADDTKLLEILSQ--SPPGTVNRTNKYGRTAIQ 47
Heliocidaris erythrogramma         MADEETIKAMTQAAATADDAKLADKLHQ--TPPGTVNRPNRYGRTAIQ 46
Heliocidaris tuberculata           MADEETINAMTQAAATADDATLADILDR--APPGTVNRPNRYGRTAIQ 46
Lytechinus pictus                  MAAA---APVDVLDQATINEMTQAAANGNHGRIEEILAQAQHPRDAVNKVNKYGRTAIQ 56
Lytechinus variegatus (X1~2)       MAAP---DPVDVLDQATINEMTQAAANGNHGRIEEILAQANPRDAVNEVNNYGRTAIQ 56
Patiria miniata  (X1)              ------------TCLI-------------LLSA--F---------WRSFTPLRTSSPSP-------------------IGMI-------STESRRLELPSVFESYFIDVAIMAANANHQQ--PAAVNPQDEFTRAAANGDMNRLRELSGQ---VGVDVDGRNKYGRTAIQ 120
Patiria miniata  (X2)              M-------------NYSG--A---------FSVFGCLRNLAVNG-------------------NGVR-------NN----QKSSSTEWERELDVAIMAANANHQQ--PAAVNPQDEFTRAAANGDMNRLRELSGQ---VGVDVDGRNKYGRTAIQ 96
Patiria miniata  (X3)              MWNLAVNG-------------------NGVR-------NN----QKSSSTEWERELDVAIMAANANHQQ--PAAVNPQDEFTRAAANGDMNRLRELSGQ---VGVDVDGRNKYGRTAIQ 84
Patiria miniata  (X4)              MAANANHQQ--PAAVNPQDEFTRAAANGDMNRLRELSGQ---VGVDVDGRNKYGRTAIQ 54
Acanthaster planci (X1)            ------------SEEV-------------DLAQ--VRPHSDPFAKLSEVICKSTGQSRI-------------------PGMV-------SPVSRPLQLQIVFKGYLIDVAMIAANAPRQL--S-AATLQDEFTRAAANGDMEQLLELKGI---ANVDINGQNKYGRSAIQ 144
Acanthaster planci (X2)            ------------SEEV-------------D-----------------------------------------------------------------------------VAMIAANAPRQL--S-AATLQDEFTRAAANGDMEQLLELKGI---ANVDINGQNKYGRSAIQ 94
Acanthaster planci (X3)            MIAANAPRQL--S-AATLQDEFTRAAANGDMEQLLELKGI---ANVDINGQNKYGRSAIQ 54
Patiriella regularis               MAANANNQQ--RATASPQDDFTQAAARGDMGRLRQLSGQ---ANVEVNGRNRYGRTAIQ 54
Asterias rubens  (X1~2)            MAR--V----SLSLTFSVFECHHKWQG---------------------KHVT-------ADDKSPLTLDLLLADSDVSVIMANNNN-NQH--L-GTDVQNDFTKAAANGNLQRLQELSDH---QDVNINGRNRYGRNAIQ 99
Asterias rubens  (X3)              MANNNN-NQH--L-GTDVQNDFTKAAANGNLQRLQELSDH---QDVNINGRNRYGRNAIQ 53
Pisaster brevispinus               MADNI-NQH--L-GTDAQNEFTRAAANGNLQRLQELSGH---QNVSINGRNRYGRTAIQ 52
Pisaster ochraceus                 MADNT-NQH--L-GTDAQNEFTRAAANGNLQRLQELSGH---QDVSINGRNKYGRTAIQ 52
Plazaster borealis                 MADNA-NQQ--V-GTDAQNQFTQAAANGDLQRLHELSEH---QDVNINGHNQYGRTAIQ 52
Ophioderma brevispina              GAEHEECPRDGIARVNDLTQAAARGDIDQVRQLLSK----GVDPNALNKYGRTALQ 52
Ophionereis fasciata               MTESDRMTDLMTEAAAKGHVERVRELLNQ----CVHPDSTNKYGRTALQ 45
Anneissia japonica                 MMSKQHPNQLTVASTRGNLNEVRSLLEQ----GFPVDSPNKYQHTALQ 44
Nesometra (Dorometra) sesokonis    MSEDQNPLTIAASSGNLDVVRSLLEK----GFPVDSTNEFQHTALQ 42
Onychophoran
Epiperipatus broadwayi             MENPHNGGDLVEELTTAASQGNYDKVNQLISK----GVNVNATNQFGRTALQ 48
Euperipatoides rowelli             MEEVPEQRELVESLSRAAADGVHEEVIRLIEM----GVNVNETNRFGRTALQ 48
Annelida
Lamellibrachia luymesi             MDNGGFG---NFAD-----VNSLSTANDKPRVDDLTTAAARGDFPEVTRLIRD----GVGVNDVNIYGRTALQ 61
Lamellibrachia satsuma             MDNGGFG---KFAD-----VNSLSTANDKPRVDDLTTAAARGDFPEVTRLIRD----GVGVNDVNIYGRTALQ 61
Ridgeia piscesae                   MDNGGIG---NLAD-----ANSLSTASDKPRVDDLTTAASRGDFPEVTRLIEA----GVGVNEVNVYGRTALQ 61
Paraescarpia echinospica           MDNGGIG---NLAD-----ANSLSTANDEPGVDDLTTAAARGDFPEVTRLIEA----GVGVNEVNIYGRTALQ 61
Dimorphilus gyrociliatus           SE----------MGVKHVPIRVAEGAEVPVLSDTVFRPVSNE------RLCFSPSKLRQAHARRELRYTPVEGRSSSSPISHHNS--------TPSGRPISQKR---AW---LE--HVSPVCGTPELPSDELSWAAACGDLETVIKLIKE----GRNVNTPNSFGRAPLQ 166
Sipunculus nudus                   MGLE------QANRLN-STAT---QVKMAG---SDEPDEVVIITPADHLSTAAARGDIKEVRKLLEA----GVDVNERNSFGRTALQ 70
Mollusca
Marisa cornuarietis                VKGDDHQPSE-LQCMKLKPIDRQHE--PDILAQTDFKQRS--------ELHFHQHTAHP--KSSDQESVSYETSHSEFQRIAQISYQQN-YVSSSRQLASSSAS---QFSNSA---NSWSDCEAAPPDVDALTTAAAGDIAMVKTLLMN----GVDVNTPNSFGRTAVQ 324
Pomacea canaliculata               VKGDDQQSSE-LSCMELKPMDRQHE--HDILAQTDFKQRN--------ELHFHQHAAHP--ESSDQESVSDEISHSEFQHTAQISYPPN-SVSS---------------SNLT---NSWSACEAALPDVDALTTAAAGDIATVKTLLMK----GVDVNTPNRFGRTAVQ 312
Pomacea maculata                   VKGDDQQPSE-LSCMELKPMDRQHE--NDILAQTDFKQRN--------ELHFHQHAAHP--ESSDQESVSDEISHSEFQHTAQISYQPN-SVSS---------------SNLT---NSWSACEAALPDVDALTTAAAGDIATVKTLLMK----GVDVNTPNRFGRTAVQ 312
Alviniconcha marisindica           VA-----AAVPISCEPFESGLEAKQ--DTVVAQ---------------TTQF--SATSPTEPS--------------------------WEAPTPDSLDS---S---NW-------SLSFGQTAVAPGVDALTTAAAGDMMKVKALLAD----GVDVNAVNTFGRTAIQ 141
Babylonia areolata                 MNS---S---IL-------SV--GQSAAAPEADALTTAAAGDLKKVRALLAE----GVSVNAINTFGRTAVQ 54
Conus betulinus                    ILRRKK--QTAVPS---------------RLCK--PPRHPVTPC--------------------------RDAPTPQSLDS---S---VW-------SL--GQSAVATEVDALTTAAAGDLKKVRSLLSE----DVNVNAVNTFGRTAIQ 87
Conus consors                      MQ--TTTSPSHLC--------------------------RDAPTPQSLDS---S---VW-------SL--GQSAVATEVDALTTAAAGDLKKVRSLLSE----GMNVNAVNTFGRTAIQ 73
Conus tribblei                     SA-----DAVRQQCNNFSPFNFEKK--TDSCPQ---------------PLVQ--TTTSPSHPC--------------------------RDAPTPQSLDS---S---VW-------SL--GQSAVATEVDALTTAAAGDLKKVRSLLSE----GVNVNAVNTFGRTAIQ 157
Conus ventricosus                  DNGCKNFSSLNFEKK--ADSCPQ---------------PFVQ--TTTSPSHPC--------------------------REAPTPQSLDS---S---VW-------SL--GQSAVATEVDALTTAAAGDLKKVRSLLSE----DVNVNAVNTFGRTAIQ 96
Patella depressa                   ME----RMKDQDRRRRNSVDELTSAAAVGDLDQVKYLLSV----GVGVNGRNSYGRTAIQ 52
Patella pellucida                  ME----RMKDKDRRRRNNVDELTSAAAVGDLDQVKYLLSV----GVDVNGKNSYGRTAIQ 52
Patella vulgata                    ME----RMKDKDRRRRNNVDELTSAAAVGDIDQVKYLLSL----GVDVNGRNSYGRTAIQ 52
Haliotis cracherodii               M----AREGEIRCGEFTVDDLTTAANGDLAKVRDILQT----GVDVNGRNSFGRTAVQ 51
Haliotis laevigata                 M----AKEGEVRCGEFTVDDLTTAANGDLAKVRDILRT----GVDVNGRNSFGRTAIQ 51
Haliotis rubra                     M----AKEGEVRCGEFTVDDLTTAANGDLAKVRDILRT----GVDVNGRNSFGRTAIQ 51
Haliotis rufescens                 M----AREGEIRCGEFTVDDLTTAANGDLTKVRDILQT----GVDVNGRNSFGRTAVQ 51
Gibbula magus                      M----AKEGETHCHGFTVDDLTTAAASGDLDRVREVLRT----GVDVNGRNRFGRTAIQ 51
Phorcus lineatus                   M----AKEGETHCQEFTVDDLTTAAASGDLDRVREVLRT----GVDVNGRNRFGRTAIQ 51
Steromphala cineraria              MA----MVEGEAHFQDYTVDDLTTAAASGDLDRVREVLQT----GVDVNGRNRFGRTAIQ 52
Nodopelta heminoda                 AAANGDLENVQVMLLA----GVDVNGRNSFGRTAIQ 32
Nautilus pompilius                 MDPEAGETNCHHISLQYSADDLTNAAAVGNFELVQEIIAS----GLPVNSRNQYGRTALQ 56
Brachiopoda/Phoronida
Lingula anatina                    MHLYYIMFTMVDGVLDHKEMTEEHDASLEEFFNAAVNGNAHRLQQLLDE---HRINATNDWGRTAIQ 66
Phoronis australis                 MQNSSGSRMDELCDACSQGNVERMSQLFDL----DLDVNGTNRYGRTPLQ 46
Phoronis pallida                   MDDLCNACAQGKTAEVLRLLDS----GLPVNGKNSYGRTPIQ 38
```

**Figure S1a**

```
CDKN2A                                      a     aaaaaaaaaa                    bb  aaaaaaaa  aaaaaaaaaa          bb  aaaaaaaa     aaaaaaaaaaa
Homo sapiens                            51  VMMMGSARVAELLLLHGAEPNCADPA----------TLTRPVHDAAREGFLDTLVVLHRAGARLDVRD-AWGRLPVDLAEELGH---RDVARYLRAAAG--GTRG----SNHARIDAAEG--P--SDIPD* 156
CDKN2B
Homo sapiens                            53  VMMMGSARVAELLLLHGAEPNCADPA----------TLTRPVHDAAREGFLDTLVVLHRAGARLDVRD-AWGRLPVDLAEERGH---RDVAGYLRTATG--D* 138
CDKN2C                                      a     aaaaaaaaa          b           b  aaaaaaaa  aaaaaaaaa           bb  aaaaaaa      aaaaaaaaaa          bb  aaaaaaa       aaaaaaaaa
Homo sapiens                            44  VMKLGNPEIARRLLLRGANPDLKDRT-----------GFAVIHDAARAGFLDTLQTLLEFQADVNIED-NEGNLPLHLAAKEGH---LRVVEFLVKHTASNVGHRNHKGDTACDLARLYG--R--NEVVSLMQANGA-GGATNLQ* 168
CDKN2D                                      a     aaaaaaaaa          b           b  aaaaaaa   aaaaaaaaa           bb  aaaaaaaa     aaaaaaaaa           bb  aaaaaaa       aaaaaaaaa
Homo sapiens                            47  VMMFGSTAIALELLKQGASPNVQDTS-----------GTSPVHDAARTGFLDTLKVLVEHGADVNVPD-GTGALPIHLAVQEGH---TAVVSFLAAESD--LHRRDARGLTPLELALQRG--A--QDLVDILQGHMVAPL* 165

Cdkn2
Cephalochordate
Branchiostoma belcheri                  75  VMQFGHDLIVKELLENGADPNVRDPGR---------KGQYPVHDAARGGWDEALKLLFDYHAELDVKD-DEDMTPAHFAAKKGH---VRILEFLRAKSA-DLWAKNRQGQTPRDLARIHG--R--TDVTAWFRGSFPEQHTEQQGQ* 202
Branchiostoma floridae                  77  VMEFGHHIIAKELLENGADPNVRDVE----------TGLSPVHDAARDGWEDTLKLLFEYHADLDVKD-LQDMTPAHLAAEKGH---VRILEFLRAKST-DLWAKNRRGQTPRDLARIHG--K--TDVTAWFRGSFPEQHAEQQGQ* 203
Branchiostoma lanceolatum               233 VMKLGHHFIAKELLENGADPNVRDEE----------TGLSPVHDAARDGWEESLKLLFEYHAILDVKD-KEDMTPAHLAATKGH---VRILEFLRAKSA-DLWAKNRRGQTPRDLARIHG--K--TDVTAWFRGSFPEQHAEQQGQ* 359
Hemichordate
Ptychodera flava                        66  AMNFSRVDIARLLLQRGADPNEGDSV-----------RRAPIHDAAQAGYVDTLIVMAEYGANLELRD-NFGNIPAHVTAMEGN---EQ 139
Saccoglossus kowalevskii                65  AMNFAHPEIARMLLRRGADPNVADST-----------GRAPIHDAAEGGYVNTVQVFVEYGGDLTLRD-NWGNTPAHIAADSDN---LQALELLVKGSN--IYNRNNRNQTVVDLAISKP--R--V--SQFISDYAETPEPRSLKHL----CRLAIR--------KSYDVRVNQVN-VSCRSL---PASLIDYVMMG* 222
Echinoderm
Strongylocentrotus purpuratus (X1)      47  VMNSDSPSTARILIDHGADLDRPDPS----------VQRTILHDIAERGCIETLNLLLEHGANATARD-NMGNTPAHLAAEKGGSYRIQILDRLSQNCT--LTEVNAAGKTPLDLANEGN--H--REVVEWLQNFYTRVFPL--QFL----VRRVI-------------TM--RVR-VINNLHNLLPSILIDYLHGRRPK* 207
Strongylocentrotus purpuratus (X2)      47  VMNSDSPSTARILIDHGADLDRPDPS----------VQRTILHDIAERGCIETLNLLLEHGANATARD-NMGNTPAHLAAEKGGSYRIQILDRLSQNCT--LTEVNAAGKTPLDLANEGN--H--REVVEWLQNFCM* 166
Mesocentrotus franciscanus              47  VMNSSSLSTARILIQHDADLNRLDPS----------VQRTILHDIAEKGCIETLILLLEHGADATARD-NMGNTPAHLAAREGGHSRIEILRRLSQNCT--LTEVNAAGKTPLDLANEGN--H--GEIVEWLQNFRM* 166
Hemicentrotus pulcherrimus              47  -----------------ADLNRPDPA----------VQRTILHDIAETGCIETLELLLEHGADATARD-NMGNTPAHLAARKGSSYRIRVLERLSQNCT--LTEVNTAGKTPLDLANEGN--H--GVIIEWLQNFHTRVFPL--QFL----VRRVI-------------RK--RVR-IISNLQNILPSIVINYLYGR* 233
Paracentrotus lividus                   48  VMNSMSPSTARILLQHGADPNIPDPD----------VRRTILHDIAESGCFQILELLLDHGANATARD-NQGNTPAHLAARQGGSSRIEILERLSQSCT--LSEMNTAGETPLDLATQNN--H--GDIVTWLQNFRM* 167
Heliocidaris erythrogramma              47  VVNLMSPNTVHILLQHGADPNIPDQN----------TQKTILHDMAEKGCPDSLELLLIHGANATARD-NMGNTPAHLAAREGGPSRIEILERLSLHCT--FTEVNNAGKTPLDIAMEKN--H--GDIVELLQNC-NQVFPL--VFL----ARRVI-------------RM--RVR-IVQDLQHLLPPILIDYLNGR* 203
Heliocidaris tuberculata                47  VMNSMSPSTARILLQHGAKPNVRDPD----------VQRTILHDIAESGCLGTLQLLLQHGADATAQD-NMGNTPAHLAAREGGQSRIEILERLSQSCY--LNEVNRAGETPLALAIQNN--H--RDIVEWLQNFHNQIFPL--VFL----ARRVI-------------RM--RVR-IMQDLQDLLPSILIDYLNGR* 204
Lytechinus pictus                       57  VMNSMSESTARTLINHGADVNIPDPD----------VQRTPLHEVAETGSIEVLDLLLQNGADVIARD-KWGNTPAHLAASKGR--DIKVLERLSQFCT--LREVNTEGKTPLDLAREKG--H--QHVVEWIQNHHTRLYSL--QFL----VRRLI-------------RI--RVR-VIGNSVGNLPRPLIDYLHGK* 212
Lytechinus variegatus (X1)              57  VMNTMCESTARTLINHGANVNIPDPD----------VRRTILHDVAEKGSIEVLDLLLQNGADVIARD-NWGNTPAHLAASEGR--DIKVLERLSQFCT--LREVNTEGNTPLDLAREKG--H--QQVVEWIQNHHTRVYSL--QFL----VRRLI-------------RI--RVR-VIGNSVGNFPRPLMDYLHGR* 212
Lytechinus variegatus (X2)              57  VMNTMCESTARTLINHGANVNIPDPD----------VRRTILHDVAEKGSIEVLDLLLQNGADVIARD-NWGNTPAHLAASEGR--DIKVLERLSQFCT--LREVNTEGNTPLDLAREKG--H--QQVVEWIQNHRT* 174
Eucidaris tribuloides                   1   AMNGMCPRTAEILLAHGADPNVHDPT----------TGNTPLHDIAYNGCLETLNLLLQSDADPLARN-NRGDTPAHLAAEYGTQSRIA 78
Apostichopus japonicus                  1   AANFAIPEVVEYLLQKGANPNVYDMNP---DSDHRPFLRTPLHDALHHGNEDSAILLIQHGADLTAVD-NRGDLPLHLAVQK---------------CS--LRVRNTAR-------------------PSWLCQVYNQTFYL* 102
Patiria miniata  (X1~4)                     VMTMATPMVAQFLLEKGADPNVPDPD----------VGRTPLHDAAEHGHEDTVRWLLAHGADPTAREWRRGNTPAHLAAEEGRPV--GILSLLSRGCS--FWETNNAGQTPLDGAIANN--R--QATVQWITDYHSSGEGRTLKHL----CRLTIH--------RALGTQ--RMRRVGELTSFPLPETLVDYLKMTR* 220~286
Acanthaster planci (X1~3)                   VMTMAIPEVAEFLLKSGADPNVRDRD----------VGRTPLHDAAEHGHEDTVRLLLKHGADPLVRD-MRGDTPAHLAAVEGRPP--QILELLSRGCS--MWERNNAGRTPLDEADRNE--R--RETKRWIQNYHSSGKGTALKHL----CRLAIH--------RVLGTQ--RLPRVRELIGSTLPKALVEYLEMAR* 219~309
Patiriella regularis                    55  VMTMAMPDVCQFLLELGAEPNEPDPD----------VGRTPLHDAAEHGHEDTVRLLLAHGADPVVTE-KHGNTPAHLAAGEGRPA--R 130
Asterias rubens  (X1, X3)                   VMTMTRPEVAEFLLGLGANPNEPDPD----------VGRTPLHDAAEHGHPDTVELLLKYGADPRLRDSARGNTPAHLAAGEGRPL--RILELLSNGCT--LRERNSAGLTPHDVAIANG--N--EDAAEWIASYHSSGRGVRLMHL----CRLTIH--------QVLGTQ--RLCRIRELSNYPLPDMLVDFIKMYR* 219~265
Asterias rubens  (X2)                   100 VMTMTRPEVAEFLLGLGANPNEPDPD----------VGRTPLHDAAEHGHPDTVELLLKYGADPRLRDSARGNTPAHLAAGEGRPL--RILELLSNGCT--LRERNSAGLTPHDVAIANG--N--EDAAEWIASYHSTFG* 221
Pisaster brevispinus                    53  VMTMTRPEVAEFLLGLGANPNEPDPN----------VGRTPLHDAAEHGHPETVELLLRYGADSKLRDIARGNTPAHLAAGEGRPL--RILELLSNGCT--LRERNNAGLTPHDVAIAND--N--EEAAEWIASYHSSGRGLRLMHL----CRLTVH--------QVLGTQ--RLCRIRELSNYPLPDLIVDYLKTCR* 218
Pisaster ochraceus                      53  VMTMTRPEVAEFLLGLGANPNEPDPN----------VGRTPLHDAAEHGHPETVELLLRYGADPKLRDIARGNIPAHLAAGEGRPL--RILELLSNGCT--LRERNNAGLTPHDVAIANN--N--EEAAEWIASYHSSGRGFRLMHL----CRLTVH--------QVLGTR--RLCRIRELSNYPLPDLIVDYLKTCR* 218
Plazaster borealis                      53  VMTMTRPEVAEFLLGLGANPNEPDPD----------VGRTPLHDAAEHGHPDTVELLLRYGADPMLRDIARGNTPAHLAAGEGRPL--RILELLSSGCT--LRERNNAGLTPHDVATANG--N--EEAAEWIASYHSSGRGVRLTHL----CRLTIR--------QVLGTQ--RLCRIQELSIYPLPDMIVDYIKISR* 218
Ophioderma brevispina                   53  TMNFHYPEIAQILLDEGADPNVRDPD----------VGRTPLHDAVEYGIPSTVMCLLQGGADVTAKENLFGNTPVHIAAEEGKEL 128
Ophionereis fasciata                    46  TMKFTYTEIAKILLDEGADPNVKDSD----------VGRTPLHDAVQYGTLSTVMCLLQGGADVTIKEDVFGNTPVHIAA 115
Anneissia japonica                      45  AVPVSCPEIVSILLEYGANPNLKEPV----------MGGIPLHQAAREGATATLELMLQFGANVNAQDK-NGDTPAHLAARKGYLEAFKILN---PLAD--MTIRNHNGDTPLSIAD--R--H--PEFRDWINVANVDGQRNT* 165
Nesometra (Dorometra) sesokonis         43  AVPASKFKIASILLEYGANPHLPEPV----------MGGIPLHNAAREGATTTLELLLQYGSNVNAQDKANGDTAAHLAAKKGHLEAFKILS---SITD--MTIKNHSGDTPLSIIE--Q--H--E* 147
Onychophora
Epiperipatus broadwayi                  49  VMSFSRMFAVEILLKAGANVNVSDV-----------TLRTPVHDAAEAGFLDVVQNLIAYGADITIQD-FKGNTPAHLASKKGH---FEVVQFLAQGMS--LMLQNMEGQTVFDTDAMKKLSENSPVHKKWIVTYLG--LYKLYIYMIKFIYLYIY--V* 186
Euperipatoides rowelli                  49  AVRLSNLKCVDSLLSAGANPNICDR-----------TLRTPVHDAAEAGFLDVLQCLLANGGVIVNGD-QWGNTPAHLAAKKGF---INVVQFLASGMD--LNLINNDGSTVFDIEAMKRQRANSSCDKLWMDSYRG--LH* 170
Annelida
Lamellibrachia luymesi                  62  VTKSSRVDIFEVLLRNGADPNVRDRS--YP--GHQDTLVTPAHDVARAGYTDVLKYLLRYGADVNIRD-SWGNTPAHLAAKYGH---YDVVRYLSHATD-LRKAVNMHGVTPLDYHTGRQ--GKDDSQWEWVQETKV--NPLRLDDLATIRVRTAM-------------TG--RL-----RQCHHLPVADKLK---ERLFLQFGDD* 231
Lamellibrachia satsuma                  62  VTKSSRVDIFEVLLRNGADPNVRDRS--YP--GHQDTLVTPAHDVARAGYTDVLKCLLRYGADVNIRD-SWGNTPAHLAAKYGH---YDVVRYL----D-LRKAVNMHGVTPLDYHTGRQ--GKDGSQWEWVQETKV--NPLRLDDLATIRVRTAM-------------TG--RL-----RQCHHLPVADKLK---ERLFLQFGDD* 227
Ridgeia piscesae                        62  VTKSSRLEIFEILLRNGADPNARDRS--YP--GHQDTLVTPAHDVARAGYLDVLKCLLRFGADVNIRD-SWGNTPAHLAAKYGH---YDVVMYLSHATE-LRNAVNMHGLTPLDYVTGSE--GKDGGHCEWMQEMKV--NPLRLDDLATISVRAAM-------------TG--RL-----RQCRHLPIADKLK---AQLLLQFGDD* 231
Paraescarpia echinospica                62  VTKSSRLDIFEILLRNGADPNARDRS--YP--GHQDTLVTPAHDVARAGYLDVLKCLLRYGADVDIRD-SWGNTPAHLAAKYGH---YDVVMYLSHATD-LRKAVNMHGLTPMDYSTGSQ--GQDGGQCEWMQEMKV--NPLTLEDLTTICVRAAM-------------TG--RL-----RQCRCLPIADKLK---AQLLLQFGDD* 231
Dimorphilus gyrociliatus                167 VMNLSYPDVAEQLLKAGANPNVQDRG--IG--NGRCLLLTPGHDAARVGYVRTLKVLLKYGADTNIRD-GHGNTLAHLAAKHGQ---LSVLKVLARGSD-L-SIRNLAGQTPLDYLMPEH--ARSESEVAFLTELKT--QPRPLKYL----AVTSIR--------KHLGTE--RLA-LLHRLPLPP--PLKDNLILK* 332
Sipunculus nudus                        71  VMNFSQVSLATLLLEAGADPNVVDHS--KN--NKKDTLLTPAHDVARQGYLDTLKVLLWYGADISLRD-ARGNTPAHLAARYGH---YEILNYLAEGVD-LRSPCNQAGLSPLDVFEKTK--A-GVRHQAWILNQRG-------ELT----CVYSHA* 204
Mollusca
Marisa cornuarietis                     325 VMAHDNADLAEVLLEAGGDPNALDVG--KR--TDTSTLRTIAHDVAASGYTDVLRCLLRHGADISLQD-RWGNTPLHLAAKAGH---LSTVRFLWKGAS-L-YIPNHLGKTALNIMEESK----NLHIQQWVKKRKC--TVPRLKIL----CTLAVR--------KELGPH--GLR-WWRKHAP--DEAIQRDIFLQSSKAATLMPSA---SASLSTEYS
Pomacea canaliculata                    313 VMAHDNADLAEVLLEAGGDPNALDIG--KR--TDTSTLRTVAHDVAASGYTDVLRCLLRHGADISLQD-RWGNTPLHLAAKAGH---LSTVRYLRRGAS-L-NIPNHLGKTALNIMEESK----NLQIQQWVKKRKC--TVPRLKTL----CTLAVR--------KELGAH--GLR-CWRKHVP--DEAIWRQIFLQSSRAATFMPSA---SASLSTEYS
Pomacea maculata                        313 VMAHDNADLAEVLLEAGGDPNALDIG--KR--TDTSTLRTVAHDVAASGYTDVLRCLLCHGADISLQD-RWGNTPLHLAAKAGH---LSTVRYLRRGAS-L-NIPNHLGKTALNIMEESK----NLQIQQWVKKRKC--TVPRLKTL----CTLAVR--------KELGAH--GLR-CWRKHVP--DEAIWRQIFLQSSRAATFMPSA---SASLSTEYS
Alviniconcha marisindica                142 VMTHDNTDLAEVLLRHGADPNVQDKG--RR--AANSTYCTIAHDVAASGYTDTLRCLLRHGADVNLQD-RWGNTPLHIAAKEGH---LATVRLLRRGSS-L-YLPNQQGKTPMMLLEAAG----NSAAKQWIKLKKT--TVSRLQVL----CVLVVR--------RELGPR--GLR-CWRKHAPRDDVQLQRQVLLQSRGGRPPPPLA-SASTLLSSRHS
Babylonia areolata                      55  VMSVNNTDLAEVLLKAGANPNMQDKG--RR--AATSTLRTIAHDVADSGNTDMLRCLLRHGADVNLQD-RWGNTPLHLAAKGGH---LAAVRLLRRGTS-L-DQPNLQGKTPIMLLEESR----DSAIREWVAMKKS--TVSRLQTL----CVLVVR--------RRLGPD--RLR-LWKKQAM-TDVKLQRQVLLQNQGSRPHPHASTSTSTILSSQHS
Anentome helena                         1   VMSVTNTDVAEVLLKAGADPNVQDQG--RR--AATSTFRTIAHDVADSGNTDMLRCLLRHGADVSL 62
Conus betulinus                         88  VMSVNNTDVAEVLLKAGADPNMQDKG--RR--AATSTLRTIAHDVADTGNTDMLRCLLRHGADVNLQD-RWGNTPLHLAAKGGH---LAAVRLLRRGTS-L-YLPNLQGKSPMMLFGRNE* 197
Conus consors                           74  VMSVNNTDVAEVLLKAGANPNMQDKG--RR--AATSTLRTIAHDVADTGNTDMLRCLLRHGADVNL 135
Conus tribblei                          158 VMSVNNTDVAEVLLKAGANPNMQDKG--RR--AATSTLRTIAHDVADTGNTDMLRCLLRHGADVNL 219
Conus ventricosus                       97  VMSVNNTDVAEVLLKAGANPNMQDKG--RR--AATSTLRTIAHDVADTGNTDMLRCLLRHGADVNLQD-RWGNTPLHLAAKGGH---LAAVRLLRRGTS-L-DLPNLQGKSPMMLLEEMD----DSAIREWVKMKKS--TVGRLQTQ----CVLVLR--------QQLGPE--GLQ-HWRKHA--PDVNLQRQVLLQSQGTRSHHTS---TSTIHSSQHS
Nodopelta heminoda                      33  VMTHSYVDLTEALLRAGADPNVKDVS--LRQDGGSSTFRTICHDVAESGALSTLRCLLRHGADINARD-KFGNTPAHVAAKGGH---FEVVQYLSCGMS--QNVVNIYERTPLDSLENIV--D--PAFRQW 151
Patella depressa                        53  VMAHNIPELATVLLHAGANPNIQDIS--TRDTDCLSTYRTICHDVAASGYLDTLHVLLKYGTDTNMVD-RWGNTPAHSAAKAGH---FKIVQYLACGMA--MTAVNSLGSTPLDYLADLD--D--VACQQWIDKQRS--KFCENKIWY------LYIYSNHVSSKP* 198
Patella pellucida                       53  VMAHNIPELATVLLHAGANPNIQDIS--TRDTDCLSTYRTICHDVAASGYLDTLQVLLKYGTDTNMVD-RWGNTPAHSAAKAGH---FKIVQYLACGME--MTVVNSLGSTPLDYLADLD--D--VDCQQWVDKQRS--KFYEN* 182
Patella vulgata                         53  VMAHNIPELATVLLHSGANPNNQDIS--TRDTDCLSTYRTICHDVAASGYLETLHVLLKYGTDT--VD-RWGNTPAHSAAKAGH---FKIVQYLACGMA--MTAVNSLGSTPLDYLADLD--D--VVCQQWIDKQKS--KFPEKINEMAFNVLLFFSYTGLMKTGIRHTVC--YEG-DFARQCHYGLSFTCMPIYTQDLGFSSHPND* 240
Haliotis cracherodii                    52  VMTHTYVDLADELLCAGADPNVKDIS--HRHIGGTSTYRTICHDVAESGSLPTLQCLLRHGADISLRD-KRGNTPAHVAAKSGH---FKLVQYLLCGMA--TNVTNDLKRTPLDYLDSKE--D--PGCRQWVERRRS--TPATLRTL----CVVVVR--------SALGPG--RLK-DLSXXCS------CR--GQLSAVYSFKTCVL-------SRQR* 226
Haliotis laevigata                      52  VMTHTYVDLADELLCAGADPNVKDIS--HRHIDGTSTYRTICHDVAESGSMPTLQCLLRHGADISLRD-KRGNTPAHIAAKSGH---FKLVQYLLCGMA--TNVTNDMKRTPLDYLESKE--D--PGCQQWVERRRS--TPATLRTL----CVVVVR--------SALGPG--RLK-DLSPACS------CR--DQLSAVYSFKTCVH* 222
Haliotis rubra                          52  VMTHTYVNLADELLCAGADPNVKDIS--HRHIDGTSTYRTICHDVAESGSLPTLQCLLRHGADISLRD-KRGNTPAHIAAKSGH---FKLVQYLLCGMA--TNVTNDMKRTPLDYLESKE--D--PGCQQWVERRRS--TPATLRTL----CVVVVR--------SALGPG--RLK-DLSPDCS------CR--DQLSAVYSFKTCVH* 222
Haliotis rufescens                      52  VMTHTYVDLADELLCAGADPNVKDIS--HRHIGGTSTYRTICHDVAESGSLPTLQCLLRHGADISLRD-KRGNTPAHVAAKSGH---FKLVQYLLCGMA--TNVTNDLKRTPLDYLDSKE--D--PGCRQWVERRRS--TPATLRTL----CVVVVR--------SALGPG--RLK-DLSRLQLPGPVVSCVQLQNLRALEAEIDC* 228
Gibbula magus                           52  VMTHSYVELAEEELLCAGADPNVKDIS---LHNGGTSTYRTICHDVAESGALLTLKSLLRHGADINLRD-QMGNTPAHIAAKGGH---LQIVQYLLCGMS--QNVTNHNQFSPLDYLDRIE--D--PGCKQWVDRRRS--TPLTLRSL----CVVVLR--------AALGPG--RLG-DLTSLQLPTPAVSCIQLQNRHTLDSEIESRY-------DL* 231
Phorcus lineatus                        52  VMTHSYVELAEEELLCAGADPNVKDIS---LHNGGTSTYRTICHDVAESGALTTLKSLLRHGADINLRD-KMGNTAAHIAAKGGH---LQIVQYLLCGMS--QNVNNHNQCTPLDYLDRIE--D--PGCKQWVDRRRS--TPLTLRSL----CVVVVR--------AALGPG--RLA-DLTTLQLPNPAVSCIQLQNQHTLDSEIDC* 227
Steromphala cineraria                   53  VMTHSYVELAEEELLCAGADPNVKDIS---IHNGGTSTYRTICHDVAESGALSTLKLLLRHGADINLRD-KMGNTPAHIAAKGGH---LQIVQYLLCGMS--QNVNNHNQCTPLDYLDRIE--D--PGCKQWIDRRRS--TPLTLRSQ----CVVVIR--------EALGPG--RLA-DLTSLQLPNPAVSCIQLQNQHTLDSEIDC* 227
Nautilus pompilius                      57  VMNRCKPELAEALLRAGADPNLMEGD----------TRRTICHDVAESGYLDTLRCLLKHGADLSRRD-RWGDTPAHCGAKRRQ---LSVVQYLSCGFP--IGAVNGQGRTVLDILADFD--DC-SAFRSWVAEKRG--KKLFADPSPGVLFLLLFTAACIQS--RERGGG--VMG-----EREVRPARSCRPVLT* 222
Brachiopoda/Phoronida
Lingula anatina                         67  VFKCSRPDIAEILLKHGADPNIQDKH---FQDLHLSSYRTPAHDIAEQGFLDTMKVLLKHGADVTLRS-KMGYSPAHIAAEKGR---LDTLEYLSGAID-LMNCRADDGSTPFDLIPKSI--TLTDRQRTWVEAEKN-AKPKSLKYL----TRLRIR--------HCMRHNLKRLQ-TFPTL----PLSIKKYLLLQD* 236
Phoronis australis                      47  VTKTSCHHIVQELLERGADPDVPDVG----ENLERSTYRTITHDVCSHGFLRTLQVLAAYGANLSAVD-KYGNCPAHYAARKSN---ITILRFISKGVF-LSRQINGEGKSPLDIIEETG--DIDDSDCSWVSSERS-KC---LLRL----CTM* 181
Phoronis ovalis                         1   VMKLSDPRIARILLEHGADPNLKERSPHHEADPSRSLLRTPAHNTCCEGFLDTLKTLLRCGANIIQDF-K-GDTPAHCAAKYNH---FDIVQYLSVGMI--GNETNHSGETVVDILYKSN--L-TADEKSWIQQQIG--IIKFSHFF-------VSTSHCI* 142
Phoronis pallida                        39  VMNTSCHDIAEILLMRGADPNIQDT----------STLRTILHDVCEFGYMDTLQCLISFGANLQVCD-KWGNTPTHYAAKKGN---YDMVQYLLEHGS-SVNQRNSDGQSPLDILEICG--TLSEQQKTWISQERG-EL-G---------FILTGD* 167

(cont'd)
Marisa cornuarietis                         TKSQKSTS---------QDDAPGPSFTLRSEHPLEMLSLRDLVKRRLSSGENTSLEVQNCSSIHSGDDMSL--GAAVTGSLLSETSSVFENTFSSVDPS-HDESYFSSDHFKDELGGIKKVQK* 619
Pomacea canaliculata                        TKTQKFTI---------QEDAPGPSFTFRSEHPLEMLSLRDLVPRRLSSGENTSLEVQNCNSIHSGDDMSL--GAAVTGSLLSETSSVFENTFSSVDPS-HDESLFSSDQFTEELGGIKKVASAK* 609
Pomacea maculata                            TKRQKSTI---------QEDAPGPSFTFRSEHPLEMLSLRDLVPRRLSSGENTSLEVQNCNSIHSGDDMSL--GAAVTGSLLSETSSVFENTFSSVDPS-HDESLFSSDQFTEELGGIKKVASAK* 609
Alviniconcha marisindica                    AGSRSLQSTRGQANGGGCRPVLPPGLSAMSRHVAEFLSLTDLVPRRLSTGENTSLEVQNTSSGQSNDDVSL--GAHATSSLASETGSLFETNLSSIGAASRSADELSSPSLDQSV* 442
Babylonia areolata                          TVSRSLMA-RGRSNNSNGHPLLP-GMST-GKQMTDFLTLRDLIPRRLSTGENTSLEVHNTSSGQSNDDVSIVAGAQATSSLASETGSLFETNLSSIET-SQSADDVSLPSFDHSL* 353
Conus ventricosus                           AGSRSLTX--QGRSNSNGHPLLP-GMST-GKQMTDFLSLRDLIPRRLSTGENTSLEVHNTSSGQSNDDVSL--GAHATSSLASETGSLFETNLSSIEA-SHSADDLSLPSFDQSL* 364
```

**Figure S1b**

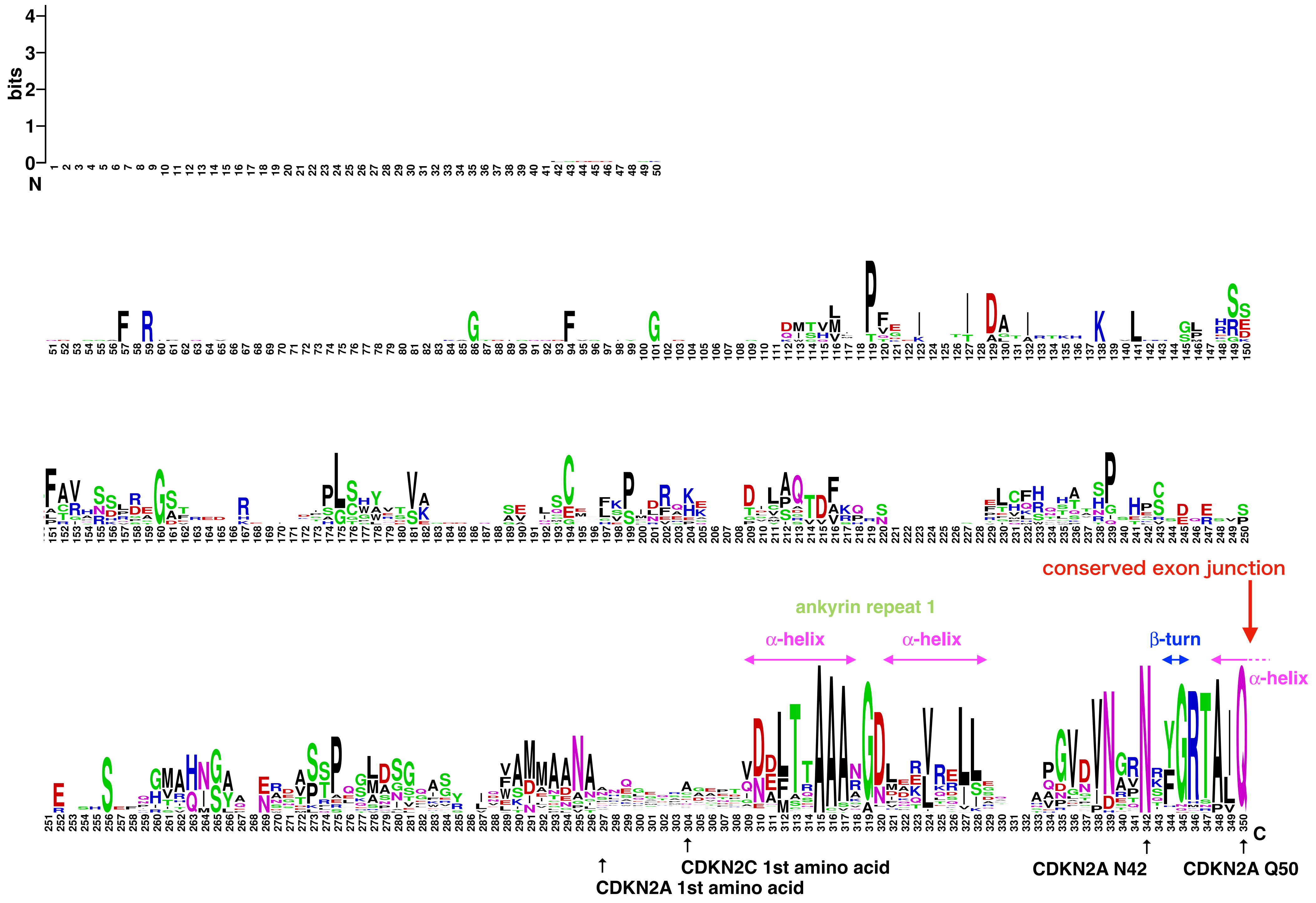
bits
N
C
conserved exon junction
ankyrin repeat 1
α-helix
α-helix
β-turn
α-helix
CDKN2C 1st amino acid
CDKN2A 1st amino acid
CDKN2A N42
CDKN2A Q50

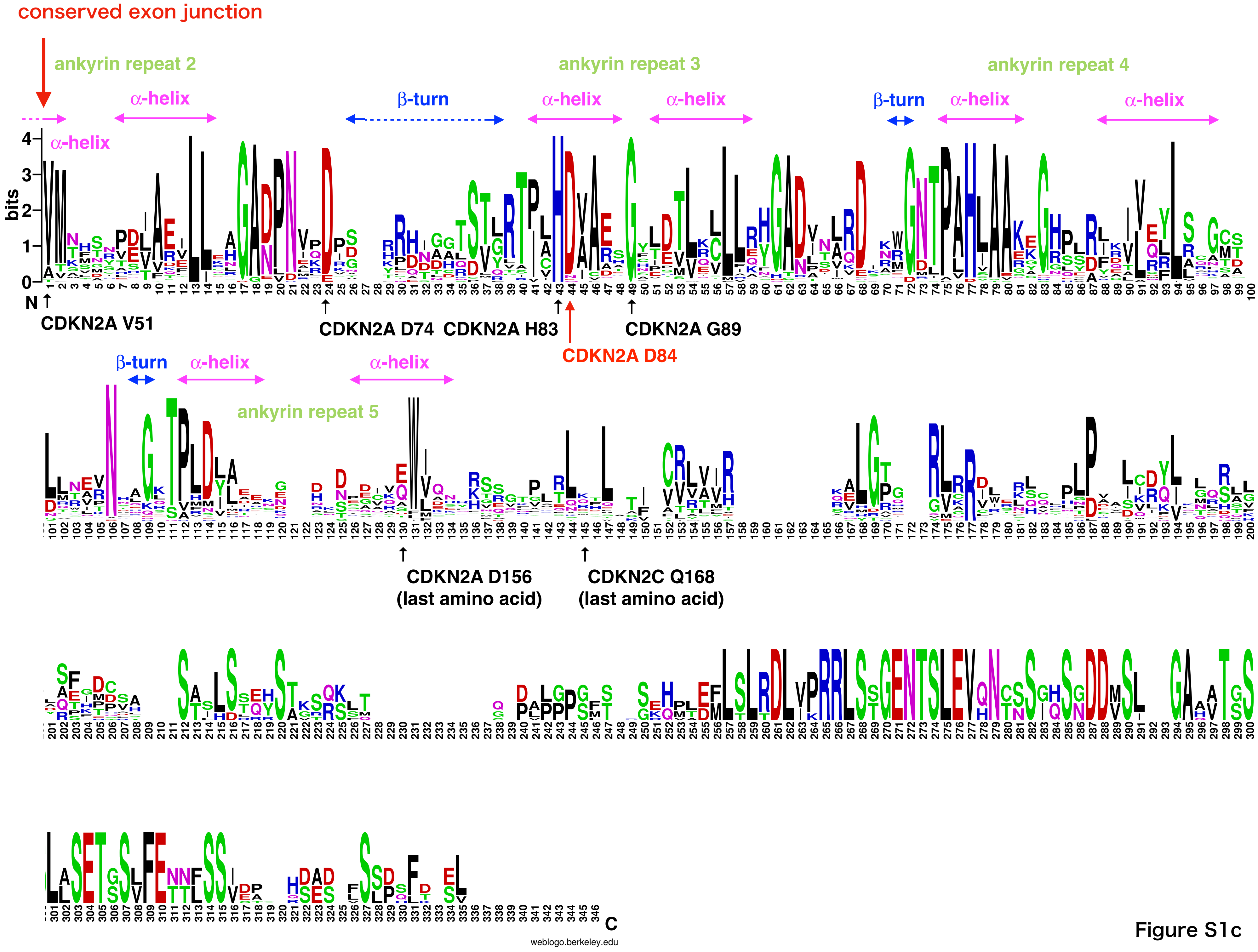
conserved exon junction
ankyrin repeat 2
ankyrin repeat 3
ankyrin repeat 4
ankyrin repeat 5
α-helix
β-turn
bits
N
C
CDKN2A V51
CDKN2A D74
CDKN2A H83
CDKN2A D84
CDKN2A G89
CDKN2A D156
(last amino acid)
CDKN2C Q168
(last amino acid)
weblogo.berkeley.edu

Figure S1c

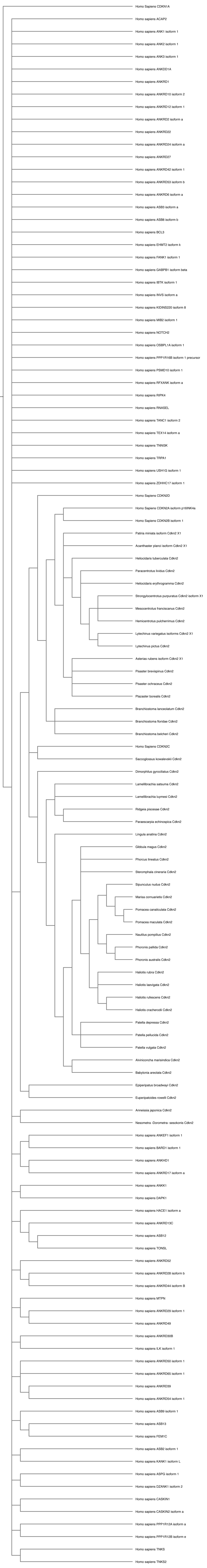
Homo Sapiens CDKN1A
Homo sapiens ACAP2
Homo sapiens ANK1 isoform 1
Homo sapiens ANK2 isoform 1
Homo sapiens ANK3 isoform 1
Homo sapiens ANKDD1A
Homo sapiens ANKRD1
Homo sapiens ANKRD10 isoform 2
Homo sapiens ANKRD12 isoform 1
Homo sapiens ANKRD2 isoform a
Homo sapiens ANKRD22
Homo sapiens ANKRD24 isoform a
Homo sapiens ANKRD27
Homo sapiens ANKRD42 isoform 1
Homo sapiens ANKRD53 isoform b
Homo sapiens ANKRD6 isoform a
Homo sapiens ASB3 isoform a
Homo sapiens ASB8 isoform b
Homo sapiens BCL3
Homo sapiens EHMT2 isoform k
Homo sapiens FANK1 isoform 1
Homo sapiens GABPB1 isoform beta
Homo sapiens IBTK isoform 1
Homo sapiens INVS isoform a
Homo sapiens KIDINS220 isoform 8
Homo sapiens MIB2 isoform 1
Homo sapiens NOTCH2
Homo sapiens OSBPL1A isoform 1
Homo sapiens PPP1R16B isoform 1 precursor
Homo sapiens PSMD10 isoform 1
Homo sapiens RFXANK isoform a
Homo sapiens RIPK4
Homo sapiens RNASEL
Homo sapiens TANC1 isoform 2
Homo sapiens TEX14 isoform a
Homo sapiens TNNI3K
Homo sapiens TRPA1
Homo sapiens USH1G isoform 1
Homo sapiens ZDHHC17 isoform 1
Homo Sapiens CDKN2D
Homo Sapiens CDKN2A isoform p16INK4a
Homo Sapiens CDKN2B isoform 1
Patiria miniata isoform Cdkn2 X1
Acanthaster planci isoform Cdkn2 X1
Heliocidaris tuberculata Cdkn2
Paracentrotus lividus Cdkn2
Heliocidaris erythrogramma Cdkn2
Strongylocentrotus purpuratus Cdkn2 isoform X1
Mesocentrotus franciscanus Cdkn2
Hemicentrotus pulcherrimus Cdkn2
Lytechinus variegatus isoforms Cdkn2 X1
Lytechinus pictus Cdkn2
Asterias rubens isoform Cdkn2 X1
Pisaster brevispinus Cdkn2
Pisaster ochraceus Cdkn2
Plazaster borealis Cdkn2
Branchiostoma lanceolatum Cdkn2
Branchiostoma floridae Cdkn2
Branchiostoma belcheri Cdkn2
Homo Sapiens CDKN2C
Saccoglossus kowalevskii Cdkn2
Dimorphilus gyrociliatus Cdkn2
Lamellibrachia satsuma Cdkn2
Lamellibrachia luymesi Cdkn2
Ridgeia piscesae Cdkn2
Paraescarpia echinospica Cdkn2
Lingula anatina Cdkn2
Gibbula magus Cdkn2
Phorcus lineatus Cdkn2
Steromphala cineraria Cdkn2
Sipunculus nudus Cdkn2
Marisa cornuarietis Cdkn2
Pomacea canaliculata Cdkn2
Pomacea maculata Cdkn2
Nautilus pompilius Cdkn2
Phoronis pallida Cdkn2
Phoronis australis Cdkn2
Haliotis rubra Cdkn2
Haliotis laevigata Cdkn2
Haliotis rufescens Cdkn2
Haliotis cracherodii Cdkn2
Patella depressa Cdkn2
Patella pellucida Cdkn2
Patella vulgata Cdkn2
Alviniconcha marisindica Cdkn2
Babylonia areolata Cdkn2
Epiperipatus broadwayi Cdkn2
Euperipatoides rowelli Cdkn2
Anneissia japonica Cdkn2
Nesometra -Dorometra- sesokonis Cdkn2
Homo sapiens ANKEF1 isoform 1
Homo sapiens BARD1 isoform 1
Homo sapiens ANKHD1
Homo sapiens ANKRD17 isoform a
Homo sapiens ANKK1
Homo sapiens DAPK1
Homo sapiens HACE1 isoform a
Homo sapiens ANKRD13C
Homo sapiens ASB12
Homo sapiens TONSL
Homo sapiens ANKRD52
Homo sapiens ANKRD28 isoform b
Homo sapiens ANKRD44 isoform B
Homo sapiens MTPN
Homo sapiens ANKRD29 isoform 1
Homo sapiens ANKRD49
Homo sapiens ANKRD30B
Homo sapiens ILK isoform 1
Homo sapiens ANKRD50 isoform 1
Homo sapiens ANKRD65 isoform 1
Homo sapiens ANKRD39
Homo sapiens ANKRD54 isoform 1
Homo sapiens ASB9 isoform 1
Homo sapiens ASB13
Homo sapiens FEM1C
Homo sapiens ASB2 isoform 1
Homo sapiens KANK1 isoform L
Homo sapiens ASPG isoform 1
Homo sapiens DZANK1 isoform 2
Homo sapiens CASKIN1
Homo sapiens CASKIN2 isoform a
Homo sapiens PPP1R12A isoform a
Homo sapiens PPP1R12B isoform e
Homo sapiens TNKS
Homo sapiens TNKS2

**Figure S1d**

**Figure S1** Amino acid alignment of invertebrate Cdkn2s. (a) N-terminal half of Cdkn2. (b) C-terminal half. The residue corresponding to D84 is in green if negatively charged, or in magenta if not. (c) Consensus residues in invertebrate Cdkn2. **a** and **b** represent amino acids of $\alpha$-helices and $\beta$-turns in *Homo sapiens* CDKN2s, respectively (Baumgartner et al., 1998). (d) Phylogenetic analysis of invertebrate Cdkn2 candidate sequences.
