## Supplementary Tables S1-S11 and Figures S1-S6 for "Evolution of the *Cdk4/6*–*Cdkn2* system in invertebrates": Figure S2.pdf

**Cdkn2a**
**Mammals**
**<Placentarians>**

| | |
|---|---|
| *Homo sapiens* | LHGAEPNCADPATLTRPVHDAAREGFLDTLVVLHRAGARL |
| *Acinonyx jubatus* | LHGADPNCADPATLTRPVHDAAREGFLDTLVVLHRAGARL |
| *Ailuropoda melanoleuca* | LHGADPNCADPTTLTRPAHDAAREGFLDTLVVLHRAGARL |
| *Aotus nancymaae* | LHGADPNCADPATLTRPVHDAAREGFLDTLVVLHRAGARL |
| *Artibeus jamaicensis* | LHGADPNCADPATLATPAHDAAREGFLDTLVALHRAGARL |
| *Arvicanthis niloticus* | FYGAESNCADPTTLSRPVHDAAREGFLDTLVVLHQAGARL |
| *Arvicola amphibius* | LYGAEPNCEDPATLSRPVHDAAREGFLDTLVVLHQAGARL |
| *Balaenoptera acutorostrata scammoni* | LHGADPNCADPDTLTRPVHDAAREGFLDTLVALHRAGARL |
| *Balaenoptera musculus* | LHGADPNCADPATLTRPVHDAAREGFLDTLVALHRAGARL |
| *Bison bison bison* | LHGADPNCADPATLTRPVHDAAREGFLDTLVALHRAGARL |
| *Bos indicus* | LHGADPNCADPATLTRPVHDAAREGFLDTLVALHRAGARL |
| *Bos mutus* | LHGADPNCADPATLTRPVHDAAREGFLDTLVALHRAGARL |
| *Bos taurus* | LHGADPNCADPATLTRPVHDAAREGFLDTLVALHRAGAQL |
| *Bubalus bubalis* | FHGADANCADPATLTRPVHDAAREGFLDTLVALHRAGARL |
| *Callithrix jacchus* | LHGAEPNCADPATLTRPVHDAAREGFLDTLVALHRAGARL |
| *Callorhinus ursinus* | CHGAEPNCADPTTLTRPVHDAAREGFLDTLVVLHRAGARL |
| *Camelus bactrianus* | LHGADPNCADPATLTRPVHDAAREGFLDTLVALHRAGARL |
| *Camelus dromedarius* | LHGADPNCADPATLTRPVHDAAREGFLDTLVALHRAGARL |
| *Camelus ferus* | LHGADPNCADPATLTRPVHDAAREGFLDTLVALHRAGARL |
| *Canis lupus familiaris* | LHGADPNCADPVTLTRPVHDAAREGFLDTLVVLHRAGARL |
| *Capra hircus* | LHGADPNCADPATLTRPVHDAAREGFLDTLVALHRAGARL |
| *Carlito syrichta* | LHGAEPNCADPATLTRPVHDAAREGFLDTLVALHRAGARL |
| *Castor canadensis* | LHGAEANCPDPATLTRPVHDAAREGFLDTLVALHGAGARL |
| *Cavia porcellus* | LHGADPNCADPVTLARPVHDAAREGFLDTLVELHQAGARL |
| *Cebus imitator* | LHGAEPNCADPATLTRPVHDAAREGFLDTLVVLHRAGARL |
| *Ceratotherium simum simum* | LHGADPNRADPATLTRPVHDAAREGFLDTLVALHRAGARL |
| *Cercocebus atys* | LHGAEPNCADPATLTRPVHDAAREGFLDTLAVLHRAGARL |
| *Cervus canadensis* | LHGADPNCADPATLTRPVHDAAREGFLDTLVALHRAGGRL |
| *Cervus elaphus* | LHGADPNCADPATLTRPVHDAAREGFLDTLVALHRAGGRL |
| *Cervus hanglu yarkandensis* | LHGADPNCADPATLTRPVHDAAREGFLDTLVALHRAGGRL |
| *Chinchilla lanigera* | LHGADPNCADPATLARPVHDAAREGFLDTVVALHQAGARL |
| *Chlorocebus sabaeus* | LHGAEPNCADPATLTRPVHDAAREGFLDTLVVLHRAGARL |
| *Choloepus didactylus* | LHGAEPNCADPVTLARPVHDAAREGFLDTLLALHRAGARL |
| *Chrysochloris asiatica* | LHGAEPNLADPTTLTHPVHDAAREGFLETLRLLHRAGARL |
| *Colobus angolensis palliatus* | LHGAEPNCADPATLSRPVHDAAREGFLDTLAVLHRAGAQL |
| *Condylura cristata* | LHGADPNCADPATLTRPVHDAAREGFLDTLVALHQAGARL |
| *Cricetulus griseus* | LYGAEPNCEDPATLSRPVHDAAREGFLETLKILHQAGARL |
| *Crocuta crocuta* | LHGADPNCADPATLTRPVHDAAREGFLDTLVVLHRAGARL |
| *Dasypus novemcinctus* | VHGADPNCADPATLTRPVHDAAREGFLDTLLELHRAGARL |
| *Delphinapterus leucas* | LHGADPNCADPATLTRPVHDAAREGFLDTLVALHRAGAQL |
| *Desmodus rotundus* | LHGADPNCADPATLNRPVHDAAREGFLDTLVALHRAGARL |
| *Dipodomys ordii* | LHGAEPNRVDPATLTRPVHDAAREGFLDTLVMLHRAGAQL |
| *Dipodomys spectabilis* | LHGAEPNWVDPATLTRPVHDAAREGFLDTLVMLHRAGAQL |
| *Echinops telfairi* | RHGADPNCVDPTTLTRPVHDAAREGFLDTLKMLHQAGAHL |
| *Elephantulus edwardii* | SYGAEPNLADPVTLTRPVHDAAREGFLDTLMVLHRNGARL |
| *Enhydra lutris kenyoni* | LHGANPNCADPITLTRPVHDAAREGFLDTLLVLHRAGAQL |
| *Eptesicus fuscus* | THGADPNRPDPATLTRPVHDAAREGFLDTLVALHRAGARL |
| *Equus asinus* | LQGADPNRADPVTLTRPVHDAAREGFLDTLVALHRAGARL |
| *Equus caballus* | LHGADPNRADPVTLTRPVHDAAREGFLDTLVALHRAGARL |
| *Equus quagga* | LHGADPNRADPVTLTRPVHDAAGEGFLDTLVALHRAGARL |
| *Erinaceus europaeus* | LHGAEPNCADPATLTRPVHDAAREGFLDTVMALHRAGARL |
| *Eumetopias jubatus* | CHGAEPNCADPTTLTRPVHDAAREGFLDTLVVLHRAGARL |
| *Felis catus* | LHGADPNCADPATLTRPVHDAAREGFLDTLVVLHRAGARL |
| *Fukomys damarensis* | LHGADPNCADPVTLTRPVHDAAREGFLDTLVALHRAGARL |
| *Galemys pyrenaicus* | LHGADPNCADPDTLTRPVHDAAREGFLDTLVVLHQAGARL |
| *Galeopterus variegatus* | LHGAEPNCADPATLTRPVHDAAREGFLETLVALHRAGARL |
| *Globicephala melas* | LHGADPNCADPATLTRPVHDAAREGFLDTLVALHRAGARL |
| *Gorilla gorilla gorilla* | LHGAEPNCADPATLTRPVHDAAREGFLDTLVVLHRAGARL |
| *Grammomys surdaster* | FYGAESNCADPTTLSRPVHDAAREGFLDTLVVLHQAGARL |
| *Gulo gulo luscus* | LHGADPNCADPITLTRPVHDAAREGFLDTLLVLHRAGARL |
| *Halichoerus grypus* | CHGAEPNCADPTTLTRPVHDAAREGFLDTLVLLHRAGARL |
| *Heterocephalus glaber* | LHGADPNCADPVTLTLPVHDAARAGFLDTLVALHRAGARL |
| *Hipposideros armiger* | LHGADPNCADPATLTRPVHDAAREGFLDTLAVLHRAGARL |
| *Hyaena hyaena* | LHGADPNCADPATLTRPVHDAAREGFLDTLVVLHRAGARL |
| *Hylobates moloch* | LHGAEPNCADPATLTRPVHDAAREGFLDTLAVLHRAGARL |
| *Ictidomys tridecemlineatus* | LHGAEPNCADPATLTRPVHDAAREGFLDTLVALHLAGARL |
| *Jaculus jaculus* | LHGAEPNCADPATLTRPVHDAAREGFLDTLAALHRAGARL |
| *Lagenorhynchus obliquidens* | LHGADPNCADPATLTRPVHDAAREGFLDTLVALHRAGARL |
| *Lemur catta* | LHGAEPNRADPATLTRPVHDAAREGFLDTLVALHRAGARL |
| *Leopardus geoffroyi* | LHGADPNCADPATLTRPVHDAAREGFLDTLVVLHRAGARL |
| *Leptonychotes weddellii* | CHGAEPNCADPTTLTRPVHDAARDGFLDTLVVLHRAGARL |
| *Lipotes vexillifer* | LHGADPNCADPATLTRPVHDAAREGFLDTLVALHRAGARL |
| *Lontra canadensis* | LHGADPNRADPITLTRPVHDAAREGFLDTLLVLHRAGARL |
| *Loxodonta africana* | LHGAEPNCADPATLTRPVHDAAREGFLDTLMVLHRAGARL |
| *Lynx canadensis* | LHGADPNCADPATLTRPVHDAAREGFLDTLVVLHRAGARL |
| *Lynx pardinus* | LHGADPNCADPATLTRPVHDAAREGFLDTLVVLHRAGARL |
| *Macaca fascicularis* | LHGAEPNCADPATLTRPVHDAAREGFLDTLAVLHRAGARL |
| *Macaca mulatta* | LHGAEPNCADPATLTRPVHDAAREGFLDTLAVLHRAGARL |
| *Macaca nemestrina* | LHGAEPNCADPATLTRPVHDAAREGFLDTLAVLHRAGARL |
| *Mandrillus leucophaeus* | LHGAEPNCADPATLTRPVHDAAREGFLDTLAVLHRAGARL |
| *Manis javanica* | LHGADPNCTDPTTLTRPVHDAAREGFLDTVVALHRAGAQL |
| *Manis pentadactyla* | LHGADPNCTDPTTLTRPVHDAAREGFLDTVVALHRAGARL |
| *Marmota flaviventris* | LHGAEPNCADPATLTRPVHDAAREGFLDTLVALHLAGARL |
| *Marmota marmota marmota* | LHGAEPNCADPATLTRPVHDAAREGFLDTLVALHLAGARL |
| *Marmota monax* | LHGAEPNCADPATLTRPVHDAAREGFLDTLVALHLAGARL |
| *Mastomys coucha* | LYGADSNCADATTLSRPVHDAAREGFLDTLVVLHRAGARL |
| *Meles meles* | LHGADPNCADPITLTRPVHDAAREGFLDTLLVLHRAGARL |
| *Meriones unguiculatus* | FHGADPNCEDPATLSRPVHDAAREGFLDTLVVLHQAGARL |
| *Mesocricetus auratus* | LYGAEPNCEDPATLSRPVHDAAREGFLETLAILHQAGARL |
| *Microcebus murinus* | LHGAEPNCADPATLTRPVHDAAREGFLDTLVALHRAGARL |
| *Microtus ochrogaster* | LYGAEPNCEDPATLSRPVHDAAREGFLDTLVVLHQAGARL |
| *Microtus oregoni* | LYGAEPNCEDPTTLSRPVHDAAREGFLDTLVVLHQAGAQL |
| *Miniopterus natalensis* | THGADPNRADPTTLTRPLHDAAREGFLDTVAALHRAGARL |
| *Mirounga angustirostris* | CHGAEPNCADPTTLTRPVHDAARDGFLDTLVVLHRAGARL |
| *Mirounga leonina* | CHGAEPNCADPTTLTRPVHDAARDGFLDTLVVLHRAGARL |
| *Molossus molossus* | LHGADPNCADPDTLTRPVHDAAREGYLETLVVLHRGGARL |
| *Monodon monoceros* | LHGADPNCADPATLTRPVHDAAREGFLDTLVALHRAGAQL |
| *Mus caroli* | NYGADSNCEDPTTLSRPVHDAAREGFLDTLVVLHASGAWL |
| *Mus musculus* | NYGADSNCEDPTTFSRPVHDAAREGFLDTLVVLHGSGARL |
| *Mus pahari* | FYGGDSNCEDPTTLSRPVHDAAREGFLDTLVLLHRSGARL |
| *Mustela erminea* | LHGADPNCADPVTLTRPVHDAAREGFLDTLVVLHRAGARL |
| *Mustela putorius furo* | LHGADPNCADPVTLTRPVHDAAREGFLDTLVALHRAGARL |
| *Myotis davidii* | IHGADPNCADPATLTRPVHDAAREGFLDTLVALHRAGARL |
| *Myotis lucifugus* | IHGADPNCADPATLTRPVHDAAWEGFLDTLVALHRAGAQL |
| *Myotis myotis* | IHGADPNCADPATLTRPVHDAAREGFLDTLVALHRAGARL |
| *Nannospalax galili* | LHGADPNCADPATLTRPVHDAAREGFLDTLVALHQAGARL |
| *Neogale vison* | LHGADPNCADPVTLTRPVHDAAWEGFLDTLVVLHRAGARL |
| *Neomonachus schauinslandi* | CHGAEPNCADPTTLTRPVHDAARDGFLDTLVVLHRAGARL |
| *Neophocaena asiaeorientalis asiaeorientalis* | LHGADPNCADPATLTRPVHDAAREGFLDTLVALHRAGARL |
| *Neosciurus carolinensis* | LHGAEPNRADPATLTRPVHDAAREGFLDTLVALHQAGARL |
| *Neotoma lepida* | LYGAEPNCEDPTTLSRPVHDAAREGFLDTLVILHQAGARL |
| *Nomascus leucogenys* | LHGAEPNCADPATLTRPVHDAAREGFLDTLAVLHRAGARL |
| *Nyctereutes procyonoides* | LHGADPNCADPATLTRPVHDAAREGFLDTLVVLHRAGARL |
| *Ochotona curzoniae* | LHGAEPNYADPVTLTRPVHDAAREGFLDTLVALHRAGARL |
| *Ochotona princeps* | LHGAEPNYADPVTLTRPVHDAAREGFLDTLVALHRAGARL |
| *Octodon degus* | LHGADPNCADPVSLSRPVHDAAREGFLDTLVALHQAGARL |
| *Odobenus rosmarus divergens* | CHGAEPNCADPTTLTRPVHDAAREGFLDTLVVLHRAGARL |
| *Odocoileus virginianus texanus* | LHGADPNCADPATLTRPVHDAAREGFLDTLVALHRAGARL |
| *Onychomys torridus* | LYGAEPNREDPATLSRPVHDAAREGFLDTLVVLHQAGARL |
| *Orcinus orca* | LHGADPNCADPATLTRPVHDAAREGFLDTLVALHRAGARL |
| *Orycteropus afer afer* | LHGAEPNRADPTTLTRPVHDAAREGFLDTLVVLHRGGARL |
| *Oryctolagus cuniculus* | LHGAEPNCADPATLSRPVHDAAREGFLDTLVALHRAGARL |
| *Oryx dammah* | LHGADPNCADPATLTRPVHDAAREGFLDTLVALHRAGARL |
| *Otolemur garnettii* | LHGADPNRADPVTLTRPVHDAAREGFLDTLMALHQAGAQL |
| *Ovis aries* | LHGADPNCADPATLTRPVHDAAREGFLDTLVALHRAGARL |
| *Pan paniscus* | LHGAEPNCADPATLTRPVHDAAREGFLDTLVVLHRAGARL |
| *Pan troglodytes* | LHGAEPNCADPATLTRPVHDAAREGFLDTLVVLHRAGARL |
| *Panthera leo* | LHGADPNCADPATLTRPVHDAAREGFLDTLVVLHRAGARL |
| *Panthera pardus* | LHGADPNCADPATLTRPVHDAAREGFLDTLVVLHRAGARL |
| *Panthera tigris* | LHGADPNCADPATLTRPVHDAAREGFLDTLVVLHRAGARL |
| *Papio anubis* | LHGAEPNCADPATLTRPVHDAAREGFLDTLAVLHRAGARL |
| *Perognathus longimembris pacificus* | LHGAEANCADPATLTRPVHDAAREGFLDTLVTLHRAGARL |
| *Peromyscus leucopus* | LYGAEPTAXDPATLSRPVHDAAREGFLDTLVVLHQAGARL |
| *Peromyscus maniculatus bairdii* | LYGAEPNREDPATLSRPVHDAAREGFLDTLVVLHQAGARL |
| *Phoca vitulina* | CHGAEPNCADPTTLTRPVHDAAREGFLDTLVLLHRAGARL |
| *Phocoena sinus* | LHGADPNCADPATLTRPVHDAAREGFLDTLVALHRAGARL |
| *Phyllostomus discolor* | RHGADPNCADPATLATPVHDAAREGFLDTLVALHRAGAQL |
| *Phyllostomus hastatus* | RYGADPNCADPTTLATPVHDAAREGFLDTLVVLHRAGAQL |
| *Physeter catodon* | LHGADPNCADPATLTRPVHDAAREGFLDTLVALHRAGARL |
| *Physeter catodon* | LHGAEPNCADPATLTRPVHDAAREGFLDTLAVLHRAGAQL |
| *Pipistrellus kuhlii* | AHGADPNCADPATRTRPVHDAAREGFLDTLAALHRAGARL |
| *Pongo abelii* | LHGAEPNCADPATLTRPVHDAAREGFLDTLAVLHRAGARL |
| *Prionailurus bengalensis* | LHGADPNCADPATLTRPVHDAAREGFLDTLVVLHRAGARL |
| *Propithecus coquereli* | LHGAEPNRADPATLTRPVHDAAREGFLDTLVALHRAGARL |
| *Pteropus alecto* | LHGADPNCADPATLTRPVHDAAREGFLDTVVALHRAGARL |
| *Pteropus giganteus* | LHGADPNCADPATLTRPVHDAAREGFLDTVVALHRAGARL |
| *Pteropus vampyrus* | LHGADPNCADPATLTRPVHDAAREGFLDTVVALHRAGARL |
| *Puma concolor* | LHGADPNCADPATLTRPVHDAAREGFLDTLVVLHRAGARL |
| *Puma yagouaroundi* | LHGADPNCADPATLTRPVHDAAREGFLDTLVVLHRAGARL |
| *Rattus norvegicus* | SYGADSNCEDPTTLSRPVHDAAREGFLDTLVVLHQAGARL |
| *Rattus rattus* | SYGADSNCEDPTTLSRPVHDAAREGFLDTLAVLHEAGARL |
| *Rhinolophus ferrumequinum* | LHGADPNCADPATLTRPVHDAAREGFLDTLAVLHRAGARL |
| *Rhinolophus ferrumequinum* | LHGADPNCADPTTLTRPVHDAAREGFLDTLVVLHQAGARL |
| *Rhinopithecus bieti* | LHGAEPNCADPATLTRPVHDAAREGFLDTLAVLHRAGARL |
| *Rhinopithecus roxellana* | LHGAEPNCADPATLTRPVHDAAREGFLDTLAVLHRAGARL |
| *Rousettus aegyptiacus* | LHGADPNCADPVTLTRPVHDAAREGFLDTLVALHRAGARL |
| *Saimiri boliviensis boliviensis* | LHGAEPNCADPATLTRPVHDAAREGFLDTLVVLHRAGARL |
| *Sapajus apella* | LHGAEPNCADPATLTRPVHDAAREGFLDTLVVLHRAGAPL |
| *Sorex araneus* | LHGAEPNCADPVTLTRPVHDAAREGFLETLVALHRAGAQL |
| *Sousa chinensis* | LHGADPNCADPATLTRPVHDAAREGFLDTLVALHRAGARL |
| *Sturnira hondurensis* | LHGADPNRVDPATLATPVHDAAREGFLDTVVALHLAGARL |
| *Suricata suricatta* | LHGADPNCADPATLTRPVHDAAREGFLDTLVVLHRAGARL |
| *Sus scrofa* | LHGADPNCEDPATLTRPVHDAAREGFLDTLVVLHRAGARL |
| *Talpa occidentalis* | LHGADPNCADPATLTRPVHDAAREGFLDTLEALHQAGARL |
| *Theropithecus gelada* | LHGAEPNCADPATLTRPVHDAAREGFLDTLAVLHRAGARL |
| *Trachypithecus francoisi* | LHGAEPNCADPATLTRPVHDAARQGFLDTLAVLHRAGARL |
| *Trichechus manatus latirostris* | LHGAEPNCADPVTLTRPVHDAAREGFLDTLMVLHRAGARL |
| *Tursiops truncatus* | LHGADPNCADPATLTRPVHDAAREGFLDTLVALHRAGARL |
| *Urocitellus parryii* | LHGAEPNCADPATLTRPVHDAAREGFLDTLVALHLAGARL |
| *Ursus americanus* | LHGADPNCADPTTLTRPAHDAAREGFLDTLVVLHRAGARL |
| *Ursus arctos* | LHGADPNCADPTTLTRPAHDAAREGFLDTLVVLHRAGARL |
| *Ursus maritimus* | LHGADPNCADPTTLTRPAHDAAREGFLDTLVVLHRAGARL |
| *Vicugna pacos* | LHGADPNCADPATLTRPVHDAAREGFLDTLVALHRAGARL |
| *Vulpes lagopus* | LHGADPNCADPATLTRPVHDAAREGFLDTLVVLHRAGARL |
| *Vulpes vulpes* | LHGADPNCADPATLTRPVHDAAREGFLDTVVVLHRAGARL |
| *Zalophus californianus* | CTARSPTAPTLPTLTRPVHDAAREGFLDTLVVAAPSRRAA |

**<Marsupials>**

| | |
|---|---|
| *Dromiciops gliroides* | QHGAEPNTPDPTTLTLPVHDAAREGFLDTLMLLHRAGARL |
| *Gracilinanus agilis* | QYGAEPNTPDPTTLTLPVHDAAREGFLDTLMLLHRAGARL |
| *Monodelphis domestica* | QYGAEPNTPDPTTLTLPVHDAAREGFLDTLMLLHRAGARL |
| *Phascolarctos cinereus* | QHGAEPNTPDPTTLTLPVHDAAREGFLDTLMLLHRAGARL |
| *Sarcophilus harrisii* | QYGAEPNTPDPXTLTLPVHDVAREGFLDTLMLLHRAGARL |
| *Trichosurus vulpecula* | QHGADPNTPDPTTLALPVHDAAREGFLDTLMLLHRAGARL |
| *Vombatus ursinus* | QHGAEPNTPDLTTLTLPVHDAAREGFLDTLILLHRAGARL |

**<Monotremes>**

| | |
|---|---|
| *Ornithorhynchus anatinus* | QHGAEPNCPDPTTLTQPAHDAAREGFLDTLVLLHEAGARL |
| *Tachyglossus aculeatus* | QHGAEPNSPDPTTLTQPAHDAAREGFLDTLVLLHGAGARL |

**Figure S2a**

**Cdkn2a**

| | |
|---|---|
| *Homo sapiens* | LHGAEPNCADPATLTRPVHDAAREGFLDTLVVLHRAGARL |

**Birds**

| | |
|---|---|
| *Acanthisitta chloris* | QRGADPNRPDPRTGCLPAHDAARAGFLETLAALHRAGARL |
| *Accipiter nisus* | QRGADPNRPDPRTGSLPAHDAARAGFLETLAALHRAGARL |
| *Amazona aestiva* | QRGADPNRPDPRTGCFPAHDAARAGFLETLAVLHRAGARL |
| *Amazona collaria* | QRGADPNRPDPRTGCFPAHDAARAGFLETLAVLHRAGARL |
| *Anas platyrhynchos* | QRGADPNRPDPRTGCLPAHDAARAGFLDTLAALHRAGARL |
| *Anas zonorhyncha* | QRGADPNRPDPRTGCLPAHDAARAGFLDTLAALHRAGARL |
| *Anser brachyrhynchus* | QRGADPNRPDPRTGCLPAHDAARAGFLDTLAALHRAGARL |
| *Anser cygnoides* | QRGADPNRPDPRTGCLPAHDAARAGFLDTLAALHRAGARL |
| *Aptenodytes patagonicus* | QRGADPNRPDPRTGCLPAHDAARAGFLETLAAL------- |
| *Apteryx haastii* | QRGADPNRADPRTGCFPAHDAARAGFLDTLAALHRAG--- |
| *Apteryx owenii* | QRGADPNRADPRTGCFPAHDAARAGFLDTLAALHRA---- |
| *Aquila chrysaetos chrysaetos* | QRGADPNRPDPRTGSLPAHDAARAGFLETLAALHRAGARL |
| *Aythya fuligula* | QRGADPNRPDPRTGCLPAHDAARAGFLDTLAALHRAGARL |
| *Balearica regulorum gibbericeps* | QRGADPNRPDPRTGCFPAHDAARAGFLETLAALHRAGARL |
| *Cairina moschata domestica* | QRGADPNRPDPRTGCLPAHDAARAGFLDTLAALHRAGARL |
| *Calidris pugnax* | QRGADPNRPDPRTGCFPAHDAARAGFLETLAALHRAGARL |
| *Calidris pygmaea* | QRGADPNRPDPRTGCFPAHDAARAGFLETLAALHRAGARL |
| *Calypte anna* | QRGADPNRPDPRTGCLPVHDAARTGFLETLAVLHRAGARL |
| *Camarhynchus parvulus* | RHGADPSRPDPRTGCLPVHDAARAGFVETLAALHRAGARL |
| *Cariama cristata* | QRGADPNRPDPRTGCLPAHDAARAGFLETLAALHRAGARL |
| *Catharus ustulatus* | RRGADPNRPDPHTGCLPVHDAARAGFLETLAALHRAGARL |
| *Chiroxiphia lanceolata* | QRGADPNRPDPSTGCLPAHDAARAGFLETLAALHRAGARL |
| *Chlamydotis macqueenii* | QRGADPNRPDPRTGCLPAHDAARAGFLETLVALHRAGARL |
| *Corapipo altera* | QRGADPNRPDPSTGCLPAHDAARAGFLETLAALHRAGARL |
| *Corvus cornix cornix* | RRGADPNRPDPRTGCLPAHDAARAGFLETLAALHRAGARL |
| *Corvus hawaiiensis* | RRGADPNRPDPRTGCLPAHDAARAGFLETLAALHRAGARL |
| *Corvus kubaryi* | RRGADPNRPDPRTGCLPAHDAARAGFLETLAALHRAGARL |
| *Corvus moneduloides* | RRGADPNRPDPRTGCLPAHDAARAGFLETLAALHRAGARL |
| *Cuculus canorus* | QRGADPNRPDPLTGCFPAHDAARAGFLETLAALHRAGARL |
| *Cyanoderma ruficeps* | RRGADPNRPDPRTGCLPAHDAARAGFLDTLVALHRAGARL |
| *Cygnus atratus* | QRGADPNRPDPRTGCFPAHDAARAGFLDTLAALHRAGARL |
| *Dromaius novaehollandiae* | QRGADPNRADPRTGCFPAHDAARAGFLDTLAALHRAGARL |
| *Dryobates pubescens* | QFGADPNRPDPSTGCFPVHDAARSGFLETVAVLHRAGARF |
| *Empidonax traillii* | QRGADPNRPDPSTGCLPAHDAARAGFLETLVALHRAGARL |
| *Erythrura gouldiae* | RHGADPSRPDPRTGCLPAHDAARAGFLETLAALHRAGARL |
| *Eudyptes chrysocome* | QRGADPNRPDPRTGCLPAHDAARAGFLETLAALHRAGARL |
| *Eudyptes filholi* | QRGADPNRPDPRTGCLPAHDAARAGFLETLAALHRAGARL |
| *Eudyptes moseleyi* | QRGADPNRPDPRTGCLPAHDAARAGFLETLAALHRAGARL |
| *Eudyptes pachyrhynchus* | QRGADPNRPDPRTGCLPAHDAARAGFLEQLA--------- |
| *Eudyptes robustus* | QRGADPNRPDPRTGCLPAHDAARAGFLETLAALHRAGX-- |
| *Eudyptes sclateri* | QRGADPNRPDPRTGCLPAHDAARAGFLETLAALHRAGARL |
| *Eudyptula novaehollandiae* | QRGADPNRPDPRTGCLPAHDAARAGFLETLAALHRAGARL |
| *Falco cherrug* | QRGADPNRPDPSTGCFPVHDAASAGFLETLEALHRAGARL |
| *Falco naumanni* | QRGADPNRPDPSTGCFPVHDAASAGFLETLEALHRAGARL |
| *Falco peregrinus* | QRGADPNRPDPSTGCFPVHDAASAGFLETLEALHRAGARL |
| *Falco rusticolus* | QRGADPNRPDPSTGCFPVHDAASAGFLETLEALHRAGARL |
| *Falco tinnunculus* | QRGADPNRPDPSTGCFPVHDAASAGFLETLEALHRAGARL |
| *Ficedula albicollis* | RRGADPNRPDPRTGCRPAHDAARAGFLETLAALHRAAHPG |
| *Gallus gallus* | QRGADPNRPDPRTGCRPAHDAARAGFLDTLAALHRAGARL |
| *Haliaeetus albicilla* | QRGADPNRPDPRTGSLPAHDAARAGFLETLAALHRAGARL |
| *Hirundo rustica* | RRGADPNRPDPRTGCFPVHDAARAGFLETLAALHRAGARL |
| *Hypotaenidia okinawae* | QRGADPNRPDPRTGCLPAHDAARAGFLETLAALHRAGARL |
| *Junco hyemalis* | LHGADPNRPDPRTGCLPVHDAARAGFLGTLAALHRAGALL |
| *Lagopus leucura* | QRGADPSRPDPRTGCRPAHDAARAGFLDALEDLHRAGARL |
| *Lepidothrix coronata* | QRGADPNRPDPSTGCLPAHDAARAGFLETLAALHRAGARL |
| *Lonchura striata domestica* | RHGADPNRPDPRTGCLPAHDAARAGFLETLAALHRAGARL |
| *Melopsittacus undulatus* | QRGADPNRPDPRTGCLPAHDAARAGFLETLAVLHRAGARL |
| *Merops nubicus* | QRGADPNRPDPRTGCLPAHDAARAGFLETLAALHRAGARL |
| *Molothrus ater* | RHGADPNRPDPRTGCLPVHDAARAGFLETLAALHRAGARL |
| *Motacilla alba alba* | RHGADPNRPDPRTGCLPVHDAARAGFLGTLAALHRAGARL |
| *Neopelma chrysocephalum* | QRGADPNRPDPSTGCLPAHDAARAGFLETLVALHRAGARL |
| *Nothoprocta perdicaria* | QRGADPNRADPRTGCYPAHDAARAGFVDTLAALHRAGARL |
| *Onychostruthus taczanowskii* | RHGADPNRPDPRTGCLPAHDAARAGFLETLAALHRAGARL |
| *Parus major* | RRGADPNRPDPRTGCLPVHDAARAGFLETLAVLHRAGAHL |
| *Passer montanus* | RHGADPNRADPRTGCLPAHDAARAGFLETLAALHRAGARL |
| *Pavo cristatus* | QRGADPNRPDPRTGCRPAHDAARAGFLDTLAALHRAGARL |
| *Pipra filicauda* | QRGADPNRPDPSTGCLPAHDAARAGFLETLAALHRAGARL |
| *Pseudopodoces humilis* | RRGADPNRPDPRTGCLPVHDAARAGFLETLAVLHRAGARL |
| *Pterocles gutturalis* | QRGADPNRPDPLTGCLPAHDAARAGFLETLAALHRAGARL |
| *Pyrgilauda ruficollis* | RHGADPNRPDPSTGCLPAHDAARAGFLETLAALHRAGARL |
| *Serinus canaria* | RHGADPNHPDPRTGCFPVHDAARAGFLETLATLHRAGARL |
| *Spheniscus demersus* | QRGADPNRPDPRTGCLPAHDAARAGFLETLAALHRAGARL |
| *Spheniscus humboldti* | QRGADPNRPDPRTGCLPAHDAARAGFLETLAA-------- |
| *Strigops habroptila* | QRGADPNRPDPRTGCLPAHDAARAGFLETLAALHRAGARL |
| *Strix occidentalis caurina* | QRGADPNRPDPRTGCLPAHDAARAGFLETLAALHRAGARL |
| *Sturnus vulgaris* | RRGADPNRPDPRTGCLPAHDAARAGFLETLAALHRAGARL |
| *Taeniopygia guttata* | RHGADPNRPDPRTGCLPAHDAARSGFLETLAALHRAGARL |
| *Tauraco erythrolophus* | ERGADPNRPDPRTGCLPAHDAARTGFLETLAALHRAGARL |
| *Turdus rufiventris* | RRGADPNRPDPHTGCLPVHDAARAGFLETLAVLHRAGARL |
| *Turnix velox* | QHGADPNRPDPRTGCLPVHDAARAGFLDTLQTLHRAGARL |
| *Tyto alba* | QRGADPNRPDPRTGCLPAHDAARAGFLETLAALHRAGARL |
| *Zonotrichia albicollis* | LHGADPNRPDPRTGCLPVHDAARAGFLGTQAALHRAGARL |
| *Zosterops borbonicus* | RRGADPNRPDPRTGCLPAHDAARAGFLETLRALHRAGARL |

**Reptiles**

| | |
|---|---|
| *Chelonia mydas* | QRGADPNRPDPRTGSLPVHDAAREGFLDTLVALHRGGARL |
| *Chelonoidis abingdonii* | ERGADPNRPDPRTGSLPVHDAAREGFLDTLVALHRGGARL |
| *Chelydra serpentina* | QRGADPNRPDPRTGSLPVHDAARGGFLDTLVALHRGGARL |
| *Chrysemys picta bellii* | QRGADPNRPDPRTGSLPVHDAAREGFLDTLVALHRGGARL |
| *Crocodylus porosus* | RSGADPNRPDPATGSCPAHDAAREGFLDTLAALHRAGARL |
| *Crotalus adamanteus* | ERGANPNLPDPSTGSRPVHDAAREGFLDTLQVLISGGARL |
| *Crotalus tigris* | ERGANPNLPDPSTGSRPVHDAAREGFLDTLQVLISGGARL |
| *Dermochelys coriacea* | QRGADPNRPDPLTGSLPVHDAAREGFLDTLVALHRGGARL |
| *Gavialis gangeticus* | RSGADPNRPDPATGSRPAHDAAREGFLDTLVALHRAGARL |
| *Gekko japonicus* | RKGADPNRPDPTTGALPSHDAARQGFLDTLLVLREGGARL |
| *Gopherus agassizii* | QRGADPNRPDPRTGSLPVHDAAREGFLDTLVALHRGGARL |
| *Gopherus evgoodei* | QRGADPNRPDPRTGSLPVHDAAREGFLDTLVALHRGGARL |
| *Lacerta agilis* | ERGADPNVPDPSTGSYPAHDAAHQGFLDTLRMLCSGGARV |
| *Laticauda laticaudata* | ERGANPNVPDPSTGSLPVHDAAREGFLDTLQVLVSGGARL |
| *Mauremys mutica* | QRGADPNRPDPRTGSLPVHDAAREGFLDTLVALHRGGARL |
| *Mauremys reevesii* | QRGADPNRPDPRTGSLPVHDAAREGFLNTLVALHRGGARL |
| *Naja naja* | ERGADPNVPDPSTGSLPVHDAAREGFLDTLQVLVSGGARL |
| *Ophiophagus hannah* | ERGANPNVPDPSTGSLPVHDAAREGFLDTLQVLVSGGARL |
| *Pantherophis guttatus* | ERKANPNVPDPSTGSLPVHDAAREGFLDTLQVLISGGARL |
| *Pelodiscus sinensis* | QSGADPNRPDPSTGSLPVHDAARGGFLDTLVALHRGGARL |
| *Pelusios castaneus* | QRGADPDRPDPSTGSLPVHDAAREGFLDTLVALHRGGARL |
| *Podarcis muralis* | RWGADPNVPDPSTGSCPAHDAAREGFLDTLRVLRSGGARL |
| *Pogona vitticeps* | QRGADPNLRDPTTGSFPAHDAARGGFLDTLRVLCQWGARF |
| *Pseudonaja textilis* | SGGARLDLRN-CYGRLPLDEAAERGQNQVVHFLAR----- |
| *Salvator merianae* | QHGADPNLPDPSTETLPVHDAAREGFLDTLRILRLWGARF |
| *Sceloporus undulatus* | EAGADPNLPDRATGSLPAHDAAGQGFLETLQVLHRGGARL |
| *Sphenodon punctatus* | RAGADPNIPDPATGSFPAHDAAREGFLDTLEVLHLGGARF |
| *Terrapene carolina triunguis* 1 | QRGADPNRPDPRTGSLPVHDAARYGFLDTLVALHRGGARL |
| *Terrapene carolina triunguis* 2 | QRGADPNRPDPRTGSLPVHDAAREGFLDTLVALHRGGARL |
| *Thamnophis elegans* | ERGANPNVPDRSTGSLPVHDAAREGFLDTLQVLIAGGARL |
| *Trachemys scripta elegans* | QRGADPNRPDPRTGSLPVHDAAREGFLDTLVALHRGGARL |
| *Varanus komodoensis* | QRGADPNRPDPSTGCLPAHDAAREGFLDTLRVLHGGGARF |
| *Zootoca vivipara* | ERGADPNMPDPSTGSCPAHDAAREGFLDTLRVLRSGGARF |

**Amphibians**

| | |
|---|---|
| *Geotrypetes seraphini* | RYGADPNVPDPSTGTYPAHDAAREGFLNTLQVLVMGGARL |
| *Microcaecilia unicolor* | RYGADPNVPDPTTGTCPAHDAAREGFLDTLQVLVTGGARL |
| *Rhinatrema bivittatum* | RYGADPNLPDPTTATCPAHDAAREGFLDTLRVLLRAGASL |

**Bony fish**

| | |
|---|---|
| *Ameiurus melas* | LHGANPNVVDVSTGAMPLHDAAREGFLLTVQLLVRAGADL |
| *Coryphaenoides rupestris* | SRGAGPRVADPSTGSTPLHDAARAGFPDTVRALLESSADP |
| *Gouania willdenowi* | KYGADPNVFDPTTGATPLHDAAQTGFLDTVRLLVQFQGNP |
| *Larimichthys crocea* | KHGADPNLADCSTGTTPLHDAARTGFLETVQLLVQHMADP |
| *Protopterus annectens* | AHGANPNIKD-SYDTSPVHDAARAGFLESVQLLIKYGAVA |
| *Siniperca chuatsi* | RRGADPNVKDRSTGTTPLHDAARTGFLDTARLLVQYKADP |

### Figure S2b

**Cdkn2a**

| | |
|---|---|
| *Homo sapiens* | LHGAEPNCADPATLTRPVHDAAREGFLDTLVVLHRAGARL |

**Cdkn2b**

**Mammals**

**<Placentarians>**

| | |
|---|---|
| *Acinonyx jubatus* | LHGADPNCADPATLTRPVHDAAREGFLDTLVVLHRAGARL |
| *Ailuropoda melanoleuca* | LHGADPNCADPTTLTRPAHDAAREGFLDTLVVLHRAGARL |
| *Aotus nancymaae* | LHGAEPNCADPATLTRPVHDAAREGFLDTLVMLHRAGARL |
| *Artibeus jamaicensis* | LHGADPNCADPTTLTRPVHDAAREGFLDTLVALHRAGAHL |
| *Arvicanthis niloticus* | LHGAEPNCADPATLTRPVHDAAREGFLDTLVVLHRAGARL |
| *Arvicola amphibius* | LHGAEPNCADPTTLTRPVHDAAREGFLDTLVMLHKAGAWL |
| *Balaenoptera acutorostrata scammoni* | LHGADPNCADPATLTRPVHDAAREGFLDTLAALHRAGARL |
| *Balaenoptera musculus* | LHGADPNCADPATLTRPVHDAAREGFLDTLAALHRAGARL |
| *Bison bison bison* | -------------LTRPVHDAAREGFLDTLVALHRAGGRL |
| *Bos indicus* | LHGADPNCADPATLTRPVHDAAREGFLDTLVALHRAGGRL |
| *Bos mutus* | LHGADPNCADPATLTRPVHDAAREGFLDTLVALHRAGGRL |
| *Bos taurus* | LHGADPNCADPATLTRPVHDAAREGFLDTLVALHRAGGRL |
| *Bubalus bubalis* | LHGADPNCADPATLTRPVHDAAREGFLDTLVALHRAGGRL |
| *Callithrix jacchus* | LHGAEPNCADPATLTRPVHDAAREGFLDTLVMLHRAGARL |
| *Callorhinus ursinus* | CHGAEPNCADPTTLTRPVHDAAREGFLDTLVVLHRAGARL |
| *Camelus bactrianus* | LHGADPNCADPATLTRPVHDAAREGFLDTLVALHRAGARL |
| *Camelus dromedarius* | LHGADPNCADPATLTRPVHDAAREGFLDTLVALHRAGARL |
| *Camelus ferus* | LHGADPNCADPATLTRPVHDAAREGFLDTLVALHRAGARL |
| *Canis lupus familiaris* | LHGADPNCADPATLTRPVHDAAREGFLDTLVVLHRAGARL |
| *Capra hircus* | LHGADPNCADPATLTRPVHDAAREGFLDTLVALHRAGGRL |
| *Carlito syrichta* | LHGAEPNCADPATLTRPVHDAAREGFLDTLVALHRAGARL |
| *Castor canadensis* | LYGAEPNCADSATLTRPVHDAAREGFLDTLVALHGAGARL |
| *Cavia porcellus* | LRGAEPNCADPATLTRPVHDAAREGFLDTLVALHRAGAQL |
| *Cebus imitator* | LHGAEPNCADPATLTRPVHDAAREGFLDTLVMLHRAGARL |
| *Ceratotherium simum simum* | LHGADPNRADPATLTRPVHDAAREGFLDTLVALHRAGARL |
| *Cercocebus atys* | LHGAEPNCADPATLTRPVHDAAREGFLDTLVVLHRAGARL |
| *Cervus canadensis* | LHGADPNCADPATLTRPVHDAAREGFLDTLVALHRAGGRL |
| *Cervus elaphus* | LHGADPNCADPATLTRPVHDAAREGFLDTLVALHRAGGRL |
| *Cervus hanglu yarkandensis* | LHGADPNCADPATLTRPVHDAAREGFLDTLVALHRAGGRL |
| *Chinchilla lanigera* | LRGAEPNCADPATLTRPVHDAAREGFMDTLVALHRAGAQL |
| *Chlorocebus sabaeus* | LHGAEPNCADPATLTRPVHDAAREGFLDTLVVLHRAGARL |
| *Choloepus didactylus* | LHGAEPNCADPVTLTRPVHDAAREGFLDTLLALHRAGARL |
| *Chrysochloris asiatica* | LHGAEPDCMDPATLTRPVHDAAREGFLDTLVLLHQAGAQL |
| *Colobus angolensis palliatus* | LHGAEPNCADPATLTRPVHDAAREGFLDTLVVLHRAGARL |
| *Condylura cristata* | LHGADPNCADPHTLTRPVHDAAREGFLDTLVALHQAGARL |
| *Cricetulus griseus* | LHGAEPNCADPNTLTRPVHDAAREGFLDTLVVLHRAGARL |
| *Crocuta crocuta* | LHGADPNCADPATLTRPVHDAAREGFLDTLVVLHRAGARL |
| *Dasypus novemcinctus* | LHGADPNCADPATFTRPVHDAAREGFLDTVLELHRAGAQL |
| *Delphinapterus leucas* | LHGADPNCADPATLTRPVHDAAREGFLDTLVALHRAGARL |
| *Desmodus rotundus* | LHGADPNCADPATLNRPVHDAAREGFLDTLVALHRAGARL |
| *Dipodomys ordii* | LHGAEANCADPTTLTRPVHDAAREGFLDTLVMLHRAGARL |
| *Dipodomys spectabilis* | LHGAEANCADPATLTRPVHDAAREGFLDTLVMLHRAGARL |
| *Echinops telfairi* | CHGAEPNCADPTTLARPVHDAAREGFLDTLVLLHRAGAQL |
| *Elephantulus edwardii* | LHGADPNFEDPVTRTWPVHDAAREGFLDTLAVLHRNGARL |
| *Enhydra lutris kenyoni* | LYGAEPNCADEATLTRPVHDAAREGFLDTLVVLHRAGAQL |
| *Eptesicus fuscus* | THGADPNRPDPATLTRPVHDAAREGFLDTLVALHRAGARL |
| *Equus asinus* | LHGADPNRADPDTLTRPVHDAAREGFLDTLVALHRAGARL |
| *Equus caballus* | LHGADPNRADPDTLTRPVHDAAREGFLDTLVALHRAGARL |
| *Equus przewalskii* | LHGADPNRADPDTLTRPVHDAAREGFLDTLVALHRAGARL |
| *Equus quagga* | LHGADPNRADPDTLTRPVHDAAREGFLDTLVALHRAGARL |
| *Erinaceus europaeus* | LHGAEPNCADPATLTQPVHDAAREGFLDTVMALHRAGARL |
| *Eumetopias jubatus* | CHGAEPNCADPTTLTRPVHDAAREGFLDTLVVLHRAGARL |
| *Felis catus* | LHGADPNCADPATLTRPVHDAAREGFLDTLVVLHRAGARL |
| *Fukomys damarensis* | LHGAEPNCADPATLTRPVHDAAREGFMDTLVALHRAGARL |
| *Galemys pyrenaicus* | LYGADPNCADPDTLTRPVHDAAREGFLDTLVVLHQAGARL |
| *Galeopterus variegatus* | LHGAEPNCADPATLTRPVHDAAREGFLETLVALRRAGARL |
| *Globicephala melas* | LHGADPNCADPATLTRPVHDAAREGFLDTLVTLHRAGARL |
| *Gorilla gorilla gorilla* | LHGAEPNCADPATLTRPVHDAAREGFLDTLVVLHRAGARL |
| *Grammomys surdaster* | LHGAEPNCADPATLTRPVHDAAREGFLDTLVVLHRAGARL |
| *Gulo gulo luscus* | LHGADPNCADPITLTRPVHDAAREGFLDTLLVLHRAGARL |
| *Halichoerus grypus* | CHGAEPNCADPTTLTRPVHDAAREGFLDTLVLLHRAGARL |
| *Heterocephalus glaber* | LHGAEPNCADPATLTRPVHDAAREGFIDTLVALHRAGARL |
| *Hipposideros armiger* | LHGADPNCADPATLTRPVHDAAWEGFLDTLAVLHRAGARL |
| *Homo sapiens* | LHGAEPNCADPATLTRPVHDAAREGFLDTLVVLHRAGARL |
| *Hyaena hyaena* | LHGADPNCADPATLTRPVHDAAREGFLDTLVVLHRAGARL |
| *Hylobates moloch* | LHGAEPNCADPATLTRPVHDAAREGFLDTLVVLHRAGARL |
| *Ictidomys tridecemlineatus* | LHGAEPNCADPATLTRPVHDAAREGFLDTLVALHRAGARL |
| *Jaculus jaculus* | LHGAEPNCADPATLTRPVHDAAREGFLDTLAALHRAGARL |
| *Lagenorhynchus obliquidens* | LHGADPNCADPATLTRPVHDAAREGFLDTLVTLHRAGARL |
| *Lemur catta* | LHGAEPNCADPATLTRPVHDAAREGFLDTLVALHRAGARL |
| *Leopardus geoffroyi* | LHGADPNCADPATLTRPVHDAAREGFLDTLVVLHRAGARL |
| *Leptonychotes weddellii* | CHGAEPNCADPTTLTRPVHDAARDGFLDTLVVLHRAGARL |
| *Lipotes vexillifer* | LHGADPNCADPATLTRPVHDAAREGFLDTLVALHRAGARL |
| *Lontra canadensis* | LYGAEPNCADEATLTRPVHDAAREGFLDTLVVLHRAGAQL |
| *Loxodonta africana* | LHGAEPNCADPATLTRPVHDAAREGFLDTLMVLQRAGARL |
| *Lynx canadensis* | LHGADPNCADPATLTRPVHDAAREGFLDTLVVLHRAGARL |
| *Lynx pardinus* | LHGADPNCADPATLTRPVHDAAREGFLDTLVVLHRAGARL |
| *Macaca fascicularis* | LHGAEPNCADPATLTRPVHDAAREGFLDTLVVLHRAGARL |
| *Macaca mulatta* | LHGAEPNCADPATLTRPVHDAAREGFLDTLVVLHRAGARL |
| *Macaca nemestrina* | LHGAEPNCADPATLTRPVHDAAREGFLDTLVVLHRAGARL |
| *Mandrillus leucophaeus* | LYGAEPNCADPATLTRPVHDAAREGFLDTLVVLHRAGARL |
| *Manis javanica* | LHGADPNCTDPATLTRPVHDAAREGFLDTLVALHRAGARL |
| *Manis pentadactyla* | LHGADPNCTDPATLTQPVHDAAREGFLDTLVALHRAGARL |
| *Marmota flaviventris* | LHGAEPNCADPATLTRPVHDAAREGFLDTLVALHRAGARL |
| *Marmota marmota marmota* | LHGAEPNCADPATLTRPVHDAAREGFLDTLVALHRAGARL |
| *Marmota monax* | LHGAEPNCADPATLTRPVHDAAREGFLDTLVALHRAGARL |
| *Mastomys coucha* | LHGAEPNCADPATLTRPVHDAAREGFLDTLVVLHRAGARL |
| *Meles meles* | LHGAEPNCADEATLTRPVHDAAREGFLDTLVVLHRAGAQL |
| *Meriones unguiculatus* | LHGAEPNCADPATLTRPVHDAAREGFLDTLVVLHRAGARL |
| *Mesocricetus auratus* | LHGAEPNCADPNTLTRPVHDAAREGFLDTLVVLHRAGARL |
| *Microcebus murinus* | LHGAEPNCADPATLTRPVHDAAREGFLDTLGALHRAGARL |
| *Microtus ochrogaster* | LHGAEPNCADPATLTRPVHDAAREGFLDTLVMLHKAGARL |
| *Microtus oregoni* | LHGAEPNCADPATLTRPVHDAAREGFLDTLVMLHKAGAQL |
| *Mirounga angustirostris* | CHGAEPNCADPTTLTRPVHDAARDGFLDTLVVLHRAGARL |
| *Mirounga leonina* | CHGAEPNCADPTTLTRPVHDAARDGFLDTLVVLHRAGARL |
| *Molossus molossus* | LHGADPNCADPATLTRPVHDAAREGFLDTLVVLHQAGARL |
| *Monodon monoceros* | LHGADPNCADPATLTRPVHDAAREGFLDTLVALHRAGARL |
| *Mus caroli* | LHGAEPNCADPATLTRPVHDAAREGFLDTLVVLHRAGARL |
| *Mus musculus* | LHGAEPNCADPATLTRPVHDAAREGFLDTLVVLHRAGARL |
| *Mus pahari* | LHGAEPNCADPATLTRPVHDAAREGFLDTLVVLHRAGARL |
| *Mustela erminea* | LYGAEPNCADEATLTRPVHDAAREGFLDTLMVLHRAGAQL |
| *Mustela putorius furo* | LYGAEPNCADEATLTRPVHDAAREGFLDTLVVLHRAGAQL |
| *Myotis davidii* | IHGADPNCADPATLTRPVHDAAREGFLDTLVALHRAGARL |
| *Myotis lucifugus* | IHGADPNCADPATLTRPVHDAAREGFLDTLVALHRAGAQL |
| *Myotis myotis* | IHGADPNCADPATLTRPVHDAAREGFLDTLVALHRAGARL |
| *Nannospalax galili* | LHGAEPNCTDPATLTRPVHDAAREGFLDTLVALHQAGARL |
| *Neogale vison* | LYGAEPNCADPVTLTRPVHDAAREGFLDTLVVLHRAGAQL |
| *Neomonachus schauinslandi* | CHGAEPNCADPTTLTRPVHDAARDGFLDTLVVLHRAGARL |
| *Neophocaena asiaeorientalis asiaeorientalis* | LHGADPNCADPATLTRPVHDAAREGFLDTLVALHRAGARL |
| *Neosciurus carolinensis* | FHGAEPNCADPATLTRPVHDAAREGFLDTLVALHRAGARL |
| *Neotoma lepida* | LHGAEPNCADPATLTRPVHDAAREGFLDTLVVLHRAGARL |
| *Nomascus leucogenys* | LHGAEPNCADPATLTRPVHDAAREGFLDTLVVLHRAGARL |
| *Nyctereutes procyonoides* | LHGADPNCADPATLTRPVHDAAREGFLDTLVVLHRAGARL |
| *Ochotona curzoniae* | LHGAEPNYADPVTLTRPVHDAAREGFLDTLVALHRAGARL |
| *Ochotona princeps* | LHGAEPNYADPVTLTRPVHDAAREGFLDTLVALHRAGARL |
| *Octodon degus* | LHGAEPNCADPATLTRPIHDAAREGFMDTLVALHRAGAQL |
| *Odobenus rosmarus divergens* | LHGADPNCADPATLTRPVHDAAREGFLDTLVALHRAGARL |
| *Odocoileus virginianus texanus* | LHGADPNCADPATLTRPVHDAAREGFLDTLVALHRAGARL |
| *Onychomys torridus* | LHGAEPNCADPATLTRPVHDAAREGFLDTLVVLHRAGARL |
| *Orcinus orca* | LHGADPNCADPATLTRPVHDAAREGFLDTLVALHRAGARL |
| *Orycteropus afer afer* | LFGAEPNCADPATLTQPVHDAAREGFLDTLVVLHRAGARL |
| *Oryctolagus cuniculus* | LHGAEPNCADPATLSRPVHDAAREGFLDTLVALHRAGARL |
| *Oryx dammah* | LHGADPNCADPATLTRPVHDAAREGFLDTLVALHRAGARL |
| *Otolemur garnettii* | LHGAEPNCADPTTLTRPVHDAAREGFLDTLVVLHRAGARL |
| *Ovis aries* | LHGADPNCADPATLTRPVHDAAREGFLDTLVALHRAGGRL |
| *Pan paniscus* | LHGAEPNCADPATLTRPVHDAAREGFLDTLVVLHRAGARL |
| *Pan troglodytes* | LHGAEPNCADPATLTRPVHDAAREGFLDTLVVLHRAGARL |
| *Panthera leo* | LHGADPNCADPATLTRPVHDAAREGFLDTLVVLHRAGARL |
| *Panthera pardus* | LHGADPNCADPATLTRPVHDAAREGFLDTLVVLHRAGARL |
| *Panthera tigris* | LHGADPNCADPATLTRPVHDAAREGFLDTLVVLHRAGARL |
| *Papio anubis* | LHGAEPNCADPATLTRPVHDAAREGFLDTLVVLHRAGARL |
| *Perognathus longimembris pacificus* | LHGAEANCADPATLTRPVHDAAREGFLDTLVTLHRAGARL |
| *Peromyscus leucopus* | LHGAEPNCADPATLTRPVHDAAREGFLDTLVALHRAGARL |
| *Peromyscus maniculatus bairdii* | LHGAEPNCADPATLTRPVHDAAREGFLDTLVALHRAGARL |
| *Phoca vitulina* | CHGAEPNCADPTTLTRPVHDAAREGFLDTLVLLHRAGARL |
| *Phocoena sinus* | LHGADPNCADPATLTRPVHDAAREGFLDTLVALHRAGARL |
| *Phyllostomus discolor* | LHGADPNCADPITLTRPVHDAAREGFLDTLVALHRAGARL |
| *Phyllostomus hastatus* | LHGADPNCADPITLTRPVHDAAREGFLDTLVALHRAGARL |
| *Physeter catodon* | LHGADPNCADPATLTRPVHDAAREGFLDTLVALHRAGARL |
| *Piliocolobus tephrosceles* | LHGAEPNCADPATLTRPVHDAAREGFLDTLVVLHRAGARL |
| *Pipistrellus kuhlii* | AHGADPNCADPATCTRPVHDAAREGFLDTLAALHRAGARL |
| *Pongo abelii* | LHGAEPNCADPATLTRPVHDAAREGFLDTLVVLHRAGARL |
| *Prionailurus bengalensis* | LHGADPNCADPATLTRPVHDAAREGFLDTLVVLHRAGARL |
| *Propithecus coquereli* | LHGAEPNCADPATLTRPVHDAAREGFLDTLVALHRAGARL |
| *Pteropus alecto* | LHGADPNCADPATLTRPVHDAAREGFLDTVVALHRAGARL |
| *Pteropus giganteus* | LHGADPNCADPATLTRPVHDAAREGFLDTVVALHRAGARL |
| *Pteropus vampyrus* | LHGADPNCADPATLTRPVHDAAREGFLDTVVALHRAGARL |
| *Puma concolor* | LHGADPNCADPATLTRPVHDAAREGFLDTLVVLHRAGARL |
| *Puma yagouaroundi* | LHGADPNCADPATLTRPVHDAAREGFLDTLVVLHRAGARL |
| *Rattus norvegicus* | LHGAEPNCADPATLTRPVHDAAREGFLDTLMVLHKAGARL |
| *Rattus rattus* | LHGAEPNCADPATLTRPVHDAAREGFLDTLVVLHKAGARL |
| *Rhinolophus ferrumequinum* | LHGADPNCADPTTLARPVHDAAWEGFLDTLVVLHRAGARL |
| *Rhinolophus sinicus* | LHGADPNCADPTTLARPVHDAAWEGFLDTLVVLHRAGARL |
| *Rhinopithecus roxellana* | LHGAEPNCADPATLTRPVHDAAREGFLDTLVVLHRAGARL |
| *Rousettus aegyptiacus* | LHGADPNCADSATLTRPVHDAAREGFLDTLVALHRAGARL |
| *Saimiri boliviensis boliviensis* | LHGAEPNCADPATLTRPVHDAAREGFLDTLVMLHRAGARL |
| *Sapajus apella* | LHGAEPNCADPATLTRPVHDAAREGFLDTLVMLHRAGARL |
| *Sorex araneus* | LHGAEPNCADPATLTRPVHDAAREGFLDTLVALHRAGAQL |
| *Sousa chinensis* | LHGADPNCADPATLTRPVHDAAREGFLDTLVALHRAGARL |
| *Sturnira hondurensis* | LHGADPNCADPATLTRPVHDAAREGFLDTLVALHRAGARL |
| *Suricata suricatta* | LHGADPNCADPATLTRPVHDAAREGFLDTLVVLHRAGARL |
| *Sus scrofa* | LHGADPNCADPATLTRPVHDAAREGFLDTLVALRRAGARL |
| *Talpa occidentalis* | LHGADPNCADPATLTRPVHDAAREGFLDTLVALRQAGARL |
| *Theropithecus gelada* | LHGAEPNCADPATLTRPVHDAAREGFLDTLVVLHRAGARL |
| *Trachypithecus francoisi* | LHGAEPNCADPATLTRPVHDAAREGFLDTLMVLHRAGARL |
| *Trichechus manatus latirostris* | LHGAEPNCADPVTLTRPVHDAAREGFLDMLMVLHRAGARL |
| *Tupaia chinensis* | LHGAEPNCADPATLTQPVHDAAREGFLDTRVALHRAGARL |
| *Tursiops truncatus* | LHGADPNCADPATLTRPVHDAAREGFLDTLVALHRAGARL |
| *Urocitellus parryii* | LHGAEPNCADPATLTRPVHDAAREGFLDTLVALHRAGARL |
| *Ursus americanus* | LHGAEPNCADEATLTRPVHDAAREGFLDTLVVLHRAGARL |
| *Ursus arctos* | LHGAEPNCADEATLTRPVHDAAREGFLDTLVVLHRAGARL |
| *Ursus maritimus* | LHGAEPNCADEATLTRPVHDAAREGFLDTLVVLHRAGARL |
| *Vicugna pacos* | LHGADPNCADPATLTRPVHDAAREGFLDTLVALHRAGARL |
| *Vulpes lagopus* | LHGADPNCADPATLTRPVHDAAREGFLDTLVVLHRAGARL |
| *Vulpes vulpes* | LHGADPNCADPATLTRPVHDAAREGFLDTVVVLHRAGARL |
| *Zalophus californianus* | CHGAEPNCADPTTLTRPVHDAAREGFLDTLVVLHRAGARL |

**<Marsupials>**

| | |
|---|---|
| *Gracilinanus agilis* | KHGADPNIPDPTTLTLPVHDAAREGFLDTLILLHRAGARL |
| *Monodelphis domestica* | KHGADPNIPDPTTLTLPVHDAAREGFLDTLILLHRAGARL |
| *Phascolarctos cinereus* | KHGADPNVPDPTTLTLPVHDAAREGFLDTLILLHRAGARL |
| *Trichosurus vulpecula* | KHGADPNIPDPTTLTLPVHDAAREGFLDTLILLHRAGARL |
| *Vombatus ursinus* | KHGADPNVPDPTTLTLPVHDAAREGFLDTLILLHRAGARL |

**<Monotremes>**

| | |
|---|---|
| *Dromiciops gliroides* | KYGADPNVPDPTTLTLPVHDAAREGFLDTLILLHRAGARL |
| *Ornithorhynchus anatinus* | QHGAEPNCPDPTTLTQPAHDAAREGFLDTLVLLHEAGARL |
| *Tachyglossus aculeatus* | QHGAEPNSPDPTTLTQPAHDAAREGFLDTLVLLHGAGARL |

### Figure S2c

**Cdkn2a**

| | |
|---|---|
| *Homo sapiens* | LHGAEPNCADPATLTRPVHDAAREGFLDTLVVLHRAGARL |

**Cdkn2b**

**Birds**

| | |
|---|---|
| *Acanthisitta chloris* | QRGADPNRPDPRTGCLPAHDAARAGFLETLAALHRAGARL |
| *Accipiter nisus* | QRGADPNRPDPRTGSLPAHDAARAGFLETLAALHRAGARL |
| *Amazona aestiva* | QRGADPNRPDPRTGCFPAHDAARAGFLETLAVLHRAGARL |
| *Amazona collaria* | QRGADPNRPDPRTGCFPAHDAARAGFLETLAVLHRAGARL |
| *Anas platyrhynchos* | QRGADPNRPDPRTGCLPAHDAARAGFLDTLAALHRAGARL |
| *Anas zonorhyncha* | QRGADPNRPDPRTGCLPAHDAARAGFLDTLAALHRAGARL |
| *Anser brachyrhynchus* | QRGADPNRPDPRTGCLPAHDAARAGFLDTLAALHRAGARL |
| *Anser cygnoides* | QRGADPNRPDPRTGCLPAHDAARAGFLDTLAALHRAGARL |
| *Apteryx haastii* | QRGADPNRADPRTGCFPAHDAARAGFLDTLAALHRAG--- |
| *Apteryx owenii* | QRGADPNRADPRTGCFPAHDAARAGFLDTLAALHRA---- |
| *Apteryx rowi* | QRGADPNRPDPRTGCRPAHDAARAGF-------------- |
| *Aquila chrysaetos chrysaetos* | QRGADPNRPDPRTGSLPAHDAARAGFLETLAALHRAGARL |
| *Aythya fuligula* | QRGADPNRPDPRTGCLPAHDAARAGFLDTLAALHRAGARL |
| *Balearica regulorum gibbericeps* | QRGADPNRPDPRTGCFPAHDAARAGFLETLAALHRAGARL |
| *Cairina moschata domestica* | QRGADPNRPDPRTGCLPAHDAARAGFLDTLAALHRAGARL |
| *Calidris pugnax* | QRGADPNRPDPRTGCFPAHDAARAGFLETLAALHRAGARL |
| *Calidris pygmaea* | QRGADPNRPDPRTGCFPAHDAARAGFLETLAALHRAGARL |
| *Callipepla squamata* | QRGADPNRPDPRTGCRPAHDAARAGFLDTLAALHRAGARL |
| *Calypte anna* | QRGADPNRPDPRTGCLPVHDAARTGFLETLAVLHRAGARL |
| *Camarhynchus parvulus* | RHGADPSRPDPRTGCLPVHDAARAGFVETLAALHRAGARL |
| *Cariama cristata* | QRGADPNRPDPRTGCLPAHDAARAGFLETLAALHRAGARL |
| *Catharus ustulatus* | RRGADPNRPDPRTGCLPVHDAARAGFLETLAALHRAGARL |
| *Chiroxiphia lanceolata* | QRGADPNRPDPSTGCLPAHDAARAGFLETLAALHRAGARL |
| *Colinus virginianus* | QRGADPNRPDPRTGCRPAHDAARAGFLDTLAALHRAGARL |
| *Corapipo altera* | QRGADPNRPDPSTGCLPAHDAARAGFLETLAALHRAGARL |
| *Corvus cornix cornix* | RRGADPNRPDPRTGCLPAHDAARAGFLETLAALHRAGARL |
| *Corvus hawaiiensis* | RRGADPNRPDPRTGCLPAHDAARAGFLETLAALHRAGARL |
| *Corvus kubaryi* | RRGADPNRPDPRTGCLPAHDAARAGFLETLAALHRAGARL |
| *Corvus moneduloides* | RRGADPNRPDPRTGCLPAHDAARAGFLETLAALHRAGARL |
| *Coturnix japonica* | QRGADPNRPDPRTGCRPAHDAARAGFLDTLETLHRAGARL |
| *Cuculus canorus* | QRGADPNRPDPLTGCFPAHDAARAGFLETLAALHRAGARL |
| *Cyanistes caeruleus* | RRGADPNRPDPRTGCLPVHDAARAGFL------------- |
| *Cyanoderma ruficeps* | RRGADPNRPDPRTGCLPAHDAARAGFLDTLVVLHRAGARL |
| *Cygnus atratus* | QRGADPNRPDPRTGCFPAHDAARAGFLDTLAALHRAGARL |
| *Dromaius novaehollandiae* | QRGADPNRADPRTGCFPAHDAARAGFLDTLAALHRAGARL |
| *Dryobates pubescens* | QFGADPNRPDPSTGCFPIHDAARSGFLETVAVLHRAGARL |
| *Empidonax traillii* | QRGADPNRPDPSTGCLPVHDAVRAGFLETLVALHRAGARL |
| *Erythrura gouldiae* | RHGADPSRPDPRTGCLPAHDAARAGFLETLAALHRAGARL |
| *Eudyptes chrysocome* | QRGADPNRPDPRTGCLPAHDAARAGFLETLAALHRAGARL |
| *Eudyptes filholi* | QRGADPNRPDPRTGCLPAHDAARAGFLETLAALHRAGARL |
| *Eudyptes moseleyi* | QRGADPNRPDPRTGCLPAHDAARAGFLETLAALHRAGARL |
| *Eudyptes pachyrhynchus* | QRGADPNRPDPRTGCLPAHDAARAGFLEQLA--------- |
| *Eudyptes robustus* | QRGADPNRPDPRTGCLPAHDAARAGFLETLAALHRAGARL |
| *Eudyptes sclateri* | QRGADPNRPDPRTGCLPAHDAARAGFLETLAALHRAGARL |
| *Eudyptes schlegeli* | QRGADPNRPDPRTGCLPAHDAARAGFLETLAALH------ |
| *Eudyptula albosignata* | ERGADPNRPDPRTGCLPAHDAARAGFLETLAALHR----- |
| *Falco cherrug* | QRGADPNRPDPSTGCFPVHDAASAGFLETLEALHRAGARL |
| *Falco naumanni* | QRGADPNRPDPSTGCFPVHDAASAGFLETLEALHRAGARL |
| *Falco peregrinus* | QRGADPNRPDPSTGCFPVHDAASAGFLETLEALHRAGARL |
| *Falco rusticolus* | QRGADPNRPDPSTGCFPVHDAASAGFLETLEALHRAGARL |
| *Falco tinnunculus* | QRGADPNRPDPSTGCFPVHDAASAGFLETLEALHRAGARL |
| *Ficedula albicollis* | RRGADPNRPDPRTGCRPAHDAARAGFLETLAALHRP---- |
| *Gallus gallus* | QRGADPNRPDPRTGCRPAHDAARAGFLDTLAALHRAGARL |
| *Haliaeetus albicilla* | QRGADPNRPDPRTGSLPAHDAARAGFLETLAALHRAGARL |
| *Hirundo rustica* | RRGADPNRPDPRTGCFPVHDAARAGFLETLAALHRAGARL |
| *Hypotaenidia okinawae* | QRGADPNRPDPRTGCLPAHDAARAGFLETLAALHRAGARL |
| *Lagopus leucura* | QRGADPNRPDPRTGCRPAHDAARAGFLDTLAVLHRAGARL |
| *Lepidothrix coronata* | QRGADPNRPDPSTGCLPAHDAARAGFLETLAALHRARARL |
| *Lonchura striata domestica* | RHGADPNRPDPRTGCLPAHDAARAGFLETLAALHRAGARL |
| *Melopsittacus undulatus* | QRGADPNRPDPRTGCLPAHDAARAGFLETLTALHSAGARL |
| *Merops nubicus* | QRGADPNRPDPRTGCLPAHDAARAGFLETLAALHRAGARL |
| *Molothrus ater* | RHGADPNRPDPRTGCLPVHDAARAGFLGTLAALHRAGARL |
| *Motacilla alba alba* | RHGADPSRPDPRTGCLPVHDAARAGFLGTLAALHRAGARL |
| *Neopelma chrysocephalum* | QRGADPNRPDPSTGCLPAHDAARAGFLETLAALHRAGARL |
| *Nothoprocta perdicaria* | QRGADPNRADPRTGCYPAHDAARAGFVDTLAALHRAGARL |
| *Numida meleagris* | QRGADPNRPDPLTGCRPAHDAARAGFLDTLAALHRAGARL |
| *Onychostruthus taczanowskii* | RHGADPNRPDPRTGCLPAHDAARAGFLETLAALHRAGARL |
| *Otus sunia* | QRGADPNRPDPRTGCLPAHDAARAGFLETLAALHRAGARL |
| *Oxyura jamaicensis* | QRGADPNRPDPRTGCLPAHDAARAGFLDTLAALHRAGARL |
| *Parus major* | RRGADPNRPDPRTGCLPVHDAARAGFLETLAVLHRAGAHL |
| *Passer montanus* | RHGADPNRADPRTGCLPAHDAARAGFLETLAALHRAGARL |
| *Patagioenas fasciata monilis* | -------RPDPRTGCFPAHDAARAGFLETLAALHRAGARL |
| *Pavo cristatus* | QRGADPNRPDPRTGCRPAHDAARAGFLDTLAALHRAGARL |
| *Phasianus colchicus* | QRGADPNRPDPRTGCRPAHDAARAGFLDTLAALHRAGARL |
| *Pipra filicauda* | QRGADPNRPDPSTGCLPAHDAARAGFLETLAALHRAGARL |
| *Pseudopodoces humilis* | RRGADPNRPDPRTGCLPVHDAARAGFLETLAVLHRAGAPP |
| *Pterocles gutturalis* | -----PNRPDPLTGCLPAHDAARAGFLETLAALHRAGARL |
| *Pygoscelis antarcticus* | QRGADPNRPDPRTGCLPAHDAARAGFLEALAALHRAGARL |
| *Pyrgilauda ruficollis* | RHGADPNRPDPSTGCLPAHDAARAGFLETLAALHRAGARL |
| *Spheniscus humboldti* | QRGADPNRPDPRTGCLPAHDAARAGFLETLAA-------- |
| *Strigops habroptila* | QRGADPNRPDPRTGCLPAHDAARAGFLETLAALHRAGARL |
| *Strix occidentalis caurina* | -RGADPNRPDPRTGCLPAHDAARAGFLETLAALHRAGARL |
| *Sturnus vulgaris* | RRGADPNRPDPRTGCLPAHDAARAGFLETLAALHRAGARL |
| *Taeniopygia guttata* | RHGADPNRPDPRTGCLPAHDAARSGFLETLAALHRAGARL |
| *Tauraco erythrolophus* | ERGADPNRPDPRTGCLPAHDAARTGFLETLAALHRAGARL |
| *Turnix velox* | QHGADPNRPDPSTGCLPAHDAARAGFLDTLEALHRAGARL |
| *Tyto alba* | QRGADPNRPDPRTGCLPAHDAARAGFLETLAALHRAGARL |
| *Zonotrichia albicollis* | LHGADPNRPDPRTGCLPVHDAARAGFLGTLAALHRAGARL |
| *Zosterops borbonicus* | ----DPNRPDPRTGCLPAHDAARAGFLETLRALHRAGARL |

**Reptiles**

| | |
|---|---|
| *Alligator sinensis* | RSGADPNRPDPATGSRPAHDAAREGFLDTLVALHRAGARL |
| *Anolis carolinensis* | EAGANPNRPDP-SGFLPTHDAAREGFLDTIKVLHQGGARL |
| *Chelonia mydas* | QRGADPNRPDPRTGSLPVHDAAREGFLDTLVALHRGGARL |
| *Chelonoidis abingdonii* | QRGADPNRPDPRTGSLPVHDAAREGFLDTLVALHRGGARL |
| *Chelydra serpentina* | QRGADPNRPDPRTGSLPVHDAARGGFLDTLVALYRGGARL |
| *Chrysemys picta bellii* | QRGADPNRPDPRTGSLPVHDAAREGFLDTLVALHRGGARL |
| *Crocodylus porosus* | RSGADPNRPDPATGSCPAHDAAREGFLDTLAALHRAGARL |
| *Crotalus adamanteus* | QRGADPNRPDPSTGATPAHDLAQEGFLDTLMILHHWGARF |
| *Crotalus tigris* | QRGADPNRPDPSTGATPAHDLAQEGFLDTLMILHHWGARF |
| *Dermochelys coriacea* | QRGADPNRPDPLTGSLPVHDAAREGFLDTLVALHRGGARL |
| *Gekko japonicus* | QRGADPNRPDPCTGSLPAHDAAREGFLDTLQVLRQGGARL |
| *Gopherus agassizii* | QRGADPNRADPRTGSLPVHDAAREGFLDTLVALHRGISKX |
| *Gopherus evgoodei* | QRGADPNRADPRTGSLPVHDAAREGFLDTLVALHRGGARL |
| *Lacerta agilis* | QRGAGPNRPDPSTGTFPVHDAARGGFLDTLRVLRGGSARF |
| *Laticauda laticaudata* | QKGADPNRPDPSTGATPAHDVAREGFLDTLKLLHHCGAHF |
| *Mauremys mutica* | QRGADPNRPDPRTGSLPVHDAAREGFLDTLVALHRGGARL |
| *Mauremys reevesii* | QRGADPNRPDPRTGSLPVHDAAREGFLDTLVALHRGGARL |
| *Naja naja* | RSGADPNRPDPSTGATPAHDVAWEGFLDTLKLLHHWGASF |
| *Notechis scutatus* | RRGADPNRPDPSTGATPAHDVAQEGFLDTLKLLHHWGAHL |
| *Ophiophagus hannah* | QRGADPNRPDPSTGAMPAHDVAQEGFLDTLKVLHHWGARF |
| *Pantherophis guttatus* | QRGADPNQPDPSTGAMPAHDVSREGFLDTLKLLHHWGARF |
| *Pelodiscus sinensis* | QSGADPNRPDPSTGSLPVHDAARGGFLDTLVALHRGGARL |
| *Pelusios castaneus* | QRGADPDRPDPSTGSLPVHDAAREGFLDTLVALHRGGARL |
| *Podarcis muralis* | QRGADPNRPDPSTGTFPVHDAARGGFLDTLRVLRRGSARF |
| *Pogona vitticeps* | QGGADPNLPDPTTGSLPAHDAARAGFLDTLRALRRGGACF |
| *Protobothrops mucrosquamatus* | QRGADPNRPDPSTGATPAHDLAQEGFLDTLMILHHWGARF |
| *Pseudonaja textilis* | RRGADPNRPDPSTGATPAHDVAQEGFLDTLKLLHHWGAHF |
| *Python bivittatus* | QRGADPNRPDPSTGATPAHDVAWGGFLDTLKILYQWGARF |
| *Salvator merianae* | RKGADPNRQDHATGMVPAHDAAREGFLDTLQVLRRWGARF |
| *Sceloporus undulatus* | QAGANPNMPDRTTGSLPAHDAARQGFLDTLKVLHHWGARF |
| *Sphenodon punctatus* | RAGADPNIPDPATGSFPAHDAAREGFLDTLEVLHLGGARF |
| *Terrapene carolina triunguis* 1 | QRGADPNRPDPRTGSLPVHDAARDGFLDTLVALHRGGARL |
| *Terrapene carolina triunguis* 2 | QRGADPNRPDPRTGSLPVHDAARDGFLDTLVALHRGGARL |
| *Thamnophis elegans* | PRGADPNRPDPSTGATPAHDAAREGFLDTLKLLYHRGAHL |
| *Thamnophis sirtalis* | PRGADPNRPDPSTGATPAHDAAREGFLDTLKLLYHRGAHL |
| *Trachemys scripta elegans* | QRGADPNRPDPRTGSLPVHDAAREGFLDTLVALHRGGARL |
| *Varanus komodoensis* | QRGADPNRPDPSTGCLPAHDVACEGFLDTLQVLHGGGARF |
| *Zootoca vivipara* | QRGAGPNRPDPSTGTFPVHDAARGGFLDTLRALRGGSARF |

**Amphibians**

| | |
|---|---|
| *Ambystoma maculatum* | QHGANPNRPDPTTGTCPAHDAAREGFLETLVALLEGGASL |
| *Ambystoma mexicanum* | QRGANPNRPDPTTGTCPAHDAAREGFLETLVALLEGGASL |
| *Ambystoma velasci* | QRGANPNRPDPTTGTCPAHDAAREGFLETLVALLEGGASL |
| *Bufo bufo* | DHGADPTIPDPTTGTCPAHDAIREGFVDTLVVLINGGASI |
| *Bufo gargarizans* | DHGADPTIPDPTTGTCPAHDAIREGFVDTLVVLIKGGASI |
| *Engystomops pustulosus* | DHGADPNLPDPTTGTCPAHDAIRGGFVDTLAVLLKGGASV |
| *Geotrypetes seraphini* | LYGADPNVPDPTTNTYPVHDAAREGFLDTLQVLVRGGACL |
| *Hymenochirus boettgeri* | QWGADPRAQDPRTGTSPAHDAIREGFLDTLQVLLAGGASL |
| *Leptobrachium leishanense* | KHGANPRVPDPLTGTCPAHDAAREGFLDTLLILYQGGASL |
| *Microcaecilia unicolor* | RYGADPNVPDPTTGSCPVHDAAREGFLDTLQVLVRGGARL |
| *Nanorana parkeri* | EHGAEPSLPDPSTGTCPAHDAIREGFVDTLLELLNGGASL |
| *Rana temporaria* | EHGADPLIPDPSTGTCPAHDAIREGFVDTLLELVKGGASL |
| *Rhinatrema bivittatum* | RYGADPNLPDPTTAACPAHDAAREGFLDTLRVLLRAGASL |
| *Xenopus laevis* 1 | DYGADPSVPDPSTGTCPTHDAAREGFLDTLLVLLRNGANL |
| *Xenopus laevis* 2 | DQGADPNVPDPSTGTCPAHDAAREGFLDTLLVLLRNGASL |
| *Xenopus tropicalis* | DHGADPKLPDPCTGACPVHDAAREGFLDTLLVLLNNGASL |

**Figure S2d**

**Cdkn2a**

| | |
|---|---|
| *Homo sapiens* | LHGAEPNCADPATLTRPVHDAAREGFLDTLVVLHRA-GARL |

**Cdkn2b**

**Bony fish**

| | |
|---|---|
| *Acanthochromis polyacanthus* | EHGADPNVADPSTGSTPLHDAARTGFVDTARLLVKF-LADP |
| *Acanthopagrus latus* | NHGADPNVADSSTGTTPLHDAARTGYLDTVRLLVEY-QADP |
| *Acipenser ruthenus* 1 | MHGANPNIVDSSTGLTPIHDAAREGFLDTVGMLVRN-NANS |
| *Acipenser ruthenus* 2 | LYGANPNISDSSTGLTPVHDAAREGFLDTVKMLVQN-YANT |
| *Albula glossodonta* 1 | EHGANPNVPDSHTGATPLHDAAREGFLDTVAILIQY-HADI |
| *Albula glossodonta* 2 | RHGANPNLQDSSTGSTPLHDAARQGFLDTVEILIES-RADP |
| *Alosa alosa* | EHGADPNLRDSTTGTTPLHDAARTGYLETVRLLVEF-QADA |
| *Alosa sapidissima* | EHGADPNLRDSTTGTTPLHDAARTGYLETVRLLVEF-QADA |
| *Amphilophus citrinellus* | DHGADPNVADNSTGTTPLHDAAREGFLDTVRLLERH-HADP |
| *Amphiprion ocellaris* | EHGADPNVADRSTGSTPLHDAARTGFLDTARLLVKF-LADP |
| *Amphiprion percula* | EHGADPNVADRSTGSTPLHDAARTGFLDTARLLVKF-LADP |
| *Anabarilius grahami* | KYGADPNLPDPGTGSTPLHDAARTGFMDTVQLLIRF-NADP |
| *Anabas testudineus* | KHGANPNVAEGGTGTTPLHDAARTGFLDTVRLLVEH-GADP |
| *Anarrhichthys ocellatus* | EHGADPNVTDRSTGTTPLHDAAGTGFLDTARLLVQF-RADP |
| *Anguilla anguilla* | KHGANPNLSDESTGDTPLHDAARGGFLDTVKMLIEF-NADI |
| *Archocentrus centrarchus* | DHGADPNVADNSTGTTPLHDAAREGFLDTVRLLERH-HADP |
| *Astatotilapia calliptera* | EHGANSNVGDSSTGASPLHDAARAGFLDTVHLLVQH-HADP |
| *Astyanax mexicanus* | SRGADPNVPDLDTGATPLHDAARAGFSDTVSALLEF-NANP |
| *Austrofundulus limnaeus* | QRGANPNVADSSTGNTPLHDAAQTGFLETVRLLVEA-GAEP |
| *Bagarius yarrelli* | SHGANPNQADASTGVTPLHDAAREGFLHTVRLLVRA-GADI |
| *Betta splendens* | SHGANPNVADRSTGTTPLHDAARTGFLDTVRLLVQH-GADP |
| *Boleophthalmus pectinirostris* | ESGADPNVRDRSTGATPLHDAARTGFADTARLLVQY-HADP |
| *Caranx melampygus* | RHGADPNVADRSTGATPLHDAARTGFVETVRLLVEY-GADP |
| *Carassius auratus* 1 | EYGADPNVPDPGTGSTPLHDAARTGFMDTVQLLIRF-NADP |
| *Carassius auratus* 2 | EHGADPNVSDPGTGSTPLHDAARSGFTDTVRLLLRF-KADP |
| *Carassius auratus* 3 | EHGADPNVSDPGTGSTPLHDAARSGFTDTVRLLLRF-EADP |
| *Chanos chanos* | DHGADPNVSDESTGATPLHDTARLGFLETVRILVQH-HADP |
| *Cheilinus undulatus* | THGANPNVADRDTGHTPLHDAARAGFLDTVRLLVDF-RANP |
| *Chelmon rostratus* | KHGADPNVADRSTGTTPLHDAARTGFLDTVRLLVAF-LADP |
| *Clupea harengus* | EHGAVPNVRDCSTGTTPLHDAARTGYLETVRLLVEF-HADS |
| *Collichthys lucidus* | KHGADPNVPDCSTGTTPLHDAARAGFLETVRLLVEY-MADP |
| *Colossoma macropomum* | SCGAEPNVSDRSTGASPLHDAARAGFLDTVCVLLEF-KADP |
| *Coregonus clupeaformis* 1 | MQGADPNIADRHTGTTPLHDAARMGFXDTVEILVQF-LADP |
| *Coregonus clupeaformis* 2 | MQGADPNIADRHTGTTPLHDAARMGFLDTVEILVQF-LADP |
| *Cottoperca gobio* | KHGADPNVADRSTGGTPLHDAARTGFLDTVQLLVQF-LADP |
| *Cyclopterus lumpus* | NHGAAPNVRDRSTGTTPLHDAARAGFVDTVRLLVKC-GADP |
| *Cynoglossus semilaevis* | AHGANPNVADRSTGSTPLHDAARMGFVDTVRLLVDN-GADP |
| *Cyprinodon tularosa* | EHGADPNVADSSTGSTPLHDAARAGFLDTVRLLVEA-GANL |
| *Cyprinodon variegatus* | EHGADPNVADSSTGSTPLHDAARTGFLDTVRLLVEA-GANL |
| *Cyprinus carpio* 1 | EYGADPNVPDPGTGSTPLHDAARTGFMDTVRLLIRF-NADP |
| *Cyprinus carpio* 2 | EYGADPNVADPGTGSTPLHDAARTGFMDTVRLLIRF-KADP |
| *Danio rerio* | EQGADPNVPDPDTGSTPLHDAARTGFIDTVRLLIRF-GADP |
| *Danionella translucida* | DHGADPNVVDPCCGSSPLHDASRGGFLETVRLLVRF-HADP |
| *Denticeps clupeoides* | QHGGDPNLPDRSTGDTPLHDAARGGFLDTVRALVEF-HADP |
| *Dicentrarchus labrax* | EHGADPNVADPSTGTTPLHDAARTGFLDTVRLLVRY-KADP |
| *Echeneis naucrates* | EQGADPNVADVRTRTTPLHDAARTGFLDTVRLLVEY-KADP |
| *Electrophorus electricus* | AHGADPNVVDAHAGATPLHDAARTGFLDTVKILIQF-GANP |
| *Epinephelus coioides* | RAGADPNLKDISTGATPLHDAARTGFLDTVKALVEH-GADI |
| *Epinephelus lanceolatus* | RAGADPNLKDISTGATPLHDAARTGFLDTVKALVEH-GADI |
| *Esox lucius* | QQGADPNVADRHTGTMPLHDAARMGFLDTVKILVQC-NANP |
| *Etheostoma cragini* | KHGADPNVGDPSTGTTPLHDAARAGFLDTVRLLVQF-LADP |
| *Etheostoma spectabile* | KHGADPNVGDTSTGATPLHDAARAGFLDTVRLLVQF-LADP |
| *Fundulus heteroclitus* | ELGADPNVADRSTKATPLHDAARAGFLDTVRLLVEA-GADP |
| *Gadus morhua* | ARGSRPNVADSSTGTTPLHDAAREGFLDTLRVLLEF-KADP |
| *Gambusia affinis* | SRGADPNVADRSTGATPLHDAARTGFLDTVQLLVKA-GADP |
| *Gasterosteus aculeatus aculeatus* | KHGADPNAADRSTGATPLHDAARAGFLDTVRLLVQY-RAEP |
| *Girardinichthys multiradiatus* | THGADPNVADRSTRATPLHDAARAGFLGSVRLLVEA-GADP |
| *Gouania willdenowi* | KYGADPNVFDPTTGATPLHDAAQTGFLDTVRLLVRF-QGNP |
| *Gymnodraco acuticeps* | RHGADPAVSDRSTGSSPLHDAARTGFVDTVRLLVKN-RADP |
| *Haplochromis burtoni* | EHGANSNVGDSSTGASPLHDAARTGFLDTVHLLVQH-HADP |
| *Hippocampus comes* | AGGGDPNVADARTGSTPLHDAARSGFVDTVRILVRF-GANL |
| *Hippoglossus hippoglossus* | GHGADPNVADSSTGSTPLHDAARTGHVGTVRLLVEH-QADP |
| *Hippoglossus stenolepis* | GHGADPNVADSSTGSTPLHDAARTGHVGTVRLLVEH-QADP |
| *Hucho hucho* 1 | MHGADPNIADRHTGTTPLHDAARMGFLDTVEILVQF-LADP |
| *Hucho hucho* 2 | KQGADPNIADRHTGTTPLHDAARMGFLDTVEILVQF-LADP |
| *Iconisemion striatum* | QHGADPNVADGSTRTTPLHDAACTGFVDTVRLLVAA-RADP |
| *Ictalurus punctatus* | LHGANPNILDASTGATPLHDAAREGFLLTVQLLVRA-GADV |
| *Kryptolebias marmoratus* | RHQADPNVADRSTGTSPLHDAAREGFTDTVKLLLQA-QADP |
| *Labeo rohita* | EYGADPNVADPGTGSTPLHDAARTGFMDTVQLLIRF-NADP |
| *Labrus bergylta* | THGANPNVADRSTEHTPLHDAARMGFLDTVKLLVEF-KANP |
| *Larimichthys crocea* | ---ADPNLADCSTGTTPLHDAARTGFLE------------ |
| *Lates calcarifer* | THGANPNVVDCSTGDTPLHDAARTGYLDTVRLLVEY-GADP |
| *Latimeria chalumnae* | LSGADPNRRD-PTGLTAVHDAAREGFLDTVRILCQY-GGQM |
| *Lepisosteus oculatus* 1 | SHGADPNVSDCSTFSTPLHDAAREGFLCTVKVLVQY-NADR |
| *Lepisosteus oculatus* 2 | LYGANPNLQDVTTGMTPLHDAASGGFLDTVMILDQY-HADR |
| *Liparis tanakae* | NHGAAPNVRDRSTGTTPLHDAARAGFVDTVRLLVTF-RADP |
| *Mastacembelus armatus* | KHGGDPNVADRHTGTTPLHDAARTGFLDTVRLLVNY-RADP |
| *Maylandia zebra* | EHGANSNVGDSSTGASPLHDAARTGFLDTVHLLVQH-HADP |
| *Megalobrama amblycephala* | KYGADPNLPDPGTGSTPLHDAARTGFMDTVQLLIRF-NADP |
| *Megalops atlanticus* | KHGANPNVSDCHTGATPLHDAAREGFLETVQILVQF-YANK |
| *Megalops cyprinoides* 1 | EHGANPNVSDSHTGATPLHDAARGGFLETVAILIQY-QADV |
| *Megalops cyprinoides* 2 | RHGANPNVSDCHTGATPLHDAAREGFLETVEILIQF-HADK |
| *Melanotaenia boesemani* | EHGADPNVADISTGTTPLHDAASLGFLDTVRLLLQA-QANP |
| *Micropterus dolomieu* | EHGADPNVADRSTGATPLHDAARTGFLDTVRLLVEK-QADP |
| *Micropterus salmoides* | EHGADPNVADRSTGATPLHDAARTGFLDTVRLLVEK-QADP |
| *Mola mola* | NHGADPTVTDRSTGATPLHDAARAGFLDTVQLLVLF-RADP |
| *Monopterus albus* | RHGADPNVADRRTGATPLHDAARAGFLDTVRLLVNY-KADP |
| *Morone saxatilis* | EHGADPNVADPSTGTTPLHDAARTGFLDTVRLLVRY-KADP |
| *Mugil cephalus* | SRGADPNVKDRSTGTTPLHDAARTGFVDTVRLLVES-GADP |
| *Myripristis murdjan* | RRGADPNVADRCTGATPLHDAARTGFVDTVRLLVQH-NGDP |
| *Nematolebias whitei* | QHGADPNLADCSTGSTPLHDAARAGFLDTVRLLVEA-RAEP |
| *Neogobius melanostomus* | KYGADPNVKDRSTGATPLHDAARTGFTDTVRLLVHF-HADP |
| *Neolamprologus brichardi* | EHGANPNVGDSSTGASPLHDAARTGFLDTVHLLVQH-HADP |
| *Nibea albiflora* | KHGADPTLADRSTGATPLHDAARTGFLETVRLLVEY-MADP |
| *Nothobranchius furzeri* | QHGADPNVADRSTRTTPLHDAARTGFVDMVRLLVAA-QADP |
| *Nothobranchius kadleci* | QHGADPNVADRSTRTTPLHDAARTGFVDTVRLLVAA-QADP |
| *Nothobranchius korthausae* | QHGADPNVADRSTRTTPLHDAARTGFVDTVRLLVAA-QADP |
| *Nothobranchius kuhntae* | QHGADPNVADRSTRTTPLHDAARTGFVDTVRLLVAA-QADP |
| *Nothobranchius pienaari* | QHGADPNVADTSTRTTPLHDAAQ----------------- |
| *Nothobranchius rachovii* | QHGADPNVADTSTRTTPLHDAARSGFVDTVRLLVAA-QADP |
| *Notolabrus celidotus* | EHGANPNVADCTTGHTPLHDAARAGFLDTVRLLIEF-NANP |
| *Notothenia coriiceps* | RHGADPSVRDRSTGSSPLHDAARTGFVDTVRLLVKN-RADP |
| *Oncorhynchus gorbuscha* | MQGADPNIADRHTGTTPLHDAARMGFLDTVEILVQF-LADP |
| *Oncorhynchus keta* | MQGADPNIADRHTGTTPLHDAARMGFLDTVEILVQF-LADP |
| *Oncorhynchus kisutch* | MQGADPNIADRHTGTTPLHDAARMGFLDTVEILVQF-LADP |
| *Oncorhynchus mykiss* 1 | MQGADPNIADRHTGTTPLHDAARMGFLDTVEILVQF-LADP |
| *Oncorhynchus mykiss* 2 | MQGADPNIADRHTGTTPLRDAARMGFLDTVEILVQF-LADP |
| *Oncorhynchus nerka* | MQGADPNIADRHTGTTPLHDAARMGFLDTVEILVQF-LADP |
| *Oncorhynchus tshawytscha* | MQGADPNIADRHTGTTPLHDAARMGFLDTVEILVQF-LADP |
| *Onychostoma macrolepis* | EYGADPNVPDPGTGSSPLHDAARTGFMDTVRLLIRF-NADP |
| *Oreochromis aureus* | EHGANPNVGDSSTGASPLHDAARTGFVDTVRLLVQH-HADP |
| *Oreochromis niloticus* | EHGANPNVGDSSTGASPLHDAARTGFVDTVHLLVQH-HADP |
| *Oryzias javanicus* | RHGADPNVADESTGATPLHDAARTGFLDTVRLLVRA-GADP |
| *Oryzias latipes* | RHGADPNVADSSTGATPLHDAARTGFLDTVQLLVEA-GADP |
| *Oryzias melastigma* | RHGADPNVADQTTGATPLHDAARTGFLDTVRLLVQA-GADP |
| *Oryzias sinensis* | RHGADPNVADSSTGATPLHDAARTGFLDTAQLLVEA-GADP |
| *Pangasianodon hypophthalmus* | SHGANPNMADVSTGVTPLHDAAREGFLLTVQLLVRA-GADV |
| *Paralichthys olivaceus* | SHGADPNLADSSTGSTPLHDAARTGFLDTVRLLVEH-HADP |
| *Parambassis ranga* | RHGADPNVSD-STGATPLHDAARTGFLDTVRLLLEF-QADP |
| *Paramormyrops kingsleyae* | QYGADPSLRDRDTGGTPLHDASRGGFLDTVKVLVAH-GAEV |
| *Perca flavescens* | KHGADPNVADRSTGTTPLHDAARTGFLDTVRLLVQF-LADP |
| *Perca fluviatilis* | KHGADPNVADTSTGTTPLHDAARTGFLDTVRLLVQF-LADP |
| *Periophthalmus magnuspinnatus* | KHGADPNVRDRSTGATPLHDAVRTGFVDTARLLIEF-QADP |
| *Pimephales promelas* | KYGADPNVPDPGTGSTPLHDAARAGFMDTVRLLILF-NADP |
| *Plectropomus leopardus* | KHGADPNVADGSTGSTPLHDAARTGFLDTVRLLVQH-LADP |
| *Poecilia formosa* | TSGADPNVADRSTGATPLHDAARTGFLDTVKLLVEA-GADR |
| *Poecilia latipinna* | TSGADPNVADRSTGATPLHDAARTGFLDTVKLLVEA-GADR |
| *Poecilia mexicana* | TSGADPNVADRSTGATPLHDAARTGFLDTVKLLVEA-GADR |
| *Poecilia reticulata* | TSGADPNVADRSTGATPLHDAARTGFLDTVKLLVEA-GADR |
| *Poeciliopsis prolifica* | NHGADPNVADRSTGATPLHDAARTGFLDTVRLLLEF-GADP |
| *Polyodon spathula* | LYGANPNISDSSTGLTPVHDAAREGFLDTVMVLVRN-SANT |
| *Polyodon spathula* | LHGANPNTVDSSTGLTPIHDAAREGFLDTVVMLVRN-NADT |
| *Pseudochaenichthys georgianus* | RHGADPAVRDRSTGSSPLHDAARTGFVDTVRLLVKN-RADP |
| *Pundamilia nyererei* | EHGANSNVGDSSTGASPLHDAARTGFLDTVHLLVQH-HADP |
| *Pungitius pungitius* | KHGADPNVTDRSSGATPLHDAARTGFLDTVRLLVEF-GADP |
| *Puntigrus tetrazona* | EHGADPNVPDPRTGSTPLHDAARTGFTDTVRLFVRF-NADP |
| *Pygocentrus nattereri* | SYGAEPNVSDMSTGATPLHDAARAGFTDTVRVLLAF-KADP |
| *Salarias fasciatus* | KHGADPNVADRSTGATPLHDAARTGFLDTVRLLVHF-GADP |
| *Salmo salar* 1 | MQGADPNIADRHTGTTPLHDAARMGFLDTVEILVQF-HADP |
| *Salmo salar* 2 | MQGADPNIADRHTGTTPLHDAARMGFLDTVEILVQF-LADP |
| *Salmo trutta* | MQGADPNIADRHTGTTPLHDAARMGFLDTVEILVQF-HADP |
| *Salvelinus alpinus* | MKGADPNIADRHTGTTPLHDAARMGFLDTVEILVQF-LADP |
| *Salvelinus namaycush* | MKGADPNIADRHTGTTPLHDAARMGFLDTVEILVQF-LADP |
| *Sander lucioperca* | KHGADPNVADTSTGTTPLHDAARTGFVDTVRLLVKF-QADP |
| *Scatophagus argus* | SHGADPNVKDRSTGSTPLHDAARMGFLDTVRLLVRF-QADP |
| *Scleropages formosus* 1 | SHGADPNLGDPLTGSTPLHDAARLGFLDTVRLLVRF-GARV |
| *Scleropages formosus* 2 | DHGANPEVQD-STGSSPLHDAAREGFLETVQLLCSF-QANP |
| *Scophthalmus maximus* | KHGADPNVADGRTGSSPLHDAAREGFVDTARLLVEH-LADP |
| *Sebastes umbrosus* | THGADPSVTDGSTGATPLHDAARTGFLDTVRLLVQF-LADP |
| *Seriola dumerili* | KYGADPNVADTSTGTTPLHDAARTGFVDTVRLLVEY-LADP |
| *Seriola lalandi dorsalis* | KYGADPNVADTSTGTTPLHDAARTGFVDTVRLLVEY-QAEP |
| *Silurus meridionalis* | SHGGNPNAVDASTGVTPLHDAAREGFLATVELLVRG-GANV |
| *Simochromis diagramma* | EHGANSNVGDSSTGASPLHDAARTGFLDTVHLLVQH-HADP |
| *Sinocyclocheilus anshuiensis* 1 | EYGADPNVADPGTGSTPLHDAARTGFMDTVRLLIRF-KADP |
| *Sinocyclocheilus anshuiensis* 2 | EYGADPNVPDPGTGSTPLHDAARSGFMDTVRLLIRF-NADP |
| *Sinocyclocheilus grahami* 1 | EHGADPNVADPGTGSTPLHDAARTGFTDTVRLLIRF-KADP |
| *Sinocyclocheilus grahami* 2 | EYGADPNVPDPGTGSTPLHDAARSGFMDTVRLLIRF-NADP |
| *Sinocyclocheilus rhinocerous* 1 | EYGADPNVADPGTGSTPLHDAARTGFTDTVRLLIRF-KADP |
| *Sinocyclocheilus rhinocerous* 2 | EYGADPNVPDPGTGSTPLHDAARSGFMDTVRLLIRF-NADP |
| *Solea senegalensis* | THGADPNVADTSTGRTPLHDAASAGFVDTVRLLVKH-LADP |
| *Sparus aurata* | NHGADPNVADSSTGTTPLHDAARTGYLDTVRLLVEY-QADP |
| *Sphaeramia orbicularis* | KHGADPNVADPRTGSTPLHDAARTGFLDTVRLLVEY-TADP |
| *Stegastes partitus* | EHGADPNVADGTTGSTPLHDASRTGFVDTVRLLVKF-MADP |
| *Syngnathus acus* | EGGGDPNVADSSTGSTPLHDAARTGFSDTARLLVRF-AASP |
| *Tachysurus fulvidraco* | SHGANPNVVDVSTGATPLHDAAREGFLTTVELLVRA-GADI |
| *Takifugu flavidus* | DHGADPNVADGTTGTTPLHDAARSGFLDTVRLLVRF-TADP |
| *Takifugu rubripes* | DHGADPNVADGTTGATPLHDAARSGFLDTVRLLVRF-TADP |
| *Tetraodon nigroviridis* | DHAADPNVADRSTGARPLHDAARSGFLDTVRLLVLF-GANP |
| *Thalassophryne amazonica* | ESGADPNVADRSTGTTPLHDAARTGFVDTVRLLVQF-GADP |
| *Thunnus albacares* | KHGADPNVTDRHTGATPLHDAARMGFLDTARLLVDF-LANP |
| *Thunnus maccoyii* | EHGADPNVTDRHTGATPLHDAARTGFLDTARLLVDF-LANP |
| *Toxotes jaculatrix* | KHGADPNLPDKNTGATPLHDAAMTGFLDTVRLLVAC-GANP |
| *Trematomus bernacchii* | RHGADPAVRDRSTGSSPLHDAARTGFVDTVRLLVKN-RADP |
| *Triplophysa tibetana* | EHGADPNLPDPGTGSTPLHDAARTGFLDTLKLLIHF-SADP |
| *Xiphias gladius* | KYGGDPNVADRSTGATPLHDAARAGFLDTVRLLVEY-LADP |
| *Xiphophorus couchianus* | TRGADPNVTDKSTGATPLHDAARTGFLDTVQLLVKA-GADP |
| *Xiphophorus hellerii* | TRGADPNVTDKSTGATPLHDAARTGFLDTVQLLVEA-GADP |
| *Xiphophorus maculatus* | TRGADPNVTDKSTGATPLHDAARTGFLDTVQLLVKA-GADP |

**Cartilaginous fish**

| | |
|---|---|
| *Amblyraja radiata* | NWGADPDRQD-GYSIALIHDLAREGHLDTLNVLVEVGNANI |
| *Callorhinchus milii* | ERGADPNRQD-SMGMALIHDVAREGFLDTLQVLVGFGHANP |
| *Carcharodon carcharias* | EHGADPNRQD-NQQIALIHDVAREGFLDTLKVLVEFGMADT |
| *Chiloscyllium plagiosum* | EHGADPNRQD-NHQMALIHDVAREGFLDTLQVLIEVGKADT |
| *Hydrolagus affinis* | ENGADPNRRD-STGIALVHDVAREGFLDTLRVLVEVGNAKV |
| *Rhincodon typus* | QHGADPNRQD-NQQIALIHDVAREGFLDTLKVLIEVGNADA |
| *Scyliorhinus canicula* | RHGADPNRQD-NQQIAIIHDVAREGFLDTLQILVEIGNADT |
| *Stegostoma fasciatum* | QHGADPNRQD-NQQIALIHDVAREGFLDTLQVLIEVGNADA |

**Cdkn2b/c**

**Cyclostomes**

| | |
|---|---|
| *Eptatretus burgeri* | EASAQLNVRDPSCGLTPAQDAAREEFADTLRFLAQA-GAQL |
| *Entosphenus tridentatus* | AAGADPNVRDAVLGLTPAHDAAREGFVATLAELRAA-GASL |
| *Lethenteron camtschaticum* | AAGADPNVRDAVLGLTPAHDAAREGFVAVLAELRAA-GASL |
| *Lethenteron reissneri* | AAGADPNVRDAVLGLTPAHDAAREGFVAVLAELRAA-GASL |
| *Petromyzon marinus* | AAGADPDVRDAVLGLTPAHDAAREGFVAVLAELRAA-GASL |

### Figure S2e

**Cdkn2a**

| | |
|---|---|
| *Homo sapiens* | LHGAEPNCADPATLTRPVHDAAREGFLDTLVVLHRAGARL |

**Cdkn2c**
**Mammals**
**<Placentarians>**

| | |
|---|---|
| *Acinonyx jubatus* | LRGANPDLKDRT-GFAVIHDAARAGFLDTLQTLLEFQADV |
| *Ailuropoda melanoleuca* | LRGANPDLKDRT-GFAVIHDAARAGFLDTLQTLLEFQADV |
| *Aotus nancymaae* | LKGANPDLKDRT-GFAVIHDAARAGFLDTLQTLLEFQADV |
| *Artibeus jamaicensis* | LRGANPNLKDGT-GFAVIHDAARAGFLDTLQTLLEFQADV |
| *Arvicanthis niloticus* | LRGANPNLKDRT-GFAVIHDAARAGFLDTVQALLEFQADV |
| *Arvicola amphibius* | LRGANPNLKDRT-GFAVIHDAARAGFLDTVRALLEFQADV |
| *Balaenoptera acutorostrata scammoni* | LRGANPDLKDRT-GFAVIHDAARAGFLDTLQTLLEFQADV |
| *Balaenoptera musculus* | LRGANPDLKDRT-GFAVIHDAARAGFLDTLQTLLEFQADV |
| *Bison bison bison* | LRGANPDLKDRT-GFAVIHDAARAGFLDTLQTLLEFQADV |
| *Bos indicus* | LRGANPDLKDRT-GFAVIHDAARAGFLDTLQTLLEFQADV |
| *Bos mutus* | LRGANPDLKDRT-GFAVIHDAARAGFLDTLQTLLEFQADV |
| *Bos taurus* | LRGANPDLKDRT-GFAVIHDAARAGFLDTLQTLLEFQADV |
| *Bubalus bubalis* | LRGANPDLKDRT-GFAVIHDAARAGFLDTLQTLLEFQADV |
| *Callithrix jacchus* | LRGANPDLKDRT-GFAVIHDAARAGFLDTLQTLLEFQADV |
| *Callorhinus ursinus* | LRGANPDLKDRT-GFAVIHDAARAGFLDTLQTLLEFQADV |
| *Camelus bactrianus* | LRGANPDLKDRT-GFAVIHDAARAGFLDTLQTLLEFQADV |
| *Camelus dromedarius* | LRGANPDLKDRT-GFAVIHDAARAGFLDTLQTLLEFQADV |
| *Camelus ferus* | LRGANPDLKDRT-GFAVIHDAARAGFLDTLQTLLEFQADV |
| *Canis lupus familiaris* | LRGANPDLKDRT-GFAVIHDAARAGFLDTLQTLLEFQADV |
| *Capra hircus* | LRGANPDLKDRT-GFAVIHDAARAGFLDTLQTLLEFQADV |
| *Carlito syrichta* | LRGANPDLKDRT-GFAVIHDAARAGFLDTLQTLLEFQADV |
| *Castor canadensis* | LRGANPNLKDRT-GFAVIHDAARAGFLDTLQTLLEFQADV |
| *Cavia porcellus* | LRGANPNLKDRT-GFAVIHDAARAGFLDTLQTLLEFQADV |
| *Cebus imitator* | LRGANPDLKDRT-GFAVIHDAARAGFLDTLQTLLEFQADV |
| *Ceratotherium simum simum* | LRGANPDLKDRT-GFAVIHDAARAGFLDTLQTLLEFQADV |
| *Cercocebus atys* | LRGANPDLKDRT-GFAVIHDAARAGFLDTLQTLLEFQADV |
| *Cervus canadensis* | LRGANPDLKDRT-GFAVIHDAARAGFLDTLQTLLEFQADV |
| *Cervus elaphus* | LRGANPDLKDRT-GFAVIHDAARAGFLDTLQTLLEFQADV |
| *Cervus hanglu yarkandensis* | LRGANPDLKDRT-GFAVIHDAARAGFLDTLQTLLEFQADV |
| *Chinchilla lanigera* | LRGANPNLKDRT-GFAVIHDAARAGFLDTLQTLLEFQADV |
| *Chlorocebus sabaeus* | LRGANPDLKDRT-GFAVIHDAARAGFLDTLQTLLEFQADV |
| *Choloepus didactylus* | LRGANPDLKDRT-GFAVIHDAARAGFLDTLQTLLEFQADV |
| *Chrysochloris asiatica* | LRGANPNLKDGT-GFAVIHDAARAGFLDTLQTLLEFQADV |
| *Colobus angolensis palliatus* | LRGANPDLKDRT-GFAVIHDAARAGFLDTLQTLLEFQADV |
| *Condylura cristata* | LSGANPDLKDRT-GFAVIHDAARAGFLDTLQTLLEFQADV |
| *Cricetulus griseus* | LRGANPNLKDRT-GFAVIHDAARAGFLDTVQALLEFQADV |
| *Crocuta crocuta* | LRGANPDLKDRT-GFAVIHDAARAGFLDTLQTLLEFQADV |
| *Dasypus novemcinctus* | LRGANPDLKDRT-GFAVIHDAARAGFLDTLQTLLEFQADV |
| *Delphinapterus leucas* | LRGANPDLKDRT-GFAVIHDAARAGFLDTLQTLLEFQADV |
| *Desmodus rotundus* | LRGANPNLKDGT-GFAVIHDAARAGFLDTLQTLLEFQADV |
| *Dipodomys ordii* | LKGANPNLKDGT-GFAVIHDAARAGFLDTLQTLLEFQADV |
| *Dipodomys spectabilis* | LKGANPNLKDGT-GFAVIHDAARAGFLDTLQTLLEFQADV |
| *Echinops telfairi* | LRGANPDLRDQT-GFAVIHDAARAGFLDTLQTLLEFQADV |
| *Elephantulus edwardii* | LRGANPNLKDQT-GFAVIHDAARAGFLDTIQTLLEFQADV |
| *Enhydra lutris kenyoni* | LRGANPDLKDRT-GFAVIHDAARAGFLDTLQTLLEFQADV |
| *Eptesicus fuscus* | LRGANPNLKDGT-GFAVIHDAARAGFLDTLQTLLEFQADV |
| *Equus asinus* | LRGANPDLKDRT-GFAVIHDAARAGFLDTLQTLLEFQADV |
| *Equus caballus* | LRGANPDLKDRT-GFAVIHDAARAGFLDTLQTLLEFQADV |
| *Equus przewalskii* | LRGANPDLKDRT-GFAVIHDAARAGFLDTLQTLLEFQADV |
| *Equus quagga* | LRGANPDLKDRT-GFAVIHDAARAGFLDTLQTLLEFQADV |
| *Erinaceus europaeus* | LRGANPDLKDQT-GFAVIHDAARAGFLDTLQTLLEFQADV |
| *Eumetopias jubatus* | LRGANPDLKDRT-GFAVIHDAARAGFLDTLQTLLEFQADV |
| *Felis catus* | LRGANPDLKDRT-GFAVIHDAARAGFLDTLQTLLEFQADV |
| *Fukomys damarensis* | LRGANPNLKDRT-GFAVIHDAARAGFLDTLQTLLEFQADV |
| *Galemys pyrenaicus* | LSGANPDLKDRT-GFAVIHDAARAGFLDTLQTLLEFQADV |
| *Galeopterus variegatus* | LKGANPDLKDRT-GFAVIHDAARAGFLDTLQTLLEFQADV |
| *Globicephala melas* | LRGANPDLKDRT-GFAVIHDAARAGFLDTLQTLLEFQADV |
| *Gorilla gorilla gorilla* | LRGANPDLKDRT-GFAVIHDAARAGFLDTLQTLLEFQADV |
| *Grammomys surdaster* | LRGANPNLKDRT-GFAVIHDAARAGFLDTVQALLEFQADV |
| *Gulo gulo luscus* | LRGANPDLKDRT-GFAVIHDAARAGFLDTLQTLLEFQADV |
| *Halichoerus grypus* | LRGANPDLKDRT-GFAVIHDAARAGFLDTLQTLLEFQADV |
| *Heterocephalus glaber* | LRGANPNLKDRT-GFAVIHDAARAGFLDTLQTLLEFQADV |
| *Hipposideros armiger* | LRGANPNLKDGT-GFAVIHDAARAGFLDTLQTLLEFQADV |
| *Homo sapiens* | LRGANPDLKDRT-GFAVIHDAARAGFLDTLQTLLEFQADV |
| *Hyaena hyaena* | LRGANPDLKDRT-GFAVIHDAARAGFLDTLQTLLEFQADV |
| *Hylobates moloch* | LRGANPNLKDRT-GFAVIHDAARAGFLDTLQTLLEFQADV |
| *Ictidomys tridecemlineatus* | LRGANPNLKDRT-GFAVIHDAARAGFLDTLQTLLEFQADV |
| *Jaculus jaculus* | LRGANPNLKDQT-GFAVIHDAARAGFLDTVQTLLEFQADV |
| *Lagenorhynchus obliquidens* | LRGANPDLKDRT-GFAVIHDAARAGFLDTLQTLLEFQADV |
| *Lemur catta* | LRGANPDLKDRT-GFAVIHDAARAGFLDTLQTLLEFKADV |
| *Leopardus geoffroyi* | LRGANPDLKDRT-GFAVIHDAARAGFLDTLQTLLEFQADV |
| *Leptonychotes weddellii* | LRGANPDLKDRT-GFAVIHDAARAGFLDTLQTLLEFQADV |
| *Lipotes vexillifer* | LRGANPDLKDRT-GFAVIHDAARAGFLDTLQTLLEFQADV |
| *Lontra canadensis* | LRGANPDLKDRT-GFAVIHDAARAGFLDTLQTLLEFQADV |
| *Loxodonta africana* | LRGANPNLKDRT-GFAVIHDAARAGFLDTLQTLLEFQADV |
| *Lynx canadensis* | LRGANPDLKDRT-GFAVIHDAARAGFLDTLQTLLEFQADV |
| *Lynx pardinus* | LRGANPDLKDRT-GFAVIHDAARAGFLDTLQTLLEFQADV |
| *Macaca fascicularis* | LRGANPDLKDRT-GFAVIHDAARAGFLDTLQTLLEFQADV |
| *Macaca mulatta* | LRGANPDLKDRT-GFAVIHDAARAGFLDTLQTLLEFQADV |
| *Macaca nemestrina* | LRGANPDLKDRT-GFAVIHDAARAGFLDTLQTLLEFQADV |
| *Mandrillus leucophaeus* | LRGANPDLKDRT-GFAVIHDAARAGFLDTLQTLLEFQADV |
| *Manis javanica* | LRGANPDLKDRT-GFAVIHDAARAGFLDTLQTLLEFQADV |
| *Manis pentadactyla* | LRGANPDLKDRT-GFAVIHDAARAGFLDTLQTLLEFQADV |
| *Marmota flaviventris* | LRGANPNLKDRT-GFAVIHDAARAGFLDTLQTLLEFQADV |
| *Marmota marmota marmota* | LRGANPNLKDRT-GFAVIHDAARAGFLDTLQTLLEFQADV |
| *Marmota monax* | LRGANPNLKDRT-GFAVIHDAARAGFLDTLQTLLEFQADV |
| *Mastomys coucha* | LRGANPNLKDRT-GFAVIHDAARAGFLDTVQALLEFQADV |
| *Meles meles* | LRGANPDLKDRT-GFAVIHDAARAGFLDTLQTLLEFQADV |
| *Meriones unguiculatus* | LRGANPNLKDQT-GFAVIHDAARAGFLDTVQALLEFQADV |
| *Mesocricetus auratus* | LRGANPNLKDRT-GFAVIHDAARAGFLDTVQALLEFQADV |
| *Microcebus murinus* | LRGANPDLKDRT-GFAVIHDAARAGFLDTLQTLLEFKADV |
| *Microtus ochrogaster* | LRGANPNLKDRT-GFAVIHDAARAGFLDTVQALLEFQADV |
| *Microtus oregoni* | LRGANPNLKDRT-GFAVIHDAARAGFLDTVRALLEFQADV |
| *Miniopterus natalensis* | LRGANPNLKDGT-GFAVIHDAARAGFLDTLQTLLEFQADV |
| *Mirounga angustirostris* | LRGANPDLKDRT-GFAVIHDAARAGFLDTLQTLLEFQADV |
| *Mirounga leonina* | LRGANPDLKDRT-GFAVIHDAARAGFLDTLQTLLEFQADV |
| *Molossus molossus* | LSGANPNLKDGT-GFAVIHDAARAGFLDTLQTLLEFQADV |
| *Monodon monoceros* | LRGANPDLKDRT-GFAVIHDAARAGFLDTLQTLLEFQADV |
| *Mus caroli* | LRGANPNLKDGT-GFAVIHDAARAGFLDTVQALLEFQADV |
| *Mus musculus* | LRGANPNLKDGT-GFAVIHDAARAGFLDTVQALLEFQADV |
| *Mus pahari* | LRGANPNLKDGT-GFAVIHDAARAGFLDTVQALLEFQADV |
| *Mustela erminea* | LRGANPDLKDRT-GFAVIHDAARAGFLDTLQTLLEFQADV |
| *Mustela putorius furo* | LRGANPDLKDRT-GFAVIHDAARAGFLDTLQTLLEFQADV |
| *Myotis brandtii* | LKGANPNLKDGT-GFAVIHDAARAGFLDTLQTLLEFQADV |
| *Myotis davidii* | LRGADPNLKDGT-GFAVIHDAARAGFLDTLQTLLEFQADV |
| *Myotis lucifugus* | LRGANPNLKDGT-GFAVIHDAARAGFLDTLQTLLEFQADV |
| *Myotis myotis* | LRGADPNLKDGT-GFAVIHDAARAGFLDTLQTLLEFQADV |
| *Nannospalax galili* | LRGANPNLKDRT-GFAVIHDAARAGFLDTVQTLLEFQADV |
| *Neogale vison* | LRGANPDLKDRT-GFAVIHDAARAGFLDTLQTLLEFQADV |
| *Neomonachus schauinslandi* | LRGANPDLKDRT-GFAVIHDAARAGFLDTLQTLLEFQADV |
| *Neophocaena asiaeorientalis asiaeorientalis* | LRGANPDLKDRT-GFAVIHDAARAGFLDTLQTLLEFQADV |
| *Neosciurus carolinensis* | LRGANPNLKDRT-GFAVIHDAARAGFLDTLQTLLEFQADV |
| *Neotoma lepida* | LRGANPNLKDRT-GFAAIHDAARAGFLDTVQALLEFQADV |
| *Nomascus leucogenys* | LRGANPNLKDRT-GFAVIHDAARAGFLDTLQTLLEFQADV |
| *Nyctereutes procyonoides* | LRGANPDLKDRT-GFAVIHDAARAGFLDTLQTLLEFQADV |
| *Ochotona curzoniae* | LRGANPDLKDQT-GFAVIHDAAREGFLDTLQTLLEFQADV |
| *Ochotona princeps* | LRGANPDLKDQT-GFAVIHDAAREGFLDTLQTLLEFQADV |
| *Octodon degus* | LRGANPNLKDRT-GFAVIHDAARAGFLDTLQTLLEFQADV |
| *Odobenus rosmarus divergens* | LRGANPDLKDRT-GFAVIHDAARAGFLDTLQTLLEFQADV |
| *Odocoileus virginianus texanus* | LRGANPDLKDRT-GFAVIHDAARAGFLDTLQTLLEFQADV |
| *Onychomys torridus* | LRGANPNLKDPT-GFAVIHDAARAGFLDTVQALLEFQADV |
| *Orcinus orca* | LRGANPDLKDRT-GFAVIHDAARAGFLDTLQTLLEFQADV |
| *Orycteropus afer afer* | LRGANPDLKDRT-GFAVIHDAARAGFLDTLQTLLEFQADV |
| *Oryctolagus cuniculus* | LRGANPDLKDRT-GFAVIHDAARAGFLDTLQTLLEFQADV |
| *Oryx dammah* | LRGANPDLKDRT-GFAVIHDAARAGFLDTLQTLLEFQADV |
| *Otolemur garnettii* | LRGANPDLKDRT-GFAVIHDAARAGFLDTLQTLLEFQADV |
| *Ovis aries* | LRGANPDLKDRT-GFAVIHDAARAGFLDTLQTLLEFQADV |
| *Pan paniscus* | LRGANPDLKDRT-GFAVIHDAARAGFLDTLQTLLEFQADV |
| *Pan troglodytes* | LRGANPDLKDRT-GFAVIHDAARAGFLDTLQTLLEFQADV |
| *Panthera leo* | LRGANPDLKDRT-GFAVIHDAARAGFLDTLQTLLEFQADV |
| *Panthera pardus* | LRGANPDLKDRT-GFAVIHDAARAGFLDTLQTLLEFQADV |
| *Panthera tigris* | LRGANPDLKDRT-GFAVIHDAARAGFLDTLQTLLEFQADV |
| *Papio anubis* | LRGANPDLKDRT-GFAVIHDAARAGFLDTLQTLLEFQADV |
| *Perognathus longimembris pacificus* | LKGANPNLKDGT-GFAVIHDAARAGFLDTLQTLLEFQADV |
| *Peromyscus leucopus* | LRGANPNLKDRT-GFAVIHDAARAGFLDTVQALLEFQADV |
| *Peromyscus maniculatus bairdii* | LRGANPNLKDRT-GFAVIHDAARAGFLDTVQALLEFQADV |
| *Phoca vitulina* | LRGANPDLKDRT-GFAVIHDAARAGFLDTLQTLLEFQADV |
| *Phocoena sinus* | LRGANPDLKDRT-GFAVIHDAARAGFLDTLQTLLEFQADV |
| *Phyllostomus discolor* | LGGANPNLKDGT-GFAVIHDAARAGFLDTLQTLLEFQADV |
| *Phyllostomus hastatus* | LGGANPNLKDGT-GFAVIHDAARAGFLDTLQTLLEFQADV |
| *Physeter catodon* | LRGANPDLRDRT-GFAVIHDAARAGFLDTLQTLLEFQADV |
| *Piliocolobus tephrosceles* | LRGANPDLKDRT-GFAVIHDAARAGFLDTLQTLLEFQADV |
| *Pipistrellus kuhlii* | LRGANPNLKDGT-GFAVIHDAARAGFLDTLQTLLEFQADV |
| *Pongo abelii* | LRGANPDLKDRT-GFAVIHDAARAGFLDTLQTLLEFQADV |
| *Prionailurus bengalensis* | LRGANPDLKDRT-GFAVIHDAARAGFLDTLQTLLEFQADV |
| *Procavia capensis* | LRGANPNLKDRT-GFAVIHDAARAGFLDTLQTLLEFQADV |
| *Propithecus coquereli* | LRGANPDLKDRT-GFAVIHDAARAGFLDTLQTLLEFKADV |
| *Pteropus alecto* | LRGANPNLKDGT-GFAVIHDAARAGFLDTLQTLLEFQADV |
| *Pteropus giganteus* | LRGANPNLKDGT-GFAVIHDAARAGFLDTLQTLLEFQADV |
| *Pteropus vampyrus* | LRGANPNLKDGT-GFAVIHDAARAGFLDTLQTLLEFQADV |
| *Puma concolor* | LRGANPDLKDRT-GFAVIHDAARAGFLDTLQTLLEFQADV |
| *Puma yagouaroundi* | LRGANPDLKDRT-GFAVIHDAARAGFLDTLQTLLEFQADV |
| *Rattus norvegicus* | LRGANPNLKDRT-GFAVIHDAARAGFLDTVQALLEFQADV |
| *Rattus rattus* | LRGANPNLKDRT-GFAVIHDAARAGFLDTVQALLEFQADV |
| *Rhinolophus ferrumequinum* | LRGANPNLRDGT-GFAVIHDAARAGFLDTLQTLLEFQADV |
| *Rhinolophus sinicus* | LRGANPNLRDGT-GFAVIHDAARAGFLDTLQTLLEFQADV |
| *Rhinopithecus bieti* | LRGANPDLKDRT-GFAVIHDAARAGFLDTLQTLLEFQADV |
| *Rhinopithecus roxellana* | LRGANPDLKDRT-GFAVIHDAARAGFLDTLQTLLEFQADV |
| *Rousettus aegyptiacus* | LRGANPNLKDGT-GFAVIHDAARAGFLDTLQTLLEFQADV |
| *Saimiri boliviensis boliviensis* | LRGANPDLKDRT-GFAVIHDAARAGFLDTLQTLLEFQADV |
| *Sapajus apella* | LRGANPDLKDRT-GFAVIHDAARAGFLDTLQTLLEFQADV |
| *Sorex araneus* | LRGANPDLKDGT-GFAVIHDAARAGFLDTLQILLEFQADV |
| *Sousa chinensis* | LRGANPDLKDRT-GFAVIHDAARAGFLDTLQTLLEFQADV |
| *Sturnira hondurensis* | LRGANPNLKDGT-GFAVIHDAARAGFLDTLQTLLEFQADV |
| *Suricata suricatta* | LRGANPDLKDRT-GFAVIHDAARAGFLDTLQTLLEFQADV |
| *Sus scrofa* | LRGANPDLKDRT-GFAVIHDAARAGFLDTLQTLLEFQADV |
| *Talpa occidentalis* | LSGANPDLKDRT-GFAVIHDAARAGFLDTLQTLLEFRADV |
| *Theropithecus gelada* | LRGANPDLKDRT-GFAVIHDAARAGFLDTLQTLLEFQADV |
| *Trachypithecus francoisi* | LRGANPDLKDRT-GFAVIHDAARAGFLDTLQTLLEFQADV |
| *Trichechus manatus latirostris* | LRGANPDLKDRT-GFAVIHDAARAGFLDTLQTLLEFQADV |
| *Tupaia chinensis* | LKGANPDLKDRT-GFAVIHDAARAGFLDTLQTLLEFQADV |
| *Tursiops truncatus* | LRGANPDLKDRT-GFAVIHDAARAGFLDTLQTLLEFQADV |
| *Urocitellus parryii* | LRGANPNLKDRT-GFAVIHDAARAGFLDTLQTLLEFQADV |
| *Ursus americanus* | LRGANPDLKDRT-GFAVIHDAARAGFLDTLQTLLEFQADV |
| *Ursus arctos* | LRGANPDLKDRT-GFAVIHDAARAGFLDTLQTLLEFQADV |
| *Ursus maritimus* | LRGANPDLKDRT-GFAVIHDAARAGFLDTLQTLLEFQADV |
| *Vicugna pacos* | LRGANPDLKDRT-GFAVIHDAARAGFLDTLQTLLEFQADV |
| *Vulpes lagopus* | LRGANPDLKDRT-GFAVIHDAARAGFLDTLQTLLEFQADV |
| *Vulpes vulpes* | LRGANPDLKDRT-GFAVIHDAARAGFLDTLQTLLEFQADV |
| *Zalophus californianus* | LRGANPDLKDRT-GFAVIHDAARAGFLDTLQTLLEFQADV |

**<Marsupials>**

| | |
|---|---|
| *Dromiciops gliroides* | LCGANPDLKDPT-GFAVLHDAARAGFLDTLQTLLEFQADV |
| *Gracilinanus agilis* | LRGANPDLKDRT-GFAVLHDAARAGFLDTLQTLLEFQADV |
| *Monodelphis domestica* | LRGANPDLKDRT-GFAVLHDAARAGFLDTLQTLLEFQADV |
| *Notamacropus eugenii* | LRGANPDLKDRT-GFAVLHDAARAGFLDTLQTLLEFQADV |
| *Phascolarctos cinereus* | LRGANPDLKDRT-GFAVLHDAARAGFLDTLQTLLEFQADV |
| *Sarcophilus harrisii* | LSGANPDLKDRT-GFAVLHDAARAGFLDTLQTLLEFQADV |
| *Trichosurus vulpecula* | LRGANPDLKDRT-GFAVLHDAARAGFLDTLQTLLEFQADV |
| *Vombatus ursinus* | LRGANPDLKDRT-GFAVLHDAARAGFLDTLQTLLEFQADV |

**<Monotremes>**

| | |
|---|---|
| *Ornithorhynchus anatinus* | VGGADPDLKDGT-GFAALHDAARAGYLDTLQTLLEFQADV |
| *Tachyglossus aculeatus* | LGGADPDLKDGT-GFAALHDAARAGYLDTLQTLLEFRADV |

### Figure S2f

**Cdkn2a**

*Homo sapiens* LHGAEPNCADPATLTRPVHDAAREGFLDTLVVLHRAGARL

**Cdkn2c**

**birds**

*Acanthisitta chloris* TNGANPNLKDSTGFA-VIHDVAREGFLDTLQTLLEFKADV
*Accipiter nisus* LSGADPNLKDSTGFA-VIHDVAREGFADTLQALLEFQADV
*Amazona aestiva* ISGANPNLKDSTGFA-VIHDVAREGFLDTLQTLLEFKADV
*Amazona collaria* ISGANPNLKDSTGFA-VIHDVAREGFLDTLQTLLEFKADV
*Anas platyrhynchos* MSGANPDLKDSTGFA-VIHDVARAGFLDTLQTLLEFKADV
*Anas zonorhyncha* MSGANPDLKDSTGFA-VIHDVARAGFLDTLQTLLEFKADV
*Anser brachyrhynchus* MSGANPDLKDSTGFA-VIHDVARAGFLDTLQTLLEFKADV
*Anser cygnoides* MSGANPDLKDSTGFA-VIHDVARAGFLDTLQTLLEFKADV
*Antrostomus carolinensis* MNGANPNLKDSTGFA-VMHDVAREGFLDTLQTLLEFKADV
*Apaloderma vittatum* LKGANPNLKDSTGYA-VIHDVAREGFLDTLQTLLEFQADV
*Aptenodytes forsteri* INGANPNLKDSTGFA-VIHDVAREGFLDTLQTLLEFKADV
*Aptenodytes patagonicus* INGANPNLKDSTGFA-VIHDVAREGFLDTLQTLLEFKADV
*Apteryx haastii* SRGANPNLKDSTGFA-VIHDVARAGFLDTLQTLLEFHADV
*Apteryx mantelli mantelli* SRGANPNLKDSTGFA-VIHDVARAGFLDTLQTLLEFHADV
*Apteryx owenii* SRGANPNLKDSTGFA-VIHDVARAGFLDTLQTLLEFHADV
*Apteryx rowi* SRGANPNLKDSTGFA-VIHDVARAGFLDTLQTLLEFHADV
*Aquila chrysaetos chrysaetos* LSGADPNLKDSTGFA-VIHDVAREGFADTLQALLEFQADV
*Athene cunicularia* ISGANPNLKDSTGFA-VIHDVAREGFLDTLQTLLEFKADV
*Aythya fuligula* MSGANPDLKDSTGFA-VIHDVARAGFLDTLQTLLEFKADV
*Balearica regulorum gibbericeps* INGANPNLKDSTGFA-VIHDVAREGFLDTLQTLLEFKADV
*Bubo bubo* ISGANPNLKDSTGFA-VIHDVAREGFLDTLQTLLEFKADV
*Buceros rhinoceros silvestris* INGANPNLKDSTGFA-VIHDVAREGFLDTLQTLLEFKADV
*Buteo japonicus* LSGADPNLKDSTGFA-VIHDVAREGFADTLQALLEFQADV
*Cairina moschata domestica* MSGANPDLKDSTGFA-VIHDVARAGFLDTLQTLLEFKADV
*Calidris pugnax* INGADPNLKDSTGFA-VIHDVAREGFLDTLQTLVEFQADV
*Calidris pygmaea* INGADPNLKDSTGFA-VIHDVAREGFLDTLQTLVEFKADV
*Callipepla squamata* MSGANPNLKDSTGFA-VIHDVARAGFLDTLQTLLEFHADV
*Calypte anna* MKGANPNLKDSTGFA-VIHDVAREGFLDTLQTLLEFEADV
*Camarhynchus parvulus* SNGANPNLKDSTGFA-VIHDVAREGFLDTLQTLLEFKADV
*Cariama cristata* INGANPNLKDSTGFA-VIHDVAREGFLDTLQTLLEFKADV
*Cathartes aura* INGANPNLKDSTGFA-VIHDVAREGFLDTLQTLLEFKADV
*Catharus ustulatus* SNGANPNLRDSTGFA-VIHDVAREGFLDTLQTLLEFKADV
*Centrocercus urophasianus* MSGANPNLKDSTGFA-VIHDVARAGFLDTLQTLLEFHADV
*Chaetura pelagica* LKGANPNLKDSTGFA-VIHDVAREGFLDTLQTLLEFEADV
*Charadrius vociferus* INGANPNLKDSTGFA-VIHDVAREGFLDTLQTLVEFKADV
*Chiroxiphia lanceolata* SKGANPNLKDSTGFA-IIHDVAREGFLDTLQTLLEFKADV
*Chlamydotis macqueenii* MKGANPDLKDSTGFA-VIHDVAREGFLDTLQTLLEFQADV
*Chrysolophus pictus* MSGANPNLKDSTGFA-VIHDVARAGFLDTLQTLLEFHADV
*Colinus virginianus* MSGANPNLKDSTGFA-VIHDVARAGFLDTLQTLLEFHADV
*Colius striatus* INGANPDLKDSTGFA-VIHDVAREGFLDTLQTLLEFKADV
*Columba livia* AHGANPDLRDSTGFA-VMHDVAREGFPDTLQALLEFQADV
*Corapipo altera* SKGANPNLKDSTGFA-IIHDVAREGFLDTLQTLLEFKADV
*Corvus brachyrhynchos* SNGANPNLKDSTGFA-VIHDVAREGFLDTLQTLLEFKADV
*Corvus cornix cornix* SNGANPNLKDSTGFA-VIHDVAREGFLDTLQTLLEFKADV
*Corvus hawaiiensis* SNGANPNLKDSTGFA-VIHDVAREGFLDTLQTLLEFKADV
*Corvus kubaryi* SNGANPNLKDSTGFA-VIHDVAREGFLDTLQTLLEFKADV
*Corvus moneduloides* SNGANPNLKDSTGFA-VIHDVAREGFLDTLQTLLEFKADV
*Coturnix japonica* MSGANPNLKDSTGFA-VIHDVARAGFLDTLQTLLEFHADV
*Cuculus canorus* TNGANPNLKDSTGFA-VIHDAAREGFLDTLQTLLEFKADV
*Cyanistes caeruleus* SNGANPNLKDSTGFA-VIHDVAREGFLDTLQTLLEFKADV
*Cyanoderma ruficeps* SNGANPNLKDSTGFA-VIHDVAREGFLDTLQTLLEFKADV
*Cygnus atratus* MSGANPDLKDSTGFA-VIHDVARAGFLDTLQTLLEFKADV
*Cygnus olor* MSGANPDLKDSTGFA-VIHDVARAGFLDTLQTLLEFKADV
*Dromaius novaehollandiae* SRGANPNLKDSTGFA-VIHDVARAGFLDTLQTLLEFKADV
*Dryobates pubescens* MSGANPNLKDRTGFA-VIHDVAREGFLDTLQTLLEFQADV
*Egretta garzetta* IHGANPNLKDSTGFA-VIHDVAREGFLDTLQTLLEFEADV
*Empidonax traillii* SKGANPNLKDSTGFA-IIHDVAREGFLDTLQTLLEFKADV
*Erythrura gouldiae* SNGANPNLKDSTGFA-VIHDVAREGFLDTLQTLLEFKADV
*Eudyptes chrysocome* INGADPNLKDSTGFA-VIHDVAREGFLDTLQTLLEFKADV
*Eudyptes chrysolophus* INGADPNLKDSTGFA-VIHDVAREGFLDTLQTLLEFKADV
*Eudyptes filholi* INGADPNLKDSTGFA-VIHDVAREGFLDTLQTLLEFKADV
*Eudyptes moseleyi* INGADPNLKDSTGFA-VIHDVAREGFLDTLQTLLEFKADV
*Eudyptes pachyrhynchus* INGADPNLKDSTGFA-VIHDVAREGFLDTLQTLLEFKADV
*Eudyptes robustus* INGADPNLKDSTGFA-VIHDVAREGFLDTLQTLLEFKADV
*Eudyptes schlegeli* INGADPNLKDSTGFA-VIHDVAREGFLDTLQTLLEFKADV
*Eudyptes sclateri* INGADPNLKDSTGFA-VIHDVAREGFLDTLQTLLEFKADV
*Eudyptula albosignata* INGADPNLKDSTGFA-VIHDVAREGFLDTLQTLLEFKADV
*Eudyptula minor* INGADPNLKDSTGFA-VIHDVAREGFLDTLQTLLEFKADV
*Eudyptula novaehollandiae* INGADPNLKDSTGFA-VIHDVAREGFLDTLQTLLEFKADV
*Eurypyga helias* MSGANPNLKDSTGFA-VIHDAAREGFLDTLQTLLEFKADV
*Falco cherrug* VSGANPNLKDSTGFA-VIHDVAREGFLDTLQTLLEFKADV
*Falco naumanni* VSGANPNLKDSTGFA-VIHDVAREGFLDTLQTLLEFKADV
*Falco peregrinus* VSGANPNLKDSTGFA-VIHDVAREGFLDTLQTLLEFKADV
*Falco rusticolus* VSGANPNLKDSTGFA-VIHDVAREGFLDTLQTLLEFKADV
*Falco tinnunculus* VSGANPNLKDSTGFA-VIHDVAREGFLDTLQTLLEFKADV
*Ficedula albicollis* SNGANPNLKDSTGFA-VIHDVAREGFLDTLQTLLEFKADV
*Gallus gallus* MSGANPNLKDSTGFA-VIHDVARAGFLDTLQTLLEFHADV
*Geospiza fortis* SNGANPNLKDSTGFA-VIHDVAREGFLDTLQTLLEFKADV
*Haliaeetus albicilla* LSGADPNLKDSTGFA-VIHDVAREGFADTLQALLEFQADV
*Haliaeetus leucocephalus* LSGADPNLKDSTGFA-VIHDVAREGFADTLQALLEFQADV
*Hirundo rustica* SNGANPNLKDSTGFA-VIHDVAREGFLDTLQTLLEFKADV
*Hypotaenidia okinawae* INGANPNLKDSTGFA-VIHDVAREGFLDTLQTLLEFKADV
*Junco hyemalis* SNGANPNLRDSTGFA-VIHDVAREGFLDTLQTLLEFEADV
*Lagopus leucura* MSGANPNLKDSTGFA-VIHDVARAGFLDTLQTLLEFHADV
*Lepidothrix coronata* SKGANPNLKDSTGFA-IIHDVAREGFLDTLQTLLEFKADV
*Leptosomus discolor* INGANPNLKDSTGFA-VIHDVAREGFLDTLQTLLEFKADV
*Lonchura striata domestica* SNGANPNLKDSTGFA-VIHDVAREGFLDTLQTLLEFKADV
*Malurus cyaneus samueli* SNGANPNLKDSTGFA-VIHDVAREGFLDTLQTLLEFKADV
*Manacus vitellinus* SKGANPNLKDSTGFA-IIHDVAREGFLDTLQTLLEFKADV
*Megadyptes antipodes antipodes* INGADPNLKDSTGFA-VIHDVAREGFLDTLQTLLEFKADV
*Meleagris gallopavo* MSGANPNLKDSTGFA-VIHDVARAGFLDTLQTLLEFHADV
*Melopsittacus undulatus* ISGANPNLKDSTGFA-VIHDVAREGFLDTLQTLLEFKADV
*Melospiza melodia maxima* SNGANPNLKDSTGFA-VIHDVAREGFLDTLQTLLEFEADV
*Merops nubicus* MNGANPNLKDSTGFA-VIHDVAREGFLDTLQTLLEFEADV
*Mesitornis unicolor* MSGADPDLRDGTGFA-VLPDAAREGFVDTLQTPPAFR---
*Molothrus ater* SNGANPNLKDSTGFA-VIHDVAREGFLDTLQTLLEFKADV
*Motacilla alba alba* SNGANPNLKDSTGFA-VIHDVAREGFLDTLQTLLEFKADV
*Neopelma chrysocephalum* SKGANPNLKDSTGFA-IIHDVAREGFLDTLQTLLEFKADV
*Nipponia nippon* INGADPNLKDSTGFA-VIHDAAREGFLDTLQTLLEFKADV
*Nothoprocta perdicaria* SSGADPDLKDRTGFA-VIHDVARAGFLDTLQTLLEFEADV
*Numida meleagris* MSGANPNLKDSTGFA-VIHDVARAGFLDTLQTLLEFHADV
*Onychostruthus taczanowskii* SNGANPNLKDSTGFA-VIHDVAREGFLDTLQTLLEFKADV
*Otus sunia* ISGANPNLKDSTGFA-VIHDVAREGFLDTLQTLLEFKADV
*Oxyura jamaicensis* MSGANPDLKDSTGFA-VIHDVARAGFLDTLQTLLEFKADV
*Parus major* SNGANPNLKDSTGFA-VIHDVAREGFLDTLQTLLEFKADI
*Passer montanus* SNGANPNLKDSTGFA-VIHDVAREGFLDTLQTLLEFKADV
*Patagioenas fasciata monilis* AHGANPDLRDSTGFA-VMHDVAREGFPDTLQALLEFQADV
*Pavo cristatus* MSGANPNLKDSTGFA-VIHDVARAGFLDTLQTLLEFHADV
*Pelecanus crispus* INGANPNLKDRTGFA-VIHDVAREGFLDTLQTLLEFKADV
*Phaethon lepturus* SHGANPNLKDSTGFA-VIHDVAREGFLDTLQTLLEFEADV
*Phalacrocorax carbo* INGANPNLKDSTGFA-VIHDVAREGFLDTLQTLLEFKADV
*Phasianus colchicus* MSGANPNLKDSTGFA-VIHDVARAGFLDTLQTLLEFHADV
*Pipra filicauda* SKGANPNLKDSTGFA-IIHDVAREGFLDTLQTLLEFKADV
*Pseudopodoces humilis* SNGANPNLKDSTGFA-VIHDVAREGFLDTLQTLLEFKADV
*Pterocles gutturalis* INGANPNLKDSTGFA-VIHDVAREGFLDTLQTLLEFKADV
*Pygoscelis adeliae* INGANPNLKDSTGFA-VIHDVAREGFLDTLQTLLEFKADV
*Pygoscelis antarcticus* INGANPNLKDSTGFA-VIHDVAREGFLDTLQTLLEFKADV
*Pygoscelis papua* INGANPNLKDSTGFA-VIHDVAREGFLDTLQTLLEFKADV
*Pyrgilauda ruficollis* SNGANPNLKDSTGFA-VIHDVAREGFLDTLQTLLEFKADV
*Serinus canaria* SNGANPNLKDSTGFA-VIHDVAREGFLDTLQTLLEFKADV
*Spheniscus demersus* INGADPNLKDSTGFA-VIHDVAREGFLDTLQTLLEFKADV
*Spheniscus humboldti* INGADPNLKDSTGFA-VIHDVAREGFLDTLQTLLEFKADV
*Spheniscus magellanicus* INGADPNLKDSTGFA-VIHDVAREGFLDTLQTLLEFKADV
*Spheniscus mendiculus* INGADPNLKDSTGFA-VIHDVAREGFLDTLQTLLEFKADV
*Strigops habroptila* ISGANPNLKDSTGFA-VIHDVAREGFLDTLQTLLEFKADV
*Strix occidentalis caurina* ISGANPNLKDSTGFA-VIHDVAREGFLDTLQTLLEFKADV
*Struthio camelus australis* SRGANPNLKDSTGFA-VIHDVARAGFLDTLQTLLEFKADV
*Sturnus vulgaris* SNGANPNLKDSTGFA-VIHDVAREGFLDTLQTLLEFKADV
*Taeniopygia guttata* SNGANPNLRDSTGFA-VIHDVAREGFLDTLQTLLEFKADA
*Tauraco erythrolophus* INGANPNLKDSTGFA-VIHDVAREGFLDTLQTLLEFKADV
*Tinamus guttatus* SSGADPDLKDRTGFA-VIHDVARAGFLDTLQTLLEFEADV
*Turdus rufiventris* SNGANPNLRDSTGFA-VIHDVAREGFLDTLQTLLEFKADV
*Turnix velox* IKGADPNLKDSTGFA-VIHDVAREGFLDTLQTLVEFQADI
*Tyto alba* VSGANPNLKDSTGFA-VIHDVAREGFLDTLQTLLEFKADV
*Zonotrichia albicollis* SNGANPNLRDSTGFA-VIHDVAREGFLDTLQTLLEFEADV
*Zosterops lateralis melanops* SNGANPNLRDRTGFA-VIHDAAREGFLDTLQTLLEFQADV

**reptiles**

*Alligator mississippiensis* LKGADPNLRDSTGFA-VIHDAARAGFLDTLQTLLEFKADV
*Alligator sinensis* LKGADPNLRDSTGFA-VIHDAARAGFLDTLQTLLEFKADV
*Anolis carolinensis* LRGADPDLRDAAGFS-VLHDAARAGFLDTLQTLLEFRADV
*Chelonia mydas* ITGANPDLKDSTGFA-VIHDAARAGFLDTLQTLLEFNADV
*Chelonoidis abingdonii* MTGANPDLKDSTGFA-VIHDAARAGFLDTLQTLLEFNADV
*Chelydra serpentina* ITGANPDLKDSTGFA-VIHDAARAGFLDTLQTLLEFNADV
*Chrysemys picta bellii* ISGANPDLKDSTGFA-VIHDAARAGFLDTLQTLLEFNADV
*Crocodylus porosus* QRGADPNLRDSTGFA-AIHDAARAGFLDTLQTLLEFKADV
*Crotalus adamanteus* LRGADPDLKDRTGFA-VLHDAARAGFLDTLQILLEFQADV
*Crotalus tigris* LRGADPDLKDRTGFA-VLHDAARAGFLDTLQILLEFQADV
*Dermochelys coriacea* ITGANPDLKDSTGFA-VIHDAARAGFLDTLQTLLEFNADV
*Gavialis gangeticus* LRGADPNLRDSTGFA-AIHDAARAGFLDTLQTLLEFKADV
*Gekko japonicus* EKGADPNLKDSAGFA-VLHDAARAGFLDTLQTLLDSQADV
*Gopherus agassizii* MTGANPDLKDSTGFA-VIHDAARAGFLDTLQTLLEFNADV
*Gopherus evgoodei* MTGANPDLKDSTGFA-VIHDAARAGFLDTLQTLLEFNADV
*Lacerta agilis* LRGADPNLRDRTGFA-VLHDAARAGFLDTLQTLLEFKADV
*Laticauda laticaudata* LRGADPDLKDRTGFA-VLHDAARAGFLDTLQTLLEFRADV
*Mauremys mutica* MTGANPDLKDSTGFA-VIHDVARAGFLDTLQTLLEFNADV
*Mauremys reevesii* MTGANPDLKDSTGFA-VIHDVARAGFLDTLQTLLEFNADV
*Naja naja* AAGADPDLKDRTGFA-VLHDAARAGFLDTLQTLLEFRADV
*Notechis scutatus* LRGADPDLKDRTGFA-VLHDAARAGFLDTLQTLLEFGADV
*Ophiophagus hannah* -----PDLKDRTGFA-VLHDAARAGFLDTLQTLLEFRADV
*Pantherophis guttatus* LRGADPDLKDRTGFA-VLHDAARAGFLDTLQTLLEFQADV
*Pelodiscus sinensis* NAGANPDLKDSTGFA-VIHDAARAGFLDTVQTLLEFNADV
*Pelusios castaneus* TRGADPDLKDRTGFA-VIHDVARAGFLDTLETLLEFQADV
*Podarcis muralis* RRGADPDLRDRTGFA-VLHDAARAGFLDTLQTLLEFNADV
*Pogona vitticeps* MRGADPDLKDGTGFA-VLHDVARAGFLDTLQTLLEFHADV
*Protobothrops mucrosquamatus* LRGADPDLKDRTGFA-VLHDAARAGFLDTLQTLLEFQADV
*Pseudonaja textilis* LRGAHPDLKDRTGFA-VLHDAARAGFLDTLQTLLEFGADV
*Python bivittatus* LRGADPDLKDRTGFA-VLHDAARAGFLDTLQTLLEFQADV
*Salvator merianae* LRGADPDLKDATGFA-ILHDVARAGFLDTLQTLLEFQADV
*Sceloporus undulatus* LGGADPDLKDGRGFA-VLHDAARAGFLDTLQTLLEFQADV
*Sphenodon punctatus* ARGAHPDLKDSAGFA-VIHDTARAGFLDTLQTLLEFRADV
*Terrapene carolina triunguis* ITGANPDLKDSTGFA-VIHDAARAGFLDTLQTLLEFNADV
*Thamnophis elegans* LRGADPDLKDRTGFA-VLHDAARAGFLDTLQILLEFQADV
*Thamnophis sirtalis* LRGADPDLKDRTGFA-VLHDAARAGFLDTLQILLEFQADV
*Trachemys scripta elegans* ITGANPDLKDSTGFA-VIHDAARAGFLDTLQTLLEFNADV
*Varanus komodoensis* LRGADPDLKDGTGFA-VLHDAARGGFLDTLQTLLEFQADV
*Zootoca vivipara* LRGADPDLKDRTGFA-VLHDAARAGFLDTLQTLLEFKADV

**amphibians**

*Ambystoma andersoni* DNGADPNLRDPTGFA-VLHDAAREGFLDTIRTLVVFRAEV
*Ambystoma maculatum* --GADPNLRDPTGFA-VLHDAAREGFLDTIQTLLVFRAEV
*Ambystoma mexicanum* DNGADPNLKDPTGFA-VLHDAAREGFLDTIRTLVVFRAEV
*Ambystoma velasci* DNGADPNLKDPTGFA-VLHDAAREGFLDTIRTLVVFRAEV
*Bufo bufo* RRGADPNLQDRSGFS-VLHDTARAGFLDTLQILLDFQADV
*Bufo gargarizans* RRGADPNLQDSSGFS-VLHDTARAGFLDTLQILLDFQADV
*Engystomops pustulosus* RRGADPNLRDSSGFS-VLHDTARAGFQDTMQILLDFHADV
*Geotrypetes seraphini* YKGADPNLKDRTGFA-VIHDAARAGFLDTVKTLLQFQADV
*Hymenochirus boettgeri* SQGADPNLRDRTGFS-VLHDAARAGFHDTVQTLLDFQADA
*Leptobrachium leishanense* RRGANPNLRDSTGFS-VLHDAARAGFQDTVQTLLHFQADA
*Microcaecilia unicolor* HEGADPNVKDRTGFA-VIHDAARAGFLDTVQTLLQFQADV
*Nanorana parkeri* GRGADPNLQDSCGFS-VLHDTARAGFCDTMRILLDFHADV
*Rana temporaria* GKGADPNLQDSCGFS-VVHDTARAGFCDTMRILLDFHVDV
*Rhinatrema bivittatum* HRGADPNLRDRTGFA-VMHDAARAGFLDTVQTLLQFQADV
*Xenopus laevis 1* RQGADPNMRDRTGFS-VVHDAARAGFQDTLETFFDFQADA
*Xenopus laevis 2* SQGADPNLRDRTGYS-VLHDAARAGFQDTLETLFDFQADA
*Xenopus tropicalis* SQGADPNLRDRTGYS-VLHDAARAGFQDTLKTLFDFQADA

### Figure S2g

| | |
|---|---|
| **Cdkn2a** | |
| *Homo sapiens* | LHGAEPNCADPAT-LTRPVHDAAREGFLDTLVVLHRAGARL |
| **Cdkn2c** | |
| **bony fish** | |
| *Acanthochromis polyacanthus* | WAGADVTVRDPVLNLT-VTHDAARAGFGDSVCALLNYGADV |
| *Acanthopagrus latus* | QAGADPNRPDPACGLT-VTHDAAREGFIDSVRVLLKHKAQV |
| *Acipenser ruthenus* 1 | RAKANPNERDPLLGLT-VAHDAARDGFEDTLRVLVDYGADV |
| *Acipenser ruthenus* 2 | RAKANPNERDPLFGLT-VAHDAARDGFEDTLRVLVDYGADV |
| *Albula glossodonta* | QANADPNVRDPLGGLT-ALHDTARDGYADTLQVLVDHGADV |
| *Alosa alosa* | RANADPNVRDPVLGLT-IAHDASRDGFLDTLRVLVHHGADV |
| *Alosa sapidissima* | RANADPNVRDPVLGLT-IAHDASRDGFLDTLRVLVHHGADV |
| *Ameiurus melas* | RAGADPNVPDPSGGLT-VTHDAARDGHADTLQVLLSYGADV |
| *Amphilophus citrinellus* | NAGANPNVRDPSCGLT-VTHDAAREGFLDSVSILIKGGADV |
| *Amphiprion ocellaris* | WAGADVTVRDPVLNLT-VTHDAARAGFGDSVCALLNYGANV |
| *Amphiprion percula* | W-GADVTVRDPVLNLT-VTHDAARAGFGDSVCALLNYGANV |
| *Anabarilius grahami* | RAGADPNVRDPILGLT-ISHDAARDGYLDTLQVLAQNGADV |
| *Anabas testudineus* | DAGADLSRCDPVLSLS-VTHDAAREGFVDTVRALMRHGADV |
| *Anarrhichthys ocellatus* | RAGADPDLRDPDCSLT-VMHDAARAGYVDSVRVLLDYSADT |
| *Anguilla anguilla* | QAKADPNVRDPVGGLT-VLHDAARDGYADTLQVLVNHGADV |
| *Archocentrus centrarchus* | NAGANPNVRDPSCGLT-VTHDAAREGFLDSVSILIKGGADV |
| *Astatotilapia calliptera* 1 | QAGANPNVPDPSCGLT-VTHDAAREGFVDSVRVLLNYGADV |
| *Astatotilapia calliptera* 2 | QAGANPNVPDPSCGLT-VTHDAAREGFVDSVRVLLNYGADV |
| *Astyanax mexicanus* | KKGADPNLRDPVKDLT-ISHDVAREGHADTLAVLLTYVADV |
| *Austrofundulus limnaeus* | AAGANPNERDPLLRLT-VTHDAAREGFVETVRALIEYGADV |
| *Bagarius yarrelli* | RAGADPNVHDVIGRLT-VTHDAARDGYVDMLQVLLRYGANV |
| *Betta splendens* | RNGADPNVRDPLLGLC-VIHDAAREGFVDTVRVLIGHGADV |
| *Boleophthalmus pectinirostris* | DRGADPTVRDPIFGLT-VLHDAAREGFEDTVRVLLDHGADV |
| *Caranx melampygus* | EVGANPDRLDPVCSLT-VTHDAAREGFVDTVRVLMEHGANI |
| *Carassius auratus* 1 | RAGADPNARDPILRLT-ISHDAARDGYLDTLRVLAQNGADV |
| *Carassius auratus* 2 | RAGADPNARDPILRLT-VSHDAARDGYLDTLRVLAQNGADV |
| *Carassius auratus* 3 | RAGADPNARDPILRLT-VSHDAARDGYLDTLRVLAQNGADV |
| *Channa argus* | VARADPNIRDPVLSLT-VTHDAAREGFADAVRALIQHGADV |
| *Chanos chanos* | TANADPNVRDPIRGLT-ITHDAARDGYIDTLQVLVEHGADV |
| *Cheilinus undulatus* | EAGADPNMPDPVLNLT-VTHDAARDGHKDTVRVLLDYGANP |
| *Chelmon rostratus* | EAGAHPNLRDPVCGLT-VTHDAAREGFADTVRALVDRGADP |
| *Clarias magur* | CAGANPNIRDPSGGLT-VTHDAARDGHADTLQVLLKYGADV |
| *Clupea harengus* | RANADPNVRDPVLGLT-VAHDASRDGFLDTLNVLVHHGADV |
| *Collichthys lucidus* 1 | KAGADPDRRDQVCGLT-VTHDAARQGFVDSVRLLLDYRADP |
| *Collichthys lucidus* 2 | KAGADPDRRDQVCGLT-VTHDAARQGFVDSVRLLLDYRADP |
| *Colossoma macropomum* | QANADPNARDPVKRLT-ITHDAARDGYVDMLEVLVTYGADV |
| *Coregonus clupeaformis* 1 | RANANPNVRDHVRGLT-ITHDAARDGYVDTLRVLMDYGADV |
| *Coregonus clupeaformis* 2 | RANANPNVRDHVRGLT-ITHDAARDGYADTLRILMVHGADV |
| *Coryphaenoides rupestris* | -----PDRRDPRRGLT-VAHDAARDGFGDTVRVLVDCGADV |
| *Cottoperca gobio* | EAGADTSVRDPVLNLT-VMHDAARAGYIDSVRMLMDHGADI |
| *Cyclopterus lumpus* | RAGADPNLRDPVCGLT-VMHDAAREGFIESVRALVDYGADL |
| *Cynoglossus semilaevis* | NKGANPDLRDPVLGLT-VTHDAAREGFDDTVSVLLQHGANP |
| *Cyprinodon tularosa* 1 | DKGADPNALDPVRHLS-VTHDAAREGFVDTVRVLIDHGADV |
| *Cyprinodon tularosa* 2 | DKGADPNALDPVQHLS-VTHDAAREGFVDTVRVLIHHGADV |
| *Cyprinodon variegatus* | DKGADPNALDPVQYLS-VTHDAAREGFVDTVRVLIDHGADV |
| *Cyprinus carpio* 1 | RAGADPNARDPILRLT-VSHDAARDGYLDTLRVLAQNGADV |
| *Cyprinus carpio* 2 | RAGADPNARDPILRLT-VSHDAARDGYLDTLRVLAQNGADV |
| *Danio rerio* | QAGADPNVRDPILGLT-VIHDAARDGYLDTLHVLAQNGADV |
| *Danionella translucida* | RACADPNVRDPILGLT-ITHDAARDGYLDTLQVLAQNGADV |
| *Denticeps clupeoides* | LRGADADVRDPVAGLT-VAHDAARDGHADTLRVLVGHKADV |
| *Dicentrarchus labrax* | VAGADPSVRDPVLSLT-VMHDAAREGFADSVRVLVDHGADT |
| *Echeneis naucrates* | EWGADPNLRDPDCGLT-VTHDAAREGFVDTVRLLLEYGADA |
| *Electrophorus electricus* | LAQADPNLRDPIGGLT-ITHDAARDGYVDTLQVLVNYGANV |
| *Epinephelus coioides* | EAGADPNIRDPVLRLT-VTHDAAREGFINSVHELVDHGADI |
| *Epinephelus lanceolatus* | EAGADPNIRDPVLRLT-VTHDAAREGFINSVRELVDHGADI |
| *Erpetoichthys calabaricus* | RANANPNEQDISLGVT-IAHDAAREGFLETLMVLTENGADV |
| *Esox lucius* | KANANPNVRDQVKGLT-VCHDAARDGYADTLRVLVDHGADV |
| *Etheostoma cragini* | DAGSDPNLREPFCGLT-VTHDAAREGFIDSLRVLVNHGADL |
| *Etheostoma spectabile* | EAGSDPNVREPFCGLT-VTHDAAREGFLDSLRVLVNHGADL |
| *Fundulus heteroclitus* | EAGAEPNARDDVLKLT-VVHDAARDGFLEAVRALADHGADV |
| *Gambusia affinis* | AAGADPNARDQILNLT-VTHDAAREGFVETVRVLINHGADV |
| *Gasterosteus aculeatus aculeatus* | RAGADPNLRDPVLGLT-VMHDAAREGFADSVRVLLAYGADP |
| *Girardinichthys multiradiatus* | EAGADPNVRDMILKLT-VTHDAAREGFLETVRVLVDRGADV |
| *Gouania willdenowi* | DAGADPNVRDPVRNLT-VMHDMARGGHDDTLQILLAYGADV |
| *Gymnodraco acuticeps* | SAGADFDIRDPVLGLT-PLHDAARAGYIGSVEKLVQCGSDV |
| *Haplochromis burtoni* 1 | QAGANPNVPDPSCGLT-VTHDAAREGFVDSVRVLLNYGANV |
| *Haplochromis burtoni* 2 | QAGANPNVPDPSCGLT-VTHDAAREGFVDSVRVLLNYGANV |
| *Haplochromis burtoni* 3 | QAGANPNVPDPSCGLT-VTHDAAREGFVDSVRVLLNYGANV |
| *Hippocampus comes* | DAEADPNLRDPVLGLT-VTHDAAREGFADTVRALLAHGADV |
| *Hippoglossus hippoglossus* | GGGASADVRDPDSGRT-VTHDAAREGSVDTVRALMRAEANV |
| *Hippoglossus stenolepis* | GGGASPDVRDPDSGRT-VTHDAAREGSVDTVRALMRAEANV |
| *Hucho hucho* 1 | RANARPNVRDHVRGLT-ITHDAARDGYVDTLRVLIDHGADV |
| *Hucho hucho* 2 | RANANPNVRDHVRGLT-ITHDAARDGYADTLRVLMDHGADV |
| *Ictalurus punctatus* | RAGADPNVPDPSGGLT-VTHDAARDGHADTLQVLLSYGADV |
| *Kryptolebias marmoratus* | AAGANPDARDPLLNLT-VTHDAAREGFVDTVRALIAHGADV |
| *Labeo rohita* | QAGADPNERDPILQLT-VSHDAARDGYLDTLQVLAQNGADV |
| *Labrus bergylta* | VAGADPNVPDPVLNLT-VTHDAAREGFIDSVRVLVDHGANT |
| *Larimichthys crocea* | KAGADPDRRDQVCGLT-VTHDAARQGFVDSVRLLLDYRADP |
| *Lates calcarifer* | DAGANPNLRDPTCGLT-VTHDAAREGFVDTVRVLISHGADV |
| *Latimeria chalumnae* | HAGADPNLKDR-TGYT-VVHDTARAGFCDTLQILLQYKADV |
| *Lepisosteus oculatus* | RANADPNVRDPLGGLT-VTHDTARDGYEDTLRVLVEHGADV |
| *Liparis tanakae* | RAGSDPNLRDPVCGLT-VMHDAAREGFVESVRALVDYGADL |
| *Lucifuga dentata* | RAGANPNLRDPICSLT-VTHDAARDGFLDTVRVLVEHGAEV |
| *Mastacembelus armatus* | KKGANPNLRDPVLNLT-VTHDAARDGYVDTVSALLQNGADA |
| *Maylandia zebra* 1 | QAGANPNVPDPSCGLT-VTHDAAREGFVDSVRVLLNYGADV |
| *Maylandia zebra* 2 | QAGANPNVPDPSCGLT-VTHDAAREGFVDSVRVLLNYGADV |
| *Megalobrama amblycephala* | RAGADPNVRDPILGLT-ISHDAARDGYLDTLQVLAQNGADV |
| *Megalops atlanticus* | KANADPNVRDPLGGLT-VLHDTARDGYADTLRVLVEYGADV |
| *Megalops cyprinoides* | KANADPNVRDPLGGLT-VLHDTARDGYADTLRVLVEYGADV |
| *Melanotaenia boesemani* | EAGADPNMRDRVLNLT-VTHDAAREGFVESVGLLIDHSANV |
| *Micropterus dolomieu* | LVGADINVRDPACGLT-VTHDAAREGFIDSVRVLVERGADT |
| *Micropterus salmoides* | LVGADINVRDPACGLT-VTHDAAREGFIDSVRVLVDRGADT |
| *Mola mola* | LAGADPNLRDPACGLT-VTHDAAREGFADSVRVLVEHGADV |
| *Monopterus albus* | EAGADPNLRDPVLSLT-VVHDAAREGFVDTACVLMQHGANV |
| *Morone saxatilis* | VAGADPSARDPVLSLT-VMHDAAREGFADSVRVLVDHGADT |
| *Mugil cephalus* | EAGADPNVPDPSCGLT-VTHDAAREGFLETVRTLVEYGADV |
| *Myripristis murdjan* | AAGADPNARDPTCGLT-VTHDAAREGFLDMVHVLVEHRADV |
| *Nematolebias whitei* | AAGADPNVRDQTSKLT-VMHDAAREGFAETVRTLLAHGADA |
| *Neogobius melanostomus* | RLGANPDARDPIHGIT-IIHDAAREGYEESVRVLLAHGVDV |
| *Neolamprologus brichardi* | QAGANPNVPDPSCGLT-VTHDAAREGFVDSVRVLLKYGADV |
| *Nibea albiflora* | KAGADPDRRDPVCGLT-VTHDAAREGFVDSVRLLLDYKADP |
| *Nothobranchius furzeri* | QNGANPNKRDPVLNLT-VTHDAAREGFVDSVHALIDNGADV |
| *Nothobranchius kuhntae* | QNGANPNKRDPVLNLT-VTHDAAREGFVDSVHALIDNGADV |
| *Notolabrus celidotus* | VAGADPNVPDPVLNLT-LTHDAAREGFIHTVRVLVNHGADL |
| *Notothenia coriiceps* | LAGADFNIRDPVLGLT-PLHDAARAGYIGSVEKLVQYGSDV |
| *Oncorhynchus gorbuscha* 1 | RANANPNVRDHVRGLT-ITHDAARDGYVDTLRVLIDHGADV |
| *Oncorhynchus gorbuscha* 2 | RANANPNVRDHVRGLT-ITHDAARDGYADTLHVLMDHGADV |
| *Oncorhynchus keta* 1 | RANANPNVRDHVRGLT-ITHDAARDGYVDTLRVLIDHGADV |
| *Oncorhynchus keta* 2 | RANANPNVRDNVRGLT-ITHDAARDGYADTLHVLMDHGADV |
| *Oncorhynchus kisutch* 1 | RANANPNVRDHVRGLT-ITHDAARDGYVDTLRVLIDHGADV |
| *Oncorhynchus kisutch* 2 | RANANPNVRDHVRGLT-ITHDAARDGYADTLHVLMDHGADV |
| *Oncorhynchus mykiss* 1 | RANANPNVRDHVRGLT-ITHDAARDGYVDTLRVLIDHGADV |
| *Oncorhynchus mykiss* 2 | RANANPNVRDHVRGLT-ITHDAARDGYADTLHVLMDHGADV |
| *Oncorhynchus nerka* 1 | RANANPNVRDHVRGLT-ITHDAARDGYVDTLRVLIDHGADV |
| *Oncorhynchus nerka* 2 | RANANPNVRDHVRGLT-ITHDAARDGYADTLHVLMDHGADV |
| *Oncorhynchus tshawytscha* 1 | RANANPNVRDHVRGLT-ITHDAARDGYVDTLRVLIDHGADV |
| *Oncorhynchus tshawytscha* 2 | RANANPNVRDHVRGLT-ITHDAARDGYADTLHVLMDHGADV |
| *Onychostoma macrolepis* | RAGADPNARDPILRLT-VSHDAARDGYLDTLQVLAQNGADV |
| *Oreochromis aureus* 1 | QAGANPNVPDPSCGLT-VTHDAAREGFVDSVRALLTYGADV |
| *Oreochromis aureus* 2 | QAGANPNVPDPSCGLT-VTHDAAREGFVDSVRALLTYGADV |
| *Oreochromis niloticus* 1 | QAGANPNVPDPSCGLT-VTHDAAREGFVDSVRALLTYGADV |
| *Oreochromis niloticus* 2 | QAGANPNVPDPSCGLT-VTHDAAREGFVDSVRALLTYGADV |
| *Oryzias javanicus* | QAGADPNARDPVLKLT-VTHDAAREGFADTVRALITHGADV |
| *Oryzias latipes* | HAGADPNARDPVLKLT-VTHDAAREGFVDTVRVLIRHSADV |
| *Oryzias melastigma* | HAGADPNVRDHVLKLT-VTHDAAREGFTDTVRVLINHDADV |
| *Oryzias sinensis* | HAGADPNARDRVLKLT-VTHDAAREGFVDTVRVLTERSADV |
| *Pangasianodon hypophthalmus* | RAGADPNVPDPSGGLT-VMHDAAREGHADTLQVLLRYGADA |
| *Parambassis ranga* | EAGADPNLPDPACGLT-VTHDAAREGFIDSVRTLVKHGADV |
| *Paramormyrops kingsleyae* | EAKADPNKRDPVKDLT-IAHDAARGGYLETLQVLVKYGADV |
| *Perca flavescens* | EAGSDPNVRDPFCGLT-VTHDAAREGFIDSVRVLVNHGADL |
| *Perca fluviatilis* | EAGSDPNVRDPFCGLT-VTHDAAREGFIDSVRVLVNHGADL |
| *Periophthalmus magnuspinnatus* | RAGADPSVRDPVCGLT-VLHDAAREGFEDTVRVLLDHGADL |
| *Pimephales promelas* | LAGADPNARDPILGLT-VSHDAARDGYLDTLRVLVENGADV |
| *Plectropomus leopardus* | EAGADPNIRDPVHNLT-VTHDAAREGFVDSVRALVDHGADI |
| *Poecilia formosa* | EAGADPNARDPTLNLT-VTHDAAREGFVETVRVLINHGAEV |
| *Poecilia latipinna* | EAGADPNARDPTLNLT-VTHDAAREGFVETVRVLINHGAEV |
| *Poecilia mexicana* | EAGADPNARDPTLNLT-VTHDAAREGFVETVRVLINHGAEV |
| *Poecilia reticulata* | AAGADPNAQDPTLNLT-VTHDAAREGFVETVRVLVNHGAEV |
| *Poeciliopsis prolifica* | AAGADPNARDQVLNLT-VTHDAARAGFVETVRALVNHGADV |
| *Polyodon spathula* 1 | RAKANPNERDPLWGLT-VAHDAARDGFEDTLRVLVDYGADV |
| *Polyodon spathula* 2 | RARANPNERDPLWGLT-VAHDAARDGFEDTLRVLLDYGADV |
| *Polypterus senegalus* | RANANPNEQDTSLGVT-IAHDAAREGFLETLMVLTENGADV |
| *Pseudochaenichthys georgianus* | SVGADFNIRDPVLGLT-PLHDAARAGYIGSVEKLVQCGSDV |
| *Pungitius pungitius* | RAGADPNLRDPVLALT-VMHDAAREGFADSVRVLLAYRGDP |
| *Puntigrus tetrazona* | RAGADPNARDPILQLT-VSHDAARDGYLDTLRVLAQNGADV |
| *Pygocentrus nattereri* | QAKADPNARDPVKRLT-IAHDAARDGYVDMLEVLVTYGADV |
| *Salarias fasciatus* | AAGANPNLQDPACGLT-VLHDAAVTGFRCTVRALLDHGADA |
| *Salmo salar* 1 | RANANPNVRDHVRGLT-ITHDAARDGYVDTLRVLIDHGADV |
| *Salmo salar* 2 | RANANPNVRDHVRGLT-ITHDAARDGYADTLHVLMDHGADV |
| *Salmo trutta* 1 | RANANPNVRDHVRGLT-ITHDAARDGYVDTLRVLIDHGADV |
| *Salmo trutta* 2 | RANANPNVRDHVRGLT-ITHDAARDGYADTLHVLMDHGADV |
| *Salvelinus alpinus* 1 | RANANPNVRDHVRGLT-ITHDAARDGYVDTLRVLIDHGADV |
| *Salvelinus alpinus* 2 | RANANPNVRDHVRGLT-ITHDAARDGYADTLHVLMDHGADV |
| *Salvelinus namaycush* 1 | RANANPNVRDHVRGLT-ITHDAARDGYVDTLRVLIDHGADV |
| *Salvelinus namaycush* 2 | RANANPNVRDHVRGLT-ITHDAARDGYADTLHVLMDHGADV |
| *Sander lucioperca* | EAGSDPNVRDPFCGLT-VTHDAAREGFIDAVRVLVNHGADL |
| *Scatophagus argus* | VQGADPNLRDPACGLT-VMHDAAREGFVDSVRVLLDHEADA |
| *Scleropages formosus* | KANADPNVCDA-LGLT-VTHDAARDGYLDTLRVLLEHGADV |
| *Scophthalmus maximus* | AAGARPDVRDPVLGLT-VTHDAARGGFAHTVRALLQHGANV |
| *Sebastes umbrosus* | LAGANPNLPDRCLSLT-AAHDGAREGYIDSMRVLVEHGADV |
| *Seriola dumerili* | EVGANPNIRDPACSLT-VMHDAAREGFVDTVRVLMDHGANI |
| *Seriola lalandi dorsalis* | EVGANPNIRDPACSLT-VMHDAAREGFVDTVRVLMDHGANI |
| *Silurus meridionalis* | RAGADPNVRDRCKQLT-LMHDAARDGHADTVHVLLQYGADA |
| *Simochromis diagramma* 1 | QAGANPNVPDPSCGLT-VTHDAAREGFVDSVRVLLNYGADV |
| *Simochromis diagramma* 2 | QAGANPNVPDPSCGLT-VTHDAAREGFVDSVRVLLNYGADV |
| *Siniperca chuatsi* | AAGANPNVRDPVLSLT-VTHDAARQGFIDSVRALVDGGADT |
| *Sinocyclocheilus anshuiensis* 1 | RAGADPNARDPILRLT-VSHDAARDGYLDTVRVLAQNGADV |
| *Sinocyclocheilus anshuiensis* 2 | RAGADPNTRDPILRLT-VSHDAARDGYLDTLRVLAQNGADV |
| *Sinocyclocheilus grahami* 1 | GAGADPNARDPILRLT-VSHDAARDGYLDTLRVLAQNGADV |
| *Sinocyclocheilus grahami* 2 | RAGADPNTRDPILRLT-VSHDAARDGYLDTLRMLAQNGADV |
| *Sinocyclocheilus grahami* 3 | RAGADPNTRDPILRLT-VSHDAARDGYLDTLRMLAQNGADV |
| *Sinocyclocheilus rhinocerous* 1 | LAGADPNARDPILRLT-VSHDAARDGYLDTLRVLAQNGADV |
| *Sinocyclocheilus rhinocerous* 2 | RAGADPNARDPILRLT-VSHDAARDGYLDTLRVLAQNGADV |
| *Solea senegalensis* | EAGANPDVRDPVCGLT-VTHDAARDGWIDAVRLLIDHKADP |
| *Sparus aurata* | VAGADPDIPDPACGLT-VTHDAARDGFMDTVRVLVKHRAKV |
| *Sphaeramia orbicularis* | DAGADPTVRDPVLSLT-VLHDAAREGFVDTVRVLADHRSVD |
| *Stegastes partitus* | QAGADVTVRDPVLDLT-VMHDAARAGFRDSVRALLNYEADV |
| *Syngnathus acus* | EAGADPNLRDPVLGLT-VTHDAAREGFADTVRTLLAQKADA |
| *Tachysurus fulvidraco* | RAGADPNVRDPCGGLT-VTHDAARDGHTDTLQVLLRNGADV |
| *Takifugu flavidus* | QAGANPDVADPACGLT-LLHDTSREGFDDSVQVLLEHGADA |
| *Takifugu rubripes* | QAGANPDVADPACGLT-LLHDTSREGFDDSVRVLLEHGADA |
| *Tetraodon nigroviridis* | LAGAEPNVADPACDLT-VLHDASRDGFAASVQLLLQHGADA |
| *Thalassophryne amazonica* | KAGGNPHVRDHVCGLM-VIHDAAREGFVETVRVLIEYGADV |
| *Thunnus albacares* | EAGANPNLRDPACGLT-VTHDAAREGFVDTVRVLLDHGADV |
| *Thunnus maccoyii* | EAGANPNLRDPACGLT-VTHDAAREGFVDTVRVLLDHGADV |
| *Toxotes jaculatrix* | QKGANPNVRDPVCELT-VIHDAARDGFADTVRVLIDHGADA |
| *Trematomus bernacchii* | SAGAEFNIRDPVLRLT-PLHDAARAGYIGSVQKLVLYGADV |
| *Triplophysa tibetana* | LAGADPNARDPIGGLA-LCHDAARDGFLDTLQVLGQNGADV |
| *Xiphias gladius* | AAGANPDVRDPVLSLT-VTHDAAREGFVDTVSVLMRHGANV |
| *Xiphophorus couchianus* | AAGADPNARDHILNLT-VTHDAAREGFAETVRELVNHGADV |
| *Xiphophorus hellerii* | AAGADPNARDHILNLT-VTHDAAREGFAETVRELVNHGADV |
| *Xiphophorus maculatus* | AAGADPNARDHILNLT-VTHDAAREGFAETVRELVNHGADV |
| **cartilaginous fish** | |
| *Amblyraja radiata* | KAGAKPNQQDH-GGFT-PAHDAAREGFLDTLKTLVDFGANV |
| *Callorhinchus milii* | KAGAKPNQQDS-NGFT-PAHDAAREGFVDTLRVLVDSGADV |
| *Carcharodon carcharias* | KAGAKPNQQDR-GGFT-PAHDAAREGFLDTLKTLVDFGANV |
| *Chiloscyllium plagiosum* | KAGAKPNQQDH-SGFT-PAHDAAREGFLDTLITLVDFGANV |
| *Hydrolagus affinis* | KAGAKPNQPDA-NGFT-PAHDAAREGFVETLRVLVDSGANV |
| *Rhincodon typus* | KAGAKPNQQDR-GGFT-PAHDAAREGFLDTLITLVDFGANV |
| *Scyliorhinus canicula* | KAGAKPNQQDP-GGYA-PAHDVAREGFLDTLKILVDFGANV |
| *Stegostoma fasciatum* | KAGAKPNEQDR-SGFT-PAHDAAREGFLDTLITLVDFGANV |

**Figure S2h**

**Cdkn2a**

| | |
|---|---|
| *Homo sapiens* | LHGAEPNCADPATLTRPVHDAAREGFLDTLVVLHRAGARL |

**Cdkn2d**

**Mammals**

**<Placentarians>**

| | |
|---|---|
| *Acinonyx jubatus* | KQGASPNVQDTS-GTTPAHDAARTGFLDTLKVLVEHGADV |
| *Ailuropoda melanoleuca* | KQGASPNVQDTT-GTTPAHDAARTGFLDTLKVLVEHGADV |
| *Aotus nancymaae* | KQGASPNVQDTF-GTSPVHDAARTGFLDTLKVLVEHGADV |
| *Artibeus jamaicensis* | KQGASPNVQDAS-GTTPAHDAARTGFLDTLKVLVEHGADV |
| *Arvicanthis niloticus* | KQGASPNVQDAS-GTSPVHDAARTGFLDTLKVLVEHGADV |
| *Arvicola amphibius* | KQGASPNVQDAS-GTSPVHDAARTGFLDTLKVLVEHGADV |
| *Balaenoptera acutorostrata scammoni* | KQGASPNVQDAS-GTTPAHDAARTGFLDTLKVLVEHGADV |
| *Balaenoptera musculus* | KQGASPNVQDAS-GTTPAHDAARTGFLDTLKVLVEHGADV |
| *Bison bison bison* | KQGASPNVQDAS-GTTPAHDAARTGFLDTLKVLVEHGADV |
| *Bos indicus* | KQGASPNVQDAS-GTTPAHDAARTGFLDTLKVLVEHGADV |
| *Bos mutus* | KQGASPNVQDAS-GTTPAHDAARTGFLDTLKVLVEHGADV |
| *Bos taurus* | KQGASPNVQDAS-GTTPAHDAARTGFLDTLKVLVEHGADV |
| *Bubalus bubalis* | KQGASPNVQDAS-GTTPAHDAARTGFLDTLKVLVEHGADV |
| *Callithrix jacchus* | KQGASPNVQDTS-GTSPIHDAARTGFLDTLKVLVEHGADV |
| *Callorhinus ursinus* | KQGASPNVQDTT-GTTPAHDAARTGFLDTLKVLVEHGADV |
| *Camelus bactrianus* | KQGASPNVQDAS-GTTPAHDAARTGFLDTLKVLVDHGADV |
| *Camelus dromedarius* | KQGASPNVQDAS-GTTPAHDAARTGFLDTLKVLVDHGADV |
| *Camelus ferus* | KQGASPNVQDAS-GTTPAHDAARTGFLDTLKVLVDHGADV |
| *Canis lupus familiaris* | KQGASPNVQDAT-GTTPAHDAARTGFLDTLKVLVEHGADV |
| *Capra hircus* | KQGASPNVQDAS-GTTPAHDAARTGFLDTLKVLVEHGADV |
| *Carlito syrichta* | KQGASPNVQDST-GASPVHDAARTGFLDTLKVLMEHGADV |
| *Castor canadensis* | KQGASPNVQDDS-GTSPIHDAARTGFLDTLKVLVEHGADV |
| *Cavia porcellus* | KQGASPNVQDAS-GTSPLHDAARTGFLDTVKILVEHSADV |
| *Cebus imitator* | KQGASPNVQDTT-GTSPVHDAARTGFLDTLKVLVEHGADV |
| *Ceratotherium simum simum* | KQGASPNIQDAS-GTTPAHDAARTGFLDTLRVLVEHGADV |
| *Cercocebus atys* | KQGASPNVQDTS-GTSPVHDAARTGFLDTLKVLVEHGADV |
| *Cervus canadensis* | KQGASPNVQDAS-GTTPAHDAARTGFLDTLKVLVEHGADV |
| *Cervus elaphus* | KQGASPNVQDAS-GTTPAHDAARTGFLDTLKVLVEHGADV |
| *Cervus hanglu yarkandensis* | KQGASPNVQDAS-GTTPAHDAARTGFLDTLKVLVEHGADV |
| *Chinchilla lanigera* | KHGASPNVQDAS-GTSPLHDAARTGFLDTLKVLVEHGADV |
| *Chlorocebus sabaeus* | KQGASPNVQDTS-GTSPVHDAARTGFLDTLKVLVEHGADV |
| *Choloepus didactylus* | KQGATPNIQDAS-GTTPAHDAARTGFLDTLKVLVEHGADV |
| *Chrysochloris asiatica* | KQGACPNVQDIS-GTTPAHDAARTGFLDTLKVLVEHGADV |
| *Colobus angolensis palliatus* | KQGASPNVQDTS-GTSPVHDAARTGFLDTLKVLVEHGADV |
| *Condylura cristata* | KQGASPNVQDAS-GTTPAHDAARTGFLDTLSILVEHGADV |
| *Cricetulus griseus* | KQGASPNVQDAS-GTSPVHDAARTGFLDTLKVLVEHGADV |
| *Crocuta crocuta* | KQGASPNVQDTS-GTTPAHDAARTGFLDTLRVLVEHGADV |
| *Dasypus novemcinctus* | KQGASVNVQDAS-GTTPAHDAARTGFLDTLKVLVDHGADV |
| *Delphinapterus leucas* | KQGASPNVQDAS-GTTPAHDAARTGFLDTLKVLVEHGADV |
| *Desmodus rotundus* | KQGASPNVQDAS-GTTPAHDAARTGFLDTLKVLVEHGADV |
| *Dipodomys ordii* | KQGASPNVQDAS-GTSPVHDAARTGFLDTLKVLVEHGADV |
| *Dipodomys spectabilis* | KQGASPNVQDAS-GTSPVHDAARTGFLDTLKVLVEHGADV |
| *Echinops telfairi* | KQGACPNVQDRS-GTTPAHDAARTGFLDTLKVLVEHGADV |
| *Elephantulus edwardii* | KQGASPNVQDAS-GISPAHDAARTGFLDTLRVLVEHGADV |
| *Enhydra lutris kenyoni* | KQGASPNVQDAT-GTTPAHDAARTGFLDTLKVLVEHGADV |
| *Eptesicus fuscus* | KQGASPNVQDAS-GTTPAHDAARTGFLDTLKVLVEHGADV |
| *Equus asinus* | KQGANPNIQDAS-GTTPAHDAARTGFLDTLRVLVEHGADV |
| *Equus caballus* | KQGANPNIQDAS-GTTPAHDAARTGFLDTLRVLVEHGADV |
| *Equus przewalskii* | KQGANPNIQDAS-GTTPAHDAARTGFLDTLRVLVEHGADV |
| *Equus quagga* | KQGANPNIQDAS-GTTPAHDAARTGFLDTLRVLVEHGADV |
| *Erinaceus europaeus* | KQGASPNVQDAW-GTTPAHDAARTGFLDTLKVLVDHGADV |
| *Eumetopias jubatus* | KQGASPNVQDTT-GTTPAHDAARTGFLDTLKVLVEHGADV |
| *Felis catus* | KQGASPNVQDTS-GTTPAHDAARTGFLDTLKVLVEHGADV |
| *Fukomys damarensis* | KQGASPNIQDAS-GTSPVHDAARTGFLDTLKVLVEYGADV |
| *Galemys pyrenaicus* | KQGASPNVQDAS-GTTPAHDAARTGFLDTLKVLVEHGADV |
| *Galeopterus variegatus* | KQGASPNVQDAS-GTSPVHDAARTGFLDTLKALVEYGADV |
| *Globicephala melas* | KQGASPNVQDAS-GTTPAHDAARTGFLDTLKVLVEHGADV |
| *Gorilla gorilla gorilla* | KQGASPNVQDTS-GTSPVHDAARTGFLDTLKVLVEHGADV |
| *Grammomys surdaster* | KQGASPNVQDAS-GTSPVHDAARTGFLDTLKVLVEHGADV |
| *Gulo gulo luscus* | KQGASPNVQDAT-GTTPAHDAARTGFLDTLKVLVEHGADV |
| *Halichoerus grypus* | KQGASPNVQDTT-GTTPAHDAARTGFLDTLKVLVEHGADV |
| *Heterocephalus glaber* | KQGASPNIQDAS-GTSPVHDAARTGFLDTLKVLVEYGADV |
| *Hipposideros armiger* | KQGASPNVQDAS-GTTPAHDAARTGFLDTLKVLVEHGADI |
| *Homo sapiens* | KQGASPNVQDTS-GTSPVHDAARTGFLDTLKVLVEHGADV |
| *Hyaena hyaena* | KQGASPNVQDTS-GTTPAHDAARTGFLDTLRVLVEHGADV |
| *Hylobates moloch* | KQGASPNVQDTS-GTSPVHDAARTGFLDTLKVLVEHGADV |
| *Ictidomys tridecemlineatus* | KQGASPNVQDAS-GTSPVHDAARTGFLDTLKVLVEHGADV |
| *Jaculus jaculus* | KQGASPNVQDAS-GTSPVHDAARTGFLDTLKVLVEHGADI |
| *Lagenorhynchus obliquidens* | KQGASPNVQDAS-GTTPAHDAARTGFLDTLKVLVEHGADV |
| *Lemur catta* | KQGASPNVQDIS-GTSPVHDAARTGFLDTLKVLVEHGADV |
| *Leopardus geoffroyi* | KQGASPNVQDTS-GTTPAHDAARTGFLDTLKVLVEHGADV |
| *Leptonychotes weddellii* | KQGASPNVQDTT-GTTPAHDAARTGFLDTLKVLVQHGADV |
| *Lipotes vexillifer* | KQGASPNVQDAS-GTTPAHDAARTGFLDTLKVLVEHGADV |
| *Lontra canadensis* | KQGASPNVQDAT-GTTPAHDAARTGFLDTLKVLVEHGADV |
| *Loxodonta africana* | KQGASPNVQDVS-GTTPAHDAARTGFLDTLKVLVEHGADV |
| *Lynx canadensis* | KQGASPNVQDTS-GTTPAHDAARTGFLDTLKVLVEHGADV |
| *Lynx pardinus* | KQGASPNVQDTS-GTTPAHDAARTGFLDTLKVLVEHGADV |
| *Macaca fascicularis* | KQGASPNVQDTS-GTSPVHDAARTGFLDTLKVLVEHGADV |
| *Macaca mulatta* | KQGASPNVQDTS-GTSPVHDAARTGFLDTLKVLVEHGADV |
| *Macaca nemestrina* | KQGASPNVQDTS-GTSPVHDAARTGFLDTLKVLVEHGADV |
| *Mandrillus leucophaeus* | KQGASPNVQDTS-GTSPVHDAARTGFLDTLKVLVEHGADV |
| *Manis javanica* | KQGASPNVQDAS-GTTPAHDAARTGFLDTLKVLVEHGADV |
| *Manis pentadactyla* | KQGASPNVQDAS-GTTPAHDAARTGFLDTLKVLVEHGADV |
| *Marmota flaviventris* | KQGASPNVQDAS-GTSPVHDAARTGFLDTLKVLVEHGADV |
| *Marmota marmota marmota* | KQGASPNVQDAS-GTSPVHDAARTGFLDTLKVLVEHGADV |
| *Marmota monax* | KQGASPNVQDAS-GTSPVHDAARTGFLDTLKVLVEHGADV |
| *Mastomys coucha* | KQGASPNVQDAS-GTSPVHDAARTGFLDTLKVLVEHGADV |
| *Meles meles* | KQGASPNVQDAT-GTTPAHDAARTGFLDTLKVLVEHGADV |
| *Meriones unguiculatus* | KQGASPNVQDAS-GTSPVHDAARTGFLDTLKVLVEHGADV |
| *Mesocricetus auratus* | KQGASPNVQDAS-GTSPVHDAARTGFLDTLKVLVEHGADV |
| *Microcebus murinus* | KQGASPNVQDIS-GTSPVHDAARTGFLDTLKVLVEHGADV |
| *Microtus ochrogaster* | KQGASPNVQDAS-GTSPVHDAARTGFLDTLKVLVEHGADV |
| *Microtus oregoni* | KQGASPNVQDAS-GTSPVHDAARTGFLDTLKVLVEHGADV |
| *Miniopterus natalensis* | KQGASPNVQDAS-GITPAHDAARTGFLDTLKVLVEHGADV |
| *Mirounga angustirostris* | KQGASPNVQDTT-GTTPAHDAARTGFLDTLKVLVEHGADV |
| *Mirounga leonina* | KQGASPNVQDTT-GTTPAHDAARTGFLDTLKVLVEHGADV |
| *Molossus molossus* | KQGASPNVQDTS-GTTPAHDAARTGFLDTLKVLVEHGADV |
| *Monodon monoceros* | KQGASPNVQDAS-GTTPAHDAARTGFLDTLKVLVEHGADV |
| *Mus caroli* | KQGASPNVQDAS-GTSPVHDAARTGFLDTLKVLVEHGADV |
| *Mus musculus* | KQGASPNVQDAS-GTSPVHDAARTGFLDTLKVLVEHGADV |
| *Mus pahari* | KQGASPNVQDAS-GTSPVHDAARTGFLDTLKVLVEHGADV |
| *Mustela erminea* | KQGASPNVQDAT-GTTPAHDAARTGFLDTLKVLVEHGADV |
| *Mustela putorius furo* | KQGASPNVQDAT-GTTPAHDAARTGFLDTLKVLVEHGADV |
| *Myotis brandtii* | KQGASPNVQDTS-GTTPAHDAARTGFLDTLKVLVEHGADV |
| *Myotis davidii* | KQGASPNVQDTS-GTTPAHDAARTGFLDTLKVLVEHGADV |
| *Myotis lucifugus* | KQGASPNVQDTS-GTTPAHDAARTGFLDTLKVLVEHGADV |
| *Myotis myotis* | KQGASPNVQDTS-GTTPAHDAARTGFLDTLKVLVEHGADV |
| *Nannospalax galili* | KQGASPNVQDAS-GTSPVHDAARTGFLDTLKVFLDHGADV |
| *Neogale vison* | KQGASPNVQDAT-GTTPAHDAARTGFLDTLKVLVEHGADV |
| *Neomonachus schauinslandi* | KQGASPNVQDTT-GTTPAHDAARTGFLDTLKVLVEHGADV |
| *Neophocaena asiaeorientalis asiaeorientalis* | KQGASPNVQDAS-GTTPAHDAARTGFLDTLKVLVEHGADV |
| *Neosciurus carolinensis* | KQGASPNVQDAS-GTSPVHDAARTGFLDTLKVLVEHGADV |
| *Neotoma lepida* | KQGASPNVQDAS-GTSPVHDAARTGFLDTLKVLVEHGADV |
| *Nomascus leucogenys* | KQGASPNVQDTS-GTSPVHDAARTGFLDTLKVLVEHGADV |
| *Nyctereutes procyonoides* | KQGASPNVQDAT-GTTPAHDAARTGFLDTLKVLVEHGADV |
| *Ochotona curzoniae* | KQGASPNVQDAW-GTSPAHDAARTGFLDTLKVLVDHGADI |
| *Ochotona princeps* | KQGASPNVQDAW-GTSPAHDAARTGFLDTLKVLVDHGADI |
| *Octodon degus* | KQGASPNVQDAS-GTSPLHDAARTGFLDTLKVLVEHGADV |
| *Odobenus rosmarus divergens* | KQGASPNVQDTT-GTTPAHDAARTGFLDTLKVLVEHGADV |
| *Odocoileus virginianus texanus* | KQGASPNVQDAS-GTTPAHDAARTGFLDTLKVLVEHGADV |
| *Onychomys torridus* | KQGASPNVQDAS-GTSPVHDAARTGFLDTLKVLVEHGADV |
| *Orcinus orca* | KQGASPNVQDAS-GTTPAHDAARTGFLDTLKVLVEHGADV |
| *Orycteropus afer afer* | KQGASPNVQDNS-GTTPAHDAARTGFLDTLKILVEHGADV |
| *Oryctolagus cuniculus* | KQGASPNVQDAS-GTAPAHDAARTGFLDTLKVLVDHGADV |
| *Oryx dammah* | KQGASPNVQDAS-GTTPAHDAARTGFLDTLKVLVEHGADV |
| *Otolemur garnettii* | KQGASPNIQDMS-GTSPVHDAARTGFLDTLKVLVEHGADV |
| *Ovis aries* | KQGASPNVQDAS-GTTPAHDAARTGFLDTLKVLVEHGADV |
| *Pan paniscus* | KQGASPNVQDTS-GTSPVHDAARTGFLDTLKVLVEHGADV |
| *Pan troglodytes* | KQGASPNVQDTS-GTSPVHDAARTGFLDTLKVLVEHGADV |
| *Panthera leo* | KQGASPNVQDTS-GTTPAHDAARTGFLDTLKVLVEHGADV |
| *Panthera pardus* | KQGASPNVQDTS-GTTPAHDAARTGFLDTLKVLVEHGADV |
| *Panthera tigris* | KQGASPNVQDTS-GTTPAHDAARTGFLDTLKVLVEHGADV |
| *Papio anubis* | KQGASPNVQDTS-GTSPVHDAARTGFLDTLKVLVEHGADV |
| *Perognathus longimembris pacificus* | KQGASPNVQDAS-GTSPAHDAARTGFLDTLKVLVEHGADV |
| *Peromyscus leucopus* | KQGASPNVQDAS-GTSPVHDAARTGFLDTLKVLVEHGADV |
| *Peromyscus maniculatus bairdii* | KQGASPNVQDAS-GTSPVHDAARTGFLDTLKVLVEHGADV |
| *Phoca vitulina* | KQGASPNVQDTT-GTTPAHDAARTGFLDTLKVLVEHGADV |
| *Phocoena sinus* | KQGASPNVQDAS-GTTPAHDAARTGFLDTLKVLVEHGADV |
| *Phyllostomus discolor* | KQGASPNVQDAS-GTTPAHDAARTGFLDTLKVLVEHGADV |
| *Phyllostomus hastatus* | KQGASPNVQDAS-GTTPAHDAARTGFLDTLKVLVEHGADV |
| *Physeter catodon* | KQGASPNVQDAS-GTTPAHDAARTGFLDTLKVLVEHGADV |
| *Piliocolobus tephrosceles* | KQGASPNVQDTS-GTSPVHDAARTGFLDTLKVLVEHGADV |
| *Pipistrellus kuhlii* | KQGASPNVQDAA-GTTPAHDAARTGFLDTLKVLVEHGADV |
| *Pongo abelii* | KQGASPNVQDTS-GTSPVHDAARTGFLDTLKVLVEHGADV |
| *Prionailurus bengalensis* | KQGASPNVQDTS-GTTPAHDAARTGFLDTLKVLVEHGADV |
| *Procavia capensis* | KQGASPNVQDAS-GTTPAHDAARTGFLDTLKVLVEHGADV |
| *Propithecus coquereli* | KQGASPNVQDIS-GTSPVHDAARTGFLDTLKVLVEHGADV |
| *Pteropus alecto* | KQGASPNVQDAS-GTTPAHDAARTGFLDTLKVLVEHGADV |
| *Pteropus giganteus* | KQGASPNVQDAS-GTTPAHDAARTGFLDTLKVLVEHGADV |
| *Pteropus vampyrus* | KQGASPNVQDAS-GTTPAHDAARTGFLDTLKVLVEHGADV |
| *Puma concolor* | KQGASPNVQDTS-GTTPAHDAARTGFLDTLKVLVEHGADV |
| *Puma yagouaroundi* | KQGASPNVQDTS-GTTPAHDAARTGFLDTLKVLVEHGADV |
| *Rattus norvegicus* | KQGASPNVQDAS-GTSPVHDAARTGFLDTLKVLVEHGADV |
| *Rattus rattus* | KQGASPNVQDAS-GTSPVHDAARTGFLDTLKVLVEHGADV |
| *Rhinolophus ferrumequinum* | KQGASPNVQDAS-GTTPAHDAARTGFLDTLKVLVEHGADV |
| *Rhinolophus sinicus* | KQGASPNVQDAS-GTTPAHDAARTGFLDTLKVLVEHGADV |
| *Rhinopithecus bieti* | KQGASPNVQDTS-GTSPVHDAARTGFLDTLKVLVEHGADV |
| *Rhinopithecus roxellana* | KQGASPNVQDTS-GTSPVHDAARTGFLDTLKVLVEHGADV |
| *Rousettus aegyptiacus* | KQGASPNVQDAS-GTTPAHDAARTGFLDTLKVLVEHGADV |
| *Saimiri boliviensis boliviensis* | KQGASPNVQDTS-GTSPVHDAARTGFLDTLKVLVEHGADV |
| *Sapajus apella* | KQGASPNVQDTT-GTSPVHDAARTGFLDTLKVLVEHGADV |
| *Sorex araneus* | KQGACPNVQDTS-GATPAHDAARTGFLDTLRLLVDHGADV |
| *Sousa chinensis* | KQGASPNVQDAS-GTTPAHDAARTGFLDTLKVLVEHGADV |
| *Suricata suricatta* | KQGASPNVQDTS-GTTPAHDAARTGFLDTLKVLVEHGADV |
| *Sus scrofa* | KQGASPNVQDAS-GTTPAHDAARTGFLDTLKVLVEHGADV |
| *Talpa occidentalis* | KQGASPNVQDAS-GTTPAHDAARTGFLDTLKVLVEHGADV |
| *Theropithecus gelada* | KQGASPNVQDTS-GTSPVHDAARTGFLDTLKVLVEHGADV |
| *Trachypithecus francoisi* | KQGASPNVQDTS-GTSPVHDAARTGFLDTLKVLVEHGADV |
| *Trichechus manatus latirostris* | KQGASPNVQDIS-GTTPAHDAARTGFLDTLKVLVEHGADV |
| *Tupaia chinensis* | KQGASPNVQDAS-GTSPIHDAARTGFLDTLKVLVDHGADV |
| *Tursiops truncatus* | KQGASPNVQDAS-GTTPAHDAARTGFLDTLKVLVEHGADV |
| *Urocitellus parryii* | KQGASPNVQDAS-GTSPVHDAARTGFLDTLKVLVEHGADV |
| *Ursus americanus* | KQGASPNVQDTT-GTTPAHDAARTGFLDTLKVLVEHGADV |
| *Ursus arctos* | KQGASPNVQDTT-GTTPAHDAARTGFLDTLKVLVEHGADV |
| *Ursus maritimus* | KQGASPNVQDTT-GTTPTHDAARTGFLDTLKVLVEHGADV |
| *Vicugna pacos* | KQGASPNVQDAS-GTTPAHDAARTGFLDTLKVLVEHGADV |
| *Vulpes lagopus* | KQGASPNVQDAT-GTTPAHDAARTGFLDTLKVLVEHGADV |
| *Vulpes vulpes* | KQGASPNVQDAT-GTTPAHDAARTGFLDTLKVLVEHGADV |
| *Zalophus californianus* | KQGASPNVQDTT-GTTPAHDAARTGFLDTLKVLVEHGADV |

**<Marsupials>**

| | |
|---|---|
| *Dromiciops gliroides* | KQGASPNVQDGS-GTSPAHDAARTGFLDTLRILVEHGADV |
| *Gracilinanus agilis* | KQGASPNVQDGS-GTSPAHDAARTGFLDTLRILVEHGADV |
| *Monodelphis domestica* | KQGASPNVQDGS-GTSPAHDAARTGFLDTLRVLVEHGADV |
| *Notamacropus eugenii* | KQGASPNVQDGS-GTSPAHDAARTGFLDTLRVLVEHGADV |
| *Phascolarctos cinereus* | KQGASPNVQDGS-GTSPAHDAARTGFLDTLRVLVEHGADV |
| *Sarcophilus harrisii* | KQGASPNVQDGS-GTSPAHDAARTGFLDTLRILVEHGADV |
| *Trichosurus vulpecula* | KQGASPNVQDGS-GTSPAHDAARTGFLDTLRVLVEHGADV |
| *Vombatus ursinus* | KQGASPNVQDGS-GTSPAHDAARTGFLDTLRVLVEHGADV |

**<Monotremes>**

| | |
|---|---|
| *Ornithorhynchus anatinus* | KQGANPNVQDAT-GTTPVHDAARTGFLDTLQALVEHGADV |
| *Tachyglossus aculeatus* | KQGASPNVQDAS-GITPAHDAARTGFLDTLQALVEHGADV |

**Figure S2i**

**Cdkn2a**

| | |
|---|---|
| *Homo sapiens* | LHGAEPNCADPATLTRPVHDAAREGFLDTLVVLHRAGARL |

**Cdkn2d**

**Reptiles**

| | |
|---|---|
| *Alligator mississippiensis* | KQGASANVQD-AGGRAPAHDAARGGFADTLRVLLEHGADA |
| *Anolis carolinensis* | KHGANPNLQD-ASGTAPAHDAARTGFLDTLKVLVEYGADV |
| *Chelonia mydas* | KQGASPNIQD-EAGRAPAHDAARTGFLDTLRVLVEHGADV |
| *Chelonoidis abingdonii* | KQGASPNIQD-EAGRAPAHDAARTGFLDTLRVLVEHGADV |
| *Chelydra serpentina* | KQGASPNIQD-EAGRAPAHDAARTGFLDTLRVLVEHGADV |
| *Chrysemys picta bellii* | KQGASPNIQD-EAGRAPVHDAARTGFLDTLRVLVEHGADV |
| *Crocodylus porosus* | KQGASANVQD-AGGRAPAHDAARGGFADTLRVLLEHGADA |
| *Crotalus adamanteus* | KQGANPNIQD-GSGTTPAHDAARTGFLDTLKILVEHGADV |
| *Crotalus tigris* | KQGANPNIQD-GSGTTPAHDAARTGFLDTLKILVEHGADV |
| *Dermochelys coriacea* | KQGASPNIQD-EAGRAPAHDAARTGFLDTLRVLVEHGADV |
| *Gavialis gangeticus* | KQGASANVQD-AGGRAPAHDAARGGFADTLRVLLEHGGDA |
| *Gekko japonicus* 1 | KQGVSPNVQD-ESGTAPAHNAARTGFLDTLSILVEHGADV |
| *Gekko japonicus* 2 | --------QD-ESGTAPAHDAARTGFLDTLSVLVEHGADV |
| *Gopherus agassizii* | KQGASPNIQD-EAGRAPAHDAARTGFLDTLRVLVEHGADI |
| *Gopherus evgoodei* | KQGASPNIQD-EAGRAPAHDAARTGFLDTLRVLVEHGADI |
| *Lacerta agilis* | KQGANPNVRD-ESGTAPAHDAARTGFLDTLKILVEYGADV |
| *Laticauda laticaudata* | KQGANPNIQD-GSGTTPAHDAARTGFLDTLKILVEHGADV |
| *Mauremys mutica* | KQGASPNIQD-EAGRAPAHDAARTGFLDTLRVLVEHGADV |
| *Mauremys reevesii* | KQGASPNIQD-EAGRAPAHDAARTGFLDTLRVLVEHGADV |
| *Naja naja* | KQGANPNIQD-ASGTTPAHDAARTGFLDTLKILVEHGADV |
| *Notechis scutatus* | KQGANPNIQD-GSGTTPAHDAARTGFLDTLKILVEHGADV |
| *Ophiophagus hannah* | KQGANPNIQD-GSGTTPAHDAARTGFLDTLKILVEHGADV |
| *Pantherophis guttatus* | KQGANPNIQD-GSGTTPAHDAARTGFLDTLKILVEHGADV |
| *Pelodiscus sinensis* | KQGASPNIQD-EVGRAPAHDAARTGFLDTLRVLVEYGADV |
| *Pelusios castaneus* | KQGASPNIQD-EAGRAPAHDAARTGFLDTLCVLVDHGADV |
| *Podarcis muralis* | KQGANPNVRD-ESGTAPAHDAARTGFLDTLKILVEYGADV |
| *Pogona vitticeps* | KEGAHPNVQD-ESGTTPAHDAARTGFLDTLKILVEYGADV |
| *Protobothrops mucrosquamatus* | KQGANPNIQD-GSGTTPAHDAARTGFLDTLKILVEHGADV |
| *Pseudonaja textilis* | KQGANPNIQD-GSGTTPAHDAARTGFLDTLKILVEHGADV |
| *Python bivittatus* | KQGANPNIQD-GSGTTPAHDAARTGFLDTLKILVEHGADV |
| *Salvator merianae* | KQGANPNIQD-KSGTAPAHDAARTGFLDTLKVLVEHGADV |
| *Sceloporus undulatus* | KQGANPNVQD-ESSTAPAHDAARTGFIDTLKILVEHGADV |
| *Sphenodon punctatus* | KQGASPNIQD-ETGTAPVHDAARTGFLDTLQLLVEHGADI |
| *Thamnophis elegans* | KQGANPNVRD-GSGTTPAHDAARTGFLDTLKVLVEHGADV |
| *Thamnophis sirtalis* | KQGANPNVRD-GSGTTPAHDAARTGFLDTLKVLVEHGADV |
| *Varanus komodoensis* | KQGANPNIQD-ESGTVPAHDAARTGFLDTLKILVDHGADV |
| *Zootoca vivipara* | KQGANPNVRD-ESGTAPAHDAARTGFLDTLKILVEYGADV |

**Amphibians**

| | |
|---|---|
| *Ambystoma andersoni* | KQGASVTIQD-EYGTTPAHDAARTGFLQTIQLLVEHGADI |
| *Ambystoma maculatum* | KQGASVTIQD-EYGTTPAHDAARTGFLQTIQLLVEHGADI |
| *Ambystoma mexicanum* | KQGASVNIQD-EYGTTPAHDAARTGFLQTIQLLVEHGADI |
| *Ambystoma velasci* | KQGASVTIQD-EYGTTPAHDAARTGFLQTIQLLVEHGADI |
| *Bufo bufo* | KQGASPNIQD-SYGISPAHDAARMGFLDTLQVLVQYGADV |
| *Bufo gargarizans* | KQGASPNIQD-SYGISPAHDAARMGFLDTLQVLVQYGADV |
| *Engystomops pustulosus* | KQGASPNIQD-SYGITPAHDAARMGFLDTLQVLVQYGADV |
| *Geotrypetes seraphini* | KQGACANVQD-SYGTTPAHDAARTGFLETLQLLVEHGADI |
| *Hymenochirus boettgeri* | KQGASPNIQD-AYGTSPAHDAARSGFLDTLQVLVQHGAEV |
| *Leptobrachium leishanense* | KQGSSPNIQD-AYGITPAHDAARCGFLDTLQVLVQYGADL |
| *Microcaecilia unicolor* | KQGACANVQD-AYGTTPAHDAARTGFLETLQLLVEHGADV |
| *Nanorana parkeri* | KQGATANIQD-PYGITPAHDAARTGFLDTLQILVEYGADL |
| *Rana temporaria* | KQGATANIQD-SHGITPAHDAARTGFLDTLQILVQYGADL |
| *Rhinatrema bivittatum* | KQGACANVQDPGCGTTPAHDAARTGFPGHPAAAGGARADV |
| *Xenopus laevis* 1 | KQGATPNIQD-AYGTTPAHDAARCGFLDTLQVLVQHGAEV |
| *Xenopus laevis* 2 | KQGASPNIQD-SFGTTPAHDAARCGFLDTLQVLLQHGAEV |
| *Xenopus tropicalis* | KQGASPNIQD-AYGTTPAHDAARCGFLDTLQVLVQHGAEI |

**Bony fish**

| | |
|---|---|
| *Acanthochromis polyacanthus* | EKGAEPNIQD-KHGIAPIHDAARTGFLDTLQVLVEYGASV |
| *Acanthopagrus latus* | EKGADPNVQD-KHGIAPVHDAARTGFLDTLQVLVEYGASV |
| *Acipenser ruthenus* 1 | KHGASANVQD-GLGISPAHDAARTGFVDTLRVLVEYGASV |
| *Acipenser ruthenus* 2 | KHGASANVQD-GLGISPAHDAARTGFVDTLRVLVEYGASV |
| *Albula glossodonta* | EHGANANVQD-RQGVAPAHDAARTGFVDTLRVLVQFGASV |
| *Alosa alosa* | EHGADPNIQD-RFGVTPAHDAARGGFLDTLRALVDFGASV |
| *Alosa sapidissima* | EHGADPNIQD-RFGVTPAHDAARGGFLDTLRALVDFGASV |
| *Ameiurus melas* | ENGADPNIQD-RFGITPAHDAARTGFLDTLCVLVDYGASV |
| *Amphilophus citrinellus* | EKGAEPNIQD-RHGITPVHDAARTGFLDTVQVLVEYGASV |
| *Amphiprion ocellaris* | EKGAEPNVQD-KHGIAPIHDAARTGFLDTLQVLVEYGASV |
| *Amphiprion percula* | EKGAEPNVQD-KHGIAPIHDAARTGFLDTLQVLVEYGASV |
| *Anabarilius grahami* | DHGADPNVQD-RCGVTPAHDAARTGFLDTVRVLVDYGASV |
| *Anabas testudineus* | EKGADPNIQD-KQGITPVHDAARTGFLDTVQVLVEYGASV |
| *Anarrhichthys ocellatus* | EKGADPNVQD-KHGIAPVHDAARTGFLDTLQVLVEYGASV |
| *Anguilla anguilla* | EHGANANVQD-RLGVTPAHDAARAGFVDTLRVLVQFGASV |
| *Archocentrus centrarchus* | EKGAEPNIQD-RHGITPVHDAARTGFLDTVQVLVEYGASV |
| *Astatotilapia calliptera* | EKGANPNVQD-RHGITPVHDAARTGFLDTVEVLVEYGASV |
| *Astyanax mexicanus* | EKGADPNIPD-RFGITPAHDAARTGFLETLHVLVDYGASV |
| *Austrofundulus limnaeus* | EKGANPNVQN-KHGIAPAHDAARTGFLDTLQVLVEHGASV |
| *Bagarius yarrelli* | ENGADPNIQD-RFGITPAHDAARTGFLDTLCVLVDYGASV |
| *Betta splendens* | EKGADPNVQD-KQGITPLHDAARTGFLDTVQVLVEYGALV |
| *Boleophthalmus pectinirostris* | QKGANPNLQD-KHGISPAHDAARTGFLDTLRVLVEHGASV |
| *Caranx melampygus* | EKGANPNVQD-KYGIAPVHDAARTGFLDTLRVLVEYGASV |
| *Carassius auratus* 1 | DHGADPNVQD-RCGVTPAHDAARTGFLDTLRVLVEHGASV |
| *Carassius auratus* 2 | DHGADPNVQD-RCGVTPAHDAARTGFLDTLRVLVEHGASV |
| *Channa argus* | EKGADPNIQD-KHGIAPVHDAARTGFLDTLKVLVEFGASV |
| *Chanos chanos* | ENGADPNIQD-CFGVTPAHDAARTGFLDTLRALVLYGASV |
| *Cheilinus undulatus* | EKGADPNIQD-KHGIAPVHDAARTGFLDTLQVLVEYGASV |
| *Chelmon rostratus* | EKGADPNRQD-KYGIAPVHDAARTGFLDTLQVLVEYGASV |
| *Clarias magur* | ENGADPNIQD-HFGITPAHDAARTGFLDTLCVLVDYGASV |
| *Clupea harengus* | EHGADPNIQD-RFGVTPAHDASRGGFLDTLRALVDFGASV |
| *Collichthys lucidus* | EKGADPNLQD-KHGIAPVHDAARTGFLDTLQVLVEYGASV |
| *Colossoma macropomum* | ESGANPNIQD-RFGITPAHDAARTGFLDTLRVLVDFGADV |
| *Coregonus clupeaformis* | EHGADPNITD-RRGVSPAHDAARTGFVDTLRVLVEFGASV |
| *Coryphaenoides rupestris* | EHGADPNAQD-QHGIAPAHDAARTGFLDTLRALVEHGASV |
| *Cottoperca gobio* | EKGADPNIQD-KHGIAPVHDAARTGFVDTLQVLVEYGASV |
| *Cyclopterus lumpus* | EKGADPNVQD-KHGIAPIHDAARTGFPDTLQVLVEYGASV |
| *Cynoglossus semilaevis* | EKGAQPNIQD-RHGIAPVHDAARTGFLDTVGVLVEYGASV |
| *Cyprinodon tularosa* | EKGAEPNIQD-KHGIAPVHDAAQTGFLETLEVLVEHGASV |
| *Cyprinodon variegatus* | EKGAEPNIQD-KHGIAPVHDAAQTGFLETLEVLVEHGASV |
| *Cyprinus carpio* 1 | DHGADPNVQD-RCGVTPAHDAAPHRIPDTLRVLVDHGASV |
| *Cyprinus carpio* 2 | DHGADPNVQD-RCGVTPAHDAARTGFLDTLRVLVDHGASV |
| *Danio rerio* | DFGADPNVQD-RSGVTPAHDAARTGFLETLRVLVDGGASV |
| *Danionella translucida* | DHGANPNLQD-RCGVAPIHDAARTGFLDTVRVLVEGGASV |
| *Denticeps clupeoides* | EHGANPNAQD-RFGVTPVHDAARTGFLNTLRALVDHGASV |
| *Dicentrarchus labrax* | EKGADPNMQD-KHGIAPVHDAARTGFLDTVQVLVEYGASV |
| *Echeneis naucrates* | EKGADPNVQD-KYGIAPVHDAARTGFLDTLRVLVEYGASV |
| *Electrophorus electricus* | ENGADPNIQD-NFGITPAHDAARTGFLDTLHVLVDYGASV |
| *Epinephelus coioides* | EKGADPNVQD-KHGIAPVHDAARTGFLDTLQVLVEYGASV |
| *Epinephelus lanceolatus* | EKGADPNVQD-KHGIAPVHDAARTGFLDTLQVLVEYGASV |
| *Erpetoichthys calabaricus* | THGATPNVQD-SLGITPAHDAARTGFLETLKVLVEHGASV |
| *Esox lucius* | EHGANPNITD-KRGISPAHDAAHTGFLDTLQVLVEFGASV |
| *Etheostoma cragini* | EKGADPNVQD-KHGIAPVHDAARTGFLDTLQVLVEYGASV |
| *Etheostoma spectabile* | EKGADPNVQD-KHGIAPVHDAARTGFLDTLQVLVEYGASV |
| *Fundulus heteroclitus* | EKGAEPNVQD-KYGIAPVHDAARTGFLETLQVLVEHGASV |
| *Gadus morhua* | EHGANPNAQD-RHGITPAHDAAQTGFLETLQVLVEHGASV |
| *Gambusia affinis* | EKGAEPNIQD-KHGIAPVHDAARTGFLETLQVLVEHGASV |
| *Gasterosteus aculeatus aculeatus* | EKGADPNVQD-NHGIAPVHDAARTGFPDTLRVLVEYGASV |
| *Girardinichthys multiradiatus* | EKGAEPNIQD-KHGIAPVHDAAQTGFLETLQVLVEYGASV |
| *Gouania willdenowi* | EKGADPNVQD-KHGVAPVHDAAQTGFLDTLQVLVEYGASV |
| *Gymnodraco acuticeps* | EKGANPNVQD-KHGIAPVHDAARTGFLDTLQVLVEYGASV |
| *Haplochromis burtoni* | EKGANPNVQD-RHGITPVHDAARTGFLDTVEVLVEYGASV |
| *Hippocampus comes* | ERGADPNVQD-AGGTAPVHDAARTGFVDTLKVLVEHGASV |
| *Hippoglossus hippoglossus* | EKGAEPNVQD-RHGIAPVHDAARTGFLDTVLALVAHGASV |
| *Hippoglossus stenolepis* | EKGAEPNVQD-RHGIAPVHDAARTGFLDTVLALVAHGASV |
| *Hucho hucho* 1 | EHGADPNITD-RRGVSPAHDAARTGFVDTLRVLVEFGASV |
| *Hucho hucho* 2 | EHGADPNITD-RRGVSPAHDAARTGFVDTLRVLVEFGASV |
| *Hypomesus transpacificus* | EHGADPNIQD-RQGVTPAHDAARTGFLDTLRVLVEFGASV |
| *Iconisemion striatum* | EKGADPNLQD-RHGIAPVHDAARTGFLNTLQVLVEHGASV |
| *Ictalurus punctatus* | ENGADPNIQD-RFGITPVHDAARTGFLDTLCVLVDYGASV |
| *Kryptolebias marmoratus* | EKGADPNVQD-KHGIAPAHDAARTGFLDTLQVLVEHGASV |
| *Labeo rohita* | EHGADPNVQD-RCGVTPAHDAARTGFLDTLRVLVDYGASV |
| *Labrus bergylta* | EKGADPNIQD-KHGIAPVHDAARTGFLDTLRVLVDYGASV |
| *Larimichthys crocea* | EKGADPNLQD-KHGIAPVHDAARTGFLDTLQVLVEYGASV |
| *Lates calcarifer* | EKGADPNVQD-KHGIAPVHDAARTGFVDTLQVLVEYGASV |
| *Latimeria chalumnae* | KCGADPNVQD-AHGNTPAHDAVRTGFLETLQVLVENGANV |
| *Lepisosteus oculatus* 1 | ERGAEPNVRD-RLGITPAHDAARTGFLDTLRVLVQFGAAV |
| *Lepisosteus oculatus* 2 | ERGADPNVQD-HSGVTPAHDAARTGFLETLRILVEFGAGV |
| *Liparis tanakae* | EKGADPNVQD-KHGIAPVHDAARTGFPDTLQVLVEYGASV |
| *Lucifuga dentata* | EKGANPNVQD-HHGVAPVHDAARTGFLDTVEVLVEYGASV |
| *Mastacembelus armatus* | EKGANPNVQD-RHGIAPIHDAARTGFLDTVQVLVEYGASV |
| *Maylandia zebra* | EKGANPNVQD-RHGITPVHDAARTGFLDTVEVLVEYGASV |
| *Megalobrama amblycephala* | DHGADPNVQD-RCGVTPAHDAARTGFLDTVRVLVDYGASV |
| *Megalops atlanticus* | EHGANANVQD-RHGVTPAHDAARTGFVDTLRVLVEFGASV |
| *Megalops cyprinoides* | EHGANANVQD-RHGVTPAHDAARTGFVDTLRVLVEFGASV |
| *Melanotaenia boesemani* | EKGADPNIQD-KHGITPVHDAARTGFLDTLEVLVEHGASV |
| *Micropterus dolomieu* | EKGADPNVQD-KHGIAPIHDAARTGFLDTVQVLVEYGALV |
| *Micropterus salmoides* | EKGADPNVQD-KHGIAPIHDAARTGFLDTVQVLVEYGALV |
| *Mola mola* | EKGADPNMQD-KHGIVPVHDAARTGFLDTLQVLVEYGASV |
| *Monopterus albus* | EKGADPNVQD-RHGIAPVHDAARTGFLDTLQVLVEYGASV |
| *Morone saxatilis* | EKGADPNMQD-KHGIAPVHDAARTGFLDTVQVLVEYGASV |
| *Mugil cephalus* | EKGADPSVQD-KQGIAPVHDAARTGFLDTLQVLVEYGASV |
| *Myripristis murdjan* | EYGADPNIQD-KHGIAPVHDAARTGFLDTLRVLVEYGASV |
| *Nematolebias whitei* | EKGADPNVQD-KHGIAPAHDAARTGFLDTLQVLVEHGASV |
| *Neogobius melanostomus* | QKGANPNLQD-KHGISPAHDAARTGFLDTLQVLVEHGASV |
| *Neolamprologus brichardi* | EKGANPNVQD-RHGITPVHDAARTGFLDTVEVLVEYGASV |
| *Nibea albiflora* | EKGADPNLQD-KHGIAPVHDAARTGFLDTLQVLVEYGASV |
| *Nothobranchius furzeri* | EKGADPNLQD-RHGIAPIHDAARTGFLDTLQVLVEHGASV |
| *Nothobranchius kadleci* | EKGADPNLQD-RHGIAPIHDAARTGFLDTLQVLVEHGASV |
| *Nothobranchius korthausae* | EKGADPNLQD-RHGIAPIHDAARTGFLDTLQVLVEHGASV |
| *Nothobranchius kuhntae* | EKGADPNLQD-RHGIAPIHDAARTGFLDTLQVLVEHGASV |
| *Nothobranchius pienaari* | EKGADPNLQD-RQGIAPIHDAARTGFLDTLQVLVEHGASV |
| *Nothobranchius rachovii* | EKGADPNLQD-RQGIAPIHDAARTGFLDTLQVLVEHGASV |
| *Notolabrus celidotus* | EKGADPNIQD-KHGIAPVHDAARTGFLDTLQVLVEYGASV |
| *Notothenia coriiceps* | EKGANPNVQD-KHGIAPVHDAARTGFLDTLQVLVEYGASV |
| *Oncorhynchus gorbuscha* | EHGADPNITD-RRGVSPAHDAARTGFVDTLRVLVEFGASV |
| *Oncorhynchus keta* 1 | EHGADPNITD-RRGVSPAHDVARTGFVDTLRVLVEFGASV |
| *Oncorhynchus keta* 2 | EHGADPNITD-RRGVSPAHDVARTGFVDTLRVLVEFGASV |
| *Oncorhynchus kisutch* 1 | EHGADPNITD-RRGVSPAHDAARTGFVDTLRVLVEFGGSV |
| *Oncorhynchus kisutch* 2 | EHGADPNITD-RRGVSPAHDAARTGFVDTLRVLVEFGGSV |
| *Oncorhynchus mykiss* 1 | EHGADPNITD-RRGVSPAHDAARTGFVDTLRVLVEFGASV |
| *Oncorhynchus mykiss* 2 | EYGADPNITD-RRGVSPAHDAARTGFVDTLRVLVEFGASV |
| *Oncorhynchus nerka* | EHGADPNITD-RRGVSPAHDAARTGFVDTLRVLVEFGASV |
| *Oncorhynchus tshawytscha* | EHGADPNITD-RRGVSPAHDAARTGFVDTLRVLVEFGASV |
| *Onychostoma macrolepis* | DHGADPNVQD-RCGVTPAHDAARTGFLDTLRVLVDHGASV |
| *Oreochromis aureus* | EKGANPNVQD-RHGITPVHDAARTGFLDTVEVLVEYGASV |
| *Oreochromis niloticus* | EKGANPNVQD-RHGITPVHDAVRTGFLDTVEVLVEYGASV |
| *Oryzias javanicus* | EKGADPNVQD-KHGIAPVHDAARTGFLDTLQVLVENGASV |
| *Oryzias latipes* | EKGADANVQD-KHGIAPVHDAARTGFLDTLQVLVENGASV |
| *Oryzias melastigma* | EKGADPNVQD-KHGIAPVHDAARTGFLDTLQVLVENGASV |
| *Oryzias sinensis* | EKGADANVQD-KHGIAPVHDAARTGFLDTLQVLVENGASV |
| *Pangasianodon hypophthalmus* | ENGADPNIQD-RFGITPAHDAARTGFLDTLCVLVDYGASV |
| *Paralichthys olivaceus* | EKGAEPNVQD-RHGIAPVHDAARTGFLDTVLVLVAHGASV |
| *Parambassis ranga* | EKGANPNIQD-RHGITPARDAACTGFLDTLQILVEYGASV |
| *Paramormyrops kingsleyae* 1 | EHGANANVQD-RHGVTPAHDAARSGFVDTLRVLVEFGASV |
| *Paramormyrops kingsleyae* 2 | EHGANANVRD-QQGITPAHDAARTGFTDTLSVLVQFGASV |
| *Perca flavescens* | EKGADPNVQD-KHGIAPVHDAARTGFLDTLQVLVEYGASV |
| *Perca fluviatilis* | EKGADPNVQD-KHGIAPVHDAARTGFLDTLQVLVEYGASV |
| *Periophthalmus magnuspinnatus* | QKGANPNVQD-KYGISPAHDAARTGFVDTLQVLVEHGASV |
| *Pimephales promelas* | DHGADPNVQD-RCGVTPAHDAARTGFLDTVRVLVDYGASV |
| *Plectropomus leopardus* | EKGADPNIQD-KHGIAPVHDAARTGFLDTLQVLVEYGASV |
| *Poecilia formosa* | EKGAEPNVQD-KHGITPVHDAAQTGFLETLQVLVEHGASV |
| *Poecilia latipinna* | EKGAEPNVQD-KHGIAPVHDAAQTGFLETLQVLVEHGASV |
| *Poecilia mexicana* | EKGAEPNVQD-KHGIAPVHDAAQTGFVETLQVLVEHGASV |
| *Poecilia reticulata* | EKGAEPNVQD-KHGIAPVHDAAQTGFLETLQVLVEHGASV |
| *Poeciliopsis prolifica* | EKGAEPNVQD-KHGIAPVHDAAQTGFLETLQVLVEHGASV |
| *Polyodon spathula* | KHGASVNVQD-GLGVTPAHDAARTGFVDTLRVLVEYGASV |
| *Polypterus senegalus* | THGATPNVQD-SLGITPAHDAARTGFLDTLKVLVEHGASV |
| *Protopterus annectens* | KNGANPNVQD-SYGITPSHDATRTGFLDTLKVLVEHGADV |
| *Pseudochaenichthys georgianus* | EKGADPNVQD-KHGIAPVHDAARTGFLDTLQVLVEYGASV |
| *Pundamilia nyererei* | EKGANPNVQD-RHGITPVHDAARTGFLDTVEVLVEYGASV |
| *Pungitius pungitius* | EKGADPNVQD-KHGIAPVHDAARTGFPDTVRVLVEYGASV |
| *Puntigrus tetrazona* | EHGADPNVQD-RCGVTPAHDAARTGFLDTLRVLVEHGASV |
| *Pygocentrus nattereri* | ESGANPNIQD-RFGITPAHDAARTGFLDTLRVLVDFGADV |
| *Salarias fasciatus* | EKGADPNVQD-KHGIAPVHDAARTGFLDTLQVLVEYGASV |
| *Salmo salar* 1 | EHGGDPNITD-RRGVSPAHDAARTGFVDTLRVLVEFGASV |
| *Salmo salar* 2 | EHGADPNITD-RRGVSPAHDAARTGFVDTLRVLVEFGASV |
| *Salmo trutta* 1 | EHGADPNITD-RRGVSPAHDAARTGFVDTLRVLVEFGASV |
| *Salmo trutta* 2 | EHGGDPNITD-RRGVSPAHDAARTGFVDTLRVLVEFGASV |
| *Salvelinus alpinus* 1 | EHGADPNITD-RRGVSPAHDAARTGFVDTLRALVEFGASV |
| *Salvelinus alpinus* 2 | EHGADPNITD-RRGVSPAHDAARTGFVDTLRALVEFGASV |
| *Salvelinus namaycush* | EHGADPNITD-RRGVSPAHDAARTGFVDTLRALVEFGASV |
| *Sander lucioperca* | EKGADPNVQD-KHGIAPVHDAARTGFLDTLQVLVEYGASV |
| *Scatophagus argus* | EKGADPNVQD-KHGIAPVHDVARTGFLDTLQVLVEYGASV |
| *Scleropages formosus* 1 | QHGANANVQD-GQGVTPVHDAARTGFVHTLRALVDFGASV |
| *Scleropages formosus* 2 | EHGANANVQD-QQGITPAHDAARTGFVDTLSVLHEFGASL |
| *Scophthalmus maximus* | EKGAEPNIQD-KHGIAPVHDAARTGFLDTLQVLVEYGASV |
| *Sebastes umbrosus* | EKGADPNVQD-KHGIAPVHDAARTGFPDTLQVLVEYGASV |
| *Seriola dumerili* | EKGADPNVQD-KHGIAPVHDAARTGFLDTLRVLVEYGASV |
| *Seriola lalandi dorsalis* | EKGADPNVQD-KHGIAPVHDAARTGFLDTLQVLVEYGASV |
| *Silurus meridionalis* | ESGADPNIQD-RFGIAPAHDAARTGFLDTLHVLVDYGASV |
| *Simochromis diagramma* | EKGANPNVQD-RHGITPVHDAARTGFLDTVEVLVEYGASV |
| *Siniperca chuatsi* | EKGADPNMQD-KHGIAPVHDAARTGFLDTLQVLVEYGASV |
| *Sinocyclocheilus anshuiensis* 1 | DHGADPSIQD-RCGVTPAHDEARTGFLDTLRVLVDHGASV |
| *Sinocyclocheilus anshuiensis* 2 | DHGADPNVQD-RCGVTPAHDAARTGFLDTLRVLVDHGASV |
| *Sinocyclocheilus grahami* 1 | DHGADPSVQD-RCGVTPAHDAARTGFLDTLSVLVDHGASV |
| *Sinocyclocheilus grahami* 2 | DHGADPSVQD-RCGVTPAHDAARTGFLDTLSVLVDHGASV |
| *Sinocyclocheilus rhinocerous* 1 | DHGADPNIQD-RCGVTPAHDAARTGFLDTLRVLVDHGASV |
| *Sinocyclocheilus rhinocerous* 2 | DHGADPNVQD-RCGVTPAHDAARTGFLDTLRVLVDHGASV |
| *Sparus aurata* | EKGADPNVQD-KHGIAPVHDAARTGFLDTLQVLVEYGASV |
| *Sphaeramia orbicularis* | EKGADPNVQD-KHGIAPVHDAARTGFLDTVQVLVEYGAAV |
| *Stegastes partitus* | EKGADPNVQD-KHGIAPIHDAARTGFLDTLQVLVEYGALV |
| *Syngnathus acus* | ERGADPNVQD-AGGTAPVHDAARTGFVDTLQVLVEHGASV |
| *Tachysurus fulvidraco* | ENGADPNIQD-RFGITPVHDAARTGFLDTLCVLVDYGASV |
| *Takifugu flavidus* | EKGADPNVQD-KHGIAPVHDAARTGFLDTLQVLVEYGASV |
| *Takifugu rubripes* | EKGADPNVQD-KHGIAPVHDAARTGFLDTLQVLVEYGASV |
| *Tetraodon nigroviridis* | EKGADPNVQD-KHGIAPVHDAARTGFLDTLQVLVEYGASV |
| *Thalassophryne amazonica* | QNGADANIQD-KHGIAPVHDAARTGFLDTLQVLVEFGASV |
| *Thunnus albacares* | EKGADPNIQD-KHGIAPVHDAARTGFLDTLQVLVEYGASV |
| *Thunnus maccoyii* | EKGADPNIQD-KHGIAPVHDAARTGFLDTLQVLVEYGASV |
| *Toxotes jaculatrix* | EKGADPNVQD-KHGIAPVHDAARTGFLDTLQVLVEYGASV |
| *Trematomus bernacchii* | EKGADPNVQD-KHGIAPVHDAARTGFLDTLQVLVEYGASV |
| *Triplophysa tibetana* | EHGADPNVQD-RCGVTPAHDAARTGFLETLRALVEHSASV |
| *Xiphias gladius* | EKGADPNIQD-KHGIAPVHDAARTGFVDTLQVLVEYGASV |
| *Xiphophorus couchianus* | EKGAEPNVQD-KHGIAPVHDAAQTGFLETLQVLVEHGASV |
| *Xiphophorus hellerii* | EKGAEPNVQD-KHGIAPVHDAAQTGFLETLQVLVEHGASV |
| *Xiphophorus maculatus* | EKGAEPNVQD-KHGIAPVHDAAQTGFLETLQVLVEHGASV |

**Cartilaginous fish**

| | |
|---|---|
| *Amblyraja radiata* | KNGADPNIQD-KNGFTPAHDVARTGFLDTMMVLVEYNSDM |
| *Callorhinchus milii* | KHGARPDVQD-QWGFALAHDVARCGFLDTLRVLTEHGANV |
| *Carcharodon carcharias* | KNGAEPNVQD-KWGFTPAHDVARNGFLDTMMVLTKYKADV |
| *Chiloscyllium plagiosum* | KNGAQPNVQD-RWGFTPAHDVARSGFLDTMKVLIKYKANV |
| *Rhincodon typus* | KNGAEPNVQD-KWGFTPAHDVARSGFLDTMKVLTKYKANV |
| *Scyliorhinus canicula* | KNGAKPNAQD-KWGFTPAHDVARCGFLDTMMVLTKYKADV |
| *Stegostoma fasciatum* | KNGAEPNVQD-KWGFTPAHDVARSGFLDTMKVLTKYKANV |

**Cdkn2c/d**

**Cyclostomes**

| | |
|---|---|
| *Eptatretus burgeri* | EAGAQPNVRDPSCGLTPAHDAAREGFADTLRLLAQAGAQL |
| *Entosphenus tridentatus* | EAGADPRVQDPVWGLTPAHDAARTGFTDTALWLKMYGAPM |
| *Lethenteron camtschaticum* | EAGADPGVQDPVWGLTPAHDAAREGFTDTALWLKMYGASM |
| *Lethenteron reissneri* | EAGADPGVQDPVWGLTPAHDAAREGFTDTALWLKMYGASM |
| *Petromyzon marinus* | EAGADPGVQDPVWGLTPAHDAAREGFADTALELKMYGAPM |

**Figure S2j**

**Figure S2** Cdkn2 amino acid alignment around the residue corresponding to D84 in vertebrates. Cdkn2a in (a) mammals and (b) non-mammalian vertebrates; Cdkn2b in (c) mammals, (d) non-mammalian/non-fish vertebrates, and (e) fish; Cdkn2c in (f) mammals, (g) non-mammalian/non-fish vertebrates, and (h) fish; Cdkn2d in (i) mammals and (j) non-mammalian vertebrates. The residue corresponding to Cdkn2 D84 is in green if negatively charged or in magenta, if not.
