## Supplementary Tables S1-S11 and Figures S1-S6 for "Evolution of the *Cdk4/6*–*Cdkn2* system in invertebrates": Figure S3.pdf

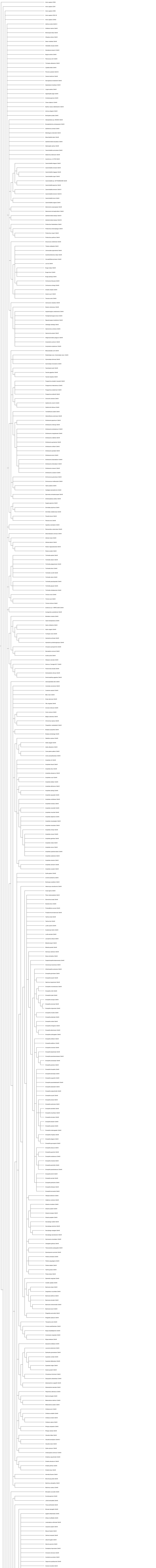
Homo sapiens CDK9
Homo sapiens CDK7
Homo sapiens CDK8
Homo sapiens CDK11B
Homo sapiens CDK20
Aethina tumida Cdk4/6 2
Hofstenia miamia Cdk4/6
Mnemiopsis leidyi Cdk4/6
Oikopleura dioica Cdk4/6
Seison nebaliae Cdk4/6
Helobdella robusta Cdk4/6
Soboliphyme baturini Cdk4/6
Bugula neritina Cdk4/6
Planococcus citri Cdk4/6
Trichoplax adhaerens Cdk4/6
Capitella teleta Cdk4/6
Phoronis australis Cdk4/6 2
Owenia fusiformis Cdk4/6
Saccoglossus kowalevskii Cdk4/6
Epiperipatus broadwayi Cdk4/6
Lingula anatina Cdk4/6
Gigantopelta aegis Cdk4/6
Anneissia japonica Cdk4/6
Cloeon dipterum Cdk4/6
Bacillus rossius redtenbacheri Cdk4/6
Ischnura elegans Cdk4/6
Brachyptera putata Cdk4/6
Halicephalobus sp- NKZ332 Cdk4/6
Bursaphelenchus okinawaensis Cdk4/6
Aphelenchus avenae Cdk4/6
Meloidogyne enterolobii Cdk4/6
Mesorhabditis belari Cdk4/6
Aphelenchoides bicaudatus Cdk4/6
Diploscapter pachys Cdk4/6
Caenorhabditis auriculariae Cdk4/6
Ditylenchus destructor Cdk4/6
Auanema sp- JU1783 Cdk4/6
Caenorhabditis elegans Cdk4/6
Caenorhabditis remanei Cdk4/6
Caenorhabditis briggsae Cdk4/6
Caenorhabditis nigoni Cdk4/6
Caenorhabditis sp- 36 PRJEB534/66 Cdk4/6
Caenorhabditis japonica Cdk4/6
Caenorhabditis brenneri Cdk4/6 1
Caenorhabditis brenneri Cdk4/6 2
Caenorhabditis bovis Cdk4/6
Caenorhabditis angaria Cdk4/6
Steinernema carpocapsae Cdk4/6
Steinernema hermaphroditum Cdk4/6
Aphelenchoides besseyi Cdk4/6 1
Aphelenchoides besseyi Cdk4/6 2
Pristionchus fissidentatus Cdk4/6
Pristionchus entomophagus Cdk4/6
Pristionchus mayeri Cdk4/6
Pristionchus pacificus Cdk4/6
Dracunculus medinensis Cdk4/6
Thelazia callipaeda Cdk4/6
Litomosoides sigmodontis Cdk4/6
Acanthocheilonema viteae Cdk4/6
Cercopithifilaria johnstoni Cdk4/6
Loa loa Cdk4/6
Brugia malayi Cdk4/6
Brugia timori Cdk4/6
Brugia pahangi Cdk4/6
Onchocerca flexuosa Cdk4/6
Onchocerca ochengi Cdk4/6
Anisakis simplex Cdk4/6
Ascaris suum Cdk4/6
Toxocara canis Cdk4/6
Cylicocyclus nassatus Cdk4/6
Necator americanus Cdk4/6
Angiostrongylus costaricensis Cdk4/6

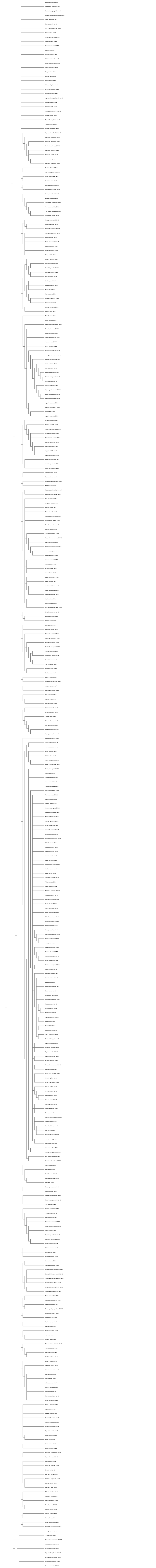
Esperia sulphurella Cdk4/6
Aproaerema taeniolella Cdk4/6
Pectinophora gossypiella Cdk4/6
Hofmannophila pseudospretella Cdk4/6
Apoda limacodes Cdk4/6
Euproctis similis Cdk4/6
Schrankia costaestrigalis Cdk4/6
Orgyia antiqua Cdk4/6
Hypena proboscidalis Cdk4/6
Catocala fraxini Cdk4/6
Lymantria monacha Cdk4/6
Euclidia mi Cdk4/6
Laspeyria flexula Cdk4/6
Trisateles emortualis Cdk4/6
Herminia tarsipennalis Cdk4/6
Carcina quercana Cdk4/6
Pyrgus malvae Cdk4/6
Zeuzera pyrina Cdk4/6
Erynnis tages Cdk4/6
Urbanus simplicius Cdk4/6
Iphiclides podalirius Cdk4/6
Parnassius apollo Cdk4/6
Agonopterix subpropinquella Cdk4/6
Leptidea sinapis Cdk4/6
Limenitis camilla Cdk4/6
Anthocharis cardamines Cdk4/6
Vanessa cardui Cdk4/6
Nymphalis polychloros Cdk4/6
Vanessa atalanta Cdk4/6
Vanessa tameamea Cdk4/6
Gymnoscelis rufifasciata Cdk4/6
Eupithecia centaureata Cdk4/6
Eupithecia abbreviata Cdk4/6
Eupithecia dodoneata Cdk4/6
Eupithecia exiguata Cdk4/6

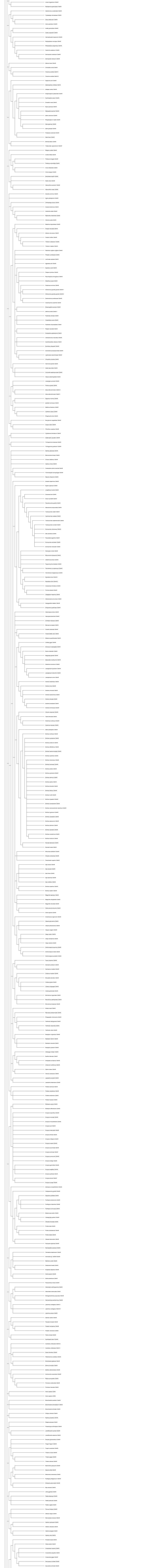

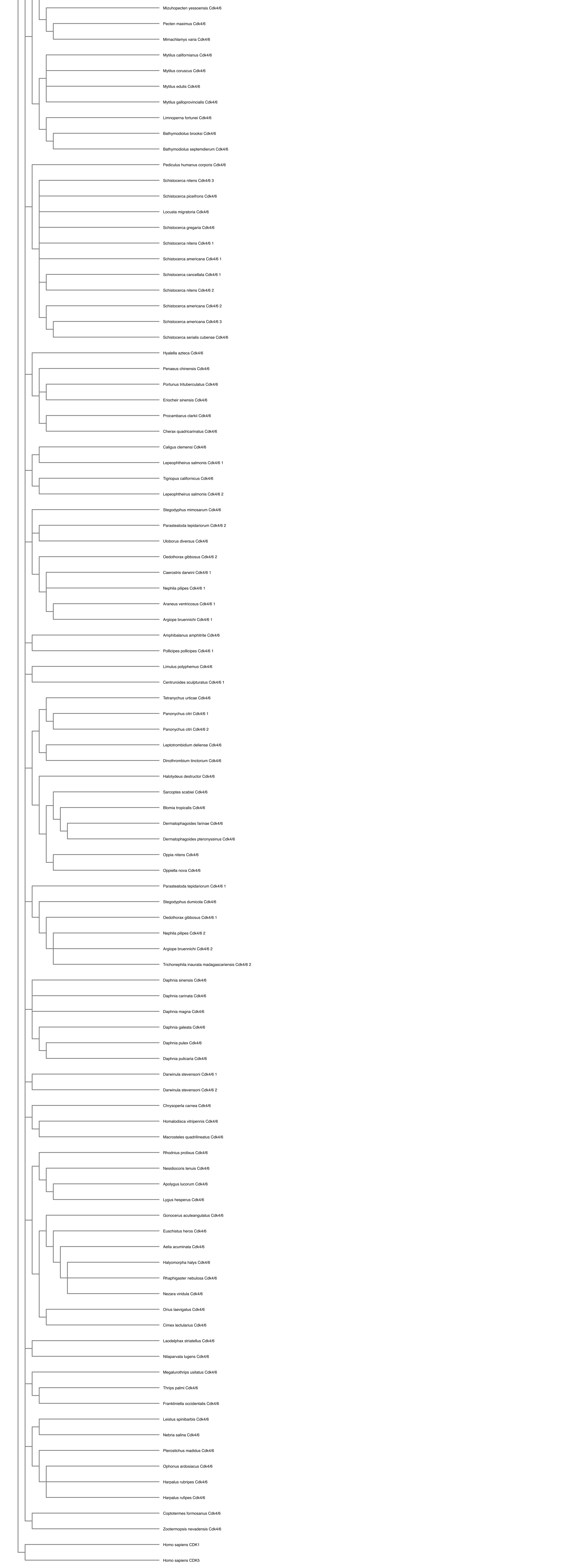
Mizuhopecten yessoensis Cdk4/6
Pecten maximus Cdk4/6
Mimachlamys varia Cdk4/6
Mytilus californianus Cdk4/6
Mytilus coruscus Cdk4/6
Mytilus edulis Cdk4/6
Mytilus galloprovincialis Cdk4/6
Limnoperna fortunei Cdk4/6
Bathymodiolus brooksi Cdk4/6
Bathymodiolus septemdierum Cdk4/6
Pediculus humanus corporis Cdk4/6
Schistocerca nitens Cdk4/6 3
Schistocerca piceifrons Cdk4/6
Locusta migratoria Cdk4/6
Schistocerca gregaria Cdk4/6
Schistocerca nitens Cdk4/6 1
Schistocerca americana Cdk4/6 1
Schistocerca cancellata Cdk4/6 1
Schistocerca nitens Cdk4/6 2
Schistocerca americana Cdk4/6 2
Schistocerca americana Cdk4/6 3
Schistocerca serialis cubense Cdk4/6
Hyalella azteca Cdk4/6
Penaeus chinensis Cdk4/6
Portunus trituberculatus Cdk4/6
Eriocheir sinensis Cdk4/6
Procambarus clarkii Cdk4/6
Cherax quadricarinatus Cdk4/6
Caligus clemensi Cdk4/6
Lepeophtheirus salmonis Cdk4/6 1
Tigriopus californicus Cdk4/6
Lepeophtheirus salmonis Cdk4/6 2
Stegodyphus mimosarum Cdk4/6
Parasteatoda tepidariorum Cdk4/6 2
Uloborus diversus Cdk4/6
Oedothorax gibbosus Cdk4/6 2
Caerostris darwini Cdk4/6 1
Nephila pilipes Cdk4/6 1
Araneus ventricosus Cdk4/6 1
Argiope bruennichi Cdk4/6 1
Amphibalanus amphitrite Cdk4/6
Pollicipes pollicipes Cdk4/6 1
Limulus polyphemus Cdk4/6
Centruroides sculpturatus Cdk4/6 1
Tetranychus urticae Cdk4/6
Panonychus citri Cdk4/6 1
Panonychus citri Cdk4/6 2
Leptotrombidium deliense Cdk4/6
Dinothrombium tinctorium Cdk4/6
Halotydeus destructor Cdk4/6
Sarcoptes scabiei Cdk4/6
Blomia tropicalis Cdk4/6
Dermatophagoides farinae Cdk4/6
Dermatophagoides pteronyssinus Cdk4/6
Oppia nitens Cdk4/6
Oppiella nova Cdk4/6
Parasteatoda tepidariorum Cdk4/6 1
Stegodyphus dumicola Cdk4/6
Oedothorax gibbosus Cdk4/6 1
Nephila pilipes Cdk4/6 2
Argiope bruennichi Cdk4/6 2
Trichonephila inaurata madagascariensis Cdk4/6 2
Daphnia sinensis Cdk4/6
Daphnia carinata Cdk4/6
Daphnia magna Cdk4/6
Daphnia galeata Cdk4/6
Daphnia pulex Cdk4/6
Daphnia pulicaria Cdk4/6
Darwinula stevensoni Cdk4/6 1
Darwinula stevensoni Cdk4/6 2
Chrysoperla carnea Cdk4/6
Homalodisca vitripennis Cdk4/6
Macrosteles quadrilineatus Cdk4/6
Rhodnius prolixus Cdk4/6
Nesidiocoris tenuis Cdk4/6
Apolygus lucorum Cdk4/6
Lygus hesperus Cdk4/6
Gonocerus acuteangulatus Cdk4/6
Euschistus heros Cdk4/6
Aelia acuminata Cdk4/6
Halyomorpha halys Cdk4/6
Rhaphigaster nebulosa Cdk4/6
Nezara viridula Cdk4/6
Orius laevigatus Cdk4/6
Cimex lectularius Cdk4/6
Laodelphax striatellus Cdk4/6
Nilaparvata lugens Cdk4/6
Megalurothrips usitatus Cdk4/6
Thrips palmi Cdk4/6
Frankliniella occidentalis Cdk4/6
Leistus spinibarbis Cdk4/6
Nebria salina Cdk4/6
Pterostichus madidus Cdk4/6
Ophonus ardosiacus Cdk4/6
Harpalus rubripes Cdk4/6
Harpalus rufipes Cdk4/6
Coptotermes formosanus Cdk4/6
Zootermopsis nevadensis Cdk4/6
Homo sapiens CDK1
Homo sapiens CDK5

**Figure S3** Phylogenetic analysis of invertebrate Cdk4/6 candidate sequences and human representative Cdk members (CDK1, 4, 5, 6, 7, 8, 9, 11B, and 20).
