## Supplementary Tables S1-S11 and Figures S1-S6 for "Evolution of the *Cdk4/6*–*Cdkn2* system in invertebrates": Figure S4.pdf

| | |
|---|---|
| **Cdk4** | |
| *Homo sapiens* | RYEPVAEIGVGAYGTVYKARDPHSG---HFVALKSVRVPN |
| **Cdk4/6** | |
| **Urochordata** | |
| *Phallusia fumigata* | RYMPLEEIGVGAYGIVYKARDTSPDTDHGVVAMKCLRVEN |
| *Phallusia mammillare* | RYRPMEEIGVGAYGIVYKARDTSPDADHGVVAMKCLRVEN |
| *Ciona intestinalis* | RYHTLSEIGVGAYGVVFKAQDMQSDDENKFVAIKCVRVEN |
| *Ciona savignyi* | RYHTLTEIGVGAYGVVFKAQDRHSEEENKFVAMKCIRVEN |
| *Corella inflata* | RYKPVEKIGVGAYGVVFKAKDLRKD--GEFVAMKCLRVEN |
| *Molgula oculata* | KYEKDEEIGVGAYGTVYRGKDLGNL--GKYVALKCLRVQT |
| *Botrylloides leachii* | NYEMVAEIGVGAYGVVYRARDLEND--RKLVALKCLRVQN |
| *Styela clava* | NYEMVAEIGVGAYGIVYRARDLENN--GRFVALKCLRVQN |
| *Halocynthia aurantium* | NYEMVAEIGVGAYGIVYRARDLEND--NRYVALKCLRVQN |
| *Halocynthia roretzi* | NYEMVAEIGVGAYGIVYRARDLEND--NRYVALKCLRVQN |
| *Oikopleura dioica* | RWVDMEEIGGGAYGVVFRARDRQHD--NRVVALKRLRVQA |
| **Cephalochordata** | |
| *Branchiostoma lanceolatum* | QYEKVAKIGTGAYGTVYKARDLKNG--GEFVALKRVRVQT |
| *Branchiostoma floridae* | QYEKVAKIGTGAYGTVYKARDLKNG--GEFVALKRVRVQT |
| *Branchiostoma belcheri* | QYEKVAKIGTGAYGTVYKARDLKNG--GEFVALKRVRVQT |
| **Hemichordata** | |
| *Saccoglossus kowalevskii* | QYEEVAEIGTGAYGTVYKARDLETG---RYVALKKVRVQT |
| **Echinodermata** | |
| *Strongylocentrotus purpuratus* | RYVQAAEIGSGAYGIVYKARDTETG---HFVALKSVRIPI |
| *Hemicentrotus pulcherrimus* | RYVQAAEIGSGAYGIVYKARDTETG---HFVALKSVRIPI |
| *Paracentrotus lividus* | RYVRAAEIGSGAYGIVYKARDTETG---QFVALKKVRIPI |
| *Heliocidaris erythrogramma* | RYVKAAEIGSGAYGIVYKARDTETG---HFVALKSVRIPI |
| *Heliocidaris tuberculata* | RYVKAAEIGSGAYGIVYKARDTETG---HFVALKSVRIPI |
| *Lytechinus pictus* | RYVQAAEIGSGAYGIVYKARDTETG---HFVALKSVRIPI |
| *Lytechinus variegatus* 1 | RYVQAAEIGSGAYGIVYKARDTETG---HFVALKSVRIPI |
| *Lytechinus variegatus* 2 | RYVQAAEIGSGAYGIVYKARDTETG---HFVALKSVRIPI |
| *Patiria miniata* | NYSVMSEIGTGAYGTVYKARDCESG---KVVALKKVRLHT |
| *Acanthaster planci* | NYAVISEIGTGAYGTVYKARDLETG---KVVALKKVRLHT |
| *Asterias rubens* | NYSVVAEIGTGAYGTVYKARDTENG---KVVALKKVRLHT |
| *Pisaster brevispinus* | NYSVVAEIGTGAYGTVYKARDTENG---RVVALKRVRLHT |
| *Pisaster ochraceus* | NYSVVAEIGTGAYGTVYKARDTENG---RVVALKKVRLHT |
| *Plazaster borealis* | NYSVVAEIGTGAYGTVYKARDTENG---KVVALKKVRLHT |
| *Anneissia japonica* | RYCEIEKIGTGAYGTVYRGRDRVSG---DVVALKRIRVQA |
| **Xenacoelomorpha** | |
| *Hofstenia miamia* | HYSVLEKVGVGAYGEVYRAKDPRTG---GHVALKRVRVFD |

### Figure S4a

| | |
|---|---|
| **Cdk4** | |
| *Homo sapiens* | RYEPVAEIGVGAYGTVYKARDPHS-GHFVALKSVRVPN |
| **Cdk4/6** | |
| **Nematoda** | |
| **<Chromadorea>** | |
| *Pristionchus entomophagus* | TYKQENIIGKGAYGVVYTVTHIHT-GQIFAMKKMEMRV |
| *Pristionchus fissidentatus* | TYKQEAVLGKGAYGVVYTVTHLPT-GRKYAMKKMEMRV |
| *Pristionchus mayeri* | MYKQENVIGKGAYGVVYTVTHLPT-GRKFAMKKMEMRV |
| *Pristionchus pacificus* | TYKQENVIGKGAYGVVYTVTHLPT-GRKFAMKKMEMRV |
| *Mesorhabditis belari* | EYREIKLIGQGAYGAVYQVEHRAT-GQTFALKKVVIRV |
| *Caenorhabditis angaria* | HFQFFQQLGRGAYGVVHHVRSKID-HQEYALKQMEIIS |
| *Caenorhabditis auriculariae* | DYIISEELGKGAYGVVYQVKEIKT-NKEYALKQINVSV |
| *Caenorhabditis bovis* | DYHIVNVLGKGSYGVVYQVRHKNS-HQVYALKQMEFRS |
| *Caenorhabditis brenneri* 1 | DFEILQALGKGAYGNVYCVRNIHD-GKDYALKQMHVAA |
| *Caenorhabditis brenneri* 2 | DFLMKQALGKGAYGNVYRVQNIRD-GKDYALKQISIHA |
| *Caenorhabditis briggsae* | DFKVGEALGKGAYGHVYRVRNMQD-GKDYALKQIVMSA |
| *Caenorhabditis elegans* | DFQIHQALGKGAYGNVYRVRSLHD-GKDYALKQIMISS |
| *Caenorhabditis japonica* | DFSVSHALGKGAYGNVYRVRNNRD-GKDYALKQILISS |
| *Caenorhabditis nigoni* | DFKVGEALGKGAYGHVYRVRNMQD-GKDYALKQIVMSA |
| *Caenorhabditis remanei* | DFQIQQALGKGSYGHVYRVRSVRD-GKDYALKQIYMSS |
| *Caenorhabditis* sp. 36 PRJEB53466 | DFAIQKALGKGAYGNVYRVRNNRD-GREYALKQILISS |
| *Auanema sp. JU1783* | EYQQVSLLGKGAYGHVYLVNHMPT-GRQFALKKIQVTV |
| *Diploscapter pachys* | HFDIKEVLGRGAYGMVYRVIDKYT-RKEYALKKVVVQL |
| *Ancylostoma caninum* | EYRELNLLGKGAYGVVYHVTHTPT-MTEYALKKIVVGV |
| *Ancylostoma ceylanicum* | EYRELNLLGKGAYGVVYHVTHTPT-MTEYALKKIVVGV |
| *Necator americanus* | EYRELNLLGKGAYGVVYHVTHTPT-MTEYALKKIVVGV |
| *Heligmosomoides polygyrus* | EYRELNLLGKGAYGVVYHVTHTPT-MTEYALKKIVVGV |
| *Nippostrongylus brasiliensis* | EYRELNLLGKGAYGVVYHVTHTPT-MTEYALKKIVVGV |
| *Angiostrongylus costaricensis* | EYRELNVLGKGAYGVVYHVTHTPT-MTEYALKKIVVGV |
| *Parelaphostrongylus tenuis* | EYRELNLLGKGAYGVVYHVTHTPT-MTEYALKKIVVGV |
| *Cylicocyclus nassatus* | EYRELNLLGKGAYGVVYHVTHTPT-MTEYALKKIVVGV |
| *Haemonchus contortus* | EYRELNLLGKGAYGVVYHVTHTPT-MTEYALKKIVVGV |
| *Haemonchus placei* | EYRELNLLGKGAYGVVYHVTHTPT-MTEYALKKIVVGV |
| *Ostertagia ostertagi* | EYRELNLLGKGAYGVVYHVTHTPT-MNEYALKKIVVGV |
| *Anisakis simplex* | EYEEHGIIGKGAYGVVYLVTHLPT-NKQFALKKMIVKI |
| *Ascaris suum* | DYEEHGIIGKGAYGIVYLVTHLAS-NKQYALKKIIVKI |
| *Toxocara canis* | DYEEHGIIGKGAYGIVYLVTHLPT-NKQFALKKIIVKI |
| *Dracunculus medinensis* | EYEERGVIGRGAYGVVHHVTHKKT-KKEYALKKITVQ- |
| *Acanthocheilonema viteae* | EYEEHRVIGKGAYGVVYLVTHIPS-NIQYALKKMIVRI |
| *Brugia malayi* | EYEEHRVIGKGAYGVVYLVTHIPS-NIQYALKKMIVRI |
| *Brugia pahangi* | EYEEHRVIGKGAYGVVYLVTHIPS-NIQYALKKMIVRI |
| *Brugia timori* | EYEEHRVIGKGAYGVVYLVTHIPS-NIQYALKKMIVRI |
| *Cercopithifilaria johnstoni* | EYEEHRVIGKGAYGVVYLVTHIPS-NIQYALKKMIVRI |
| *Litomosoides sigmodontis* | EYEEHRVIGKGAYGVVYLVTHIPS-NIQYALKKMIVRI |
| *Loa loa* | EYEEHRVIGKGAYGVVYLVTHIPS-NIQYALKKMIVRI |
| *Onchocerca flexuosa* | EYEEHRVIGKGAYGIVYLVTHIPS-NIQYALKKMIVRI |
| *Onchocerca ochengi* | EYEEHRVIGKGAYGIVYLVTHIPS-NIQYALKKMIVRI |
| *Thelazia callipaeda* | DYEEHRVIGKGAYGVVYLVTHIPS-NVQYALKKMVVRV |
| *Halicephalobus* sp. NKZ332 | DFYRGREIGRGAYGVVYDVTEKQT-GDSYVMKVITFPS |
| *Steinernema carpocapsae* | QYHIKGELGKGAYGTVYHAIDRLT-NEEFAMKTIVMRL |
| *Steinernema hermaphroditum* | RYQILRELGKGAYGTVYHARDVDN-GLEYAMKTILMRM |
| *Aphelenchus avenae* | KYMLGIELGRGAYGTVFEITDKFN-GTKHALKQIPMKV |
| *Aphelenchoides besseyi* 1 | NYDIHDELGRGAYGCVYLVTNIKT-GLKFALKQITMKP |
| *Aphelenchoides besseyi* 2 | NYQVQNEIGRGAYGCVYIVTNMQT-KVKFALKRITMKP |
| *Aphelenchoides bicaudatus* | EYETKNRIGQGAYGTVFLVKHLES-QHEYALKRINMKQ |
| *Bursaphelenchus okinawaensis* | NYHIGNTIGIGAYGRVYECTDSET-QRRLVMKVTRIET |
| *Ditylenchus destructor* | DYEMLDRLGEGAYGTVYLVTHLPT-KSQYALKKIQMKS |
| *Meloidogyne enterolobii* | DYKQIRVLGKGAYGEVWLVSNKENPDEQFALKKIPVKI |
| **<Enoplea>** | |
| *Soboliphyme baturini* | EYEEIQEIGAGAYGIVMKGRHRVT-GEVIALKKIPIVA |
| *Trichinella britovi* | RYSELGTIGSGAYGMVIKVKNVDT-GEMVAVKKIVVPI |
| *Trichinella murrelli* | RYSELGTIGSGAYGMVIKVKNVDT-GEMVAVKKIVVPI |
| *Trichinella nativa* | RYSELGTIGSGAYGMVIKVKNVDT-GEMVAVKKIVVPI |
| *Trichinella nelsoni* | RYSELGTIGSGAYGMVIKVKNVDT-GEMVAVKKIVVPI |
| *Trichinella papuae* | RYSELGTIGSGAYGMVIKVKNVDT-GEMVAVKKIVVPI |
| *Trichinella patagoniensis* | RYSELGTIGSGAYGMVIKVKNVDT-GEMVAVKKIVVPI |
| *Trichinella pseudospiralis* | RYSELGTIGSGAYGMVIKVKNVDT-GEMVAVKKIVVPI |
| *Trichinella spiralis* | RYSELGTIGSGAYGMVIKVKNVDT-GEMVAVKKIVVPI |
| *Trichinella zimbabwensis* | RYSELGTIGSGAYGMVIKVKNVDT-GEMVAVKKIVVPI |
| *Trichuris muris* | QYVELSKIGSGAYGLVMKGKNTKT-GEVIALKRMVVRL |
| *Trichuris suis* | QYVELSKIGSGAYGLVMKGRNTKT-GEVIALKRMVVRL |
| *Trichuris trichiura* | QYVELSKIGSGAYGLVMKGRNTKT-GEVIALKRMVVRL |

### Figure S4b

| | |
|---|---|
| **Cdk4** | |
| *Homo sapiens* | RYEPVAEIGVGAYGTVYKARDPHS-G-HFVALKSVRVPN |
| **Cdk4/6** | |
| **Arthropoda** | |
| **<Chelicerata>** | |
| *Blomia tropicalis* | QYEELGLIGSGAYGTVFKARDRNNHG-QMVALKKIRMPL |
| *Dermatophagoides farinae* | KYEEVALIGSGAYGTVYKATDRHNEG-QFVALKKIRIPM |
| *Dermatophagoides pteronyssinus* | KYEQIALIGSGAYGTVYKAADRQNEG-RFVALKKIRIPT |
| *Sarcoptes scabiei* | NYEKIALIGSGAYGMVYKARDLNNEG-QFVALKKIRIPL |
| *Oppia nitens* | NYEEVALIGSGAYGTVFKARDLNNDG-QLVALKKVKVPL |
| *Oppiella nova* | NYEEVALIGSGAYGTVYKARDLNNDG-QLVALKKVRVPL |
| *Leptotrombidium deliense* | ------MIGSGAYGTVYKARDLMNDG-QIVALKKIRVPL |
| *Dinothrombium tinctorium* | HYEEVALIGSGAYGTVYKAKDLSKEG-QIVALKKIRVPL |
| *Panonychus citri* 1 | NYEEIGFIGEGTYGTVFKARDLVNEG-QSVAIKKVNVAL |
| *Panonychus citri* 2 | NYEEIGFIGEGTYGTVFKARDLVNEG-QSVAIKKVNVAL |
| *Tetranychus urticae* | NYEEIGVIGSGAYGTVYKARDLVNEG-QSVAIKKVNVAL |
| *Halotydeus destructor* | NYEEVALIGSGAYGTVYKARDLSNRG-EVVALKKVRIPL |
| *Haemaphysalis longicornis* | AYEEIAQIGTGAFGTVYKARDRQNEG-RFVALKKVRVSL |
| *Hyalomma asiaticum* | AYEEIAQIGTGAFGTVYKARDRQNEG-RFVALKKVRVSL |
| *Ixodes persulcatus* | PYEEVAHIGTGAFGTVYKARDRHNDG-RMVALKKVRVPL |
| *Ixodes scapularis* | PYEEVAHIGTGAFGTVYKARDRHNDG-RMVALKKVRVPL |
| *Dermacentor andersoni* | AYEEIAQIGTGAFGTVYKARDRQNEG-RFVALKKVRVSL |
| *Dermacentor silvarum* | AYEEIAQIGTGAFGTVYKARDRQNEG-RFVALKKVRVSL |
| *Rhipicephalus microplus* | AYEEIAQIGTGAFGTVYKARDRQNEG-RFVALKKVRVSL |
| *Rhipicephalus sanguineus* | AYEEIAQIGTGAFGTVYKARDRQNEG-RFVALKKVRVSL |
| *Tropilaelaps mercedesae* | NYDNFDHIGTGAFGTVFKARDKKNEG-RFVALKKVRMAI |
| *Varroa destructor* | NYDDFNHIGTGAFGTVFKARDKKNEG-RFVALKKVRMAI |
| *Varroa jacobsoni* | NYDDFNHIGTGAFGTVFKARDKKNEG-RFVALKKVRMAI |
| *Galendromus occidentalis* | NYDDFNHIGTGAFGTVYKARDKKNEG-RFVALKKVRVAI |
| *Stegodyphus dumicola* | NYEEVSLIGTGAYGTVYKAKDLANEG-QFVALKKIRVPL |
| *Stegodyphus mimosarum* | SFDDMTLIGNGAYGTVYKARDLANSG-QFVALKKVKVPL |
| *Araneus ventricosus* 1 | ----MTVIGNGAYGTVYKARDLASNG-QYVALKKVTIPL |
| *Argiope bruennichi* 1 | NYDDMTVIGNGAYGTVYKARDLASNG-QYVALKKVTIPL |
| *Argiope bruennichi* 2 | NYEEVSLIGTGAYGTVYKARDLANEG-QFVALKKIRVPL |
| *Caerostris darwini* 1 | NYDEMTVIGNGAYGTVYKARDLASNG-QYVALKKVTIPL |
| *Oedothorax gibbosus* 1 | NYEEVGLIGTGAYGTVYKARDLKKEG-RFVALKKVRVPL |
| *Oedothorax gibbosus* 2 | NYDDMTVIGNGAYGTVYKARDLASNG-QYVALKKVKIPL |
| *Nephila pilipes* 1 | SYDEMTVIGTGAYGTVYKARDLASNG-QYVALKKVTIPL |
| *Nephila pilipes* 2 | NYEEVSLIGTGAYGTVYKARDLANEG-QFVALKKIRVPL |
| *Trichonephila inaurata madagascariensis* 2 | NYEEVSLIGTGAYGTVYKARDLANEG-QFVALKKIRVPL |
| *Parasteatoda tepidariorum* 1 | NYEEVSLIGSGAYGTVYKAKDLAHEG-QFVALKKVRVPL |
| *Parasteatoda tepidariorum* 2 | NFDDVTFIGNGAYGTVYKARDLANNG-QFVALKKVKIPI |
| *Uloborus diversus* | SYEDMTLIGNGAYGTVYKAKDLANNG-QFVALKKVRVPL |
| *Centruroides sculpturatus* 1 | NYEEICVIGTGAYGTVYKARDLANKG-QLVALKKVRVPL |
| *Limulus polyphemus* | KYDGLNFIATGAYGTVYKARDLLNDG-QTVALKKVRIPL |
| **<Crustacea>** | |
| *Daphnia carinata* | HYEEIALIGNGAYGTVYKARDTANEG-QMVALKKVRVPL |
| *Daphnia galeata* | HYEEIALIGNGAYGTVYKARDTANEG-QMVALKKVRVPL |
| *Daphnia magna* | HYEEIALIGNGAYGTVYKARDTANEG-QMVALKKVRVPL |
| *Daphnia pulex* | HYEEIALIGNGAYGTVYKARDTANEG-QMVALKKVRVPL |
| *Daphnia pulicaria* | HYEEIALIGNGAYGTVYKARDTANEG-QMVALKKVRVPL |
| *Daphnia sinensis* | HYEEIALIGNGAYGTVYKARDTANEG-QMVALKKVRVPL |
| *Tigriopus californicus* | HYEELSVIGNGAYGTVYRARDKSTD--TIVALKKMRFTL |
| *Caligus clemensi* | IYEEINRIGKGAYGTVYLARDTQKD--RFVALKKMKFCL |
| *Lepeophtheirus salmonis* 1 | HYEEINRIGKGAYGTVYLARDTQKE--RFVALKKMKFCL |
| *Lepeophtheirus salmonis* 2 | HYEELNVIGNGAYGTVYRARNKTDD--TIVALKKMRFSL |
| *Penaeus chinensis* | NYEEVSIIGNGAYGTVYCARDRNNPD-RVVALKKIRITL |
| *Procambarus clarkii* | NYQEISIIGNGAYGTVYYARDRSNLE-RVVALKKIRITL |
| *Cherax quadricarinatus* | SYQEISIIGNGAYGTVYYARDRNNLE-RVVALKKIRITL |
| *Portunus trituberculatus* | KYQEIRVIGNGAYGTVYFAKDRGNQE-RVVALKKIRITL |
| *Eriocheir sinensis* | ---------TGAYGTVYYARDRGNQE-RVVALKKIRITL |
| *Hyalella azteca* | NYEQINLIGNGAYGTVYLGRDLSERG-RYVALKKIKITL |
| *Amphibalanus amphitrite* | NYEEIGLIGNGAYGTVYKARDLRNEG-QMVALKKIRVRL |
| *Pollicipes pollicipes* 1 | NYEEIGMIGNGAYGTVFKARDLRNEG-QMVALKKIRVRL |
| *Darwinula stevensoni* 1 | IYEELDVIGNGAYGTVYKARDRTNPS-QIVALKKVRVPL |
| *Darwinula stevensoni* 2 | IYEKTAMIGMGAYGTVFKARDRTNPS-QNVALKRVRWPS |
| **<Hexapoda>** | |
| **<<Collembola>>** | |
| *Folsomia candida* 1 | LYKKEEAVIGNGAYGTVYKAKDINDPT-RYVALKKVRIPV |
| *Folsomia candida* 2 | LYKKEVVLGQGAYGTVYKAKDITDPT-RYVALKKVIIPV |
| *Orchesella cincta* | LYKEETVIGNGAYGTVYKAKDISHPA-KYVALKKVRIPI |
| *Allacma fusca* | VYKEEAIIGNGAYGTVYKARDINNPS-DYVALKKVRIPL |
| **<<Ephemeroptera>>** | |
| *Cloeon dipterum* | HYEEIHQIGTGAYGTVWKARDLHNPG-KFVALKKVRVPL |
| **<<Odonata>>** | |
| *Ischnura elegans* | KYEEVSVIGTGAYGTVYKAKDLLNPG-VMVALKKVRVPL |
| **<<Plecoptera>>** | |
| *Brachyptera putata* | NYEELAQIGNGAYGTVYKARDLSNNG-VIVALKKVRVPL |
| **<<Phasmatodea>>** | |
| *Bacillus rossius redtenbacheri* | DYEELAVIGNGAYGTVYKARNVASDA--VVAMKKVRVPF |
| **<<Orthoptera>>** | |
| *Locusta migratoria* | NFEMLAQIGNGAYGTVYRARSAN--G-QIVAVKKVRVNL |
| *Schistocerca americana* 1 | NFEMLAQIGNGAYGTVYRARSAS--G-QIVAVKKVRVNL |
| *Schistocerca americana* 2 | NFEMLAQIGNGAYGTVYRARSAS--G-QIVAVKKVRVNL |
| *Schistocerca americana* 3 | NFEVLAQIGNGAYGTVYRARSAS--E-QIVGVKKVRVNL |
| *Schistocerca cancellata* 1 | NFEMLVQIGNGAYGTVYRSRSAS--G-QIVAVKNVRVNP |
| *Schistocerca gregaria* | NFEMLAQIGNGAYGTVYRARSAS--G-QIVAVKKVRVNL |
| *Schistocerca nitens* 1 | NFEMLAQIGNGAYGTVYRARSAS--G-QIVAVKKVRVNL |
| *Schistocerca nitens* 2 | NFEMLAQIGNGAYGTVYRARSAS--G-QIVAVKKVRVNL |
| *Schistocerca nitens* 3 | NSEMLAQIGNGAYGTVYRARSAS--G-QIVAVKKVRVNL |
| *Schistocerca piceifrons* | NFEMLAQIGNGAYSTVYRACSTS--G-QIVAVKKVHVNL |
| *Schistocerca serialis cubense* | NFEVLAQIGNGAYGTVYRARSAS--G-QIVAVKKVRVNL |
| **<<Blattodea>>** | |
| *Coptotermes formosanus* | -----------AYGTVYKARDLSNNG-HVVALKKVRVPL |
| *Zootermopsis nevadensis* | DYEELALIGNGAYGTVYKARDLSNNG-HIVALKKVRVPL |
| **<<Psocodea>>** | |
| *Pediculus humanus corporis* | NYEDLALIGNGAYGTVYKAKNKTN-G-HIVAMKKIRVHL |
| **<<Thysanoptera>>** | |
| *Thrips palmi* | NYEELSMIGVGSYGTVFKARDRSNPN-NIVALKKVRVPL |
| *Frankliniella occidentalis* | NYEELSMIGVGSYGTVFRARDRSNPD-NIVALKKIRVPL |
| *Megalurothrips usitatus* | NYEELSTIGVGSYGTVFKARDRSNPN-NIVALKKVRVPF |
| **<<Hemiptera>>** | |
| *Homalodisca vitripennis* | KYEGLDLIGNGSHGFVYKARDLSNNG-ATVALKRIIVPS |
| *Macrosteles quadrilineatus* | NYEQTKLIGNGSHGFVYKGRDLTNNG-ATVALKRIVIPS |
| *Laodelphax striatellus* | NYEELNLIGNGAYGTVYKARDLSGNN-QIVAIKKLRVPL |
| *Nilaparvata lugens* | NYEELNLIGNGAYGTVYKARDLSGNN-QIVAIKKLRVPL |
| *Gonocerus acuteangulatus* | NYVELDVIGNGAYGTVYKARQRSNEN-AFVALKKIKVPL |
| *Orius laevigatus* | DYVELNVIGNGAYGTVYRGRRIGDPN-TVVAMKKIKVPL |
| *Cimex lectularius* | HYVELNVIGNGAYGTVYKARRGDPN-SVVAMKKIKVPL |
| *Nesidiocoris tenuis* | HYVALGRIGNGAYGTVYKARSTHD-G-AIVALKKIRIHL |
| *Apolygus lucorum* | HYMELDMIGNGAYGTVYKARSVLD-G-SIVALKKIRVQL |
| *Lygus hesperus* | HYMELDMIGNGAYGTVYKARSVLD-G-SIVALKKIRVQL |
| *Rhodnius prolixus* | SYVNLSVIGNGAYGTVYKARHISNPN-LIVAIKKIRLPL |
| *Aelia acuminata* | HYIQLNVIGNGAYGTVYKACHRTNEN-TFVALKKIKVPL |
| *Euschistus heros* | HYIQLNVIGNGAYGTVYKARHRTND--TFVALKKIKVPL |
| *Halyomorpha halys* | HYIQLNVIGNGAYGTVYKACHRTNEN-TFVALKKIKVPL |
| *Nezara viridula* | HYIQLNVIGNGAYGTVYKACHRTNEN-TFVALKKIKVPL |
| *Rhaphigaster nebulosa* | HYIQLNVIGNGAYGTVYKACHRTNEN-TFVALKKIKVPL |
| *Bemisia tabaci* | HYEEVSVIGNGAYGTVYKAKDLSNAG-SIVALKKVKIPL |
| *Trialeurodes vaporariorum* | NYEEVSVIGNGAYGTVYKAKDLSNNG-HIVALKKVRIPL |
| *Aphis craccivora* | NYEELNQIGNGAYGTVYKGRDLVNKG-KFVAMKKIKIPL |
| *Aphis glycines* | NYEELNQIGNGAYGTVYKGRDLVNKG-KFVAMKKIKIPL |
| *Aphis gossypii* | NYEELNQIGNGAYGTVYKGRDLVNKG-KFVAMKKIKIPL |
| *Melanaphis sacchari* | NYEELNQIGNGAYGTVYKGRDLVNKG-KFVAMKKIKIPL |
| *Rhopalosiphum maidis* | NYEELNQIGNGAYGTVYKGRDLVNKG-KFVAMKKIKIPL |
| *Acyrthosiphon pisum* | NYEELNQIGNGAYGTVYKGRDLVNKG-KFVAMKKIKIPL |
| *Diuraphis noxia* | NYEELNQIGNGAYGTVYKGRDLVNKG-KFVAMKKIKIPL |
| *Myzus persicae* | NYEELNQIGNGAYGTVYKGRDLVNKG-KFVAMKKIKIPL |
| *Sipha flava* | NYEELNQIGNGAYGTVYKGRDLVNKG-KFVAMKKIKIPL |
| *Periphyllus acericola* | NYEELNQIGNGAYGTVYKGRDLVNKG-KFVAMKKIKIPL |
| *Drepanosiphum platanoidis* | NYEELNQIGNGAYGTVYKGRDLVNKG-KFVAMKKIKIPL |
| *Adelges cooleyi* | NYEELNQIGNGAYGTVYKGRDLVNKG-KFVAMKKIKIPL |
| *Daktulosphaira vitifoliae* | NYEELNQIGNGAYGTVYKGRDLVNKG-KFVAMKKIKIPL |
| *Planococcus citri* | EFDNMNVIGNGAQGTVYKAREKAT-G-KYVALKRFKISV |
| *Diaphorina citri* | KYDELCQIGCGAYGTVFKARDLQNKG-NYVAMKKLRVKL |
| **<<Neuroptera>>** | |
| *Chrysoperla carnea* | NYEELAIIGTGAYGTVYKARDLSNDG-VIVALKKVRVPL |
| **<<Coleoptera>>** | |
| *Harpalus rubripes* | NYEELNLIGNGAYGTVYKARDRTN-N-LVVALKKVRVPL |
| *Harpalus rufipes* | NYEELNLIGNGAYGTVYKARDRTN-N-LVVALKKVRVPL |
| *Ophonus ardosiacus* | NYEELNLIGNGAYGTVYKARDRTN-N-LVVALKKVRVPL |
| *Pterostichus madidus* | NYEELNLIGNGAYGTVYKARDRTN-N-LVVALKKVRVPL |
| *Leistus spinibarbis* | NYEELNMIGNGAYGTVYKARDRTN-N-LIVALKKVRVPL |
| *Nebria salina* | NYEELNMIGNGAYGTVYKARDRTN-N-LIVALKKVRVPL |
| *Anoplophora glabripennis* | FYEGLNKIGRGAYGTVYKAKDKQT-G-NDVALKKVCVPL |
| *Rutpela maculata* | AYEGLSKIGRGAYGTVYKAKDKQT-G-NDVALKKVCVPL |
| *Acanthoscelides obtectus* | TYEALSRIGRGASGTVKKAKDKKT-G-NDVALKKVIIPL |
| *Bruchidius siliquastri* | TYEALSRIGRGASGTVKKAKDRKS-G-NDVALKKVIIPL |
| *Callosobruchus maculatus* | TYEALSRIGRGASGTVYKAKDRKT-G-NDVALKKVIIPL |
| *Chrysolina oricalcia* | TYQGLNRIGRGAYGTVYKATEKNT-G-LDVALKKVCVRL |
| *Gonioctena quinquepunctata* | TYQGLNRIGRGAYGTVYKATEKNT-G-IDVALKKVCVRL |
| *Phaedon cochleariae* | TYEALNRIGRGAYGTVYKAKDKKT-G-NDVALKKVCVPL |
| *Leptinotarsa decemlineata* | MYQGLNRIGRGAYGTVYKATEKTT-G-LDVALKKVCVQL |
| *Crepidodera aurea* | TYEALNRIGRGAYGTVYRAKDKKT-G-KDVALKKVFVPL |
| *Phyllotreta striolata* | TYEALNSIGRGAYGTVYRAKDKKT-G-TDVALKKVYVPL |
| *Psylliodes chrysocephala* | TYEALNRIGRGAYGTVYKAKDKKT-G-NDVALKKVWVPL |
| *Lochmaea capreae* | TYEALNRIGRGAYGTVYKAKDKKT-G-NDVALKKVRCVPL |
| *Diabrotica virgifera virgifera* | SYEPLNRIGRGAYGTVYRARDRKT-G-NLVALRKVCVHL |
| *Agelastica alni* | TYEALNRIGRGAYGTVYRAKDKKT-G-NDVALKKVCVPL |
| *Malachius bipustulatus* | FYEGFHKIGRGAYGTVYRACNKKT-N-QVVALKKICVPL |
| *Adalia bipunctata* | EYVELLTIGRGAYGTVYKARDKNT-G-KIVALKKVVVPL |
| *Coccinella septempunctata* | EYDELLTIGRGAYGTVYKARDKNT-G-KIVALKKVVVPL |
| *Harmonia axyridis* | GYEELLTIGRGAYGTVYKARDKNT-G-NIVALKKVVVSL |
| *Halyzia sedecimguttata* | EYDELLTIGRGAYGTVYKARDKNT-G-KIVALKKVVVPL |
| *Brassicogethes aeneus* | EYESLSRIGSGAYGTVYKAKDKRT-N-RVVALKKVCVAL |
| *Aethina tumida* 1 | EYESLSRIGSGAYGTVYKAKDKRT-N-RVVALKKVCVAL |
| *Aethina tumida* 2 | NYNILEKLGSGSYGTVYKATNTN--G-ELVALKNVTIEV |
| *Apoderus coryli* | VYEAINKIGRGAYGTVYKAKDKQT-G-SDVALKKVCIPL |
| *Ceutorhynchus assimilis* | TYEALNKIGRGAYGTVYKARDKQT-G-QEVALKKVCIPL |
| *Anthonomus grandis grandis* 1 | TYEAINRIGRGAYGTVYKARDKQT-G-LEVALKKVCIPM |
| *Anthonomus grandis grandis* 2 | TYEAINRIGRGAYGTVYKARDKQT-G-LEVALKKVCIPM |
| *Rhynchophorus ferrugineus* | TYEALNKIGRGAYGTVYRARDKQT-G-QEVALKKVCIPL |
| *Sitophilus oryzae* | TYEAINKIGRGAYGTVYKARDKQS-G-QEVALKKVCIPL |
| *Polydrusus cervinus* | TYEAIHKIGRGAYGTVYKAKDKDT-G-QEVALKKVCIPL |
| *Platypus cylindrus* | TYEAVSTIGRGAYGTVYRARDKQT-G-HEVALKKVCIPL |
| *Dendroctonus ponderosae* | TYEAINKIGRGAYGTVYKARNNET-G-QEVALKKVWIPL |
| *Anaspis maculata* | EEKTLIPIGRGAYGTVYKATDSKT-G-QTVALKKIIVPL |
| *Asbolus verrucosus* | SYYEMGSIGQGAYGVVYKGKHKTT-G-KIVALKKVCVPL |
| *Tribolium castaneum* | LYDEMDSIGRGAYGIVYKGKHKTT-G-KIVALKKISVPL |
| *Tribolium madens* | LYDEMDSIGRGAYGIVYKGKHKTT-G-KIVALKKICVPL |
| *Tenebrio molitor* | PYDEMCSIGRGAYGVVYKGKHKGT-G-RIVALKKICVPL |
| *Agrilus planipennis* | IYEEIDVIGKGAYGMVYKARNRSN-N-TIVALKKVKVPL |
| *Dascillus cervinus* | NYEELNIIGTGAYGMVYKARDRTT-NT-LVALKKVHVPM |
| *Cantharis rustica* | VYEDLNIIGVGAYGMVYKARNRKT-GT-YVALKKVRVAL |
| *Rhagonycha fulva* | VYEDLNIIGVGAYGMVYKARNRKT-G-TYVALKKVRVAL |
| *Malthinus flaveolus* | TYEDLNIIGIGAYGMVYKARNRKN-N-TYVALKKVRVPL |
| *Agrypnus murinus* | VYESLDMIGTGAYGMVYKARNRKN-D-TYVALKKVRVPL |
| *Ignelater luminosus* | VYESLDMIGTGAYGMVYKARNRKN-N-TFVALKKVRVPL |
| *Lamprigera yunnana* | TYEDLNVIGTGAYGTVYKARNRKN-N-SYVALKKIRVSL |
| *Photinus pyralis* | TYEDLNIIGTGAYGTVYKARNRTN-N-TYVALKKIRVSL |
| *Abscondita terminalis* 1 | TYEDLNVIGIGAYGTVYKARNRRN-D-SYVALKKIRVSL |
| *Abscondita terminalis* 2 | TYEILNVIGRGAYGTVYKARDKKN-G-TYVALKKVRVFQ |
| *Cetonia aurata* | KYEQISQIGSGAYGTVFKGKNKKD-N-KVVALKRVRIPI |
| *Oryctes borbonicus* | KYEEINQIGSGAYGTVFKGKNKRN-N-TVVALKRVRIPI |
| *Holotrichia oblita* | EYRYI----SGAYGTVFKGKNKKN-D-AVVALKRVRIPI |
| *Melolontha melolontha* | KYEEINQIGSGAYGTVFKGRNKQT-N-TLVALKRVRIPI |
| *Onthophagus taurus* | KYEEMNVIGSGAYGTVYKGRNKST-N-TFVALKRVRITL |
| *Nicrophorus vespilloides* | NYEEMHIIGTGAYGMVYKARDKTS-N-KMVALKKVKVPL |
| *Ocypus olens* | NYEEMNIIGTGAYGMVFKARDKTN-N-SIVALKKVKVPL |
| *Philonthus cognatus* | NYEEMNIIGTGAYGMVFKARDKTN-N-CIVALKKVKVPL |
| **<<Siphonaptera>>** | |
| *Ctenocephalides felis* | QYEELNVIGTGAYGTVYKARDLHQ-G-TIVALKKVRVAL |
| **<<Diptera>>** | |
| *Bibio marci* | EYEELDLIGTGAYGKVYKARDLNNGG-KIVALKKVKVPL |
| *Contarinia nasturtii* | NYEELSIIGTGAYGTVYKARDLKNPG-KIVALKKVRVAL |
| *Bradysia coprophila* | GYEELNVIGTGAYGTVYKARDRKNAD-QIVALKKVRVAL |
| *Bradysia odoriphaga* | GYEELNVIGTGAYGTVYKARDRKNAD-QIVALKKVRVAL |
| *Clunio marinus* | DFSDLDMIGNGAYGVVYRARDTS--G-RYVALKKIRISL |
| *Culicoides sonorensis* | NYEELNLIASGAYGTVYRARVTDS-G-DIVALKKVRIPL |
| *Chironomus riparius* | DFRDLDVIGNGAYGIVYRARDQTN-G-TYVALKKIRISI |
| *Polypedilum vanderplanki* | DFRDLDVIGNGAYGVVYRARDQTN-G-SYVALKKILISI |
| *Belgica antarctica* | DFRDLDVIGNGAYGIVYRARDTS--G-RYVALKKIRISL |
| *Anopheles atroparvus* | NYEELGVIGTGAYGTVYRARDLKNSG-NIVALKKVRVAL |
| *Anopheles coustani* | NYEELGVIGTGAYGTVYRARDLKNHG-NIVALKKVRVAL |
| *Anopheles ziemanni* | NYEELGVIGTGAYGTVYRARDLKNHG-NIVALKKVRVAL |
| *Anopheles sinensis* | NYEELGVIGTGAYGTVYRARDLKNHG-NIVALKKVRVAL |
| *Anopheles marshallii* | NYEELGVIGTGAYGTVYRARDLKNSG-TIVALKKVRVAL |
| *Anopheles moucheti* | NYEELGVIGTGAYGTVYRARDLKNSG-TIVALKKVRVAL |
| *Anopheles culicifacies* | NYEELEVIGTGAYGTVYRARDLKNSG-TIVALKKVRVAL |
| *Anopheles funestus* | NYEELGVIGTGAYGTVYRARDLKNSG-TIVALKKVRVAL |
| *Anopheles maculipalpis* | NYVELGVIGTGAYGTVYRARDLKNSG-TIVALKKVRVAL |
| *Anopheles stephensi* | NYEELGVIGTGAYGTVYRARDLKNSG-TIVALKKVRVAL |
| *Anopheles maculatus* | NYEELGVIGTGAYGTVYRARDLKNSG-SIVALKKVRVAL |
| *Anopheles nili* | HYEELGVIGTGAYGTVYRARDLKNSG-TIVALKKVRVAL |
| *Anopheles dirus* | NYEELGVIGTGAYGTVYRARDLKNSG-TIVALKKVRVAL |
| *Anopheles farauti* | NYEELGVIGTGAYGTVYRARDLKNAG-TIVALKKVRVAL |
| *Anopheles christyi* | NYEELGVIGTGAYGTVYRARDLKNSG-TIVALKKVRVAL |
| *Anopheles arabiensis* | NYEELGVIGTGAYGTVYRARDLKNAG-TIVALKKVRVAL |
| *Anopheles coluzzii* | NYEELGVIGTGAYGTVYRARDLKNAG-TIVALKKVRVAL |
| *Anopheles gambiae* | NYEELGVIGTGAYGTVYRARDLKNAG-TIVALKKVRVAL |
| *Anopheles melas* | NYEELGVIGTGAYGTVYRARDLKNAG-TIVALKKVRVAL |
| *Anopheles merus* | NYEELGVIGTGAYGTVYRARDLKNAG-TIVALKKVRVAL |
| *Anopheles quadriannulatus* | NYEELGVIGTGAYGTVYRARDLKNAG-TIVALKKVRVAL |
| *Anopheles bellator* | NYEELGIIGTGAYGTVYRARDLKNSG-NIVALKKVRVAL |
| *Anopheles cruzii* | NYEELGLIGTGSYGTVYRARDLKNSG-NIVALKKVRVAL |
| *Anopheles albimanus* | NYEELGVIGTGAYGTVYRARDLKNSG-NIVALKKVRVAL |
| *Anopheles aquasalis* | NYEELGVIGTGAYGTVYRARDLKNSG-NIVALKKVRVAL |
| *Anopheles darlingi* | NYEELGVIGTGAYGTVYRARDLKNSG-NIVALKKVRVAL |
| *Aedes aegypti* | NYEELNVIGTGAYGTVYKARDLSNEG-NIVALKKVRVAL |
| *Aedes albopictus* | NYEELNVIGTGAYGTVYKARDLSNEG-NIVALKKVRVAL |
| *Culex pipiens pallens* | NYEELNVIGTGAYGTVYKARDLGNEG-KIVALKKVRVAL |
| *Culex quinquefasciatus* | NYEELNVIGTGAYGTVYKARDLGNEG-KIVALKKVRVAL |
| *Sabethes cyaneus* | TYEELNVIGTGAYGTVYKARDLSNEG-NIVALKKVRVAL |
| *Machimus atricapillus* | NYEELNVIGTGAYGTVYRARDLNNPG-NIVALKKVRVFQ |
| *Machimus rusticus* | NYEELNVIGTGAYGTVYRARDLNNPG-NIVALKKVRVAL |
| *Villa cingulata* | KYEELNIIGIGAYGTVYRARDLNNPG-KIVALKKVRVAL |
| *Brachyopa scutellaris* | CFVMKHLITQGAYGTVYKARDLNNAG-NIVALKKVRVAL |
| *Eristalinus sepulchralis* | HYEELNVIGTGAYGTVYRARDLKNAG-NIVALKKVRVAL |
| *Eristalis arbustorum* | HYEELNVIGTGAYGTVYRARDLKNAG-NIVALKKVRVAL |
| *Eristalis pertinax* | HYEELNVIGTGAYGTVYRARDLKNAG-NIVALKKVRVAL |
| *Eristalis tenax* | HYEELNVIGTGAYGTVYRARDLKNSG-NIVALKKVRVAL |
| *Cheilosia soror* | NYEELNIIGTGAYGTVYRARDRNNPG-NIVALKKVRVAL |
| *Cheilosia variabilis* | NYEELNIIGTGAYGTVYRARDRNNPG-NIVALKKVRVAL |
| *Cheilosia vernalis* | ----------GAYGTVYRARDRNNPG-NIVALKKVRVAL |
| *Cheilosia vulpina* | NYEELNIIGTGAYGTVYRARDRNNPG-NIVALKKVRVAL |
| *Rhingia campestris* | NYEELNIIGTGAYGTVYRARDRNNPG-NIVALKKVRVAL |
| *Rhingia rostrata* | NYEELNIIGTGAYGTVYRARDRNNPG-NIVALKKVRVAL |
| *Volucella bombylans* 2 | NYEELNIIGTGAYGTVYRARERNNPG-NIVALKKVRVAL |
| *Volucella inanis* | NYEELNIIGTGAYGTVYRARDRNNPG-NIVALKKVRVAL |
| *Volucella inflata* | NYEELNIIGTGAYGTVYRARERNNPG-NIVALKKVRVAL |
| *Chalcosyrphus nemorum* | NYEELNIIGTGAYGTVYRARDINNPA-NIVALKKVRVAL |
| *Criorhina berberina* | FYEELNIIGTGAYGTVYRARDLRNSG-KIVALKKVRVAL |
| *Syritta pipiens* | RCYKGGSKIICAYGTVYRARDLNNAG-NIVALKKVRVGL |
| *Xylota sylvarum* | NYEELNIIGTGAYGTVYRARDINNPA-NIVALKKVRVAL |
| *Baccha elongata* | NYEELNIIGTGAYGTVYRARDLNNPG-NIVALKKVRVAL |
| *Melanostoma mellinum* | FIMSLSNFVQGAYGTVYRARDLNNPG-NIVALKKVRVAL |
| *Melanostoma scalare* | NYEELNIIGTGAYGTVYRARDLNNPG-NIVALKKVRVAL |
| *Platycheirus albimanus* | NYEELNIIGTGAYGTVYRARDLNNPG-NIVALKKVRVAL |
| *Chrysotoxum bicinctum* | NYEELNIIGTGAYGTVYRARDRNNPG-NIVALKKVRVAL |
| *Dasysyrphus albostriatus* | NYEELNIIGTGAYGTVYRARDRNNPG-NIVALKKVRVAL |
| *Epistrophe grossulariae* | TYPNIILLLYGAYGTVYRARDRNNPG-NIVALKKVRVAL |
| *Episyrphus balteatus* | NYEELNIIGTGAYGTVYRARDLNNPG-NIVALKKVRVAL |
| *Eupeodes corollae* | NIVFCFNIIQGAYGTVYRARDRNNPG-NIVALKKVRVAL |
| *Eupeodes latifasciatus* | PSTYIALAVVGAYGTVYRARDRNNPG-NIVALKKVRVAL |
| *Eupeodes luniger* | SYQSKYNTRSCAYGTVYRARDRNNPG-NIVALKKVRVAL |
| *Leucozona laternaria* | NYEELNIIGTGAYGTVYRARDRNNPG-NIVALKKVRVAL |
| *Scaeva pyrastri* | CLFVWIGSFWCAYGTVYRARDRNNPG-NIVALKKVRVAL |
| *Sphaerophoria rueppellii* | NYEELNIIGTGAYGTVYRARDRNNPG-NIVALKKVRVAL |
| *Sphaerophoria taeniata* | NYEELNIIGTGAYGTVYRARDRNNPG-NIVALKKVRVAL |
| *Conops quadrifasciatus* | NYEELNIIGTGAYGTVYRARDNNS-G-QIVALKKVRIPL |
| *Myopa tessellatipennis* | NYEELNIIGTGAYGTVYRARDNNS-G-QIVALKKVRIPL |
| *Myopa testacea* | PDPGLWLLYGSAYGTVYRARDNNS-G-QIVALKKVRIPL |
| *Thecophora atra* | NYEELNIIGTGAYGTVYRARDNSS-G-QIVALKKVRIPL |
| *Teleopsis dalmanni* | ----------GAYGTVYRARDNVT-G-RIVAIKKVRIPL |
| *Chymomyza fuscimana* | NYEELNIIGTGAYGTVYRARDNLT-G-EIVAIKKVRIAL |
| *Scaptodrosophila lebanonensis* | NYENIHVIGTGAYGTVYKARDNTN-G-RIVAIKKVRIAL |
| *Drosophila busckii* | -YTNVSLV-SGAYGTVYRGRDNLT-G-RVVAIKKVRISL |
| *Drosophila rubida* | HYKELDIIGTGAYGTVYRARDNET-G-RYVAIKKVRIPL |
| *Drosophila immigrans* | NYKELDIIGTGAYGTVYRARDNET-G-RYVAIKKVRIPL |
| *Drosophila albomicans* | NYKELDIIGTGAYGTVYRARDNET-G-RYVAIKKVRIPL |
| *Drosophila sulfurigaster* | NYKELDIIGTGAYGTVYRARDNET-G-RYVAIKKVRIPL |
| *Drosophila innubila* | NYKELDMIGKGAYGTVYRARDNHT-G-AFVAIKKLRIAL |
| *Drosophila phalerata* | NYKELDMIGKGAYGTVYRARDNLT-G-AFVAIKKLRISL |
| *Drosophila hydei* | NYKELDIIGTGAYGTVYRARDNLT-G-QFVAIKKVRIAL |
| *Drosophila arizonae* | NYKELDIIGTGAYGTVYRARDNLT-G-KFVAIKKVRITL |
| *Drosophila mojavensis* | NYKELDIIGTGAYGTVYRARDNLT-G-KFVAIKKVRITL |
| *Drosophila navojoa* | NYKELDIIGTGAYGTVYRARDNLT-G-KFVAIKKVRITL |
| *Drosophila novamexicana* | NYKELDIIGTGAYGTVYRARDNIT-G-HFVAIKKVRIAL |
| *Drosophila virilis* | NYKELDIIGTGAYGTVYRARDNIT-G-HFVAIKKVRIAL |
| *Drosophila grimshawi* | NYEELDIIGTGAYGTVYRARDNVT-G-CFVAIKKVRIAL |
| *Drosophila ironensis* | NYEELDVIGTGAYGTVWRARDVVT-G-NIVAIKKVRIAL |
| *Drosophila ananassae* | SYEELNIIGTGAYGTVYKARDVKT-G-NIVAIKKVRISL |
| *Drosophila pandora* | SYEELNIIGTGAYGTVYKARDVKT-G-NIVAIKKVRISL |
| *Drosophila bipectinata* | SYEELNIIGTGAYGTVYKARDVKT-G-NIVAIKKVRISL |
| *Drosophila pseudoananassae* | SYEELNIIGTGAYGTVYKARNIKT-G-NIVAIKKVRISL |
| *Drosophila elegans* | NYQELNIIGTGAYGTVYRARDVMS-G-NIVALKKVRISL |
| *Drosophila gunungcola* | NYQELNIIGTGAYGTVYRARDVMS-G-NIVALKKVRISL |
| *Drosophila eugracilis* | NYQELNIIGTGAYGTVYRARDVIT-G-NIVALKKVRISL |
| *Drosophila ficusphila* | NYQELNIIGTGAYGTVYRARDLIT-G-NIVALKKVRISL |
| *Drosophila erecta* | NYQELNIIGEGAYGTVYRARDVIT-G-NIVALKKVRISL |
| *Drosophila mauritiana* | NYQELNIIGEGAYGTVYRARDVVT-G-NIVALKKVRISL |
| *Drosophila melanogaster* | NYQELNIIGEGAYGTVYRARDVIT-G-NIVALKKVRISL |
| *Drosophila santomea* | NYQELNIIGEGAYGTVYRARDVIT-G-NIVALKKVRISL |
| *Drosophila sechellia* | NYQELNIIGEGAYGTVYRARDVVT-G-NIVALKKVRISL |
| *Drosophila simulans* | NYQELNIIGEGAYGTVYRARDVVT-G-NIVALKKVRISL |
| *Drosophila teissieri* | NYQELNIIGEGAYGTVYRARDVIT-G-NIVALKKVRISL |
| *Drosophila yakuba* | NYQELNIIGEGAYGTVYRARDVIT-G-NIVALKKVRISL |
| *Drosophila jambulina* | NYEELNIIGTGAYGTVFRARDTKT-G-KIVAIKKVRIAL |
| *Drosophila kikkawai* | NYEELNIIGTGAYGTVFRARDTKT-G-KIVAIKKVRIAL |
| *Drosophila birchii* | NYEELNIIGTGAYGTVFRARDMKT-G-KIVAIKKVRIAL |
| *Drosophila bunnanda* | NYEELNIIGTGAYGTVFRARDTKT-G-KIVAIKKVRIAL |
| *Drosophila serrata* | NYEELNIIGTGAYGTVFRARDMKT-G-KIVAIKKVRIAL |
| *Drosophila rhopaloa* | NYQELNIIGTGAYGTVYRARDVIT-G-NIVALKKVRISL |
| *Drosophila biarmipes* | NYQELNIIGTGAYGTVYRARDVIT-G-NIVALKKVRISL |
| *Drosophila subpulchrella* | NYQELNIIGTGAYGTVYRARDVIT-G-NIVALKKVRISL |
| *Drosophila suzukii* | NYQELNIIGTGAYGTVYRARDVIT-G-NIVALKKVRISL |
| *Drosophila pseudotakahashii* | NYQELNIIGTGAYGTVYRAKDVIT-G-NIVALKKVRISL |
| *Drosophila takahashii* | NYQELNIIGTGAYGTVYRARDVIT-G-NIVALKKVRISL |
| *Drosophila guanche* | NYEELNIIGTGAYGTVYRARDTVT-G-NIVALKKVRIAL |
| *Drosophila obscura* | NYEELNIIGTGAYGTVYRARDTIT-G-NIVALKKVRIAL |
| *Drosophila subobscura* | NYEELNIIGTGAYGTVYRARDTVT-G-NIVALKKVRIAL |
| *Drosophila miranda* | DYEELNIIGTGAYGTVYRARDTIT-G-NIVALKKIRIAL |
| *Drosophila persimilis* | DYEELNIIGTGAYGTVYRARDTVT-G-NIVALKKIRIAL |
| *Drosophila pseudoobscura* | DYEELNIIGTGAYGTVYRARDTVT-G-NIVALKKIRIAL |
| *Drosophila setifemur* | NYEELNVIGTGAYGTVYKARDVIT-G-NIVAIKKVRIAL |
| *Drosophila willistoni* | NYQELNIIGTGAYGTVYRARDMIT-G-NIVALKKVRIAL |
| *Hirtodrosophila cameraria* | NYEELDIIDQGAYGTVCRARDNLT-G-RFVAIKKLRIAL |
| *Zaprionus bogoriensis* | ----------GAYGTVYRALDLT-G-RYVAIKKVRIPL |
| *Clusia tigrina* | NYEELNVIGTGAYGTVYRARDNHS-G-KIVAIKKVRIPL |
| *Coremacera marginata* | FHFLHAFVTLRAYGTVYRARDNNS-G-QIVALKKVRIPL |
| *Heteromyza rotundicornis* | NYEELNVIGTGAYGTVYRARDNHT-G-KIVAIKKVRIPL |
| *Ceratitis capitata* | NYEELNVIGTGAYGTVYRAKDNQS-G-KIVAIKKVRIPI |
| *Bactrocera latifrons* | NYEELNVIGTGAYGTVYRAKDNQS-G-KIVAIKKVRIPI |
| *Bactrocera dorsalis* | HYEELNVIGTGAYGTVYRAKDNQS-G-KIVAIKKVRIPI |
| *Bactrocera neohumeralis* | NYEELNVIGTGAYGTVYRAKDNQS-G-KIVAIKKVRIPI |
| *Bactrocera tryoni* | NYEELNVIGTGAYGTVYRAKDNQS-G-KIVAIKKVRIPI |
| *Bactrocera oleae* | NYEELNVIGTGAYGTVYRAKDNQS-G-KIVAIKKVRIPI |
| *Zeugodacus cucurbitae* | NYEELNVIGTGAYGTVYRAKDNQS-G-KIVAIKKVRIPI |
| *Sphenella marginata* | NYEELNVIGTGAYGTVYRARDNQS-G-KIVAIKKVRIPI |
| *Rhagoletis pomonella* | NYEELNVIGTGAYGTVYRARDNQS-G-KIVAIKKVRIPI |
| *Rhagoletis zephyria* | NYEELNVIGTGAYGTVYRARDNQS-G-KIVAIKKVRIPI |
| *Glossina austeni* | NYEELNVIGTGAYGTVYRARDNHS-G-KIVAIKKIRIPL |
| *Glossina morsitans* | NYEELNVIGTGAYGTVYRARDNHS-G-KIVAIKKIRIPL |
| *Glossina fuscipes* | NYEELNVIGTGAYGTVYRARDNHS-G-KIVAIKKIRIPL |
| *Glossina palpalis* | NYEELNVIGTGAYGTVYRARDNHS-G-KIVAIKKIRIPL |
| *Eustalomyia histrio* | NYEELNVIGTGAYGTVYRARDNHT-G-KIVAIKKIRIPL |
| *Leucophora obtusa* | NYEELNVIGTGAYGTVYRARDNHT-G-KIVAIKKVRIPL |
| *Musca domestica* | NYEELNVIGTGAYGTVYRARDNLT-G-RIVAIKKVRITL |
| *Stomoxys calcitrans* | NYEELNVIGTGAYGTVYRARDNLT-G-RIVAIKKVRIPL |
| *Bellardia bayeri* | NYEELNVIGTGAYGTVYRARDNHT-G-KIVAIKKVRIPL |
| *Bellardia pandia* | NYEELNVIGTGAYGTVYRARDNHT-G-KIVAIKKVRIPL |
| *Calliphora vomitoria* | KSFLTLQNQLCAYGTVYRARDNHT-G-KIVAIKKVRIPL |
| *Protocalliphora azurea* | NYEELNVIGTGAYGTVYRARDNHT-G-KIVAIKKVRIPL |
| *Protophormia terraenovae* | NYEELNVIGTGAYGTVYRARDNHT-G-KIVAIKKVRIPL |
| *Lucilia cuprina* | NYEELNVIGTGAYGTVYRARDNHT-G-KIVAIKKVRIPL |
| *Lucilia sericata* | NYEELNVIGTGAYGTVYRARDNHT-G-KIVAIKKVRIPL |
| *Pollenia amentaria* | NYEELNVIGTGAYGTVYRARDNHT-G-KIVAIKKVRIPL |
| *Pollenia angustigena* | NYEELNVIGTGAYGTVYRARDNHT-G-KIVAIKKVRIPL |
| *Pollenia labialis* | NYEELNVIGTGAYGTVYRARDNHT-G-KIVAIKKVRIPL |
| *Stomorhina lunata* | WKCIGLQTGHCAYGTVYRARDNHT-G-KIVAIKKVRIPL |
| *Phyto melanocephala* | NYEELNVIGTGAYGTVYRARDNHT-G-KIVAIKKVRIPL |
| *Sarcophaga rosellei* | NYEELNVIGTGAYGTVYRARDNHT-G-KIVAIKKVRIPL |
| *Sarcophaga caerulescens* | NYEELNVIGTGAYGTVYRARDNHT-G-KIVAIKKVRIAL |
| *Sarcophaga subvicina* | NYEELNVIGTGAYGTVYRARDNHT-G-KIVAIKKVRIPL |
| *Sarcophaga variegata* | NYEELNVIGTGAYGTVYRARDNHT-G-KIVAIKKVRIAL |
| *Epicampocera succincta* | NYEELNVIGTGAYGTVYRARDNHT-G-KIVAIKKVRIPL |
| *Thecocarcelia acutangulata* | NYEELNVIGTGAYGTVYRARDNHT-G-KIVAIKKVRIPL |
| *Gymnosoma rotundatum* | NYEELNVIGTGAYGTVYRARDNHT-G-KIVAIKKVRIPL |
| *Cistogaster globosa* | NYEELNVIGTGAYGTVYRARDSHT-G-KIVAIKKVRIPL |
| *Phasia obesa* | NYEELNVIGTGAYGTVYRARDNHT-G-KIVAIKKVRIPL |
| *Nowickia ferox* | NYEELNVIGTGAYGTVYRARDNHT-G-KIVAIKKVRIPL |
| *Tachina fera* | NYEELNVIGTGAYGTVYRARDNHT-G-KIVAIKKVRIPL |
| *Tachina grossa* | NYEELNVIGTGAYGTVYRARDNHT-G-KIVAIKKVRIPL |
| *Tachina lurida* | NYEELNVIGTGAYGTVYRARDNHT-G-KIVAIKKVRIPL |
| *Empis stercorea* | VYEELDIIGTGAYGTVYKARDK-S-G-KIVALKKVRVNL |
| *Acrocera orbiculus* | NYEELNIIGTGAYGTVYRARDLNNPG-NIVALKKVRVAL |
| *Hermetia illucens* | HYEELNVIGTGAYGTVYRARDLKNPG-HIVALKKVRVPL |
| *Microchrysa polita* | LYEELNVIGTGAYGTVYRARDLKNPG-HIVALKKVRVPL |
| **<<Trichoptera>>** | |
| *Cheumatopsyche charites* | NYKELDVIGTGAYGTVFKAEDSLNPG-VVVAMKRVKVPC |
| *Athripsodes cinereus* | NYEELNLIGTGAYGTVYKARDLLNPG-QVVALKKVRVTL |
| *Glyphotaelius pellucidus* | NYQEINVIGTGAYGTVYKANDLMNPG-QVVALKKVRVAL |
| *Limnephilus lunatus* | NYQEINVIGTGAYGTVYKANDLMNPG-QVVALKKVRVAL |
| *Limnephilus marmoratus* | NYQEINVIGTGAYGTVYKANDLMNPG-QVVALKKVRVAL |
| *Limnephilus rhombicus* | NYQEINVIGTGAYGTVYKANDLMNPG-QVVALKKVRVAL |
| **<<Lepidoptera>>** | |
| *Carposina sasakii* | KYEELNVIGTGAYGTVYKARDLHNGG-QIVAMKKVKVAL |
| *Zeuzera pyrina* | KYEELNVIGTGAYGTVYKARDLHNSG-QIVAMKKVKVAL |
| *Choreutis nemorana* | KYEELNIIGTGAYGTVYKARDLHNGG-QIVAMKKVKVAL |
| *Sesia apiformis* | KYEELNVIGTGAYGTVYKARDLHNSG-QIVAMKKVKVAL |
| *Sesia bembeciformis* | KYEELNVIGTGAYGTVYKARDLYNSG-QIVAMKKVKVAL |
| *Bembecia ichneumoniformis* | NYEELNLIGTGAYGTVYKARDLHNNG-QIVAMKKVKVAL |
| *Synanthedon andrenaeformis* | NYEELNLIGTGAYGTVYKARDLHNNG-QIVAMKKVKVAL |
| *Synanthedon formicaeformis* | NYEELNLIGTGAYGTVYKARDLHNNG-QIVAMKKVKVAL |
| *Synanthedon myopaeformis* | NYEELNLIGTGAYGTVYKARDLHNNG-QIVAMKKVKVAL |
| *Synanthedon tipuliformis* | NYEELNLIGTGAYGTVYKARDLHNNG-QIVAMKKVKVAL |
| *Synanthedon vespiformis* | NYEELNLIGTGAYGTVYKARDLHNNG-QIVAMKKVKVAL |
| *Tortricodes alternella* | KYEELNVIGTGAYGTVYKARDLLKGG-QIVAMKKVKVAL |
| *Epinotia bilunana* | KYEELNVIGTGAYGTVYKARDLQKGG-QIVAMKKVKVAL |
| *Epinotia demarniana* | KYEELNVIGTGAYGTVYKARDLQKGG-QIVAMKKVKLAL |
| *Epinotia nisella* | KYEELNVIGTGAYGTVYKARDLQKGG-QIVAMKKVKVAL |
| *Epinotia ramella* | KYEELNVIGTGAYGTVYKARDLQKGG-QIVAMKKVKVAL |
| *Notocelia uddmanniana* | KYEELNVIGTGAYGTVYKARDLLKGG-QIVAMKKVKVAL |
| *Cydia amplana* | KYEELNVIGTGAYGTVYKARDLLKGG-QIVAMKKVKVAL |
| *Cydia strobilella* | KYEELNVIGTGAYGTVYKARDLLKGG-QIVAMKKVKVAL |
| *Grapholita molesta* | KYEELNVIGTGAYGTVYKARDLLKGG-QIVAMKKVKVAL |
| *Lathronympha strigana* | KYEELNVIGTGAYGTVYKARDLLKGG-QIVAMKKVKVAL |
| *Leguminivora glycinivorella* | KYEELNVIGTGAYGTVYKARDLLKGG-QIVAMKKVKVAL |
| *Pammene aurita* | KYEELNVIGTGAYGTVYKARDLLKGG-QIVAMKKVKVAL |
| *Apotomis betuletana* | KYEELNVIGTGAYGTVYKARDLLKGG-QIVAMKKVKVAL |
| *Apotomis capreana* | KYEELNVIGTGAYGTVYKARDLLKGG-QIVAMKKVKVAL |
| *Apotomis turbidana* | KYEELNVIGTGAYGTVYKARDLLKGG-QIVAMKKVKVAL |
| *Eudemis profundana* | KYEELNVIGTGAYGTVYKARDLLKGG-QIVAMKKVKVAL |
| *Hedya salicella* | KYEELNVIGTGAYGTVYKARDLLKGG-QIVAMKKVKVAL |
| *Archips crataeganus* | KYEELNVIGTGAYGTVYKARDLLKGG-QIVAMKKVKVAL |
| *Archips xylosteana* | KYEELNVIGTGAYGTVYKARDLLKGG-QIVAMKKVKVAL |
| *Choristoneura fumiferana* | KYEELNVIGTGAYGTVYKARDLLKGG-QIVAMKKVKVAL |
| *Pandemis cinnamomeana* | KYEELNVIGTGAYGTVYKARDLLKGG-QIVAMKKVKVAL |
| *Pandemis corylana* | KYEELNVIGTGAYGTVYKARDLLKGG-QIVAMKKVKVAL |
| *Acleris cristana* | KYEELNVIGTGAYGTVYKARDLLKGG-QIVAMKKVKVAL |
| *Acleris emargana* | KYEELNVIGTGAYGTVYKARDLLKGG-QIVAMKKVKVAL |
| *Acleris literana* | KYEELNVIGTGAYGTVYKARDLLKGG-QIVAMKKVKVAL |
| *Acleris sparsana* | KYEELNVIGTGAYGTVYKARDLLKGG-QIVAMKKVKVAL |
| *Apoda limacodes* | KY-NIPLLFLGAYGTVYKARDLHNGG-QIVAMKKVKVAL |
| *Zygaena filipendulae* | KYEELNVIGTGAYGTVYKARDLYNGG-QIVAMKKVKVAL |
| *Oegoconia quadripuncta* | KYEELNVIGTGAYGTVYKARDLHNGG-QIVAMKKVKVAL |
| *Blastobasis adustella* | KYEELNVIGTGAYGTVYKARDLHNGG-TIVAMKKVKVAL |
| *Blastobasis lacticolella* | KYEELNVIGTGAYGTVYKARDLHNGG-QIVAMKKVKVAL |
| *Carcina quercana* | KYEELNVIGTGAYGTVYKARDLHNGG-QIVAMKKVKVAL |
| *Agonopterix subpropinquella* | KYEELNVIGTGAYGTVYKARDLHNGG-QIVAMKKVKVAL |
| *Anarsia innoxiella* | KYEELNVIGTGAYGTVYKARDLHNGG-QIVAMKKVKVAL |
| *Aproaerema taeniolella* | KYEELNVIGTGAYGTVYKARDLHNGG-QIVAMKKVKVAL |
| *Pectinophora gossypiella* | KYEELNVIGTGAYGTVYKARDLHNGG-QIVAMKKVKVAL |
| *Athrips mouffetella* | KYEELNVIGTGAYGTVYKARDLHNG--QIVAMKKVKVAL |
| *Carpatolechia fugitivella* | KYEELNVIGTGAYGTVYKARDLHNAG-QIVAMKKVKVPL |
| *Phthorimaea operculella* | KYEELNVIGTGAYGTVYKARDLHNGG-QIVAMKKVKVAL |
| *Tuta absoluta* | KYEELNVIGTGAYGTVYKARDLHNGG-QIVAMKKVKVAL |
| *Esperia sulphurella* | KYEELNVIGTGAYGTVYKARDLHKGG-QIVAMKKVKVAL |
| *Hofmannophila pseudospretella* | NYEELNVIGTGAYGTVYKARDLHNGG-QIVAMKKVKVAL |
| *Bombyx mandarina* | KYEELSLIGTGAYGTVYKARDLHNGG-QIVAMKKVKVAL |
| *Bombyx mori* | KYEELSLIGTGAYGTVYKARDLHNGG-QIVAMKKVKVAL |
| *Dendrolimus kikuchii* | KYEELNVIGTGAYGTVYKARDLHNGG-QIVAMKKVKVAL |
| *Dendrolimus pini* | KLYKTSYSLSGAYGTVYKARDLHNGG-QIVAMKKVKVAL |
| *Saturnia pavonia* | KYEELNVIGTGAYGTVYKARDLHNGG-QIVAMKKVNVSL |
| *Hemaris fuciformis* | KYEELNVIGTGAYGTVYKARDLHNAG-QIVAMKKVKVPL |
| *Deilephila elpenor* | KYEELNVIGTGAYGTVYKARDLHNAG-QIVAMKKVKVPL |
| *Deilephila porcellus* | KYEELNVIGTGAYGTVYKARDLHNAG-QIVAMKKVKVPL |
| *Hyles euphorbiae* | KYEELNVIGTGAYGTVYKARDLHNAG-QIVAMKKVKVPL |
| *Hyles vespertilio* | KYEELNVIGTGAYGTVYKARDLHNAG-QIVAMKKVKVPL |
| *Amorpha juglandis* | KYEELNVIGTGAYGTVYKARDLHNAG-QIVAMKKVRVPL |
| *Laothoe populi* | KYEELNVIGTGAYGTVYKARDLHNAG-QIVAMKKVRVPL |
| *Mimas tiliae* | KYEELNVIGTGAYGTVYKARDLHNAG-QIVAMKKVRVPL |
| *Lapara coniferarum* | KYEELKIIGTGAYGTVYKARDLHNSG-QIVAMKKVRVPL |
| *Manduca sexta* | KYEELNIIGTGAYGTVYKARDLHNSG-QIVAMKKVRVPL |
| *Sphinx pinastri* | KYEELNIIGTGAYGTVYKARDLHNSG-QIVAMKKVRVPL |
| *Drepana falcataria* | KYEELNVIGTGAYGTVYKARDLHNGG-QIVAMKKVKVAL |
| *Watsonalla binaria* | KYEELNVIGTGAYGTVYKARDLHNGG-QIVAMKKVKVAL |
| *Achlya flavicornis* | KYEELNVIGTGAYGTVYKARDLHNSG-QIVAMKKVKVAL |
| *Habrosyne pyritoides* | KYEELNVIGTGAYGTVYKARDLHNSG-QIVAMKKVKVAL |
| *Ochropacha duplaris* | KYEELNVIGTGAYGTVYKARDLHNSG-QIVAMKKVKVAL |
| *Tetheella fluctuosa* | KYEELNVIGTGAYGTVYKARDLHNSG-QIVAMKKVKVAL |
| *Thyatira batis* | KYEELNVIGTGAYGTVYKARDLHNSG-QIVAMKKVKVAL |
| *Alsophila aescularia* | KYEELNVIGTGAYGTVYKARDLHNGG-QIVAMKKVRVAL |
| *Agriopis aurantiaria* | KYEELNVIGTGAYGTVYKARDLHNGG-QIVAMKKVKVAL |
| *Agriopis leucophaearia* | KYEELNVIGTGAYGTVYKARDLHNGG-QIVAMKKVKVAL |
| *Agriopis marginaria* | KYEELNVIGTGAYGTVYKARDLHNGG-QIVAMKKVKVAL |
| *Alcis repandata* | KYEELNVIGTGAYGTVYKARDLHNGG-QIVAMKKVKVAL |
| *Apeira syringaria* | KYEELNVIGTGAYGTVYKARDLHNGG-QIVAMKKLKVAL |
| *Apocheima hispidaria* | KYEELNVIGTGAYGTVYKARDLHNGG-QIVAMKKVKVAL |
| *Biston betularia* | KYEELNVIGTGAYGTVYKARDLHNGG-QIVAMKKVKVAL |
| *Campaea margaritaria* | KYEELNVIGTGAYGTVYKARDLHNGG-QIVAMKKVRVAL |
| *Crocallis elinguaria* | KYEELNVIGTGAYGTVYKARDLHNGG-QIVAMKKVKVAL |
| *Ectropis grisescens* | KYEELNVIGTGAYGTVYKARDLHNGG-QIVAMKKVKVAL |
| *Ennomos fuscantarius* | KYEELNVIGTGAYGTVYKARDLHNGG-QIVAMKKVKVAL |
| *Ennomos quercinarius* | KYEELNVIGTGAYGTVYKARDLHNGG-QIVAMKKVKVAL |
| *Erannis defoliaria* | KYEELNVIGTGAYGTVYKARDLHNGG-QIVAMKKVKVAL |
| *Hylaea fasciaria* | KYEELNVIGTGAYGTVYKARDLHNGG-QIVAMKKVRVAL |
| *Hypomecis punctinalis* | KYEELNVIGTGAYGTVYKARDLHNGG-QIVAMKKVKVAL |
| *Ligdia adustata* | KYEELNVIGTGAYGTVYKARDLHNGG-QIVAMKKVKVAL |
| *Lomographa bimaculata* | KYEELNVIGTGAYGTVYKARDLHNGG-QIVAMKKVKVAL |
| *Lycia hirtaria* | KYEELNVIGTGAYGTVYKARDLHNGG-QIVAMKKVKVAL |
| *Macaria notata* | KYEELNVIGTGAYGTVYKARDLHNGG-QIVAMKKVKVAL |
| *Opisthograptis luteolata* | KYEELNVIGTGAYGTVYKARDLHNGG-QIVAMKKVKVAL |
| *Peribatodes rhomboidaria* | KYEELNVIGTGAYGTVYKARDLHNGG-QIVAMKKVKVAL |
| *Petrophora chlorosata* | KYEELNVIGTGAYGTVYKARDLHNGG-QIVAMKKVKVAL |
| *Selenia dentaria* | KYEELNVIGTGAYGTVYKARDLHNGG-QIVAMKKVKVAL |
| *Hemistola chrysoprasaria* | KYEELNVIGTGAYGTVYKARDLHNGG-EIVAMKKVKVAL |
| *Hemithea aestivaria* | KYEELNVIGTGAYGTVYKARDLHSGG-QIVAMKKVKVAL |
| *Anticlea derivata* | KYEELNVIGTGAYGTVYKARDLHNGG-QIVAMKKVKVAL |
| *Aplocera efformata* | KYEELNVIGTGAYGVVYKARDLHNGG-QIVAMKKVKVAL |
| *Chesias legatella* | KYEELNVIGTGAYGTVYKARDLHNGG-QIVAMKKVKVAL |
| *Chloroclysta siterata* | KYEELNLIGTGAYGTVYKARDLHNGG-QIVAMKKVKVAL |
| *Colostygia pectinataria* | KYEELNVIGTGAYGTVYKARDLHNGG-QIVAMKKVKVAL |
| *Ecliptopera silaceata* | KYEELNVIGTGAYGTVYKARDLHNGG-QIVAMKKVKVAL |
| *Electrophaes corylata* | KYEELNVIGTGAYGTVYKARDLHNGG-QIVAMKKVKVAL |
| *Epirrhoe tristata* | KYEELNVIGTGAYGTVYKARDLHNGG-QIVAMKKVKVAL |
| *Epirrita christyi* | KYEELNVIGTGAYGTVYKARDLHNGG-QIVAMKKVKVAL |
| *Eulithis prunata* | KYEELNVIGTGAYGTVYKARDLHNGG-QIVAMKKVKVAL |
| *Eulithis testata* | KYEELNVIGTGAYGTVYKARDLHNGG-QIVAMKKVKVAL |
| *Eupithecia abbreviata* | KYEELNVIGTGAYGTVYKARDLHNGG-QIVAMKKVKVAL |
| *Eupithecia centaureata* | KYEELNVIGTGAYGTVYKARDLLNGG-QIVAMKKVKVAL |
| *Eupithecia dodoneata* | KYEELNVIGTGAYGTVYKARDLHNGG-QIVAMKKVKVAL |
| *Eupithecia exiguata* | KYEELNVIGTGAYGTVYKARDLLNGG-QIVAMKKVKVAL |
| *Eupithecia insigniata* | KYEELNVIGTGAYGTVYKARDLHNGG-QIVAMKKVKVAL |
| *Eupithecia subumbrata* | KYEELNVIGTGAYGTVYKARDLHNGG-QIVAMKKVKVAL |
| *Eupithecia vulgata* | KYEELNVIGTGAYGTVYKARDLHNGG-QIVAMKKVKVAL |
| *Gandaritis pyraliata* | KYEELNVIGTGAYGTVYKGRDLHNGG-QIVAMKKVKVAL |
| *Gymnoscelis rufifasciata* | KYEELSVIGTGAYGTVYKARDLQNGG-QLVAMRKVKVAL |
| *Hydriomena furcata* | KYEELNVIGTGAYGTVYKARDLHNGG-QIVAMKKVKVAL |
| *Lampropteryx suffumata* | KYEELNVIGTGAYGTVYKARDLHNNG-QIVAMKKVKVAL |
| *Lobophora halterata* | KYEELNVIGTGAYGTVYKARDLHNGG-QIVAMKKVKVAL |
| *Philereme vetulata* | KYEELNVIGTGAYGTVYKARDLHNGG-QIVAMKKVKVAL |
| *Scotopteryx bipunctaria* | KYEELNVIGTGAYGTVYKARDLHNGG-QIVAMKKVKVAL |
| *Thera britannica* | KYEELNVIGTGAYGTVYKARDLHNGG-QIVAMKKVKVAL |
| *Thera obeliscata* | KYEELNVIGTGAYGTVYKARDLHNGG-QIVAMKKVKVAL |
| *Venusia cambrica* | KYEELNVIGTGAYGTVYKARDLHNGG-QIVAMKKVKVAL |
| *Xanthorhoe spadicearia* | KYEELNVIGTGAYGTVYKARDLHNGG-QIVAMKKVKVAL |
| *Cyclophora punctaria* | KYEELNVIGTGAYGTVYKARDLHNGG-QIVAMKKVKVAL |
| *Idaea aversata* | KYEELNVIGTGAYGTVYKARDLHNGG-QIVALKKVKVAL |
| *Idaea dimidiata* | KYEELNVIGTGAYGTVYKARDLHNGG-QIVAMKKVKVAL |
| *Idaea straminata* | KYEELNVIGTGAYGTVYKARDLHNGG-QIVALKKVKVAL |
| *Urbanus simplicius* | KYEELNVIGTGAYGTVYKARDLHNGG-QIVAMKKVKVAL |
| *Hesperia comma* | KYEELNVIGTGAYGTVYKARDLHNGG-QIVAMKKVKVAL |
| *Ochlodes sylvanus* | KYEELNVIGTGAYGTVYKARDLHNGG-QIVAMKKVKVAL |
| *Thymelicus acteon* | KYEELNVIGTGAYGTVYKARDLHNGG-QIVAMKKVKVAL |
| *Carterocephalus palaemon* | KYEELNVIGTGAYGTVYKARDLHNGG-QIVAMKKVKVAL |
| *Erynnis tages* | KYEELNVIGTGAYGTVYKARDLHNGG-QIVAMKKVKVAL |
| *Pyrgus malvae* | KYEELNVIGTGAYGTVYKARDLHNGG-QIVAMKKVKVAL |
| *Arctia plantaginis* | KYEELNVIGTGAYGTVYKARDLHNGG-QIVAMKKVKVAL |
| *Tyria jacobaeae* | KYEELNVIGTGAYGTVYKARDLHNGG-QIVAMKKVKVAL |
| *Cybosia mesomella* | KYEELNVIGTGAYGTVYKARDLHNGG-QIVAMKKVKVAL |
| *Eilema caniola* | KYEELNVIGTGAYGTVYKARDLHNGG-QIVAMKKVKVAL |
| *Eilema depressum* | KYEELNVIGTGAYGTVYKARDLHNGG-QIVAMKKVKVAL |
| *Eilema sororculum* | KYEELNVIGTGAYGTVYKARDLHNGG-QIVAMKKVKVAL |
| *Miltochrista miniata* | KYEELNVIGTGAYGTVYKARDLLNGG-QIVAMKKVKVAI |
| *Thumatha senex* | KYEELNIIGTGAYGTVYKARDLLNGG-QIVAMKKVKVAI |
| *Diaphora mendica* | KYEELNVIGTGAYGTVYKARDLHNGG-QIVAMKKVKVAL |
| *Hypercompe scribonia* | KYEELNVIGTGAYGTVYKARDLHNGG-QIVAMKKVKVAL |
| *Phragmatobia fuliginosa* | KYEELNVIGTGAYGTVYKARDLHNGG-QIVAMKKVKVAL |
| *Spilarctia lutea* | KYEELNVIGTGAYGTVYKARDLHNGG-QIVAMKKVKVAL |
| *Spilosoma lubricipeda* | KYEELNVIGTGAYGTVYKARDLHNGG-QIVAMKKVKVAL |
| *Catocala fraxini* | KYEELNVIGTGAYGTVYKARDLHNGG-QIVAMKKVKVAL |
| *Euclidia mi* | KYEELNVIGTGAYGTVYKARDLHNGG-QIVAMKKVKVAL |
| *Laspeyria flexula* | KYEELNVIGTGAYGTVYKARDLHNGG-QIVAMKKVKVAL |
| *Herminia tarsipennalis* | KYEELNVIGTGAYGTVYKARDLHNGG-QIVAMKKVKVAL |
| *Hypena proboscidalis* | KYEELNVIGTGAYGTVYKARDLHNGG-QIVAMKKVKVAL |
| *Schrankia costaestrigalis* | KYEELNVIGTGAYGTVYKARDLHNGG-QIVAMKKVKVAL |
| *Euproctis similis* | KYEELNVIGTGAYGTVYKARDLHNGG-QIVAMKKVKVAL |
| *Lymantria monacha* | KYEELNVIGTGAYGTVYKARDLHNGG-QIVAMKKVKVAL |
| *Orgyia antiqua* | KYEELNVIGTGAYGTVYKARDLHNGG-QIVAMKKVKVAL |
| *Trisateles emortualis* | KYEELNVIGTGAYGTVYKARDLHNGG-QIVAMKKVKVAL |
| *Acronicta aceris* | KYEELNVIGTGAYGTVYKARDLHNGG-QIVAMKKVKVAL |
| *Acronicta psi* | KYEELNVIGTGAYGTVYKARDLHNGG-QIVAMKKVKVAL |
| *Craniophora ligustri* | KYEELNIIGTGAYGTVYKARDLHNGG-QIVAMKKVKVAL |
| *Allophyes oxyacanthae* | KYEELNVIGTGAYGTVYKARDLHNGG-QIVAMKKVKVAL |
| *Amphipyra berbera* | KYEELNVIGTGAYGTVYKARDLHNGG-QIVAMKKVKVAL |
| *Amphipyra tragopoginis* | KYEELNVIGTGAYGTVYKARDLHNGG-QIVAMKKVKVAL |
| *Asteroscopus sphinx* | KYEELNVIGTGAYGTVYKARDLHNGG-QIVAMKKVKVAL |
| *Spodoptera exigua* | KYEELNVIGTGAYGTVYKARDLHNGG-NIVAMKKVKVAL |
| *Spodoptera frugiperda* | KYEELNVIGTGAYGTVYKARDLHNGG-NIVAMKKVKVAL |
| *Spodoptera littoralis* | KYEELNVIGTGAYGTVYKARDLHNGG-NIVAMKKVKVAL |
| *Spodoptera litura* | KYEELNVIGTGAYGTVYKARDLHNGG-NIVAMKKVKVAL |
| *Xylocampa areola* | KYEELNVIGTGAYGTVYKARDLHNGG-QIVAMKKVKVAL |
| *Conistra vaccinii* | KYEELNVIGTGAYGTVYKARDLHNGG-NIVAMKKVKVAL |
| *Lithophane leautieri* | KYEELNVIGTGAYGTVYKARDLHNGG-NIVAMKKVKVAL |
| *Lithophane ornitopus* | KYEELNVIGTGAYGTVYKARDLHNGG-NIVAMKKVKVAL |
| *Lithophane semibrunnea* | KYEELNVIGTGAYGTVYKARDLHNGG-NIVAMKKVKVAL |
| *Lithophane socia* | KYEELNVIGTGAYGTVYKARDLHNGG-NIVAMKKVKVAL |
| *Shargacucullia verbasci* | KYEELNVIGTGAYGTVYKARDLHNGG-QIVAMKKVKVAL |
| *Protodeltote pygarga* | KYEELNVIGTGAYGTVYKARDLHNGG-QIVAMKRVKVAL |
| *Anorthoa munda* | KYEELNVIGTGAYGTVYKARDLHNGG-NIVAMKKVKVAL |
| *Euplexia lucipara* | KYEELNVIGTGAYGTVYKARDLHNGG-NIVAMKKVKVAL |
| *Hecatera dysodea* | KYEELNVIGTGAYGTVYKARDLHNGG-NIVAMKKVKVAL |
| *Lacanobia wlatinum* | KYEELNVIGTGAYGTVYKARDLHNGG-NIVAMKKVKVAL |
| *Mamestra brassicae* | KYEELNVIGTGAYGTVYKARDLHNGG-NIVAMKKVKVAL |
| *Melanchra persicariae* | KYEELNVIGTGAYGTVYKARDLHNGG-NIVAMKKVKVAL |
| *Mythimna albipuncta* | KYEELNVIGTGAYGTVYKARDLHNGG-NIVAMKKVKVAL |
| *Mythimna ferrago* | KYEELNVIGTGAYGTVYKARDLHNGG-NIVAMKKVKVAL |
| *Mythimna lalbum* | KYEELNVIGTGAYGTVYKARDLHNGG-NIVAMKKVKVAL |
| *Mythimna separata* | KYEELNVIGTGAYGTVYKARDLHNGG-QIVAMKKVKVAL |
| *Mythimna vitellina* | KYEELNVIGTGAYGTVYKARDLHNGG-NIVAMKKVKVAL |
| *Orthosia gothica* | KYEELNVIGTGAYGTVYKARDLHNGG-NIVAMKKVKVAL |
| *Orthosia gracilis* | KYEELNVIGTGAYGTVYKARDLHNGG-NIVAMKKVKVAL |
| *Orthosia incerta* | KYEELNVIGTGAYGTVYKARDLHNGG-NIVAMKKVKVAL |
| *Phlogophora meticulosa* | KYEELNVIGTGAYGTVYKARDLHNGG-NIVAMKKVKVAL |
| *Tholera decimalis* | KYEELNVIGTGAYGTVYKARDLHNAG-NIVAMKKVKVAL |
| *Helicoverpa armigera* | KYEELNVIGTGAYGTVYKARDLHNGG-QIVAMKKVKVAL |
| *Helicoverpa zea* | KYEELNVIGTGAYGTVYKARDLHNGG-QIVAMKKVKVAL |
| *Cosmia pyralina* | KYEELNVIGTGAYGTVYKARDLHNGG-NIVAMKRVKVAL |
| *Cosmia trapezina* | KYEELNVIGTGAYGTVYKARDLHNGG-NIVAMKRVKVAL |
| *Eupsilia transversa* | KYEELNVIGTGAYGTVYKARDLHNGG-NIVAMKKVKVAL |
| *Amphipoea lucens* | KYEELNVIGTGAYGTVYKARDLHNGG-NIVAMKKVKVAL |
| *Amphipoea oculea* | KYEELNVIGTGAYGTVYKARDLHNGG-NIVAMKKVKVAL |
| *Apamea crenata* | KYEELNVIGTGAYGTVYKARDLHNGG-NIVAMKKVKVAL |
| *Apamea epomidion* | KYEELNVIGTGAYGTVYKARDLHNGG-NIVAMKKVKVAL |
| *Apamea monoglypha* | WYK-INSYITGAYGTVYKARDLHNGG-NIVAMKKVKVAL |
| *Apamea sordens* | KYEELNVIGTGAYGTVYKARDLHNGG-NIVAMKKVKVAL |
| *Hydraecia micacea* | KYEELNVIGTGAYGTVYKARDLHNGG-NIVAMKKVKVAL |
| *Luperina testacea* | KYEELNVIGTGAYGTVYKARDLHNGG-NIVAMKKVKVAL |
| *Oligia latruncula* | KYEELNVIGTGAYGTVYKARDLHNGG-NIVAMKKVKVAL |
| *Caradrina clavipalpis* | KYEELNVIGTGAYGTVYKARDLHNGG-NIVAMKKVKVAL |
| *Caradrina kadenii* | KYEELNVIGTGAYGTVYKARDLHNGG-NIVAMKKVKVAL |
| *Eugnorisma glareosa* | KYEELNVIGTGAYGTVYKARDLHNGG-NIVAMKKVKVAL |
| *Globia sparganii* | KYEELNVIGTGAYGTVYKARDLHNGG-NIVAMKKVKVAL |
| *Agrotis exclamationis* | KYEELNVIGTGAYGTVYKARDLHNGG-NIVAMKKVKVPI |
| *Agrotis puta* | KYEELNVIGTGAYGTVYKARDLHNGG-NIVAMKKVKVAL |
| *Cerastis rubricosa* | KYEELNVIGTGAYGTVYKARDLHNGG-NIVAMKKVKVAL |
| *Diarsia brunnea* | KYEELNVIGTGAYGTVYKARDLHNGG-NIVAMKKVKVAL |
| *Diarsia dahlii* | KYEELNVIGTGAYGTVYKARDLHNGG-NIVAMKKVKVAL |
| *Diarsia rubi* | KYEELNVIGTGAYGTVYKARDLHNGG-NIVAMKKVKVAL |
| *Dicycla oo* | KYEELNVIGTGAYGTVYKARDLHNGG-NIVAMKKVKVAL |
| *Eurois occulta* | KYEELNVIGTGAYGTVYKARDLHNGG-NIVAMKKVKVAL |
| *Lycophotia porphyrea* | KYEELNVIGTGAYGTVYKARDLHNGG-NIVAMKKVKVAL |
| *Noctua fimbriata* | KYEELNVIGTGAYGTVYKARDLHNGG-NIVAMKKVKVAL |
| *Noctua janthe* | KYEELNVIGTGAYGTVYKARDLHNGG-NIVAMKKVKVAL |
| *Noctua pronuba* | KYEELNVIGTGAYGTVYKARDLHNGG-NIVAMKKVKVAL |
| *Ochropleura plecta* | KYEELNVIGTGAYGTVYKARDLHNGG-NIVAMKKVKVAL |
| *Xestia sexstrigata* | KYEELNVIGTGAYGTVYKARDLHNGG-NIVAMKKVKVAL |
| *Xestia xanthographa* | KYEELNVIGTGAYGTVYKARDLHNGG-NIVAMKKVKVAL |
| *Abrostola tripartita* | KYEELNVIGTGAYGTVYKARDLHNGG-QIVAMKKVKVAL |
| *Abrostola triplasia* | KYEELNVIGTGAYGTVYKARDLHNGG-QIVAMKKVKVAL |
| *Autographa gamma* | KYEELNVIGTGAYGTVYKARDLHNGG-QIVAMKKVKVAL |
| *Autographa pulchrina* | KYEELNVIGTGAYGTVYKARDLHNGG-QIVAMKKVKVAL |
| *Plusia festucae* | KYEELNVIGTGAYGTVYKARDLHNGG-QIVAMKKVKVAL |
| *Trichoplusia ni* | KYEELNVIGTGAYGTVYKARDLHNGG-QIVAMKKVKVAL |
| *Agrochola circellaris* | KYEELNVIGTGAYGTVYKARDLHNGG-NIVAMKKVKVAL |
| *Agrochola litura* | KYEELNVIGTGAYGTVYKARDLHNGG-NIVAMKKVKVAL |
| *Agrochola lota* | KYEELNVIGTGAYGTVYKARDLHNGG-NIVAMKKVKVAL |
| *Agrochola macilenta* | KYEELNVIGTGAYGTVYKARDLHNGG-NIVAMKKVKVAL |
| *Antitype chi* | KYEELNVIGTGAYGTVYKARDLHNGG-NIVAMKKVKVAL |
| *Aporophyla lueneburgensis* | KYEELNVIGTGAYGTVYKARDLHNGG-NIVAMKKVKVAL |
| *Aporophyla nigra* | KYEELNVIGTGAYGTVYKARDLHNGG-NIVAMKKVKVAL |
| *Atethmia centrago* | KYEELNVIGTGAYGTVYKARDLHNGG-NIVAMKKVKVAL |
| *Brachylomia viminalis* | KYEELNVIGTGAYGTVYKARDLHNGG-NIVAMKKVKVAL |
| *Charanyca ferruginea* | KYEELNVIGTGAYGTVYKARDLHNGG-NIVAMKKVKVAL |
| *Dryobota labecula* | KYEELNVIGTGAYGTVYKARDLHNGG-NIVAMKKVKVAL |
| *Dryobotodes eremita* | KYEELNVIGTGAYGTVYKARDLHNGG-NIVAMKKVKVAL |
| *Eremobia ochroleuca* | KYEELNVIGTGAYGTVYKARDLHNRG-NIVAMKKVKVAL |
| *Fissipunctia ypsillon* | KYEELNVIGTGAYGTVYKARDLHNGG-NIVAMKKVKVAL |
| *Griposia aprilina* | KYEELNVIGTGAYGTVYKARDLHNGG-SIVAMKKVKVAL |
| *Hoplodrina ambigua* | KYEELNVIGTGAYGTVYKARDLHNGG-NIVAMKKVKVAL |
| *Hoplodrina blanda* | KYEELNVIGTGAYGTVYKARDLHNGG-NIVAMKKVKVAL |
| *Mesoligia furuncula* | KYEELNVIGTGAYGTVYKARDLHNGG-NIVAMKKVKVAL |
| *Omphaloscelis lunosa* | KYEELNVIGTGAYGTVYKARDLHNGG-NIVAMKKVKVAL |
| *Polymixis flavicincta* | KYEELSVIGTGAYGTVYKARDLHNGG-NIVAMKKVKVAL |
| *Polymixis lichenea* | KYEELNVIGTGAYGTVYKARDLHNGG-NIVAMKKVKVAL |
| *Thalpophila matura* | KYEELNVIGTGAYGTVYKARDLHNGG-QIVAMKKVKVAL |
| *Tiliacea aurago* | KYEELNVIGTGAYGTVYKARDLHNGG-NIVAMKKVKVAL |
| *Xanthia icteritia* | KYEELNVIGTGAYGTVYKARDLHNGG-NIVAMKKVKVAL |
| *Pseudoips prasinana* | KYEELNVIGTGAYGTVYKARDLHNGG-QIVAMKKVKVAL |
| *Meganola albula* | KYEELNIIGTGAYGTVYKARDLHNGG-QIVAMKKVKVAL |
| *Notodonta ziczac* | KYEELNVIGTGAYGTVYKARDLHNGG-QIVAMKKVKVAL |
| *Pheosia gnoma* | KYEELNVIGTGAYGTVYKARDLHNGG-QIVAMKKVKVAL |
| *Pheosia tremula* | KYEELNVIGTGAYGTVYKARDLHNGG-QIVAMKKVKVAL |
| *Phalera bucephala* | KYEELNVIGTGAYGTVYKARDLHNGG-QIVAMKKVKVAL |
| *Ptilodon capucinus* | KYEELNVIGTGAYGTVYKARDLHNGG-QIVAMKKVKVAL |
| *Clostera curtula* | KYEELNVIGTGAYGTVYKARDLHNGG-QIVAMKKVKVAL |
| *Furcula furcula* | KYEELNVIGTGAYGTVYKARDLHNGG-QIVAMKKVKVAL |
| *Lycaena phlaeas* | KYEELNVIGTGAYGTVYKARDLHNGG-QIVAMKKVKVAL |
| *Aricia agestis* | KYEELNVIGTGAYGTVYKARDLHNGG-QIVAMKKVKVAL |
| *Aricia artaxerxes* | KYEELNVIGTGAYGTVYKARDLHNGG-QIVAMKKVKVAL |
| *Celastrina argiolus* | KYEELNVIGTGAYGTVYKARDLHNGG-QIVAMKKVKVAL |
| *Cyaniris semiargus* | KYEELNVIGTGAYGTVYKARDLHNGG-QIVAMKKVKVAL |
| *Glaucopsyche alexis* | KYEELNVIGTGAYGTVYKARDLHNGG-QIVAMKKVKVAL |
| *Lysandra bellargus* | KYEELNVIGTGAYGTVYKARDLHNGG-QIVAMKKVKVAL |
| *Lysandra coridon* | KYEELNVIGTGAYGTVYKARDLHNGG-QIVAMKKVKVAL |
| *Plebejus argus* | KYEELNVIGTGAYGTVYKARDLHNGG-QIVAMKKVKVAL |
| *Polyommatus icarus* | KYEELNVIGTGAYGTVYKARDLHNGG-QIVAMKKVKVAL |
| *Danaus chrysippus* | KYEELNVIGTGAYGTVYKARDLHNGG-QIVAMKKVKVAL |
| *Danaus plexippus plexippus* | KYEELNVIGTGAYGTVYKARDLHNGG-QIVAMKKVKVAL |
| *Melinaea menophilus* | KYEELNVIGTGAYGTVYKARDLHNGG-QIVAMKKVKVAL |
| *Melinaea marsaeus rileyi* | KYEELNVIGTGAYGTVYKARDLHNGG-QIVAMKKVKVAL |
| *Boloria selene* | KYEELNVIGTGAYGTVYKARDLHNGG-QIVAMKKVKVAL |
| *Brenthis ino* | KYEELNVIGTGAYGTVYKARDLHNGG-QIVAMKKVKVAL |
| *Fabriciana adippe* | KYEELNVIGTGAYGTVYKARDLHNGG-QIVAMKKVKVAL |
| *Dryas iulia moderata* | KYEELNVIGTGAYGTVYKARDLHNGG-QIVAMKKVKVAL |
| *Eueides isabella* | KYEELNVIGTGAYGTVYKARDLHNGG-QIVAMKKVKVAL |
| *Heliconius melpomene* | KYEELNVIGTGAYGTVYKARDLHNGG-QIVAMKKVKVAL |
| *Heliconius sara* | KYEELNVIGTGAYGTVYKARDLHNGG-QIVAMKKVKVAL |
| *Limenitis camilla* | KYEELNVIGTGAYGTVYKARDLHNGG-QIVAMKKVKVAL |
| *Euphydryas editha* | KYEELNVIGTGAYGTVYKARDLHNGG-QIVAMKKVKVAL |
| *Melitaea cinxia* | KYEELNVIGTGAYGTVYKARDLHNSG-QIVAMKKVKVPI |
| *Mellicta athalia* | KYEELNVIGTGAYGTVYKARDLHNSG-QIVAMKKVKVAL |
| *Nymphalis io (Inachis io)* | KYEELNVIGTGAYGTVYKARDLHNGG-QIVAMKKVKVAL |
| *Nymphalis urticae* | KYEELNVIGTGAYGTVYKARDLHNGG-QIVAMKKVKVAL |
| *Nymphalis polychloros* | KYEELNVIGTGAYGTVYKARDLHNGG-QIVAMKKVKVAL |
| *Vanessa atalanta* | KYEELNVIGTGAYGTVYKARDLHNGG-QIVAMKKVKVAL |
| *Vanessa cardui* | KYEELNVIGTGAYGTVYKARDLHNGG-QIVAMKKVKVAL |
| *Vanessa tameamea* | KYEELNVIGTGAYGTVYKARDLHNGG-QIVAMKKVKVAL |
| *Erebia aethiops* | KYEELNVIGTGAYGTVYKARDLHNGG-QIVAMKKVKVPI |
| *Erebia ligea* | KYEELNVIGTGAYGTVYKARDLHNGG-QIVAMKKVKVAL |
| *Maniola hyperantus* | KYEELNVIGTGAYGTVYKARDLHNGG-QIVAMKKVKVAL |
| *Maniola jurtina* | KYEELNVIGTGAYGTVYKARDLHNGG-QIVAMKKVKVAL |
| *Melanargia galathea* | KYEELNVIGTGAYGTVYKARDLHNGG-QIVAMKKVKVAL |
| *Bicyclus anynana* | KYEELNVIGTGAYGTVYKARDLHNNG-QIVAMKKVKVAL |
| *Lasiommata megera* | KYEELNVIGTGAYGTVYKARDLHNGG-QIVAMKKVKVAL |
| *Pararge aegeria* | KYEELNVIGTGAYGTVYKARDLHNGG-QIVAMKKVKVAL |
| *Hipparchia semele* | KYEELNVIGTGAYGTVYKARDLHNGG-QIVAMKKVKVAL |
| *Iphiclides podalirius* | KYEELNVIGTGAYGTVYKARDLHNGG-QIVAMKKVKVAL |
| *Papilio machaon* | KYEELNVIGTGAYGTVYKARDLHNGG-QIVAMKKVKVAL |
| *Papilio xuthus* | KYEELNVIGTGAYGTVYKARDLHNGG-QIVAMKKVKVAL |
| *Parnassius apollo* | KYEELNVIGTGAYGTVYKARDLHNGG-QIVAMKKVKVAL |
| *Colias croceus* | KYEELNVIGTGAYGTVYKARDLHNGG-QIVAMKKVKVAL |
| *Zerene cesonia* | KYEELNVIGTGAYGTVYKARDLHNGG-QIVAMKKVKVAL |
| *Leptidea sinapis* | KYEELNVIGTGAYGTVYKARDLHNGG-QIVAMKKVKVAL |
| *Anthocharis cardamines* | NYEELNVIGTGAYGTVYKARDLHNGG-QIVAMKKVKVAL |
| *Aporia crataegi* | KYEELNVIGTGAYGTVYKARDLHNGG-TIVAMKKVKVAL |
| *Pieris brassicae* | KYEELNVIGTGAYGTVYKARDLHNGG-QIVAMKKVKVAL |
| *Pieris macdunnoughi* | KYEELNVIGTGAYGTVYKARDLHNGG-QIVAMKKVKVPI |
| *Pieris napi* | KYEELNVIGTGAYGTVYKARDLHNGG-QIVAMKKVKVAL |
| *Pieris rapae* | KYEELNVIGTGAYGTVYKARDLHNGG-QIVAMKKVKVAL |
| *Emmelina monodactyla* | KYEELNVIGTGAYGTVYKARDLHNGG-QIVAMKKVKVAL |
| *Marasmarcha lunaedactyla* | KYEELNVIGTGAYGTVYKARDLHNGG-QIVAMKKVKVAL |
| *Agriphila geniculea* | KYEEINVIGTGAYGTVYKARDLLNEG-HIVAMKKVKVAL |
| *Agriphila straminella* | KYEEINVIGTGAYGTVYKARDLLNEG-HIVAMKKVKVAL |
| *Agriphila tristella* | KYEEINVIGTGAYGTVYKARDLLNEG-HIVAMKKVKVAL |
| *Calamotropha paludella* | KYEEISVIGTGAYGTVYKARDLLNDG-HIVAMKKVKVAL |
| *Callimorpha dominula* | KYEELNVIGTGAYGTVYKARDLHNGG-QIVAMKKVKVAL |
| *Chrysoteuchia culmella* | KYEEINVIGTGAYGTVYKARDLLNEG-HIVAMKKVKVAL |
| *Crambus lathoniellus* | KYEEINVIGTGAYGTVYKARDLLNEG-HIVAMKKVKVAL |
| *Diatraea saccharalis* | KYEEINVIGTGAYGTVYKARDLLNDG-HIVAMKKVKVAL |
| *Musotima nitidalis* | KYEELNVIGTGAYGTVYKARDLHNEG-QIVAMKKVKVAL |
| *Acentria ephemerella* | KYEELSIIGTGAYGTVYKARDLHNEG-QIVAMKKVKVAL |
| *Nymphula nitidulata* | KYEELSIIGTGAYGTVYKARDLHNEG-QIVAMKKVKVAL |
| *Parapoynx stratiotata* | KYEELNVIGTGAYGTVYKARDLHNEG-QIVAMKKVKVAL |
| *Anania crocealis* | KYEELNVIGTGAYGTVYKARDLHNGG-QIVAMKKVKVAL |
| *Cnaphalocrocis medinalis* | KYEELNVIGTGAYGTVYKARDLHNGG-HIVAMKKVKVAL |
| *Marasmia exigua* | KYEELNVIGTGAYGTVYKARDLHNGG-HIVAMKKVKVAL |
| *Ostrinia furnacalis* | KYEELNVIGTGAYGTVYKARDLHNGG-QIVAMKKVKVAL |
| *Pyrausta nigrata* | KYEELNVIGTGAYGTVYKARDLHNGG-QIVAMKKVKVAL |
| *Eudonia lacustrata* | KYEEINVIGTGAYGTVYKARDLHNEG-HIVAMKKVKVAL |
| *Mecyna flavalis* | KYEELNVIGTGAYGTVYKARDLHNGG-HIVAMKKVKVAL |
| *Udea ferrugalis* | KYEELNVIGTGAYGTVYKARDLHNGG-HIVAMKKVKVAL |
| *Galleria mellonella* | KYEELNVIGTGAYGTVYKARDLHNGG-QVVAMKKVKVPI |
| *Acrobasis suavella* | KYEELNVIGTGAYGTVYKARDLHNGG-QVVAMKKVKVAI |
| *Apomyelois bistriatella* | KYEELNVIGTGAYGTVYKARDLHNGG-QVVAMKKVKVAI |
| *Elegia similella* | KYEELNVIGTGAYGTVYKARDLHNGG-QVVAMKKVKVAI |
| *Ephestia elutella* | KYEELNVIGTGAYGTVYKARDLHNGG-QVVAMKKVKVAI |
| *Euzophera pinguis* | KYEELNVIGTGAYGTVYKARDLHNGG-QVVAMKKVKVAI |
| *Plodia interpunctella* | KYEELNVIGTGAYGTVYKARDLHNGG-QVVAMKKVKVAI |
| *Endotricha flammealis* | KYEELNVIGTGAYGTVYKARDLHNGG-QVVAMKKVKVPI |
| *Hypsopygia costalis* | KYEELNVIGIGAYGTVYKARDLHNGG-QVVAMKKVKVPI |
| *Eumeta japonica* | KYEELDVIGTGAYGTVYKARDLMNPE-QIVALKKVKLPL |
| *Luffia ferchaultella* | KYEELDVIGTGAYGTVYKGRDLHNPG-QIVALKKVKVPL |
| *Monopis laevigella* | KYEELNIIGTGAYGTVYKARDLHNSG-QIVAMKKVKVAL |
| *Tinea pellionella* | KYEELNVIGTGAYGTVYKARDLHNSG-QIVAMKKVKVAL |
| *Tinea semifulvella* | KYEELNVIGTGAYGTVYKARDLHNSN-QIVAMKKVKIPL |
| *Tinea trinotella* | KYEELNVIGTGAYGTVYKGRDLHNSG-QFVAMKKVKVAL |
| *Argyresthia goedartella* | KYEELNVIGTGAYGTVYKARDLHNAG-QIVAMKKVKVPL |
| *Plutella xylostella* | KYEELNVIGTGAYGTVYKARDLHNGG-QIVAMKKVKVPL |
| *Yponomeuta cagnagella* | KYEELNVIGTGAYGTVYKARDLHNGG-QIVAMKKVKVPL |
| *Yponomeuta padella* | KYEELNVIGTGAYGTVYKARDLHNGG-QIVAMKKVKVPL |
| *Yponomeuta plumbellus* | KYEELNVIGTGAYGTVYKARDLHNGG-QIVAMKKVKVPL |
| *Yponomeuta sedellus* | KYEELNVIGTGAYGTVYKARDLHNGG-QIVAMKKVKVPL |
| *Zelleria hepariella* | KYEELNVIGTGAYGTVYKARDLHNVG-QIVAMKKVKVPL |
| *Ypsolopha scabrella* | KYEELNVIGTGAYGTVYKARDLHNGG-QIVAMKKVKVPL |
| *Micropterix aruncella* | NSLKVSDAQPCAYGTVYKARDKENQG-KIVALKRLKIVL |
| **<<Hymenoptera>>** | |
| *Ampulex compressa* | LYQELSLIGNGAYGTVYKAKDLNS-G-QVVALKKVRVPL |
| *Andrena fulva* | SYQELSLIGNGAYGTVYKAKDLNT-G-QVVALKKVRVPL |
| *Andrena hattorfiana* | SYQELSLIGNGAYGTVYKAKDLNT-G-QVVALKKVRVPL |
| *Andrena minutula* | SYQELSLIGNGAYGTVYKAKDLNT-G-QVVALKKVRVPL |
| *Andrena bucephala* | SYQELSLIGHGAYGTVYKAKDLNT-G-QIVALKKVRVLL |
| *Andrena trimmerana* | SYQELSLIGHGAYGTVYKAKDLNT-G-QIVALKKVRVPL |
| *Andrena dorsata* | SYQELSLIGNGAYGTVYKAKDLNT-G-QVVALKKVRVPL |
| *Andrena haemorrhoa* | SYQELSLIGNGAYGTVYKAKDLNT-G-QVVALKKVRVPL |
| *Habropoda laboriosa* | SYQELSLIGNGAYGTVYKAKDLNT-G-QVVALKKVRVPL |
| *Apis cerana* | SYQELSLIGNGAYGTVYKAKDLNT-G-QVVALKKVRVPL |
| *Apis dorsata* | SYQELSLIGNGAYGTVYKAKDLNT-G-QVVALKKVRVPL |
| *Apis florea* | SYQELSLIGNGAYGTVYKAKDLNT-G-QVVALKKVRVPL |
| *Apis laboriosa* | SYQELSLIGNGAYGTVYKAKDLNT-G-QVVALKKVRVPL |
| *Apis mellifera* | SYQELSLIGNGAYGTVYKAKDLNT-G-QVVALKKVRVPL |
| *Bombus breviceps* | SYQELSLIGNGAYGTVYKAKDLNT-G-QVVALKKVRVPL |
| *Bombus polaris* | SYQELSLIGNGAYGTVYKAKDLNT-G-QVVALKKVRVPL |
| *Bombus ignitus* | SYQELSLIGNGAYGTVYKAKDLNT-G-QVVALKKVRVPL |
| *Bombus terrestris* | SYQELSLIGNGAYGTVYKAKDLNT-G-QVVALKKVRVPL |
| *Bombus confusus* | SYQELSLIGNGAYGTVYKAKDLNT-G-QVVALKKVRVPL |
| *Bombus cullumanus* | SYQELSLIGNGAYGTVYKAKDLNT-G-QVVALKKVRVPL |
| *Bombus soroeensis* | SYQELSLIGNGAYGTVYKAKDLNT-G-QVVALKKVRVPL |
| *Bombus consobrinus* | SYQELSLIGNGAYGTVYKAKDLNT-G-QVVALKKVRVPL |
| *Bombus hortorum* | SYQELSLIGNGAYGTVYKAKDLNT-G-QVVALKKVRVPL |
| *Bombus pyrosoma* | SYQELSLIGNGAYGTVYKAKDLNT-G-QVVALKKVRVPL |
| *Bombus superbus* | SYQELSLIGNGAYGTVYKAKDLNT-G-QVVALKKVRVPL |
| *Bombus waltoni* | SYQELSLIGNGAYGTVYKAKDLNT-G-QVVALKKVRVPL |
| *Bombus haemorrhoidalis* | SYQELSLIGNGAYGTVYKAKDLNT-G-QVVALKKVRVPL |
| *Bombus campestris* | SYQELSLIGNGAYGTVYKAKDLNT-G-QVVALKKVRVPL |
| *Bombus skorikovi* | SYQELSLIGNGAYGTVYKAKDLNT-G-QVVALKKVRVPL |
| *Bombus sylvestris* | SYQELSLIGNGAYGTVYKAKDLNT-G-QVVALKKVRVPL |
| *Bombus bifarius* | SYQELSLIGNGAYGTVYKAKDLNT-G-QVVALKKVRVPL |
| *Bombus huntii* | SYQELSLIGNGAYGTVYKAKDLNT-G-QVVALKKVRVPL |
| *Bombus hypnorum* | SYQELSLIGNGAYGTVYKAKDLNT-G-QVVALKKVRVPL |
| *Bombus impatiens* | SYQELSLIGNGAYGTVYKAKDLNT-G-QVVALKKVRVPL |
| *Bombus pratorum* | SYQELSLIGNGAYGTVYKAKDLNT-G-QVVALKKVRVPL |
| *Bombus vosnesenskii* | SYQELSLIGNGAYGTVYKAKDLNT-G-QVVALKKVRVPL |
| *Bombus vancouverensis nearcticus* | SYQELSLIGNGAYGTVYKAKDLNT-G-QVVALKKVRVPL |
| *Bombus sibiricus* | SYQELSLIGNGAYGTVYKAKDLNT-G-QVVALKKVRVPL |
| *Bombus difficillimus* | SYQELSLIGNGAYGTVYKAKDLNT-G-QVVALKKVRVPL |
| *Bombus opulentus* | SYQELSLIGNGAYGTVYKAKDLNT-G-QVVALKKVRVPL |
| *Bombus pascuorum* | SYQELSLIGNGAYGTVYKAKDLNT-G-QVVALKKVRVPL |
| *Eufriesea mexicana* | LYQELSLIGNGAYGTVYKAKDLNT-G-QVVALKKVRVPL |
| *Frieseomelitta varia* | LYQELSLIGNGAYGTVYKAKDLNT-G-QVVALKKIRVPL |
| *Heterotrigona itama* | SYQELSLIGNGAYGTVYKAKDLNT-G-QVVALKKVRVPL |
| *Melipona quadrifasciata* | LYQELSLIGNGAYGTVYKAKDLNT-G-QVVALKKVRVPL |
| *Nomada fabriciana* | MYQELSLIGNGAYGTVYKAKDLNT-G-QVVALKKVRVPL |
| *Nomada fucata* | MYQELSLIGNGAYGTVYKAKDLNT-G-QVVALKKVRVPL |
| *Ceratina calcarata* | VYQHLSLIGNGAYGTVYSAKDKNT-G-HVVALKKVRLPL |
| *Colletes gigas* | LYQDLSLIGNGAYGTVYKAKDLNT-G-QVVALKKVRVPL |
| *Megalopta genalis* | FYQELSLIGNGAYGTVYKAKDLNT-G-QVVALKKVRVPL |
| *Lasioglossum morio* | FYQELSLIGNGAYGTVYKAKDLNT-G-QVVALKKVRVPL |
| *Lasioglossum pauxillum* | YYRELSLIGNGAYGTVYKAKDSNT-G-QVVALKKVRVPL |
| *Lasioglossum lativentre* | FYQELSLIGNGAYGTVYKAKDLNT-G-QVVALKKVRVPL |
| *Seladonia tumulorum* | FYQELSLIGNGAYGTVYKAKDLNT-G-QVVALKKVRVPL |
| *Sphecodes monilicornis* | FYQELSLIGNGAYGTVYKAKDLNT-G-QVVALKKVRVPL |
| *Nomia melanderi* | LYQELSLIGNGAYGTVYKAKDLNT-G-QVVALKKVRVPL |
| *Dufourea novaeangliae* | FYQELSLIGNGAYGTVYKAKDLNT-G-QVVALKKVRVPL |
| *Stelis phaeoptera* | SYQDLSLIGNGAYGTVYKAKDLNT-G-QVVALKKVRVPL |
| *Megachile ligniseca* | LYQELSLIGNGAYGTVYKAKDLNT-G-QVVALKKVRVPL |
| *Megachile rotundata* | LYQELSLIGNGAYGTVYKAKDLNT-G-QVVALKKVRVPL |
| *Megachile willughbiella* | LYQELSLIGNGAYGTVYKAKDLNT-G-QVVALKKVRVPL |
| *Osmia lignaria* | LYQELSLIGNGAYGTVYKAKDLNT-G-QVVALKKVRVPL |
| *Osmia bicornis bicornis* | LYQELSLIGNGAYGTVYKAKDLNT-G-QVVALKKVRVPL |
| *Macropis europaea* | LYHELSLIGNGAYGTVYKAKDLNT-G-QVVALKKVRVPL |
| *Nysson spinosus* | LYQELSLIGNGAYGTVYKAKDPNS-G-LIVALKKVRVPL |
| *Ectemnius continuus* | LYQELSLIGNGAYGTVYKAKDLNT-G-QVVALKKVRVPL |
| *Ectemnius lituratus* | LYQELSLIGNGAYGTVYKAKDLNT-G-QVVALKKVRVPL |
| *Pemphredon lugubris* | LYQELSLIGNGAYGTVYKAKDLNT-G-QVVALKKVRVPL |
| *Mimumesa dahlbomi* | LYQELSLIGNGAYGTVYKAKDLNT-G-QVVALKKVRVPL |
| *Cerceris rybyensis* | LYQELSLIGNGAYGTVYKAKDLNT-G-QVVALKKVRVPL |
| *Linepithema humile* | SYEDLFLIGNGAYGTVYRAKDKTS-G-QIVALKKVRVPL |
| *Eciton burchellii* | TYEDLSIIGNGAYGTVYKAKDRSS-G-QVVALKKVRIPL |
| *Ooceraea biroi* | AYETLSSIGGGAYGTVFKAKDKSS-G-QVVALKKIRVPL |
| *Camponotus floridanus* | SYEDLSLIGNGAYGTVYKAKDKSS-G-QIVALKKVRVPL |
| *Cataglyphis hispanica* | SYEDLSLIGNGAYGTVYKAKDRSS-G-QIVALKKVRVPL |
| *Formica exsecta* | SYEDLSLIGNGAYGTVYKAKDKSS-G-QIVALKKVRVPL |
| *Nylanderia fulva* 1 | SYEDLSVIGNGAYGTVYKAKDKSS-G-QIVALKKVRVPL |
| *Nylanderia fulva* 2 | SYEDLSVIGNGAYGTVYKAKDKSS-G-QIVALKKVRVPL |
| *Acromyrmex charruanus* | SFENLSVIGNGAYGTVYKATDKNS-G-QTVALKRVRIPL |
| *Acromyrmex echinatior* | SFENLSVIGNGAYGTVYKATDKNS-G-QTVALKRVRIPL |
| *Acromyrmex insinuator* | SFENLSVIGNGAYGTVYKATDKNS-G-QTVALKRVRIPL |
| *Atta colombica* | SFENLSVIGNGAYGTVYKATDKNS-G-QTVALKRVRIPL |
| *Cyphomyrmex costatus* | SFENLSVIGNGAYGTVYKATDKNS-G-QTVALKRVRIPL |
| *Pseudoatta argentina* | SFENLSVIGNGAYGTVYKATDKNS-G-QTVALKRVRIPL |
| *Trachymyrmex cornetzi* | SFENLSVIGNGAYGTVYKATDKNS-G-QTVALKRVRIPL |
| *Trachymyrmex septentrionalis* | SFENLSVIGNGAYGTVYKATDKNS-G-QTVALKRVRIPL |
| *Trachymyrmex zeteki* | SFENLSVIGNGAYGTVYKATDKNS-G-QTVALKRVRIHL |
| *Wasmannia auropunctata* | SFENLSVIGNGAYGTVYKATDKNS-G-QTVALKRVRIPL |
| *Temnothorax curvispinosus* | SYEDLSLIGNGAYGTVYKAKDKTS-G-QIVALKKVRIPL |
| *Temnothorax longispinosus* | SYEDLSLIGNGAYGTVYKAKDKTS-G-QIVALKKVRIPL |
| *Pogonomyrmex barbatus* | SYEDLSIIGNGAYGTVYKAKDKTS-G-QIVALKKVRIPL |
| *Monomorium pharaonis* | SYEDLSLIGNGAYGTVYKAKDKTS-G-QIVALKKVRVLL |
| *Solenopsis invicta* | SYEDLSLIGNGAYGTVYKAKDRT--G-NAVALKKVRIPL |
| *Vollenhovia emeryi* | SFEDLSLIGNGAYGTVYKAKDKTS-G-QIVALKKVRIPL |
| *Dinoponera quadriceps* | SYEELSLIGNGAYGTVYKAKDKAS-G-QIVALKKVRVPL |
| *Harpegnathos saltator* | SYEELSLIGNGAYGTVYKAKDRTS-G-QVVALKKVRVPL |
| *Odontomachus brunneus* | SYEELSLIGNGAYGTVYKAKDKVS-G-QIVALKKVRVPL |
| *Pseudomyrmex gracilis* | LYEELSLIGNGAYGTVYKAKDKTS-G-QIVALKKVRIPV |
| *Anoplius nigerrimus* | LYQELSLIGNGAYGTVYKAKDLTS-G-QVVALKKVRVPL |
| *Tiphia femorata* | LYQELSLIGNGAYGTVYKAKDLTS-G-QVVALKKVRVPL |
| *Ancistrocerus nigricornis* | SYQELSLIGNGAYGTVYKAKDLTS-G-QVVALKKVRVPL |
| *Polistes canadensis* | LYIQLIYIGNGAYGTVYKARDSNT-G-QVVALKKVRVPL |
| *Polistes dominula* | LYIQLIYIGNGAYGTVYKARDSNT-G-QVVALKKVRIPL |
| *Polistes exclamans* | LYIQLIYIGNGAYGTVYKARDSNT-G-QVVALKKVRVPL |
| *Polistes fuscatus* | LYIQLIYIGNGAYGTVYKARDSNT-G-QVVALKKVRVPL |
| *Dolichovespula media* | SYEELCYIGNGAYGTVYKAKDLGN-G-QVVALKKVRVPL |
| *Dolichovespula saxonica* | SYEELSYIGNGAYGTVYKAKDLGN-G-QVVALKKVRVPL |
| *Dolichovespula sylvestris* | SYEELSYIGNGAYGTVYKAKDLGN-G-QVVALKKVRVPL |
| *Vespa crabro* | SYEELSYIGNGAYGTVYKAKDLTS-G-QVVALKKVRVPL |
| *Vespa mandarinia* | SYEELSYIGNGAYGTVYKAKDLTS-G-QVVALKKVRVPL |
| *Vespa velutina* | SYEELSYIGNGAYGTVYKAKDLTS-G-QVVALKKVRVPL |
| *Vespula germanica* | SYEELSYIGNGAYGTVYKAKDLTS-G-QVVALKKVRVPL |
| *Vespula pensylvanica* | SYEELSYIGNGAYGTVYKAKDLTS-G-QVVALKKVRVPL |
| *Vespula vulgaris* | SYEELSYIGNGAYGTVYKAKDLTS-G-QVVALKKVRVPL |
| *Gasteruption jaculator* | MYRELALIGRGAYGAVFRALNLNT-N-TDVAVKKIRIPF |
| *Aphidius gifuensis* | NYKELSLIGDGAYGTVYKAKDITS-G-NIVALKKVRVPI |
| *Chelonus insularis* | HYQELTLIGDGAYGTVYKAKDSSS-G-QVVALKKVRVPL |
| *Microctonus aethiopoides* | HYEELSMIGDGAYGTVYKAKDLSS-G-QVVALKKIRVPL |
| *Microctonus brassicae* | HYEELSMIGDGAYGTVYKAKDLSS-G-QVVALKKIRVPL |
| *Microctonus hyperodae* | HYEELSMIGDGAYGTVYKAKDLSS-G-QVVALKKIRVPL |
| *Cotesia congregata* | HYQELSQIGDGAYGTVYKAKDLSS-G-QVVALKKVRVPL |
| *Cotesia glomerata* | HYQELSQIGDGAYGTVYKAKDLSS-G-QVVALKKVRVPL |
| *Cotesia typhae* | HYEELSQIGDGAYGTVYKAKDLSS-G-QVVALKKVRVPL |
| *Microplitis demolitor* | HYQELSLIGDGAYGTVYKAKDLSS-G-QVVALKKVRVPL |
| *Diachasma alloeum* | HYQELSQIGDGAYGTVYKAKDVSS-G-QVVALKKVRVSL |
| *Diachasma muliebre* | HYQELSQIGDGAYGTVYKAKDVSS-G-QVVALKKVRVSL |
| *Fopius arisanus* | HYQELSQIGDGAYGTVYKAKDVSS-G-QVVALKKVRVSL |
| *Venturia canescens* | HYEEISVIGDGAYGTVYKAKDLTS-G-QVVALKKVRVPL |
| *Buathra laborator* | HYQELSLIGDGAYGTVYKAKDLSS-G-QVVALKKVRVPI |
| *Amblyteles armatorius* | HYQELSLIGDGAYGTVYKAKDLSS-G-QVVALKKVRVPL |
| *Ichneumon xanthorius* | HYQELSLIGDGAYGTVYKAKDLSS-G-QVVALKKVRVPI |
| *Ophion luteus* | HYEEISLIGDGAYGTVYKARDLTS-G-QVVALKKVRVPL |
| *Clistopyga incitator* | HYQELSLIGDGAYGTVYKAKDLTS-G-QVVALKKVRIPL |
| *Ceratosolen solmsi marchali* | LYHDLSVIGDGAYGTVYKAKDATS-G-QIIALKKVRVPI |
| *Copidosoma floridanum* | PYHELSKIGDGAYGKVFRAKNRVN-G-RVVALKKVRVPV |
| *Nasonia vitripennis* | LYQDLSLIGDGAYGTVYKAKDANS-G-QVVALKKVRVPI |
| *Trichomalopsis sarcophagae* | LYQELALIGDGAYGTVYKAKDATN-G-QVVALKKVRVPI |
| *Trichogramma brassicae* | HYDDLLLIGDGAYGTVYKAHDSVT-R-QVVALKKVRVPC |
| *Trichogramma pretiosum* | HYDDLLLIGDGAYGTVYKAHDSVT-R-QVVALKKVRVPC |
| *Belonocnema kinseyi* | LYEDLLLIGDGAYGTVYKGTNITN-G-NIVALKKVRVPI |
| *Leptopilina boulardi* | FYQELSLIGNGAYGTVYKAKDLTS-G-QVVALKKVRVPL |
| *Leptopilina heterotoma* | FYQELSLIGNGAYGTVYKAKDLTS-G-QVVALKKVRVPL |
| *Cephus cinctus* | LYQELSLIGNGAYGTVYKAKDLTS-G-QVVALKKVRVPL |
| *Orussus abietinus* | HYQELSLIGDGAYGTVYKAKDLTS-G-QVIALKKVRVPL |
| *Athalia rosae* | HYQELSLIGNGAYGTVYKAKDITS-G-QVVALKKVRVPI |
| *Neodiprion fabricii* | HYQELSLIGNGAYGTVYKAKDPAS-G-QVVALKKVRVPI |
| *Neodiprion lecontei* | HYQELSLIGNGAYGTVYKAKDPAS-G-QVVALKKVRVPI |
| *Neodiprion pinetum* | HYQELSLIGNGAYGTVYKAKDPAS-G-QVVALKKVRVPI |
| *Neodiprion virginianus* | HYQELSLIGNGAYGTVYKAKDPAS-G-QVVALKKVRVPI |
| *Macrophya alboannulata* | KYQDLREIGGGAYGTVYRAREGVS-G-PVVALKKVRVPI |
| *Rhogogaster chlorosoma* | EYQTLDEIGNGAYGTVYRARDVASR--QVVALKRVRVPI |
| *Tenthredo distinguenda* | EYQALDEIGNGAYGTVYRARDVTSR--QVVALKRVRVPI |

| | |
|---|---|
| *Tenthredo mesomela* | EYQALDEIGNGAYGTVYRARDVVSR--QVVAMKRVRVPI |
| *Tenthredo notha* | EYQPLDEIGNGAYGTVYRARDVANR--QVVAMKKVRVPI |
| **Onychophora** | |
| *Epiperipatus broadwayi* | PYEEVSVIGTGAYGTVYKARDTLK-G-HYVALKKLRVPY |
| **Tardigrada** | |
| *Hypsibius exemplaris* | LYTPVRELGAGAFGTVYRCVDSES-P-DAVAIKYIRIHD |
| *Ramazzottius varieornatus* | RYNSCKELGSGAFGTVYKCREIDDEAAETVAIKYIRITD |
| **Nematomorpha** | |
| *Gordionus* sp. m RMFG-2023 | AFQDVEIIGSGAEGIVYKARDPNNSD-KFVALKKLIIPL |
| *Acutogordius australiensis* | AFKDLQVIGSGAEGVVYKARDPNNPD-RFVALKKLTVPL |

**Figure S4c**

| | |
|---|---|
| **Cdk4** | |
| *Homo sapiens* | RYEPVAEIGVGAYGTVYKARDPHSGHFVALKSVRVPN |
| **Cdk4/6** | |
| **Platyhelminthes** | |
| **<Trematoda>** | |
| *Schistosoma bovis* | NYQVSNLLGSGAYGRVFKGTSLN-GTTVALKEILIPT |
| *Schistosoma curassoni* | NYQVSNLLGSGAYGRVFKGTSLN-GTTVALKEILIPT |
| *Schistosoma guineensis* | NYQVSNLLGSGAYGRVFKGTSLN-GTTVALKEILIPT |
| *Schistosoma haematobium* | NYQVSNLLGSGAYGRVFKGTSLN-GTTVALKEILIPT |
| *Schistosoma intercalatum* | NYQVSNLLGSGAYGRVFKGTSLN-GTTVALKEILIPT |
| *Schistosoma japonicum* | NYRVSNLLGSGAYGRVFKGTSMN-GTTVALKEILIPT |
| *Schistosoma mansoni* | NYQVSNLLGSGAYGRVFKGTSLN-GTTVALKEILIPT |
| *Schistosoma margrebowiei* | NYQVSNLLGSGAYGRVFKGTSLN-GTTVALKEILIPT |
| *Schistosoma mattheei* | NYQVSNLLGSGAYGRVFKGTSLN-GTTVALKEILIPT |
| *Schistosoma mekongi* | NYRVSNLLGSGAYGRVFKGTSLN-GTTVALKEILIPT |
| *Schistosoma rodhaini* | NYQVSNLLGSGAYGRVFKGTSLN-GTTVALKEILIPT |
| *Schistosoma spindale* | NYQVSNLLGSGAYGRVFKGTSLN-GTTVALKEILIPT |
| *Schistosoma turkestanicum* | NYQVSNLLGSGAYGRVFKGTSLN-GTTVALKEILIPT |
| *Heterobilharzia americana* | NYRVCNLLGSGAYGRVYKGTSLN-GTVVALKEILIPA |
| *Trichobilharzia szidati* | NYRVSTLLGSGAYGRVFKGTSMN-GATVALKEILIPS |
| *Fasciola gigantica* | MYKPSELVGKGTYGRVFKGTSPN-GAVVALKEIIVDP |
| *Fasciola hepatica* | MYKPSELVGKGTYGRVFKGTSPN-GAVVALKEIIVDP |
| *Fasciolopsis buski* | LYKPSELVGKGTYGRVFKGTAPN-GAVVALKEIIVDP |
| *Paragonimus heterotremus* | SYKLAQLLGSGAYGRVFKGTAPN-GATVALKEIILET |
| *Paragonimus kellicotti* | SYKLAQLLGSGAYGRVFKGTAPN-GATVALKEIILET |
| *Paragonimus skrjabini miyazakii* | SYKLAQLLGSGAYGRVFKGTAPN-GATVALKEIILET |
| *Paragonimus westermani* | SYKLAQLLGSGAYGRVFKGTAPN-GATVALKEIILET |
| *Clonorchis sinensis* | SYKISQLLGSGAYGKVFKGTAPN-GTTVALKEIILET |
| *Opisthorchis felineus* | SYKISQLLGSGAYGKVFKGTAPN-GTTVALKEIILET |
| *Opisthorchis viverrini* | SYKISQLLGSGAYGKVFKGTAPN-GTTVALKEIILET |
| **<Cestoda>** | |
| *Hymenolepis diminuta* | KYKLTELLGSGAYGRVYKGIAPD-NMTVAIKEITIPL |
| *Hymenolepis microstoma* | KYKLAELLGSGAYGRVYKGIAPD-NKTVAIKEITIPL |
| *Rodentolepis nana* | KYKLAELLGSGAYGRVYKGKDPD-NKTVAIKEIKIPL |
| *Mesocestoides corti* | RYQPSELLGSGAYGRVYKGFSEE-GEIVALKEVTIPA |
| *Echinococcus granulosus* | KYQPSDLLGAGAYGRVFKGTSPD-GQVVALKEILIPA |
| *Echinococcus multilocularis* | KYQPSDLLGAGAYGRVFKGTSPD-GQVVALKEILIPA |
| *Hydatigera taeniaeformis* | KYQPSDLLGAGAYGRVFKGTAPD-GQVVALKEIVIPA |
| *Taenia asiatica* | KYQPSDLLGAGAYGRVFKGSASD-GQIVALKEILIPA |
| *Spirometra erinaceieuropaei* | NYEVSELLGSGAYGRVFKGTSPN-GLTVALKEIVIPA |
| *Schistocephalus solidus* | NYEVSELLGSGAYGRVFKGTSPN-GLTVALKEIVIPA |
| **<Rhabditophora>** | |
| *Schmidtea mediterranea* | KYRLSEELGSGAYGVVYKGLREDDGKNIALKRIYITD |
| *Schmidtea polychroa* | KYRLSEELGSGAYGVVYKGLREDDGKNIALKRIYITD |
| *Dugesia japonica* | KYRLSEEIGSGAYGVVYKGLREDDGVNIALKRIFITD |
| *Polycelis tenuis* | MYKINEKIGSGAYGEVYKGVREDDGLQVALKRVILTD |
| *Planaria torva* | RYKIYEKIGSGAYGEVYKGTRDDDGLFVALKRVMITD |

**Figure S4d**

| | |
|---|---|
| **Cdk4** | |
| *Homo sapiens* | RYEPVAEIGVGAYGTVYKARDP-HSGHFVALKSVRVPN |
| **Cdk4/6** | |
| **Rotifera** | |
| *Adineta ricciae* | AYELGEIIGQGAYGCVFKSRNV-ATNETVAIKRMSMLR |
| *Adineta steineri* | VYELGEVIGQGAYGCVFKSRNV-TTNETVAIKKMSILR |
| *Rotaria magnacalcarata* | AYELGEIIGQGAYGCVFKSRNV-ATNETVAIKKMSMLR |
| *Rotaria socialis* | AYELGEIIGQGAYGCVFKSRNV-ATNETVAIKKMSMLR |
| *Didymodactylos carnosus* | KYAILEILGEGAFGCVHKARHR-ETNEYVAMKKVHMKK |
| *Brachionus plicatilis* | LYEEISVVGTGAYGTVYKGYDL-KTSNIVAMKRIRIQI |
| *Seison nebaliae* | RYEKTTVIGRGAYGIVFKGIDLVSGHDVVALKQINIPM |
| **Annelida** | |
| *Helobdella robusta* | YPPVKHTIGRGAYATVYKGRDKNVKGRYVAMKEVILPT |
| *Capitella teleta* | HYEIVAEIGRGAYGTVYKARDIYDAGKFVALKEIRIQT |
| *Owenia fusiformis* | DYEEICQIGSGAFGTVFKAKDPNNDGKFVALKKIRVQM |
| *Lamellibrachia luymesi* | VYEEVALIGNGAYGTVYKARDLQN-GRFVALKKIRIPS |
| *Lamellibrachia satsuma* | VYEEVALIGNGAYGTVYKARDLQN-GRFVALKKIRIPS |
| *Ridgeia piscesae* | GYEEVALIGNGAYGTVYKARDLKN-GRFVALKKIRIPS |
| *Paraescarpia echinospica* | VYEEVALIGNGAYGTVYKARDLEN-GRFVALKKIRIPS |
| **Mollusca** | |
| *Serripes groenlandicus* | HYEAISEIGNGAYGTVYKGRDLKNDGQFVAMKLIRIQN |
| *Fragum fragum* | QYEAVSEIGNGAYGTVYKGRDLKNDGQFVAMKLIRIQN |
| *Fragum sueziense* | QYEAVSEIGNGAYGTVYKGRDLRNDGQFVAMKLIRIQN |
| *Tridacna crocea* | RYEAVSEVGNGAYGTVYKGRDLKNDGQFVAMKLIRIQN |
| *Tridacna derasa* | RYEAVSEVGNGAYGTVYKGRDLKNDGQFVAMKLIRIQN |
| *Tridacna gigas* | RYEAVSEVGNGAYGTVYKGRDLKNDGQFVAMKLIRIQN |
| *Dreissena polymorpha* | KYEEVTVIGNGAYGTVYKGRDLNNDGRFVAMKMIRLQN |
| *Mya arenaria* | QYEEVMLIGTGAYGTVYKGRDLNNEGRFVAMKMIRLQN |
| *Mactromeris polynyma* | QYEEVSLIGKGAYGVVYKGRDLKNEGQYVAMKLIRIQN |
| *Spisula solida* | QYEEVSLIGKGAYGVVYKGRDLKNEGQYVAMKMIRIQN |
| *Mercenaria mercenaria* | QYEEVSLIGKGAYGVVYKGRDLKNEGQFVAMKMIRIQN |
| *Ruditapes philippinarum* | QYEEVSLIGKGAYGIVYKGRDLKNEGQFVAMKMIRIQN |
| *Limnoperna fortunei* | NYEVISVIGNGAYGTVYKARD--KDGQLVAMKKIRIQN |
| *Bathymodiolus brooksi* | NYQVISVIGNGAYGTVYKARD--KEGQLVAMKKIRIQN |
| *Bathymodiolus septemdierum* | NYQVISVIGNGAYGTVYKARD--KEGQLVAMKKIRIQN |
| *Mytilus californianus* | NYEVISVIGNGAYGTVYKARD--KEGQLVAMKKIRIQN |
| *Mytilus coruscus* | NYEVISVIGNGAYGTVYKARD--KEGQLVAMKKIRIQN |
| *Mytilus edulis* | NYEVISVIGNGAYGTVYKARD--KEGQLVAMKKIRIQN |
| *Mytilus galloprovincialis* | NYEVISVIGNGAYGTVYKARD--KEGQLVAMKKIRIQN |
| *Crassostrea angulata* | NYEEIDVIGNGAYGTVFKAKDPRNEGSWVAIKRIKIQN |
| *Crassostrea gigas* | NYEEIDVIGNGAYGTVFKAKDPRNEGSWVAIKRIKIQN |
| *Crassostrea virginica* | NYEETDIIGNGAYGTVFKARDPRNEGSWVAVKHIKIQN |
| *Ostrea edulis* | NYEEIDIIGNGAYGTVFKAKDPRNEGNWVAIKHIKIQN |
| *Saccostrea cucullata* | NYEEIDEIGNGAYGTVFKAKDTRNEGGWVAIKRIKIQN |
| *Saccostrea echinata* | NYEEIDEIGNGAYGTVFKAKDTRNEGGWVAIKRIKIQN |
| *Mimachlamys varia* | NYEEISMIGNGAYGTVYKARDLKNDGQIVAMKRIRIQN |
| *Mizuhopecten yessoensis* | NYEEISLIGNGAYGTVYKARDLKNDGQIVAMKRIRIQN |
| *Pecten maximus* | NYEEISMIGNGAYGTVYKARDLKNDGQIVAMKRIRIQN |
| *Pinctada fucata* | NYEEVAVIGTGAYGTVYKARDLKNDGAIVAMKRIRIQN |
| *Marisa cornuarietis* | NYEEVAIIGNGAYGTVFKARDLRNEGQFVAMKRIRICN |
| *Pomacea canaliculata* | NYEEVAIIGNGAYGTVFKARDLRNEGQFVAMKRIRICN |
| *Pomacea maculata* | NYEEVAIIGNGAYGTVFKARDLRNEGQFVAMKRIRICN |
| *Alviniconcha marisindica* | QYEEVAVIGNGAYGTVYKARDLRNDGKFVAMKRIRLCN |
| *Batillaria attramentaria* | RYEEVAVIGNGAYGTVYKARDLKNEGQFVAMKRIRLCN |
| *Candidula unifasciata* 1 | NYEFLSHVGSGAYGQVFKARDKSKNGAFVAIKQIVVRR |
| *Candidula unifasciata* 2 | HYEFLSHIGSGAYGQVFKARDTSRDGAVVAVKQIRICR |
| *Biomphalaria glabrata* | NYEFLSLVGAGAYGQVYKARDPNKEGCFVAIKQIRIQR |
| *Bulinus truncatus* | NYEFLTLVGAGAYGQVYKARDPNKEGCFVAIKQIRIQR |
| *Elysia chlorotica* | HYEFLAHVGSGAYGQVFKARDPARENVVVAIKQIRIRR |
| *Plakobranchus ocellatus* | HYEFLAHVGSGAYGQVFKARDPTRENVVVAIKQIRIRR |
| *Gigantopelta aegis* | NYEEVAVIGNGAYGTVYKAKDLKNNGNFVAMKSIRISN |
| *Lottia gigantea* | NYEEVGVIGNGAYGTVYKGRDLKNDGKFVAIKKIRIQN |
| *Patella depressa* | NYEEVAIIGNGAYGTVYKGRDLKNEGQLVAIKKIRIPN |
| *Patella pellucida* | NYEEVAIIGNGAYGTVYKGRDLKNEGQLVAIKKIRIPN |
| *Patella vulgata* | NYEEVAIIGNGAYGTVYKGRDLKNEGQLVAIKKIRIPN |
| *Haliotis cracherodii* | KYEEVAVVGNGAYGTVYKARDLKCDGQFVAMKKIRISN |
| *Haliotis laevigata* | KYEEVAVIGNGAYGTVYKARDLKCDGQFVAMKKIRIPN |
| *Haliotis rubra* | KYEEVAVIGNGAYGTVYKARDLKCDGQFVAMKKIRIPN |
| *Haliotis rufescens* | KYEEVAVVGNGAYGTVYKARDLKCDGQFVAMKKIRISN |
| *Gibbula magus* | KYEEVAVIGNGAYGTVYKARDLKNEGQFVAMKKIRISN |
| *Phorcus lineatus* | KYEEVAVIGNGAYGTVYKARDLKNEGQFVAMKKIRISN |
| *Steromphala cineraria* | KYEEVAVIGNGAYGTVYKARDLKNEGQFVAMKKIRISN |
| *Sepia pharaonis* | EFDQLSEIGRGAYGTVYKARH-RISGQLVAMKQIPLPV |
| *Octopus sinensis* | NFDKLSEIGRGAYGTVYKARD-RKSGQYVAMKQIPLPN |
| *Nautilus pompilius* | HFEKVAVIGRGAYGTVYKARD-LKSGQMVAMKQIPLPN |
| **Nemertea** | |
| *Lineus longissimus* | NYESLTVIGHGAYGTVYKGIDKRNGEKTVALKEIKIQQ |
| *Notospermus geniculatus* | NYESLTVIGHGAYGTVYKGIDKRNGEKTVALKEIKIQQ |
| **Brachiopoda/Phoronida** | |
| *Lingula anatina* | NYEEVAVIGNGAYGTVYKARDLRNDGQMVALKKIRIQT |
| *Phoronis australis* 2 | RYEVVAEIGQGAFGAVYKGRDR-SGGGEVAIKMIEIST |
| **Bryozoa** | |
| *Bugula neritina* | NYESLNLVGTGAYGSVYRARDLRESDRFVALKKIRMKL |

### Figure S4e

**Cdk4**

| | |
|---|---|
| *Homo sapiens* | RYEPVAEIGVGAYGTVYKARD-PHSGHFVALKSVRVPN |

**Cdk4/6**

**Cnidaria**

| | |
|---|---|
| *Actinia equina* | KYEEVAEIGTGAFGTVYKARDLLNDGHFVALKRVRIIN |
| *Actinia tenebrosa* | KYEEVAEIGTGAFGTVYKARDLLNDGQFVALKRVRIIN |
| *Exaiptasia diaphana* | NYEEIAEIGTGAFGTVYKAKDLKNDGQFVALKRVRIIN |
| *Nematostella vectensis* | KYEEVAEIGTGAYGTVYKAKDLLNDGKFVALKRVRIQN |
| *Telmatactis stephensoni* | NYEEIAEIGTGAFGTVYKARDLQNDGQFVALKRVRIVN |
| *Actinostola* sp. cb2023 | NYEEIAQIGTGAFGTVYKARDLQNDGQFVALKRVRIIN |
| *Diadumene lineata* | NYEEIGEIGTGAFGTVYKARDLKNDGQFVALKRVRFIN |
| *Metridium senile* | NYEEIAQIGTGAFGTVYKARDLQNDGQFVALKRVRIIN |
| *Acropora acuminata* | KYEEVAEIGNGAYGIVYKARDLQNEGRFVALKRVRIVN |
| *Acropora awi* | KYEEVAEIGNGAYGIVYKARDLQNEGRFVALKRVRIVN |
| *Acropora cervicornis* | KYEEVAEIGNGAYGIVYKARDLQNEGRFVALKRVRIVN |
| *Acropora digitifera* | KYEEVAEIGNGAYGIVYKARDLQNEGRFVALKRVRIVN |
| *Acropora echinata* | KYEEVAEIGNGAYGIVYKARDLQNEGRFVALKRVRIVN |
| *Acropora florida* | KYEEVAEIGNGAYGIVYKARDLQNEGRFVALKRVRIVN |
| *Acropora gemmifera* | KYEEVAEIGNGAYGIVYKARDLQNEGRFVALKRVRIVN |
| *Acropora hyacinthus* | KYEEVAEIGNGAYGIVYKARDLQNEGRFVALKRVRIVN |
| *Acropora intermedia* | KYEEVAEIGNGAYGIVYKARDLQNEGRFVALKRVRIVN |
| *Acropora microphthalma* | KYEEVAEIGNGAYGIVYKARDLQNEGRFVALKRVRIVN |
| *Acropora millepora* | KYEEVAEIGNGAYGIVYKARDLQNEGRFVALKRVRIVN |
| *Acropora muricata* | KYEEVAEIGNGAYGIVYKARDLQNEGRFVALKRVRIVN |
| *Acropora nasuta* | KYEEVAEIGNGAYGIVYKARDLQNEGRFVALKRVRIVN |
| *Acropora palmata* | KYEEVAEIGNGAYGIVYKARDLQNEGRFVALKRVRIVN |
| *Acropora selago* | KYEEVAEIGNGAYGIVYKARDLQNEGRFVALKRVRIVN |
| *Acropora tenuis* | KYEEVAEIGNGAYGIVYKARDLQNEGRFVALKRVRIVN |
| *Acropora yongei* | KYEEVAEIGNGAYGIVYKARDLQNEGRFVALKRVRIVN |
| *Astreopora myriophthalma* | KYEEVAEIGNGAYGTVYKARDLQNEGRFVALKRVRIVN |
| *Montipora cactus* | NYEEVAEIGNGAYGIVYKARDLQNEGRFVALKRVRIVN |
| *Montipora efflorescens* | NYEEVAEIGNGAYGIVYKARDLQNEGRFVALKRVRIVN |
| *Pocillopora damicornis* | KYEEVAEIGNGAYGTVYKARDLQNEGRFVALKRVRIVN |
| *Pocillopora meandrina* | KYEEVAEIGNGAYGTVYKARDLQNEGRFVALKRVRIVN |
| *Pocillopora verrucosa* | KYEEVAEIGNGAYGTVYKARDLQNEGRFVALKRVRIVN |
| *Stylophora pistillata* | KYEEVAEIGNGAYGTVYKARDLQNEGRFVALKRVRIVN |
| *Catalaphyllia jardinei* | KYEEVAEIGNGAYGTVYKARDLQNEGRFVALKRVRIVN |
| *Cladopsammia gracilis* | KYEEVAEIGNGAYGTVYKARDLQNEGRFVALKRVRIVN |
| *Dendrophyllia cribrosa* | KYEEVAEIGNGAYGTVYKARDLQNEGRFVALKRVRIVN |
| *Galaxea fascicularis* | KYQEIAEIGNGAYGTVYKARDLQNEGRFVALKRVRIVN |
| *Blastomussa wellsi* | KYEEVAEIGNGAYGTVYKARDLQNEGRFVALKRVRIVN |
| *Orbicella faveolata* | KYEEVAEIGNGAYGTVYKARDLQNEGRFVALKRVRIVN |
| *Pachyseris speciosa* | KYEEVAEIGNGAYGTVYKARDLQNEGRFVALKRVRIVN |
| *Porites australiensis* | KYEEVAMIGNGAYGTVYKARDLQNEGRFVALKRVRIIV |
| *Porites lobata* | KYEEVAMIGNGAYGTVYKARDLQNEGRFVALKRVRIIV |
| *Porites lutea* | KYEEVAMIGNGAYGTVYKARDLQNEGRFVALKRVRIIV |
| *Dendronephthya gigantea* | KYEEVGEIGTGAYGTVYKAKDLLNNNRYVALKRVRITN |
| *Paramuricea clavata* | KYEEVGEIGTGAYGTVYKARDLLNNNRYVALKRVRITN |
| *Xenia* sp. Carnegie-2017 | KYEEVGEIGTGAYGTVYKARDLLNNNRYVALKRTLITN |
| *Heliopora coerulea* | KYEEVGEIGTGAYGTVYKARDLEHNGRFVALKRVRILN |
| *Morbakka virulenta* | NYEQQALIGRGAYGTVFRAVDHLNN-RIVALKRVRLKQ |
| *Hydra viridissima* | KYEELSIIGRGAYGTVFKARDIEND-RVVALKRVRIGQ |
| *Hydra vulgaris* | EFEELSIIGRGAYGTVFKARDVEHN-RVVALKRVCISQ |
| *Hydractinia echinata* | NYEDLSIIGKGAYGTVFKAFDTEND-RIVALKRVRIIN |
| *Hydractinia symbiolongicarpus* | NYEDLSIIGKGAYGTVFKAFDAEND-RIVALKRVRIIN |
| *Turritopsis rubra* | NYEELSIIGKGAYGTVFKAIDHEND-RTVALKRVRILS |
| *Clytia hemisphaerica* | NYEELAIIGRGAYGTVFKARDVEND-RTVALKRVRIVN |
| *Nemopilema nomurai* | NYEELDLIGRGAYGTVFRARDHLND-RMVALKRVRIPT |
| *Chrysaora quinquecirrha* | KYKELGLIGRGAYGAVFKARNNVT-GSMVALKRIKILA |
| *Aurelia aurita* | NYEELGLIGRGAYGSVFRARDHLN-GRMVALKRVKIAT |

**Ctenophora**

| | |
|---|---|
| *Mnemiopsis leidyi* | NYKLLEKIGCGTFGVVYRARHEPTQ-EIVAVKQARIQQ |

**Placozoa**

| | |
|---|---|
| *Trichoplax adhaerens* | NYVQVTEIANGAYGRVYKARDL-NHNRLVALKKIAVIN |

### Figure S4f

**Figure S4** Cdk4/6 amino acid alignment around the residue corresponding to R24/R31 in invertebrates. (a) deuterostome invertebrates, (b) Nematoda, (c) other Ecdysozoa, (d) Platyhelminthes, (e) other Lophotrochozoa, and (f) basal invertebrates. The residue corresponding to R24/31 in Cdk4/6 is in green if positively charged and in magenta, if not.
