## Supplementary Tables S1-S11 and Figures S1-S6 for "Evolution of the *Cdk4/6*–*Cdkn2* system in invertebrates": Figure S5.pdf

**Cdk4**
**mammal**
**<placentarian>**

| | |
|---|---|
| *Homo sapiens* | RYEPVAEIGVGAYGTVYKARDPHSGHFVALKSVRVPN |
| *Acinonyx jubatus* | RYEPVAEIGVGAYGTVYKARDPHSGHFVALKSVRVPN |
| *Ailuropoda melanoleuca* | RYEPVAEIGVGAYGTVYKARDPHSGHFVALKSVRVPN |
| *Aotus nancymaae* | RYEPVAEIGVGAYGTVYKARDPHSGHFVALKSVRVPN |
| *Artibeus jamaicensis* | RYEPVAEIGVGAYGTVYKARDPNSGHFVALKSVRVPN |
| *Arvicanthis niloticus* | RYEPVAEIGVGAYGTVYKARDPHSGHFVALKSVRVPN |
| *Arvicola amphibius* | RYEPVAEIGVGAYGTVYKARDPHSGHFVALKSVRVPN |
| *Balaenoptera acutorostrata scammoni* | RYEPVAEIGVGAYGTVYKARDPHSGHFVALKSVRVPN |
| *Balaenoptera musculus* | RYEPVAEIGVGAYGTVYKARDPHSGHFVALKSVRVPN |
| *Bison bison bison* | RYEPVAEIGVGAYGTVYKARDPHSGHFVALKSVRVPN |
| *Bos indicus* | -YEPVAEIGVGAYGTVYKARDPHSGHFVALKSVRVPN |
| *Bos mutus* | RYEPVAEIGVGAYGTVYKARDPHSGHFVALKSVRVPN |
| *Bos taurus* | RYEPVAEIGVGAYGTVYKARDPHSGHFVALKSVRVPN |
| *Bubalus bubalis* | RYEPVAEIGVGAYGTVYKARDPHSGHFVALKSVRVPN |
| *Callithrix jacchus* | RYEPVAEIGVGAYGTVYKARDPHSGHFVALKSVRVPN |
| *Callorhinus ursinus* | RYEPVAEIGVGAYGTVYKARDPHSGHFVALKSVRVPN |
| *Camelus bactrianus* | RYEPVAEIGVGAYGTVYKARDPHSGHFVALKSVRVPN |
| *Camelus dromedarius* | RYEPVAEIGVGAYGTVYKARDPHSGHFVALKSVRVPN |
| *Camelus ferus* | RYEPVAEIGVGAYGTVYKARDPHSGHFVALKSVRVPN |
| *Canis lupus familiaris* | RYEPVAEIGVGAYGTVYKARDPHSGHFVALKSVRVPN |
| *Capra hircus* | RYEPVAEIGVGAYGTVYKARDPHSGHFVALKSVRVPN |
| *Carlito syrichta* | RYEPVAEIGVGAYGTVYKARDPHSGHFVALKSVRVPN |
| *Castor canadensis* | RYEPVAEIGVGAYGTVYKARDPHSGHFVALKSVRVPN |
| *Cavia porcellus* | RYEPVAEIGVGAYGTVYKARDPHSGHFVALKSVRVPN |
| *Cebus imitator* | RYEPVAEIGVGAYGTVYKARDPHSGHFVALKSVRVPN |
| *Ceratotherium simum simum* | RYEPVAEIGVGAYGTVYKARDPHSGHFVALKSVRVPN |
| *Cercocebus atys* | RYEPVAEIGVGAYGTVYKARDPHSGHFVALKSVRVPN |
| *Cervus canadensis* | RYEPVAEIGVGAYGTVYKARDPHSGHFVALKSVRVPN |
| *Cervus elaphus* | RYEPVAEIGVGAYGTVYKARDPHSGHFVALKSVRVPN |
| *Cervus hanglu yarkandensis* | RYEPVAEIGVGAYGTVYKARDPHSGHFVALKSVRVPN |
| *Chinchilla lanigera* | RYEPVAEIGVGAYGTVYKARDPHSGHFVALKSVRVPN |
| *Chlorocebus sabaeus* | RYEPVAEIGVGAYGTVYKARDPHSGHFVALKSVRVPN |
| *Choloepus didactylus* | RYEPVAEIGVGAYGTVYKARDPHSGHFVALKSVRVPN |
| *Chrysochloris asiatica* | RYEPVAEIGVGAYGTVYKARDPHSGHFVALKSVRVPN |
| *Colobus angolensis palliatus* | RYEPVAEIGVGAYGTVYKARDPHSGHFVALKSVRVPN |
| *Condylura cristata* | RYEPVAEIGVGAYGTVYKARDPHSGHFVALKSVRVPN |
| *Cricetulus griseus* | RYEPVAEIGVGAYGTVYKARDPHSGHFVALKSVRVPN |
| *Crocuta crocuta* | RYEPVAEIGVGAYGTVYKARDPHSGHFVALKSVRVPN |
| *Dasypus novemcinctus* | RYEPVAEIGVGAYGTVYKARDPHSGHFVALKSVRVPN |
| *Delphinapterus leucas* | RYEPVAEIGVGAYGTVYKARDPHSGHFVALKSVRVPN |
| *Desmodus rotundus* | RYEPVAEIGVGAYGTVYKARDPHSGHFVALKSVRVPN |
| *Dipodomys ordii* | RYEPVAEIGVGAYGTVYKARDPHSGHFVALKSVRVPN |
| *Dipodomys spectabilis* | RYEPVAEIGVGAYGTVYKARDPHSGHFVALKSVRVPN |
| *Echinops telfairi* | RYEPVAEIGVGAYGTVYKARDPHSGHFVALKSVRVPN |
| *Elephantulus edwardii* | RYEPVAEIGVGAYGTVYKARDPNSGHFVALKSVRVPN |
| *Enhydra lutris kenyoni* | RYEPVAEIGVGAYGTVYKARDPHSGHFVALKSVRVPN |
| *Eptesicus fuscus* | RYEPVAEIGVGAYGTVYKARDPHSGHFVALKSVRVPN |
| *Equus asinus* | RYEPVAEIGVGAYGTVYKARDPHSGHFVALKSVRVPN |
| *Equus caballus* | RYEPVAEIGVGAYGTVYKARDPHSGHFVALKSVRVPN |
| *Equus przewalskii* | RYEPVAEIGVGAYGTVYKARDPHSGHFVALKSVRVPN |
| *Equus quagga* | RYEPVAEIGVGAYGTVYKARDPHSGHFVALKSVRVPN |
| *Erinaceus europaeus* | RYEPVAEIGVGAYGTVYKARDPHSGHFVALKSVRVPN |
| *Eumetopias jubatus* | RYEPVAEIGVGAYGTVYKARDPHSGHFVALKSVRVPN |
| *Felis catus* | RYEPVAEIGVGAYGTVYKARDPHSGHFVALKSVRVPN |
| *Fukomys damarensis* | RYEPVAEIGVGAYGTVYKARDPHSGHFVALKSVRVPN |
| *Galemys pyrenaicus* | RYEPVAEIGVGAYGTVYKARDPHSGHFVALKSVRVPN |
| *Galeopterus variegatus* | RYEPVAEIGVGAYGTVYKARDPHSGHFVALKSVRVPN |
| *Globicephala melas* | RYEPVAEIGVGAYGTVYKARDPHSGHFVALKSVRVPN |
| *Gorilla gorilla gorilla* | RYEPVAEIGVGAYGTVYKARDPHSGHFVALKSVRVPN |
| *Grammomys surdaster* | RYEPVAEIGVGAYGTVYKARDPHSGHFVALKSVRVPN |
| *Gulo gulo luscus* | RYEPVAEIGVGAYGTVYKARDPHSGHFVALKSVRVPN |
| *Halichoerus grypus* | RYEPVAEIGVGAYGTVYKARDPHSGHFVALKSVRVPN |
| *Heterocephalus glaber* | RYEPVAEIGVGAYGTVYKARDPHSGHFVALKSVRVPS |
| *Hipposideros armiger* | RYVPVAEIGVGAYGTVYKARDPHSGHFVALKSVRVPN |
| *Hyaena hyaena* | RYEPVAEIGVGAYGTVYKARDPHSGHFVALKSVRVPN |
| *Hylobates moloch* | RYEPVAEIGVGAYGTVYKARDPHSGHFVALKSVRVPN |
| *Ictidomys tridecemlineatus* | RYEPVAEIGVGAYGTVYKARDPHSGHFVALKSVRVPN |
| *Jaculus jaculus* | RYEPVAEIGVGAYGTVYKARDPHSGHFVALKSVRVPN |
| *Lagenorhynchus obliquidens* | RYEPVAEIGVGAYGTVYKARDPHSGHFVALKSVRVPN |
| *Lemur catta* | RYEPVAEIGVGAYGTVYKARDPHSGHFVALKSVRVPN |
| *Leopardus geoffroyi* | RYEPVAEIGVGAYGTVYKARDPHSGHFVALKSVRVPN |
| *Leptonychotes weddellii* | RYEPVAEIGVGAYGTVYKARDPHSGHFVALKSVRVPN |
| *Lipotes vexillifer* | RYEPVAEIGVGAYGTVYKARDPHSGHFVALKSVRVPN |
| *Lontra canadensis* | RYEPVAEIGVGAYGTVYKARDPHSGHFVALKSVRVPN |
| *Loxodonta africana* | RYEPVAEIGVGAYGTVYKARDPHSGHFVALKSVRVPN |
| *Lynx canadensis* | RYEPVAEIGVGAYGTVYKARDPHSGHFVALKSVRVPN |
| *Lynx pardinus* | RYEPVAEIGVGAYGTVYKARDPHSGHFVALKSVRVPN |
| *Macaca fascicularis* | RYEPVAEIGVGAYGTVYKARDPHSGHFVALKSVRVPN |
| *Macaca mulatta* | RYEPVAEIGVGAYGTVYKARDPHSGHFVALKSVRVPN |
| *Macaca nemestrina* | RYEPVAEIGVGAYGTVYKARDPHSGHFVALKSVRVPN |
| *Mandrillus leucophaeus* | RYEPVAEIGVGAYGTVYKARDPHSGHFVALKSVRVPN |
| *Manis javanica* | RYEPVAEIGVGAYGTVYKARDPHSGHFVALKSVRVPN |
| *Manis pentadactyla* | RYEPVAEIGVGAYGTVYKARDPHSGHFVALKSVRVPN |
| *Marmota flaviventris* | RYEPVAEIGVGAYGTVYKARDPHSGHFVALKSVRVPN |
| *Marmota marmota marmota* | RYEPVAEIGVGAYGTVYKARDPHSGHFVALKSVRVPN |
| *Marmota monax* 1 | RYEPVAEIGVGAYGTVYKARDPHSGHFVALKSVRVPN |
| *Marmota monax* 2 | RYEPVAEIGVGAYGTVYKARDPHSGHFVALKSVRVPN |
| *Mastomys coucha* | RYEPVAEIGVGAYGTVYKARDPHSGHFVALKSVRVPN |
| *Meles meles* | RYEPVAEIGVGAYGTVYKARDPHSGHFVALKSVRVPN |
| *Meriones unguiculatus* | RYEPVAEIGVGAYGTVYKARDPHSGHFVALKSVRVPN |
| *Mesocricetus auratus* | RYEPVAEIGVGAYGTVYKARDPHSGHFVALKSVRVPN |
| *Microcebus murinus* | RYEPVAEIGVGAYGTVYKARDPHSGHFVALKSVRVPN |
| *Microtus ochrogaster* | RYEPVAEIGVGAYGTVYKARDPHSGHFVALKSVRVPN |
| *Microtus oregoni* | RYEPVAEIGVGAYGTVYKARDPHSGHFVALKSVRVPN |
| *Miniopterus natalensis* | RYEPVAEIGVGAYGTVYKARDPHSGHFVALKSVRVPN |
| *Mirounga angustirostris* | RYEPVAEIGVGAYGTVYKARDPHSGHFVALKSVRVPN |
| *Mirounga leonina* | RYEPVAEIGVGAYGTVYKARDPHSGHFVALKSVRVPN |
| *Molossus molossus* | RYEPVAEIGVGAYGTVYKARDPHSGHFVALKSVRVPN |
| *Monodon monoceros* | RYEPVAEIGVGAYGTVYKARDPHSGHFVALKSVRVPN |
| *Mus caroli* | RYEPVAEIGVGAYGTVYKARDPHSGHFVALKSVRVPN |
| *Mus musculus* | RYEPVAEIGVGAYGTVYKARDPHSGHFVALKSVRVPN |
| *Mus pahari* | RYEPVAEIGVGAYGTVYKARDPHSGHFVALKSVRVPN |
| *Mustela erminea* | RYEPVAEIGVGAYGTVYKARDPHSGHFVALKSVRVPN |
| *Mustela putorius furo* | RYEPVAEIGVGAYGTVYKARDPHSGHFVALKSVRVPN |
| *Myotis brandtii* | RYEPVAEIGVGAYGTVYKARDPHSGHFVALKSVRVPN |
| *Myotis davidii* | RYEPVAEIGVGAYGTVYKARDPHSGHFVALKSVRVPN |
| *Myotis lucifugus* | RYEPVAEIGVGAYGTVYKARDPHSGHFVALKSVRVPN |
| *Myotis myotis* | RYEPVAEIGVGAYGTVYKARDPHSGHFVALKSVRVPN |
| *Nannospalax galili* | RYEPVAEIGVGAYGTVYKARDPHSGHFVALKSVRVPN |
| *Neogale vison* | RYEPVAEIGVGAYGTVYKARDPHSGHFVALKSVRVPN |
| *Neomonachus schauinslandi* | RYEPVAEIGVGAYGTVYKARDPHSGHFVALKSVRVPN |
| *Neophocaena asiaeorientalis asiaeorientalis* | RYEPVAEIGIGAYGTVYKARDPHSGHFVALKSVRVPN |
| *Neosciurus carolinensis* | RYEPVAEIGVGAYGTVYKARDPHSGHFVALKSVRVPN |
| *Neotoma lepida* | RYEPVAEIGVGAYGTVYKARDPHSGHFVALKSVRVPN |
| *Nomascus leucogenys* | RYEPVAEIGVGAYGTVYKARDPHSGHFVALKSVRVPN |
| *Nyctereutes procyonoides* | RYEPVAEIGVGAYGTVYKARDPHSGHFVALKSVRVPN |
| *Ochotona curzoniae* | RYEPVAEIGVGAYGTVYKARDPLSGHFVALKSLRVPS |
| *Ochotona princeps* | RYEPVAEIGVGAYGTVYKARDPLSGHFVALKSLRVPS |
| *Octodon degus* | RYEPVAEIGVGAYGTVYKARDPHSGHFVALKSVRVPN |
| *Odobenus rosmarus divergens* | RYEPVAEIGVGAYGTVYKARDPHSGHFVALKSVRVPN |
| *Odocoileus virginianus texanus* | RYEPVAEIGVGAYGTVYKARDPHSGHFVALKSVRVPN |
| *Onychomys torridus* | RYEPVAEIGVGAYGTVYKARDPHSGHFVALKSVRVPN |
| *Orcinus orca* | RYEPVAEIGVGAYGTVYKARDPHSGHFVALKSVRVPN |
| *Orycteropus afer afer* | RYEPVAEIGVGAYGTVYKARDPHSGHFVALKSVRVPN |
| *Oryctolagus cuniculus* | RYEPVAEIGVGAYGTVYKARDPHSGHFVALKSVRVPN |
| *Oryx dammah* | RYEPVAEIGVGAYGTVYKARDPHSGHFVALKSVRVPN |
| *Otolemur garnettii* | RYEPVAEIGVGAYGTVYKARDPHSGHFVALKSVRVPN |
| *Ovis aries* | RYEPVAEIGVGAYGTVYKARDPHSGHFVALKSVRVPN |
| *Pan paniscus* | RYEPVAEIGVGAYGTVYKARDPHSGHFVALKSVRVPN |
| *Pan troglodytes* | RYEPVAEIGVGAYGTVYKARDPHSGHFVALKSVRVPN |
| *Panthera leo* | RYEPVAEIGVGAYGTVYKARDPHSGHFVALKSVRVPN |
| *Panthera pardus* | RYEPVAEIGVGAYGTVYKARDPHSGHFVALKSVRVPN |
| *Panthera tigris* | RYEPVAEIGVGAYGTVYKARDPHSGHFVALKSVRVPN |
| *Papio anubis* | RYEPVAEIGVGAYGTVYKARDPHSGHFVALKSVRVPN |
| *Perognathus longimembris pacificus* | RYEPVAEIGVGAYGTVYKARDPHSGHFVALKSVRVPN |
| *Peromyscus leucopus* | RYEPVAEIGVGAYGTVYKARDPHSGHFVALKSVRVPN |
| *Peromyscus maniculatus bairdii* | RYEPVAEIGVGAYGTVYKARDPHSGHFVALKSVRVPN |
| *Phoca vitulina* | RYEPVAEIGVGAYGTVYKARDPHSGHFVALKSVRVPN |
| *Phocoena sinus* | RYEPVAEIGVGAYGTVYKARDPHSGHFVALKSVRVPN |
| *Phyllostomus discolor* | RYEPVAEIGVGAYGTVYKARDPNSGHFVALKSVRVPN |
| *Phyllostomus hastatus* | RYEPVAEIGVGAYGTVYKARDPNSGHFVALKSVRVPN |
| *Physeter catodon* | RYEPVAEIGVGAYGTVYKARDPHSGHFVALKSVRVPN |
| *Piliocolobus tephrosceles* | RYEPVAEIGVGAYGTVYKARDPHSGHFVALKSVRVPN |
| *Pipistrellus kuhlii* | RYEPVAEIGVGAYGTVYKARDPHSGHFVALKSVRVPN |
| *Pongo abelii* | RYEPVAEIGVGAYGTVYKARDPHSGHFVALKSVRVPN |
| *Prionailurus bengalensis* | RYEPVAEIGVGAYGTVYKARDPHSGHFVALKSVRVPN |
| *Propithecus coquereli* | RYEPVAEIGVGAYGTVYKARDPHSGHFVALKSVRVPN |
| *Pteropus alecto* | RYEPVAEIGVGAYGTVYKARDPHSGHFVALKSVRVPN |
| *Pteropus giganteus* | RYEPVAEIGVGAYGTVYKARDPHSGHFVALKSVRVPN |
| *Pteropus vampyrus* | RYEPVAEIGVGAYGTVYKARDPHSGHFVALKSVRVPN |
| *Puma concolor* | RYEPVAEIGVGAYGTVYKARDPHSGHFVALKSVRVPN |
| *Puma yagouaroundi* | RYEPVAEIGVGAYGTVYKARDPHSGHFVALKSVRVPN |
| *Rattus norvegicus* | RYEPVAEIGVGAYGTVYKARDPHSGHFVALKSVRVPN |
| *Rattus rattus* | RYEPVAEIGVGAYGTVYKARDPHSGHFVALKSVRVPN |
| *Rhinolophus ferrumequinum* | RYVPVAEIGVGAYGTVYKARDPHSGHFVALKSVRVPN |
| *Rhinolophus sinicus* | RYVPVAEIGVGAYGTVYKARDPHSGHFVALKSVRVPN |
| *Rhinopithecus bieti* | RYEPVAEIGVGAYGTVYKARDPHSGHFVALKSVRVPN |
| *Rhinopithecus roxellana* | RYEPVAEIGVGAYGTVYKARDPHSGHFVALKSVRVPN |
| *Rousettus aegyptiacus* | RYEPVAEIGVGAYGTVYKARDPHSGHFVALKSVRVPN |
| *Saimiri boliviensis boliviensis* | RYEPVAEIGVGAYGTVYKARDPHSGHFVALKSVRVPN |
| *Sapajus apella* | RYEPVAEIGVGAYGTVYKARDPHSGHFVALKSVRVPN |
| *Sorex araneus* | RYEPVAEIGVGAYGTVYKARDPHSGHFVALKSVRVPN |
| *Sousa chinensis* | RYEPVAEIGVGAYGTVYKARDPHSGHFVALKSVRVPN |
| *Sturnira hondurensis* | RYEPVAEIGVGAYGTVYKARDPNSGHFVALKSVRVPN |
| *Suricata suricatta* | RYEPVAEIGVGAYGTVYKARDPHSGHFVALKSVRVPN |
| *Sus scrofa* | RYEPVAEIGVGAYGTVYKARDPHSGHFVALKSVRVPN |
| *Talpa occidentalis* | RYEPVAEIGVGAYGTVYKARDPHSGHFVALKSVRVPN |
| *Theropithecus gelada* | RYEPVAEIGVGAYGTVYKARDPHSGHFVALKSVRVPN |
| *Trachypithecus francoisi* | RYEPVAEIGVGAYGTVYKARDPHSGHFVALKSVRVPN |
| *Trichechus manatus latirostris* | RYEPVAEIGVGAYGTVYKARDPHSGHFVALKSVRVPN |
| *Tupaia chinensis* | RYEPVAEIGVGAYGTVYKARDPHSGHFVALKSVRVPN |
| *Tursiops truncatus* | RYEPVAEIGVGAYGTVYKARDPHSGHFVALKSVRVPN |
| *Urocitellus parryii* | RYEPVAEIGVGAYGTVYKAHDPHSGHFVALKSVRVPN |
| *Ursus americanus* | RYEPVAEIGVGAYGTVYKARDPHSGHFVALKSVRVPN |
| *Ursus arctos* | RYEPVAEIGVGAYGTVYKARDPHSGHFVALKSVRVPN |
| *Ursus maritimus* | RYEPVAEIGVGAYGTVYKARDPHSGHFVALKSVRVPN |
| *Vicugna pacos* | RYEPVAEIGVGAYGTVYKARDPHSGHFVALKSVRVPN |
| *Vulpes lagopus* | RYEPVAEIGVGAYGTVYKARDPHSGHFVALKSVRVPN |
| *Vulpes vulpes* | RYEPVAEIGVGAYGTVYKARDPHSGHFVALKSVRVPN |
| *Zalophus californianus* | RYEPVAEIGVGAYGTVYKARDPHSGHFVALKSVRVPN |
| **<marsupial>** | |
| *Dromiciops gliroides* | RYEPVAEIGVGAYGTVYKARDPHSGHFVALKSVKVPT |
| *Gracilinanus agilis* | RYEPVAEIGVGAYGTVYKARDPHSGHFVALKSVKVPT |
| *Notamacropus eugenii* | RYEPVAEIGVGAYGTVYKARDPHSGHFVALKSVKVPT |
| *Phascolarctos cinereus* | RYEPVAEIGVGAYGTVYKARDPHSGHFVALKSVKVPT |
| *Sarcophilus harrisii* | RYEPVAEIGVGAYGTVYKARDPHSGHFVALKSVKVPT |
| *Trichosurus vulpecula* 1 | RYEPVAEIGVGAYGTVYKARDPHSGHFVALKSVKVPT |
| *Trichosurus vulpecula* 2 | RYEPVAEIGVGAYGTVYEVRDPHSGHFVALKSVKVPT |
| *Vombatus ursinus* | RYEPVAEIGVGAYGTVYKARDPHSGHFVALKSVKVPT |

**Figure S5a**

| | |
|---|---|
| **Cdk4** | |
| *Homo sapiens* | RYEPVAEIGVGAYGTVYKARDPHSGHFVALKSVRVPN |
| **bird** | |
| *Accipiter nisus* | QYEPVAEIGVGAYGTVYKARDMQSGKFVALKNVRVQT |
| *Anas zonorhyncha* | -YEPVAEIGVGAYGTVYKARDLRSGRFVALKNVRVTA |
| *Anser cygnoides* | QYEPVAEIGVGAYGTVYKARDLQSGRFVALKNVRVQA |
| *Apteryx mantelli mantelli* | QYEPVAEIGVGAYGTVYKARDLQSGKFVALKNVRVQT |
| *Apteryx rowi* | QYEPVAEIGVGAYGTVYKARDLQSGKFVALKNVRVQT |
| *Balearica regulorum gibbericeps* | QYEPVAEIGVGAYGTVYKARDLQSGKFVALKNVRVQM |
| *Calidris pugnax* | QYEPVAEIGVGAYGTVYKARDRASGRFVALKNVRVTG |
| *Calidris pygmaea* 1 | -YEPVAEIGVGAYGTVYKARDRASGRFVALKNVRVTG |
| *Calidris pygmaea* 2 | -YEPVAEIGVGAYGTVYKAWDRASGRFVALKKVRMMG |
| *Cariama cristata* | QYEPVAEIGVGAYGTVYKARDLQSGKFVALKNVRVQT |
| *Catharus ustulatus* | QYEPVAEPGVGAHGTAFKARHRLSGRCVALKNLRVPT |
| *Corvus hawaiiensis* | QYEPVAELGVGAHGAVFKARDRLSGRFVALKNVRVPT |
| *Corvus moneduloides* | QYEPVAELGVGAHGAVFKARDRLSGRFVALKNVRVPT |
| *Coturnix japonica* | RYEPVSTLGSGSSCSVFRARDRRNGRLVALKRVRVGS |
| *Dromaius novaehollandiae* | QYEPVAEIGVGAYGTVYKARDLQSGKFVALKNVRVQT |
| *Eudyptes chrysocome* | QYEPVAEIGVGAYGTVYKARDLQSGKFVALKNVRVQT |
| *Eudyptes filholi* | QYEPVAEIGVGAYGTVYKARDLQSGKFVALKNVRVQT |
| *Eudyptes moseleyi* | QYEPVAEIGVGAYGTVYKARDLQSGKFVALKNVRVQT |
| *Eudyptes pachyrhynchus* | QYEPVAEIGVGAYGTVYKARDLQSGKFVALKNVRVQT |
| *Eudyptes robustus* | QYEPVAEIGVGAYGTVYKARDLQSGKFVALKNVRVQT |
| *Eudyptes schlegeli* | QYEPVAEIGVGAYGTVYKARDLQSGKFVALKNVRVQT |
| *Eudyptes sclateri* | QYEPVAEIGVGAYGTVYKARDLQSGKFVALKNVRVQT |
| *Eudyptula minor* | QYEPVAEIGVGAYGTVYKARDLQSGKFVALKNVRVQT |
| *Falco naumanni* | QYEPVAEIGVGAYGTVYKARDRRSGRFVALKNVRVQA |
| *Falco peregrinus* | QYEPVAEIGVGAYGTVYKARDRRSGRFVALKNVRVQA |
| *Falco tinnunculus* | QYEPVAEIGVGAYGTVYKARDRRSGRFVALKNVRVQA |
| *Haliaeetus albicilla* | QYEPVAEIGVGAYGTVYKARDMQSGKFVALKNVRVQT |
| *Hypotaenidia okinawae* | QYEPVAEIGVGAYGTVYKARDLRSGKFVALKNVRVRL |
| *Merops nubicus* | QYEPVAEIGVGAYGTVYKARDLQSGKFVALKNVRVQT |
| *Nothoprocta perdicaria* | QYEPVAEIGVGAYGTVYKARDLQSGKFVALKNVRVQT |
| *Otus sunia* | QYEPVAEIGGGAYGTVYKARDRRSGRFVALKNVRVQT |
| *Parus major* | QYEPVAELGVGAHGSVFKARDRLSGRFVALKNVRVPT |
| *Pseudopodoces humilis* | QYEPVAELGVGAHGSVFKARDRLSGRFVALKNVRVPT |
| *Spheniscus humboldti* | QYEPVAEIGVGAYGTVYKARDLQSGKFVALKNVRVQT |
| *Strix occidentalis caurina* | QYEPVAEIGVGAYGTVYKARDRQSGKFVALKNVRVQT |
| *Sturnus vulgaris* | QYEPVAELGVGAHGTVFKARDRLSGRFVALKNVRVPT |
| *Tyto alba* | QYEPVAEIGVGAYGTVYKARDRQSGKFVALKNVRVQT |
| *Zosterops borbonicus* | QYEPVAELGVGAHGTVFKARDRLSGRFVALKNVRVPT |
| **reptile** | |
| *Alligator mississippiensis* | QYEPVAEIGVGAYGTVYKARDLQSGRFVALKNVRVPS |
| *Alligator sinensis* | --DPVAEIGVGAYGTVYKARDLQSGRFVALKNVRVPS |
| *Anolis carolinensis* | QYEPVAEIGIGAYGTVFKARDLQSGKFVALKNVRFQN |
| *Chelonia mydas* | QYEPVAEIGVGAYGTVYKARDLQSGKFVALKNVRVQT |
| *Chelonoidis abingdonii* | QYEPVAEIGVGAYGTVYKARDLQSGKFVALKNVRVQT |
| *Chelydra serpentina* | QYEPVAEIGVGAYGTVYKARDLQSGKFVALKNVRVQT |
| *Chrysemys picta bellii* | QYEPVAEIGVGAYGTVYKARDLQTGKFVALKNVRVQT |
| *Crocodylus porosus* | QYEPVAEIGVGAYGTVYKARDLQSGRFVALKNVRVPS |
| *Crotalus adamanteus* | QYEPVAEIGIGAYGTVFKARDLQSGKFVALKNVRVQN |
| *Crotalus tigris* | QYEPVAEIGIGAYGTVFKARDLQSGKFVALKNVRVQN |
| *Dermochelys coriacea* | QYEPVAEIGVGAYGTVYKARDLQSGKFVALKNVRVQT |
| *Gavialis gangeticus* | QYEPVAEIGVGAYGTVYKARDLQSGRFVALKNVRVPS |
| *Gekko japonicus* | QYEPVAEIGIGAYGTVFKARDLQSGKFVALKNVRVQS |
| *Gopherus agassizii* | QYEPVAEIGVGAYGTVYKARDLQSGKFVALKNVRVQT |
| *Gopherus evgoodei* | QYEPVAEIGVGAYGTVYKARDLQSGKFVALKNVRVQT |
| *Lacerta agilis* | QYEPVAEIGIGAYGTVFKARDLLSGKFVALKNVRVQN |
| *Laticauda laticaudata* | QYEPVAEIGIGAYGTVFKARDLQSGKFVALKNVRVQN |
| *Mauremys mutica* | QYEPVAEIGVGAYGTVYKARDLQSGKFVALKNVRVQT |
| *Mauremys reevesii* | QYEPVAEIGVGAYGTVYKARDLQSGKFVALKNVRVQT |
| *Naja naja* | QYEPVAEIGIGAYGTVFKARDLQSGKFVALKNVRVQN |
| *Notechis scutatus* | QYEPVAEIGIGAYGTVFKARDLQSGKFVALKNVRVQN |
| *Ophiophagus hannah* | QYEPVAEIGIGAYGTVFKARDLQSGKFVALKNVRVQN |
| *Pantherophis guttatus* | QYEPVAEIGIGAYGTVFKARDLQSGKFVALKNVRVQN |
| *Pelusios castaneus* | QYEPVAEIGVGAYGTVYKARDLQSGKFVALKNVRVQT |
| *Podarcis muralis* | QYEPVAEIGIGAYGTVFKARDLQSGKFVALKNVRVQN |
| *Pogona vitticeps* | QYEPVAEIGIGAYGTVFKARDLQSGKFVALKNVRVQN |
| *Protobothrops mucrosquamatus* | QYEPVAEIGIGAYGTVFKARDLQSGKFVALKNVRVQN |
| *Pseudonaja textilis* | QYEPVAEIGIGAYGTVFKARDLQSGKFVALKNVRVQN |
| *Python bivittatus* | QYEPVAEIGIGAYGTVFKARDLQSGKFVALKNVRVQN |
| *Salvator merianae* | QYEPVAEIGIGAYGTVFKARDLQSGKFVALKNVRVQN |
| *Sceloporus undulatus* | QYEPVAEIGIGAYGTVFKARDLQSGKFVALKNVRVQN |
| *Sphenodon punctatus* | QYVPVAEIGVGAYGTVYKARDLQSGKFVALKNVRVQS |
| *Terrapene carolina triunguis* | QYEPVAEIGVGAYGTVYKARDLQSGKFVALKNVRVQT |
| *Thamnophis elegans* | QYEPVAEIGIGAYGTVFKARDLQSGKFVALKNVRVQN |
| *Thamnophis sirtalis* | QYEPVAEIGIGAYGTVFKARDLQSGKFVALKNVRVQN |
| *Trachemys scripta elegans* | QYEPVAEIGVGAYGTVYKARDLQTGKFVALKNVRVQT |
| *Varanus komodoensis* | QYEPVAEIGIGAYGTVFKARDLQSGKFVALKNVRVQN |
| *Zootoca vivipara* | QYEPVAEIGIGAYGTVFKARDLQSGKFVALKNVRVQN |
| **amphibian** | |
| *Ambystoma andersoni* | KYEPVAEIGVGAYGVVYKARDLQSGKFVALKNVRVQT |
| *Ambystoma maculatum* | KYEPVAEIGVGAYGVVYKARDLQSGKFVALKNVRVQT |
| *Ambystoma mexicanum* | KYEPVAEIGVGAYGVVYKARDLQSGKFVALKNVRVQT |
| *Ambystoma velasci* | KYEPVAEIGVGAYGVVYKARDLQSGKFVALKNVRVQT |
| *Bufo bufo* | QYEPVAEIGVGAYGTVYKARDLQSGKFVALKNVRVQT |
| *Bufo gargarizans* | QYEPVAEIGVGAYGTVYKARDLQSGKFVALKNVRVQT |
| *Engystomops pustulosus* | QYEPVAEIGVGAYGTVYKARDLQSGKFVALKNVRVQT |
| *Geotrypetes seraphini* | QYELVAEIGVGAYGTVYKARDMQSGKFVALKNVRVQT |
| *Hymenochirus boettgeri* | QYEPVAEIGVGAYGTVYKARDLQSGKFVALKNVRVQT |
| *Leptobrachium leishanense* | QYEPVAEIGVGAYGTVYKARDLQSGKFVALKNVRVQT |
| *Microcaecilia unicolor* | QYEPVAEIGVGAYGTVYKARDLQSGKFVALKNVRVQT |
| *Nanorana parkeri* | QYEPVAEIGVGAYGTVYKARDLQSGKFVALKNVRVQT |
| *Rana temporaria* | QYEPVAEIGVGAYGTVYKARDLRSGKFVALKNVRVQT |
| *Rhinatrema bivittatum* 1 | QYEPVAEIGVGAYGTVYKARDLQSGKFVALKNVRVQT |
| *Rhinatrema bivittatum* 2 | QYEPVAEIGVGAYGTVYKARDLQSGKFVALKNVRVQT |
| *Xenopus laevis* 1 | QYEPVAEIGVGAYGTVYKARDLQSGKFVALKNVRVQT |
| *Xenopus laevis* 2 | QYEPVAEIGVGAYGTVYKARDLHSGKFVALKNVRVQT |
| *Xenopus tropicalis* | QYEPVAEIGVGAYGTVYKARDLQSGKFVALKNVRVQT |

**Figure S5b**

| | |
|---|---|
| **Cdk4** | |
| *Homo sapiens* | RYEPVAEIGVGAYGTVYKARDPHSGHFVALKSVRVPN |
| **bony fish** | |
| *Acanthochromis polyacanthus* | QYEPVAEIGGGAYGTVYKARDTESGQFVALKSVRVQT |
| *Acanthopagrus latus* | QYEPVAEIGGGAYGTVYKARDTESGQFVALKSVRVQT |
| *Acipenser ruthenus* 1 | QYEPVAEIGGGAYGTVYKARDRQSGKFVALKSVRVQT |
| *Acipenser ruthenus* 2 | QYEPVAEIGGGAYGTVYKARDRQSGKFVALKSVRVQT |
| *Albula glossodonta* | QYEPVAEIGGGAYGTVYKARDRQSGQFVALKSVRVQT |
| *Alosa alosa* | QYEPVAQIGGGAYGTVYKARDRESGQFVALKSVRVQT |
| *Alosa sapidissima* | QYEPVAQIGGGAYGTVYKARDRESGQFVALKSVRVQT |
| *Ameiurus melas* | QYEPVAEIGGGAYGTVYKARDKESGKFVALKSVRVQT |
| *Amphilophus citrinellus* | QYEPVAEIGGGAYGTVYKARDTESGQFVALKSVRVRT |
| *Amphiprion ocellaris* | QYEPVAEIGGGAYGTVYKARDTESGQFVALKSVRVQT |
| *Amphiprion percula* | QYEPVAEIGGGAYGTVYKARDTESGQFVALKSVRVQT |
| *Anabarilius grahami* | QYEPVAEIGGGAYGTVYKARDRDSGQFVALKSVRVQT |
| *Anabas testudineus* | QYEPVAEIGGGAYGTVYKARDTESGKFVALKSVRVPT |
| *Anarrhichthys ocellatus* | QYEPVAEIGGGAYGTVYKARDLESGQFVALKSVRVQT |
| *Anguilla anguilla* | QYERVAEIGGGAYGTVYKARDRLSGQFVALKSVRVQT |
| *Archocentrus centrarchus* | QYEPVAEIGGGAYGTVYKARDTESGQFVALKSVRVRT |
| *Astatotilapia calliptera* | QYDPVAEIGGGAYGTVYKARDIESGQFVALKSVHVRT |
| *Astyanax mexicanus* | QYEPVAEIGGGAYGTVYKARDTQSGQFVALKSVRVQT |
| *Austrofundulus limnaeus* | QYEPVAEIGGGAYGTVYKARDTESGKFVALKNVRVQA |
| *Bagarius yarrelli* | QYEPVAEIGGGAYGTVYKARDKESGKFVALKSVRVQT |
| *Betta splendens* | QYEPVAEIGGGAYGTVYKARDTESGKFVALKSVRVQT |
| *Boleophthalmus pectinirostris* | PYEPVAEIGGGAYGTVYKARDTTSGEFVALKSVRVQT |
| *Caranx melampygus* | QYEPVAEIGGGAYGTVYKARDTESGQFVALKSVRVQT |
| *Carassius auratus* 1 | QYEPVAEIGGGAYGTVYKARDRDSGQFVALKSVRVQT |
| *Carassius auratus* 2 | QYEPVAEIGGGAYGTVYKARDRDSGQFVALKSVRVQT |
| *Channa argus* | QYEPVAEIGGGAYGTVYKARDTESGQFVALKSVRVQT |
| *Chanos chanos* | QYEPVAEIGGGAYGTVYKARDKESGQFVALKSVRVQT |
| *Cheilinus undulatus* | QYEPVAEIGGGAYGTVYKARDTESGEFVALKSVRVQT |
| *Chelmon rostratus* | QYEPVAEIGGGAYGTVYKARDTESGQFVALKSVRVQT |
| *Clarias magur* | QYEPVAEIGGGAYGTVYKARDKESGKFVALKSVRVQT |
| *Clupea harengus* | QYEPVAQIGGGAYGTVYKARDRESGQFVALKSVRVQT |
| *Collichthys lucidus* | QYEPVAEIGGGAYGTVYKARDIESGQFVALKSVRVQT |
| *Colossoma macropomum* | QYEPVAEIGGGAYGTVYKARDRESGQFVALKSVRVQT |
| *Coregonus clupeaformis* 1 | QYEPVAEIGGGAYGTVYKARDLESGQFVALKSVRVQT |
| *Coregonus clupeaformis* 2 | QYEPVAEIGGGAYGTVYKARDLESGQFVALKSVRVQT |
| *Coryphaenoides rupestris* | PYEQVAEIGGGAYGTVYKARDLESGKFVALKSVRVQT |
| *Cottoperca gobio* | QYEPVAEIGGGAYGTVYKARDTESGQFVALKSVRVQT |
| *Cyclopterus lumpus* | QYEPVAEIGGGAYGTVYKARDTESGQFVALKSVRVQT |
| *Cynoglossus semilaevis* | QYERVAEIGGGAYGTVYKARDTESGQFVALKSVRVQT |
| *Cyprinodon tularosa* | QYEPVAEIGGGAYGTVYKARDTESGKFVALKNVRVQT |
| *Cyprinodon variegatus* | QYEPVAEIGGGAYGTVYKARDTESGKFVALKNVRVQT |
| *Cyprinus carpio* 1 | QYEPVAEIGGGAYGTVYKARDRDSGQFVALKSVRVQT |
| *Cyprinus carpio* 2 | QYEPVAEIGGGAYGTVYKARDRDSGQFVALKSVRVQT |
| *Danio rerio* | QYEPVAEIGGGAYGTVYKARDRDSGQFVALKSVRVQT |
| *Denticeps clupeoides* | KYEPVAQIGGGAYGTVYKARDRVSGQFVALKSVRVQT |
| *Dicentrarchus labrax* | QYEPVAEIGGGAYGTVYKARDTESGQFVALKSVRVQT |
| *Echeneis naucrates* | QYEPVAEIGGGAYGTVYKARDIESGQFVALKSVRVQT |
| *Electrophorus electricus* | QYEPVAEIGGGAYGTVYKARDKQSGQFVALKSVRVQT |
| *Epinephelus coioides* | -----AEIGGGAYGTVYKARDTESGQFVALKSVRVQT |
| *Epinephelus lanceolatus* | QYEPVAEIGGGAYGTVYKARDTESGQFVALKSVRVQT |
| *Erpetoichthys calabaricus* | QYEPVAEIGGGAYGTVYKARDLQSGKFVALKSVRVQR |
| *Esox lucius* | QYEPVAEIGGGAYGTVYKARHVESGQFVALKSVRVQT |
| *Etheostoma cragini* | QYEPVAEIGGGAYGTVYKARDTESGQFVALKSVRVQT |
| *Etheostoma spectabile* | QYEPVAEIGGGAYGTVYKARDTESGQFVALKSVRVQT |
| *Fundulus heteroclitus* | QYEPVAEIGGGAYGTVYKARDTESGKFVALKNVRVQT |
| *Gadus morhua* | PYEEVAEIGGGAYGTVYKARDTESGKFVALKSVRVQT |
| *Gambusia affinis* | QYEPVAEIGGGAYGTVYKARDTESGKFVALKNVRVQT |
| *Gasterosteus aculeatus aculeatus* | QYEPVAEIGGGAYGTVYKARDTESGKFVALKSVRVQA |
| *Girardinichthys multiradiatus* | QYEPVAEIGGGAYGTVYKARDTESGEFVALKNVRVQT |
| *Gymnodraco acuticeps* | QYEPVAEIGGGAYGTVYKARDTESGQFVALKSVRVQT |
| *Haplochromis burtoni* | QYDPVAEIGGGAYGTVYKARDIESGQFVALKSVRVRT |
| *Hippocampus comes* | QYERVAEIGGGAYGTVYKARDTESGRFVALKSVRVQT |
| *Hippoglossus hippoglossus* | QYEPVAEIGGGAYGTVYKARDTESGQFVALKSVRVQT |
| *Hippoglossus stenolepis* | QYEPVAEIGGGAYGTVYKARDTESGQFVALKSVRVQT |
| *Hucho hucho* 1 | QYEPVAEIGGGAYGTVYKARDLESGQFVALKSVRVQT |
| *Hucho hucho* 2 | QYEPVAEIGGGAYGTVYKARDLESGQFVALKSVRVQT |
| *Hypomesus transpacificus* | QYEPVAEIGGGAYGTVYKARDMASGQFVALKSVRVQT |
| *Iconisemion striatum* | PYEPVAEIGGGAYGTVYKARDTESGKFVALKNVRVQT |
| *Ictalurus punctatus* | QYEPVAEIGGGAYGTVYKARDKESGKFVALKSVRVQT |
| *Kryptolebias marmoratus* | QYEPVAEIGGGAYGTVYKARDTESGKFVALKNVRVQT |
| *Labeo rohita* | QYEPVAEIGGGAYGTVYKARDKDSGQFVALKSVRVQT |
| *Labrus bergylta* | QYEPVAEIGGGAYGTVYKARDTESGQFVALKSVRVQT |
| *Larimichthys crocea* | QYEPVAEIGGGAYGTVYKARDIESGQFVALKSVRVQT |
| *Lates calcarifer* | QYEPVAEIGGGAYGTVYKARDTESGQFVALKSVRVQT |
| *Latimeria chalumnae* | LYEPVAKIGGGAYGTVYKARDLQSGKFVALKSVKVHT |
| *Lepisosteus oculatus* | QYEPVAEIGGGAYGTVYKARDRQSGQFVALKSVRVQT |
| *Liparis tanakae* | QYEPVAEIGGGAYGTVYKARDTESGQFVALKSVRVQT |
| *Lucifuga dentata* | QYEPVAEIGGGAYGTVYKARDTESGQFVALKNVRVQT |
| *Mastacembelus armatus* | QYEPVAEIGGGAYGTVYKARDTESGQFVALKSVRVQT |
| *Maylandia zebra* | QYDPVAEIGGGAYGTVYKARDIESGQFVALKSVHVRT |
| *Megalobrama amblycephala* | QYEPVAEIGGGAYGTVYKARDRDSGQFVALKSVRVQT |
| *Megalops atlanticus* | QYEPVAEIGGGAYGTVYKARDRQSGQFVALKSVRVQT |
| *Megalops cyprinoides* | QYEPVAEIGGGAYGTVYKARDRQSGQFVALKSVRVQT |
| *Melanotaenia boesemani* | HYEKVAEIGGGAYGTVYKARDTESGKFVALKHVRVQT |
| *Micropterus dolomieu* | QYEPLAEIGGGAYGTVYKARDTESGQFVALKSVRVQT |
| *Micropterus salmoides* | QYEPLAEIGGGAYGTVYKARDTESGQFVALKSVRVQT |
| *Mola mola* | QYERVAEIGGGAYGTVYKARDTESGRFVALKSVRVQT |
| *Monopterus albus* | QYEPVAEIGGGAYGTVFKARDIQSGQFVALKSVRVQT |
| *Morone saxatilis* | QYEPVAEIGGGAYGTVYKARDTESGQFVALKSVRVQT |
| *Mugil cephalus* | QYEPVAEIGGGAYGTVYKARDTESGQFVALKSVRVQT |
| *Myripristis murdjan* | QYEPVAEIGGGAYGTVYKARDMESGQFVALKSVRVQT |
| *Nematolebias whitei* | HYEPVAEIGGGAYGTVYKARDTESGKFVALKNVRVQT |
| *Neogobius melanostomus* | FYEPVAEIGGGAYGTVYKARDKESGQFVALKSVRVQT |
| *Neolamprologus brichardi* | QYDPVAEIGGGAYGTVYKARDIESGQFVALKSVRVRT |
| *Nibea albiflora* | QYEPVAEIGGGAYGTVYKARDTESGQFVALKSVRVQT |
| *Nothobranchius furzeri* | PYEPVAEIGGGAYGTVYKARDTESGKFVALKNVRVQT |
| *Nothobranchius kadleci* | PYEPVAEIGGGAYGTVYKARDTESGKFVALKNVRVQT |
| *Nothobranchius korthausae* | PYEPVAEIGGGAYGTVYKARDTESGKFVALKNVRVQT |
| *Nothobranchius kuhntae* | PYEPVAEIGGGAYGTVYKARDTESGKFVALKNVRVQT |
| *Nothobranchius pienaari* | PYEPVAEIGGGAYGTVYKARDTESGKFVALKNVRVQT |
| *Nothobranchius rachovii* | PYEPVAEIGGGAYGTVYKARDTESGKFVALKNVRVQT |
| *Notolabrus celidotus* | QYEPVAEIGGGAYGTVYKARDTESGQFVALKSVRVQT |
| *Notothenia coriiceps* | QYEPVAEIGGGAYGTVYKARDTESGQFVALKSVRVQT |
| *Oncorhynchus gorbuscha* 1 | QYEPVAEIGGGAYGTVYKARDLESGQFVALKSVRVQT |
| *Oncorhynchus gorbuscha* 2 | QYEPVAEIGGGAYGTVYKARDLESGQFVALKSVRVQT |
| *Oncorhynchus keta* 1 | QYEPVAEIGGGAYGTVYKARDLESGQFVALKSVRVQT |
| *Oncorhynchus keta* 2 | QYEPVAEIGGGAYGTVFKARDLESGQFVALKSVRVQT |
| *Oncorhynchus kisutch* 1 | QYEPVAEIGGGAYGTVYKARDLESGQFVALKSVRVQT |
| *Oncorhynchus kisutch* 2 | QYEPVAEIGGGAYGTVYKARDLESGQFVALKSVRVQT |
| *Oncorhynchus mykiss* 1 | QYEPVAEIGGGAYGTVYKARDLESGQFVALKSVRVQT |
| *Oncorhynchus mykiss* 2 | QYEPVAEIGGGAYGTVYKARDLESGQFVALKSVRVQT |
| *Oncorhynchus nerka* 1 | QYEPVAEIGGGAYGTVYKARDLESGQFVALKSVRVQT |
| *Oncorhynchus nerka* 2 | QYEPVAEIGGGAYGTVYKARDLESGQFVALKSVRVQT |
| *Oncorhynchus tshawytscha* 1 | QYEPVAEIGGGAYGTVYKARDLESGQFVALKSVRVQT |
| *Oncorhynchus tshawytscha* 2 | QYEPVAEIGGGAYGTVYKARDLESGQFVALKSVRVQT |
| *Onychostoma macrolepis* | QYEPVAEIGGGAYGTVYKARDRDSGQFVALKSVRVQT |
| *Oreochromis aureus* | QYDPVAEIGGGAYGTVYKARDIESGQFVALKSVHVRT |
| *Oreochromis niloticus* | QYDPVAEIGGGAYGTVYKARDIESGQFVALKSVHVRT |
| *Oryzias javanicus* | QYEPVAEIGGGAYGTVYKARDTESGKFVALKNVRVQT |
| *Oryzias latipes* | QYEPVAEIGGGAYGTVYKARDIESGKFVALKNVRVQT |
| *Oryzias melastigma* | QYEPVAEIGGGAYGTVYKARDIESGKFVALKNVRVQT |
| *Oryzias sinensis* | QYEPVAEIGGGAYGTVYKARDIESGKFVALKNVRVQT |
| *Paralichthys olivaceus* | QYEPVAEIGGGAYGTVYKARDTESGQFVALKSVRVQT |
| *Parambassis ranga* | QYEPVAEIGGGAYGTVYKARDTESGKFVALKSVRVQT |
| *Paramormyrops kingsleyae* | QYERVAEIGGGAYGTVYKARDRQSGQFVALKSVRVQT |
| *Perca flavescens* | QYEPLAEIGGGAYGTVYKARDTESGQFVALKSVRVQT |
| *Perca fluviatilis* | QYEPLAEIGGGAYGTVYKARDTESGQFVALKSVRVQT |
| *Periophthalmus magnuspinnatus* | HYEPVAEIGGGAYGTVYKARNTESGQFVALKSVRVQT |
| *Pimephales promelas* | QYEPVAEIGGGAYGTVYKARDRDSGQFVALKSVRVQT |
| *Plectropomus leopardus* | QYEPVAEIGGGAYGTVYKARDTESGQFVALKSVRVQT |
| *Poecilia formosa* | QYEPVAEIGGGAYGTVYKARDTESGKFVALKNVRVQT |
| *Poecilia latipinna* | QYEPVAEIGGGAYGTVYKARDTESGKFVALKNVRVQT |
| *Poecilia mexicana* | QYEPVAEIGGGAYGTVYKARDTESGKFVALKNVRVQT |
| *Poecilia reticulata* | QYEPVAEIGGGAYGTVYKARDTESGKFVALKNVRVQT |
| *Poeciliopsis prolifica* | QYEPVAEIGGGAYGTVYKARDTESGKFVALKNVRVQT |
| *Polyodon spathula* | QYEPVAEIGGGAYGTVYKARDLQSGKFVALKSVRVQT |
| *Polypterus senegalus* | QYEPVAEIGGGAYGTVYKARDLQSGKFVALKSVRVQR |
| *Protopterus annectens* | QYEPVAEIGGGAYGTVYKARDLQSGKFVALKNVRVQT |
| *Pseudochaenichthys georgianus* | QYEPVAEIGGGAYGTVYKARDTESGQFVALKSVRVQT |
| *Pundamilia nyererei* | QYDPVAEIGGGAYGTVYKARDIESGQFVALKSVRVRT |
| *Pungitius pungitius* | QYEPVAEIGGGAYGTVYKARDTESGKFVALKSVRVQA |
| *Puntigrus tetrazona* 1 | QYEPVAEIGGGAYGTVYKARDRDSGQFVALKSVRVQT |
| *Puntigrus tetrazona* 2 | QYEPVAEIXGGAYGTVYKARDRDSGQFVALKSVRVQT |
| *Pygocentrus nattereri* | QYEPVAEIGGGAYGTVYKARDRESGQFVALKSVRVQT |
| *Salarias fasciatus* | QYEPVAEIGGGAYGTVYKARDTESGQFVALKSVRVQT |
| *Salmo salar* 1 | QYEPVAEIGGGAYGTVYKARDLESGQFVALKSVRVQT |
| *Salmo salar* 2 | QYEPVAEIGGGAYGTVYKARDLESGQFVALKSVRVQT |
| *Salmo trutta* 1 | QYEPVAEIGGGAYGTVYKARDLESGQFVALKSVRVQT |
| *Salmo trutta* 2 | QYEPVAEIGGGAYGTVYKARDLESGQFVALKSVRVQT |
| *Salvelinus alpinus* 1 | QYEPVAEIGGGAYGTVYKARDLESGQFVALKSVRVQT |
| *Salvelinus alpinus* 2 | QYEPVAEIGGGAYGTVYKARDLESGQFVALKSVRVQT |
| *Salvelinus namaycush* 1 | QYEPVAEIGGGAYGTVYKARDLESGQFVALKSVRVQT |
| *Salvelinus namaycush* 2 | QYEPVAEIGGGAYGTVYKARDLESGQFVALKIVRVQT |
| *Sander lucioperca* | QYEPLAEIGGGAYGTVYKARDTESGQFVALKSVRVQT |
| *Scatophagus argus* | QYEPVAEIGGGAYGTVYKARDTESGQFVALKSVRVQT |
| *Scleropages formosus* | QYERVAEIGGGAYGTVYKARDRQSGQFVALKSVRVQT |
| *Scophthalmus maximus* | QYEPVAEIGGGAYGTVYKARDTESGQFVALKSVRVQT |
| *Sebastes umbrosus* | QYEPVAEIGGGAYGTVFKARDIESGQFVALKSVRVQT |
| *Seriola dumerili* | QYEPVAEIGGGAYGTVYKARDTESGQFVALKSVRVQT |
| *Seriola lalandi dorsalis* | QYEPVAEIGGGAYGTVYKARDTESGQFVALKSVRVQT |
| *Silurus meridionalis* | QYERVAEIGGGAYGTVYKARDTESGKFVALKSVRVHT |
| *Simochromis diagramma* | QYDPVAEIGGGAYGTVYKARDIESGQFVALKSVRVRT |
| *Siniperca chuatsi* | QYEPVAEIGGGAYGTVYKARDTESGQFVALKSVRVQT |
| *Sinocyclocheilus anshuiensis* 1 | QYEPVAEIGGGAYGMVYKARDRNTGQFVALKSVRVQT |
| *Sinocyclocheilus anshuiensis* 2 | QYEPVAEIGGGAYGTVYKARDRDSGQFVALKSVRVQT |
| *Sinocyclocheilus grahami* 1 | QYEPVAEIGGGAYGTVYKARDRDSGQFVALKSVRVQT |
| *Sinocyclocheilus grahami* 2 | QYEPVAEIGGGAYGTVYKARDRDSGQFVALKSVRVQT |
| *Sinocyclocheilus rhinocerous* 1 | QYEPVAEIGGGAYGTVYKARDRDSGQFVALKSVRVQT |
| *Sinocyclocheilus rhinocerous* 2 | QYEPVAEIGGGAYGTVYKARDRDSGQFVALKSVRVQT |
| *Solea senegalensis* | QYERVAEIGGGAYGTVYKARDMESGQFVALKSVRVQT |
| *Sparus aurata* | QYEPVAEIGGGAYGTVYKARDTESGQFVALKSVRVQT |
| *Sphaeramia orbicularis* | QYEPVAEIGGGAYGTVYKARDTESGQFVALKSVRVQT |
| *Stegastes partitus* | QYEPVAEIGGGAYGTVYKARDTESGQFVALKSVRVQT |
| *Syngnathus acus* | QYERVAEIGGGAYGTVYKARDMESGRFVALKSVRVKT |
| *Tachysurus fulvidraco* | QYEPVAEIGGGAYGTVYKARDKESGKFVALKSVRVQT |
| *Takifugu flavidus* | QYEPVAEIGGGAYGTVYKARDTESGKFVALKSVRVQT |
| *Takifugu rubripes* | QYEPVAEIGGGAYGTVYKARDTESGKFVALKSVRVQT |
| *Tetraodon nigroviridis* | QYEPVAEIGGGAYGTVYKARDTESGKFVALKSVRVQT |
| *Thalassophryne amazonica* | QYEPVAEIGGGAYGTVYKARDTESGQFVALKSVRVQT |
| *Thunnus albacares* | QYEPVAEIGGGAYGTVYKARDTESGQFVALKSVRVQT |
| *Thunnus maccoyii* | QYEPVAEIGGGAYGTVYKARDTESGQFVALKSVRVQT |
| *Toxotes jaculatrix* | QYEPVAEIGGGAYGTVYKARDTESGQFVALKSVRVQT |
| *Trematomus bernacchii* | QYEPVAEIGGGAYGTVYKARDTESGQFVALKSVRVQT |
| *Triplophysa tibetana* | QYEPVAEIGGGAHGTVYKARDKDSGQFVALKSVRVQT |
| *Xiphias gladius* | QYEPVAEIGGGAYGTVYKARDTESGQFVALKSVRVQT |
| *Xiphophorus couchianus* | QYEPVAEIGGGAYGTVYKARDTESGKFVALKNVRVQT |
| *Xiphophorus hellerii* | QYEPVAEIGGGAYGTVYKARDTESGKFVALKNVRVQT |
| *Xiphophorus maculatus* | QYEPVAEIGGGAYGTVYKARDTESGKFVALKNVRVQT |
| **cartilaginous fish** | |
| *Amblyraja radiata* | QYEPVAEIGGGAYGTVYKARDSRSGKFVALKNVRVHR |
| *Callorhinchus milii* | QYEPVAEIGGGAYGTVYKARDSRSGKFVALKIVRVQM |
| *Carcharodon carcharias* | PYEPVAEIGGGAYGTVYKARDSRSGKFVALKNVRVQR |
| *Chiloscyllium plagiosum* | -YEPVAEIGEGAYGTVYKARDSLSGKFVALKNVRVQR |
| *Rhincodon typus* | QYEPVAEIGEGAYGTVYKARDSLSGKFVALKNVRVQR |
| *Stegostoma fasciatum* | QYEPMAEIGEGAYGTVYKARDSLSGKFVALKNVRVQR |

**Figure S5c**

| | |
|---|---|
| **Cdk4** | |
| *Homo sapiens* | RYEPVAEIGVGAYGTVYKARD-PHSGHFVALKSVRVPN |
| **Cdk6** | |
| **mammal** | |
| **<placentarian>** | |
| *Acinonyx jubatus* | QYECVAEIGEGAYGKVFKARDLKNGGRFVALKRVRVQT |
| *Ailuropoda melanoleuca* | QYECVAEIGEGAYGKVFKARDLKNGGRFVALKRVRVQT |
| *Aotus nancymaae* | QYECVAEIGEGAYGKVFKARDLKNGGRFVALKRVRVQT |
| *Artibeus jamaicensis* | QYECVAEIGEGAYGKVFKARDLKNGGRFVALKRVRVQT |
| *Arvicanthis niloticus* | QYECVAEIGEGAYGKVFKARDLKNGGRFVALKRVRVQT |
| *Arvicola amphibius* | QYECVAEIGEGAYGKVFKARDLKNGGRFVALKRVRVQT |
| *Balaenoptera acutorostrata scammoni* | QYECVAEIGEGAYGKVFKARDLKNGGRFVALKRVRVQT |
| *Balaenoptera musculus* | QYECVAEIGEGAYGKVFKARDLKNGGRFVALKRVRVQT |
| *Bison bison bison* | QYECVAEIGEGAYGKVFKARDLKNGGRFVALKRVRVQT |
| *Bos indicus* | QYECVAEIGEGAYGKVFKARDLKNGGRFVALKRVRVQT |
| *Bos mutus* | QYECVAEIGEGAYGKVFKARDLKNGGRFVALKRVRVQT |
| *Bos taurus* | QYECVAEIGEGAYGKVFKARDLKNGGRFVALKRVRVQT |
| *Bubalus bubalis* | QYECVAEIGEGAYGKVFKARDLKNGGRFVALKRVRVQT |
| *Callithrix jacchus* | QYECVAEIGEGAYGKVFKARDLKNGGRFVALKRVRVQT |
| *Callorhinus ursinus* | QYECVAEIGEGAYGKVFKARDLKNGGRFVALKRVRVQT |
| *Camelus bactrianus* | QYECVAEIGEGAYGKVFKARDLKNGGRFVALKRVRVQT |
| *Camelus dromedarius* | QYECVAEIGEGAYGKVFKARDLKNGGRFVALKRVRVQT |
| *Camelus ferus* | QYECVAEIGEGAYGKVFKARDLKNGGRFVALKRVRVQT |
| *Canis lupus familiaris* | QYECVAEIGEGAYGKVFKARDLKNGGRFVALKRVRVQT |
| *Capra hircus* | QYECVAEIGEGAYGKVFKARDLKNGGRFVALKRVRVQT |
| *Castor canadensis* | QYECVAEIGEGAYGKVFKARDLKNGGRFVALKRVRVQT |
| *Cavia porcellus* | QYECVAEIGEGAYGKVFKARDLKNGGRFVALKRVRVQT |
| *Cebus imitator* | QYECVAEIGEGAYGKVFKARDLKNGGRFVALKRVRVQT |
| *Ceratotherium simum simum* | QYECVAEIGEGAYGKVFKARDLKNGGRFVALKRVRVQT |
| *Cercocebus atys* | QYECVAEIGEGAYGKVFKARDLKNGGRFVALKRVRVQT |
| *Cervus canadensis* | QYECVAEIGEGAYGKVFKARDLKNGGRFVALKRVRVQT |
| *Cervus elaphus* | QYECVAEIGEGAYGKVFKARDLKNGGRFVALKRVRVQT |
| *Cervus hanglu yarkandensis* | QYECVAEIGEGAYGKVFKARDLKNGGRFVALKRVRVQT |
| *Chinchilla lanigera* | QYECVAEIGEGAYGKVFKARDLKNGGRFVALKRVRVQT |
| *Chlorocebus sabaeus* | QYECVAEIGEGAYGKVFKARDLKNGGRFVALKRVRVQT |
| *Choloepus didactylus* | QYECVAEIGEGAYGKVFKARDLKNGGRFVALKRVRVQT |
| *Chrysochloris asiatica* | QYECVAEIGEGAYGKVFKARDLKNGGRFVALKRVRVQT |
| *Colobus angolensis palliatus* | QYECVAEIGEGAYGKVFKARDLKNGGRFVALKRVRVQT |
| *Condylura cristata* | QYECVAEIGEGAYGKVFKARDLKNGGRFVALKRVRVQT |
| *Cricetulus griseus* | QYECVAEIGEGAYGKVFKARDLKNGGRFVALKRVRVQT |
| *Crocuta crocuta* | QYECVAEIGEGAYGKVFKARDLKNGGRFVALKRVRVQT |
| *Dasypus novemcinctus* | QYECVAEIGEGAYGKVFKARDLKNGGRFVALKRVRVQT |
| *Delphinapterus leucas* | QYECVAEIGEGAYGKVFKARDLKNGGRFVALKRVRVQT |
| *Desmodus rotundus* | QYECVAEIGEGAYGKVFKARDLKNGGRFVALKRVRVQT |
| *Dipodomys ordii* | QYECVAEIGEGAYGKVFKARDLKNGGRFVALKRVRVQT |
| *Dipodomys spectabilis* | QYECVAEIGEGAYGKVFKARDLKNGGRFVALKRVRVQT |
| *Echinops telfairi* | QYECVAEIGEGAYGKVFKARDLKNGGRFVALKRVRVQT |
| *Elephantulus edwardii* | QYECVAEIGEGAYGKVFKARDLKNGGRFVALKRVRVQT |
| *Enhydra lutris kenyoni* | QYECVAEIGEGAYGKVFKARDLKNGGRFVALKRVRVQT |
| *Eptesicus fuscus* | QYECVAEIGEGAYGKVFKARDLKNGGRFVALKRVRVQT |
| *Equus asinus* | QYECVAEIGEGAYGKVFKARDLKNGGRFVALKRVRVQT |
| *Equus caballus* | QYECVAEIGEGAYGKVFKARDLKNGGRFVALKRVRVQT |
| *Equus przewalskii* | QYECVAEIGEGAYGKVFKARDLKNGGRFVALKRVRVQT |
| *Equus quagga* | QYECVAEIGEGAYGKVFKARDLKNGGRFVALKRVRVQT |
| *Erinaceus europaeus* | QYECVAEIGEGAYGKVFKARDLKNGGRFVALKRVRVQT |
| *Eumetopias jubatus* | QYECVAEIGEGAYGKVFKARDLKNGGRFVALKRVRVQT |
| *Felis catus* | QYECVAEIGEGAYGKVFKARDLKNGGRFVALKRVRVQT |
| *Fukomys damarensis* | AATRQAEFRSGGGGG---ARDLKNGGRFVALKRVRVQT |
| *Galemys pyrenaicus* | QYECVAEIGEGAYGKVFKARDLKNGGRFVALKRVRVQT |
| *Galeopterus variegatus* | QYECVAEIGEGAYGKVFKARDLKNGGRFVALKRVRVQT |
| *Globicephala melas* | QYECVAEIGEGAYGKVFKARDLKNGGRFVALKRVRVQT |
| *Gorilla gorilla gorilla* | QYECVAEIGEGAYGKVFKARDLKNGGRFVALKRVRVQT |
| *Grammomys surdaster* | QYECVAEIGEGAYGKVFKARDLKNGGRFVALKRVRVQT |
| *Gulo gulo luscus* | QYECVAEIGEGAYGKVFKARDLKNGGRFVALKRVRVQT |
| *Halichoerus grypus* | QYECVAEIGEGAYGKVFKARDLKNGGRFVALKRVRVQT |
| *Heterocephalus glaber* | QYECVAEIGEGAYGKVFKARDLKNGGRFVALKRVRVQT |
| *Hipposideros armiger* | QYECVAEIGEGAYGKVFKARDLKNGGRFVALKRVRVQT |
| *Homo sapiens* | QYECVAEIGEGAYGKVFKARDLKNGGRFVALKRVRVQT |
| *Hyaena hyaena* | QYECVAEIGEGAYGKVFKARDLKNGGRFVALKRVRVQT |
| *Hylobates moloch* | QYECVAEIGEGAYGKVFKARDLKNGGRFVALKRVRVQT |
| *Ictidomys tridecemlineatus* | QYECVAEIGEGAYGKVFKARDLKNGGRFVALKRVRVQT |
| *Jaculus jaculus* | QYECVAEIGEGAYGKVFKARDLKNGGRFVALKRVRVQT |
| *Lagenorhynchus obliquidens* | QYECVAEIGEGAYGKVFKARDLKNGGRFVALKRVRVQT |
| *Lemur catta* | QYECVAEIGEGAYGKVFKARDLKNGGRFVALKRVRVQT |
| *Leopardus geoffroyi* | QYECVAEIGEGAYGKVFKARDLKNGGRFVALKRVRVQT |
| *Leptonychotes weddellii* | QYECVAEIGEGAYGKVFKARDLKNGGRFVALKRVRVQT |
| *Lipotes vexillifer* | QYECVAEIGEGAYGKVFKARDLKNGGRFVALKRVRVQT |
| *Lontra canadensis* | QYECVAEIGEGAYGKVFKARDLKNGGRFVALKRVRVQT |
| *Loxodonta africana* | QYECVAEIGEGAYGKVFKARDLKNGGRFVALKRVRVQT |
| *Lynx canadensis* | QYECVAEIGEGAYGKVFKARDLKNGGRFVALKRVRVQT |
| *Macaca fascicularis* | QYECVAEIGEGAYGKVFKARDLKNGGRFVALKRVRVQT |
| *Macaca mulatta* | QYECVAEIGEGAYGKVFKARDLKNGGRFVALKRVRVQT |
| *Macaca nemestrina* | QYECVAEIGEGAYGKVFKARDLKNGGRFVALKRVRVQT |
| *Mandrillus leucophaeus* | QYECVAEIGEGAYGKVFKARDLKNGGRFVALKRVRVQT |
| *Manis javanica* | QYECVAEIGEGAYGKVFKARDLKNGGRFVALKRVRVQT |
| *Manis pentadactyla* | QYECVAEIGEGAYGKVFKARDLKNGGRFVALKRVRVQT |
| *Marmota flaviventris* | QYECVAEIGEGAYGKVFKARDLKNGGRFVALKRVRVQT |
| *Marmota marmota marmota* | QYECVAEIGEGAYGKVFKARDLKNGGRFVALKRVRVQT |
| *Marmota monax* | QYECVAEIGEGAYGKVFKARDLKNGGRFVALKRVRVQT |
| *Mastomys coucha* | QYECVAEIGEGAYGKVFKARDLKNGGRFVALKRVRVQT |
| *Meles meles* | QYECVAEIGEGAYGKVFKARDLKNGGRFVALKRVRVQT |
| *Meriones unguiculatus* | QYECVAEIGEGAYGKVFKARDLKNGGRFVALKRVRVQT |
| *Mesocricetus auratus* | QYECVAEIGEGAYGKVFKARDLKNGGRFVALKRVRVQT |
| *Microcebus murinus* | QYECVAEIGEGAYGKVFKARDLKNGGRFVALKRVRVQT |
| *Microtus ochrogaster* | QYECVAEIGEGAYGKVFKARDLKNGGRFVALKRVRVQT |
| *Microtus oregoni* | QYECVAEIGEGAYGKVFKARDLKNGGRFVALKRVRVQT |
| *Miniopterus natalensis* | QYECVAEIGEGAYGKVFKARDLKNGGRFVALKRVRVQT |
| *Mirounga angustirostris* | QYECVAEIGEGAYGKVFKARDLKNGGRFVALKRVRVQT |
| *Mirounga leonina* | QYECVAEIGEGAYGKVFKARDLKNGGRFVALKRVRVQT |
| *Molossus molossus* | QYECVAEIGEGAYGKVFKARDLKNGGRFVALKRVRVQT |
| *Monodon monoceros* | QYECVAEIGEGAYGKVFKARDLKNGGRFVALKRVRVQT |
| *Mus caroli* | QYECVAEIGEGAYGKVFKARDLKNGGRFVALKRVRVQT |
| *Mus musculus* | QYECVAEIGEGAYGKVFKARDLKNGGRFVALKRVRVQT |
| *Mus pahari* | QYECVAEIGEGAYGKVFKARDLKNGGRFVALKRVRVQT |
| *Mustela erminea* | QYECVAEIGEGAYGKVFKARDLKNGGRFVALKRVRVQT |
| *Mustela putorius furo* | QYEYVAEIGEGAYGKVFKARDLKNGGRFVALKRVRVQT |
| *Myotis brandtii* | QYECVAEIGEGAYGKVFKARDLKNGGRFVALKRVRVQT |
| *Myotis lucifugus* | QYECVAEIGEGAYGKVFKARDLKNGGRFVALKRVRVQT |
| *Myotis myotis* | QYECVAEIGEGAYGKVFKARDLKNGGRFVALKRVRVQT |
| *Nannospalax galili* | QYECVAEIGEGAYGKVFKARDLKNGGRFVALKRVRVQT |
| *Neogale vison* | QYECVAEIGEGAYGKVFKARDLKNGGRFVALKRVRVQT |
| *Neomonachus schauinslandi* | QYECVAEIGEGAYGKVFKARDLKNGGRFVALKRVRVQT |
| *Neophocaena asiaeorientalis asiaeorientalis* | QYECVAEIGEGAYGKVFKARDLKNGGRFVALKRVRVQT |
| *Neosciurus carolinensis* | QYECVAEIGEGAYGKVFKARDLKNGGRFVALKRVRVQT |
| *Neotoma lepida* | QYECVAEXGEGAYGKVFKARDLKNGGRFVALKRVRVQT |
| *Nomascus leucogenys* | QYECVAEIGEGAYGKVFKARDLKNGGRFVALKRVRVQT |
| *Nyctereutes procyonoides* | QYECVAEIGEGAYGKVFKARDLKNGGRFVALKRVRVQT |
| *Ochotona curzoniae* | QYECVAEIGEGAYGKVFKARDLKNGGRFVALKRVRVQT |
| *Ochotona princeps* | QYECVAEIGEGAYGKVFKARDLKNGGRFVALKRVRVQT |
| *Octodon degus* | QYECVAEIGEGAYGKVFKARDLKNGGRFVALKRVRVQT |
| *Odobenus rosmarus divergens* | QYECVAEIGEGAYGKVFKARDLKNGGRFVALKRVRVQT |
| *Odocoileus virginianus texanus* | QYECVAEIGEGAYGKVFKARDLKNGGRFVALKRVRVQT |
| *Onychomys torridus* | QYECVAEIGEGAYGKVFKARDLKNGGRFVALKRVRVQT |
| *Orcinus orca* | QYECVAEIGEGAYGKVFKARDLKNGGRFVALKRVRVQT |
| *Orycteropus afer afer* | QYECVAEIGEGAYGKVFKARDLKNGGRFVALKRVRVQT |
| *Oryctolagus cuniculus* | QYECVAEIGEGAYGKVFKARDLKNGGRFVALKRVRVQT |
| *Oryx dammah* | QYECVAEIGEGAYGKVFKARDLKNGGRFVALKRVRVQT |
| *Otolemur garnettii* | QYECVAEIGEGAYGKVFKARDLKNGGRFVALKRVRVQT |
| *Ovis aries* | QYECVAEIGEGAYGKVFKARDLKNGGRFVALKRVRVQT |
| *Pan paniscus* | QYECVAEIGEGAYGKVFKARDLKNGGRFVALKRVRVQT |
| *Pan troglodytes* | QYECVAEIGEGAYGKVFKARDLKNGGRFVALKRVRVQT |
| *Panthera leo* | QYECVAEIGEGAYGKVFKARDLKNGGRFVALKRVRVQT |
| *Panthera pardus* | QYECVAEIGEGAYGKVFKARDLKNGGRFVALKRVRVQT |
| *Panthera tigris* | QYECVAEIGEGAYGKVFKARDLKNGGRFVALKRVRVQT |
| *Papio anubis* | QYECVAEIGEGAYGKVFKARDLKNGGRFVALKRVRVQT |
| *Perognathus longimembris pacificus* | QYECVAEIGEGAYGKVFKARDLKNGGRFVALKRVRVQT |
| *Peromyscus leucopus* | QYECVAEIGEGAYGKVFKARDLKNGGRFVALKRVRVQT |
| *Peromyscus maniculatus bairdii* | QYECVAEIGEGAYGKVFKARDLKNGGRFVALKRVRVQT |
| *Phoca vitulina* | QYECVAEIGEGAYGKVFKARDLKNGGRFVALKRVRVQT |
| *Phocoena sinus* | QYECVAEIGEGAYGKVFKARDLKNGGRFVALKRVRVQT |
| *Phyllostomus discolor* | QYECVAEIGEGAYGKVFKARDLKNGGRFVALKRVRVQT |
| *Phyllostomus hastatus* | QYECVAEIGEGAYGKVFKARDLKNGGRFVALKRVRVQT |
| *Physeter catodon* | QYECVAEIGEGAYGKVFKARDLKNGGRFVALKRVRVQT |
| *Piliocolobus tephrosceles* | QYECVAEIGEGAYGKVFKARDLKNGGRFVALKRVRVQT |
| *Pipistrellus kuhlii* | QYECVAEIGEGAYGKVFKARDLKNGGRFVALKRVRVQT |
| *Pongo abelii* | QYECVAEIGEGAYGKVFKARDLKNGGRFVALKRVRVQT |
| *Prionailurus bengalensis* | QYECVAEIGEGAYGKVFKARDLKNGGRFVALKRVRVQT |
| *Procavia capensis* | QYECVAEIGEGAYGKVFKARDLKNGGRFVALKRVRVQT |
| *Propithecus coquereli* | QYECVAEIGEGAYGKVFKARDLKNGGRFVALKRVRVQT |
| *Pteropus alecto* | QYECVAEIGEGAYGKVFKARDLKNGGRFVALKRVRVQT |
| *Pteropus giganteus* | QYECVAEIGEGAYGKVFKARDLKNGGRFVALKRVRVQT |
| *Pteropus vampyrus* | QYECVAEIGEGAYGKVFKARDLKNGGRFVALKRVRVQT |
| *Puma concolor* | QYECVAEIGEGAYGKVFKARDLKNGGRFVALKRVRVQT |
| *Puma yagouaroundi* | QYECVAEIGEGAYGKVFKARDLKNGGRFVALKRVRVQT |
| *Rattus norvegicus* | QYECVAEIGEGAYGKVFKARDLKNGGRFVALKRVRVQT |
| *Rattus rattus* | QYECVAEIGEGAYGKVFKARDLKNGGRFVALKRVRVQT |
| *Rhinolophus ferrumequinum* | QYECVAEIGEGAYGKVFKARDLKNGGRFVALKRVRVQT |
| *Rhinolophus sinicus* | QYECVAEIGEGAYGKVFKARDLKNGGRFVALKRVRVQT |
| *Rhinopithecus bieti* | QYECVAEIGEGAYGKVFKARDLKNGGRFVALKRVRVQT |
| *Rhinopithecus roxellana* | QYECVAEIGEGAYGKVFKARDLKNGGRFVALKRVRVQT |
| *Rousettus aegyptiacus* | QYECVAEIGEGAYGKVFKARDLKNGGRFVALKRVRVQT |
| *Saimiri boliviensis boliviensis* | QYECVAEIGEGAYGKVFKARDLKNGGRFVALKRVRVQT |
| *Sapajus apella* | QYECVAEIGEGAYGKVFKARDLKNGGRFVALKRVRVQT |
| *Sorex araneus* | QYECVAEIGEGAYGKVFKARDLKNGGRFVALKRVRVQT |
| *Sousa chinensis* | QYECVAEIGEGAYGKVFKARDLKNGGRFVALKRVRVQT |
| *Sturnira hondurensis* | QYECVAEIGEGAYGKVFKARDLKNGGRFVALKRVRVQT |
| *Suricata suricatta* | QYECVAEIGEGAYGKVFKARDLKNGGRFVALKRVRVQT |
| *Sus scrofa* | QYECVAEIGEGAYGKVFKARDLKNGGRFVALKRVRVQT |
| *Talpa occidentalis* | QYECVAEIGEGAYGKVFKARDLKNGGRFVALKRVRVQT |
| *Theropithecus gelada* | QYECVAEIGEGAYGKVFKARDLKNGGRFVALKRVRVQT |
| *Trachypithecus francoisi* | QYECVAEIGEGAYGKVFKARDLKNGGRFVALKRVRVQT |
| *Trichechus manatus latirostris* | QYECVAEIGEGAYGKVFKARDLKNGGRFVALKRVRVQT |
| *Tupaia chinensis* | QYECVAEIGEGAYGKVFKARDLKNGGRFVALKRVRVQT |
| *Tursiops truncatus* | QYECVAEIGEGAYGKVFKARDLKNGGRFVALKRVRVQT |
| *Urocitellus parryii* | QYECVAEIGEGAYGKVFKARDLKNGGRFVALKRVRVQT |
| *Ursus americanus* | QYECVAEIGEGAYGKVFKARDLKNGGRFVALKRVRVQT |
| *Ursus arctos* | QYECVAEIGEGAYGKVFKARDLKNGGRFVALKRVRVQT |
| *Ursus maritimus* | QYECVAEIGEGAYGKVFKARDLKNGGRFVALKRVRVQT |
| *Vicugna pacos* | QYECVAEIGEGAYGKVFKARDLKNGGRFVALKRVRVQT |
| *Vulpes lagopus* | QYECVAEIGEGAYGKVFKARDLKNGGRFVALKRVRVQT |
| *Vulpes vulpes* | QYECVAEIGEGAYGKVFKARDLKNGGRFVALKRVRVQT |
| *Zalophus californianus* | QYECVAEIGEGAYGKVFKARDLKNGGRFVALKRVRVQT |
| **<marsupial>** | |
| *Dromiciops gliroides* | QYECVAEIGEGAYGKVFKARDLKNGGRFVALKRVRVQT |
| *Gracilinanus agilis* | QYECVAEIGEGAYGKVFKARDLKNGGRFVALKRVRVQT |
| *Monodelphis domestica* | QYECVAEIGEGAYGKVFKARDLKNGGRFVALKRVRVQT |
| *Phascolarctos cinereus* | QYECVAEIGEGAYGKVFKARDLKNGGRFVALKRVRVQT |
| *Sarcophilus harrisii* | QYECVAEIGEGAYGKVFKARDLKNGGRFVALKRVRVQT |
| *Trichosurus vulpecula* | QYECVAEIGEGAYGKVFKARDLKNGGRFVALKRVRVQT |
| *Vombatus ursinus* | QYECVAEIGEGAYGKVFKARDLKNGGRFVALKRVRVQT |
| **<monotreme>** | |
| *Ornithorhynchus anatinus* | QYECVAEIGEGAYGKVFKARDLKNGGRFVALKRVRVQT |
| *Tachyglossus aculeatus* | QYECVAEIGEGAYGKVFKARDLKNGGRFVALKRVRVQT |

**Figure S5d**

**Cdk4**

| | |
|---|---|
| *Homo sapiens* | RYEPVAEIGVGAYGTVYKARDPHSG-HFVALKSVRVPN |

**Cdk6**

**bird**

| | |
|---|---|
| *Acanthisitta chloris* | QYECVAEIGEGAYGKVFKARDLKNGGRFVALKRVRVQT |
| *Accipiter nisus* | QYECVAEIGEGAYGKVFKARDLKNGGRFVALKRVRVQT |
| *Amazona aestiva* | QYECVAEIGEGAYGKVFKARDLKNGGRFVALKRVRVQT |
| *Amazona collaria* | QYECVAEIGEGAYGKVFKARDLKNGGRFVALKRVRVQT |
| *Anas platyrhynchos* | QYECVAEIGEGAYGKVFKARDLKNGGRFVALKRVRVQT |
| *Anas zonorhyncha* | QYECVAEIGEGAYGKVFKARDLKNGGRFVALKRVRVQT |
| *Anser brachyrhynchus* | QYECVAEIGEGAYGKVFKARDLKNGGRFVALKRVRVQT |
| *Anser cygnoides* | QYECVAEIGEGAYGKVFKARDLKNGGRFVALKRVRVQT |
| *Antrostomus carolinensis* | QYECVAEIGEGAYGKVFKARDLKNGGRFVALKRVRVQT |
| *Apaloderma vittatum* | QYECVAEIGEGAYGKVFKARDLKNGGRFVALKRVRVQT |
| *Aptenodytes forsteri* | QYECVAEIGEGAYGKVFKARDLKNGGRFVALKRVRVQT |
| *Aptenodytes patagonicus* | QYECVAEIGEGAYGKVFKARDLKNGGRFVALKRVRVQT |
| *Apteryx haastii* | QYECVAEIGEGAYGKVFKARDLKNGGRFVALKRVRVQT |
| *Apteryx mantelli mantelli* | QYECVAEIGEGAYGKVFKARDLKNGGRFVALKRVRVQT |
| *Apteryx owenii* | QYECVAEIGEGAYGKVFKARDLKNGGRFVALKRVRVQT |
| *Apteryx rowi* | QYECVAEIGEGAYGKVFKARDLKNGGRFVALKRVRVQT |
| *Aquila chrysaetos chrysaetos* | QYECVAEIGEGAYGKVFKARDLKNGGRFVALKRVRVQT |
| *Athene cunicularia* | QYECVAEIGEGAYGKVFKARDLKNGGRFVALKRVRVQT |
| *Aythya fuligula* | QYECVAEIGEGAYGKVFKARDLKNGGRFVALKRVRVQT |
| *Balearica regulorum gibbericeps* | QYECVAEIGEGAYGKVFKARDLKNGGRFVALKRVRVQT |
| *Bubo bubo* | QYECVAEIGEGAYGKVFKARDLKNGGRFVALKRVRVQT |
| *Buceros rhinoceros silvestris* | QYECVAEIGEGAYGKVFKARDLKNGGRFVALKRVRVQT |
| *Buteo japonicus* | QYECVAEIGEGAYGKVFKARDLKNGGRFVALKRVRVQT |
| *Cairina moschata domestica* | QYECVAEIGEGAYGKVFKARDLKNGGRFVALKRVRVQT |
| *Calidris pugnax* | QYECVAEIGEGAYGKVFKARDLKNGGRFVALKRVRVQT |
| *Calidris pygmaea* | QYECVAEIGEGAYGKVFKARDLKNGGRFVALKRVRVQT |
| *Callipepla squamata* | QYECVAEIGEGAYGKVFKARDLKNGGRFVALKRVRVQT |
| *Calypte anna* | QYECVAEIGEGAYGKVFKARDLKNGGRFVALKRVRVQT |
| *Camarhynchus parvulus* | QYECVAEIGEGAYGKVFKARDLKNGGRFVALKRVRVQT |
| *Cariama cristata* | QYECVAEIGEGAYGKVFKARDLKNGGRFVALKRVRVQT |
| *Cathartes aura* | QYECVAEIGEGAYGKVFKARDLKNGGRFVALKRVRVQT |
| *Catharus ustulatus* | QYECVAEIGEGAYGKVFKARDLKNGGRFVALKRVRVQT |
| *Centrocercus urophasianus* | QYECVAEIGEGAYGKVFKARDLKNGGRFVALKRVRVQT |
| *Chaetura pelagica* | QYECVAEIGEGAYGKVFKARDLKNGGRFVALKRVRVQT |
| *Charadrius vociferus* | QYECVAEIGEGAYGKVFKARDLKNGGRFVALKRVRVQT |
| *Chiroxiphia lanceolata* | QYEYVAEIGEGAYGKVFKARDLKNGGRFVALKRVRVQT |
| *Chlamydotis macqueenii* | QYECVAEIGEGAYGKVFKARDLKNGGRFVALKRVRVQT |
| *Chrysolophus pictus* | QYECVAEIGEGAYGKVFKARDLKNGGRFVALKRVRVQT |
| *Colinus virginianus* | QYECVAEIGEGAYGKVFKARDLKNGGRFVALKRVRVQT |
| *Colius striatus* | QYECVAEIGEGAYGKVFKARDLKNGGRFVALKRVRVQT |
| *Columba livia* | QYECVAEIGEGAYGKVFKARDLKNGGRFVALKRVRVQT |
| *Corapipo altera* | QYEYVAEIGEGAYGKVFKARDLKNGGRFVALKRVRVQT |
| *Corvus brachyrhynchos* | QYECVAEIGEGAYGKVFKARDLKNGGRFVALKRVRVQT |
| *Corvus cornix cornix* | QYECVAEIGEGAYGKVFKARDLKNGGRFVALKRVRVQT |
| *Corvus hawaiiensis* | QYECVAEIGEGAYGKVFKARDLKNGGRFVALKRVRVQT |
| *Corvus kubaryi* | QYECVAEIGEGAYGKVFKARDLKNGGRFVALKRVRVQT |
| *Corvus moneduloides* | QYECVAEIGEGAYGKVFKARDLKNGGRFVALKRVRVQT |
| *Coturnix japonica* | QYECVAEIGEGAYGKVFKARDLKNGGRFVALKRVRVQT |
| *Cuculus canorus* | QYECVAEIGEGAYGKVFKARDLKNGGRFVALKRVRVQT |
| *Cyanistes caeruleus* | QYECVAEIGEGAYGKVFKARDLKNGGRFVALKRVRVQT |
| *Cyanoderma ruficeps* | QYECVAEIGEGAYGKVFKARDLKNGGRFVALKRVRVQT |
| *Cygnus atratus* | QYECVAEIGEGAYGKVFKARDLKNGGRFVALKRVRVQT |
| *Cygnus olor* | QYECVAEIGEGAYGKVFKARDLKNGGRFVALKRVRVQT |
| *Dromaius novaehollandiae* | QYECVAEIGEGAYGKVFKARDLKNGGRFVALKRVRVQT |
| *Dryobates pubescens* | QYECVAEIGEGAYGKVFKARDLKNGGRFVALKRVRVQT |
| *Egretta garzetta* | QYECVAEIGEGAYGKVFKARDLKNGGRFVALKRVRVQT |
| *Empidonax traillii* | QYEYVAEIGEGAYGKVFKARDLKNGGRFVALKRVRVQT |
| *Erythrura gouldiae* | RYECVAEIGEGAYGKVFKARDLKNGGRFVALKRVRMQT |
| *Eudyptes chrysocome* | QYECVAEIGEGAYGKVFKARDLKNGGRFVALKRVRVQT |
| *Eudyptes chrysolophus* | QYECVAEIGEGAYGKVFKARDLKNGGRFVALKRVRVQT |
| *Eudyptes filholi* | QYECVAEIGEGAYGKVFKARDLKNGGRFVALKRVRVQT |
| *Eudyptes moseleyi* | QYECVAEIGEGAYGKVFKARDLKNGGRFVALKRVRVQT |
| *Eudyptes pachyrhynchus* | QYECVAEIGEGAYGKVFKARDLKNGGRFVALKRVRVQT |
| *Eudyptes robustus* | QYECVAEIGEGAYGKVFKARDLKNGGRFVALKRVRVQT |
| *Eudyptes schlegeli* | QYECVAEIGEGAYGKVFKARDLKNGGRFVALKRVRVQT |
| *Eudyptes sclateri* | QYECVAEIGEGAYGKVFKARDLKNGGRFVALKRVRVQT |
| *Eudyptula albosignata* | QYECVAEIGEGAYGKVFKARDLKNGGRFVALKRVRVQT |
| *Eudyptula minor* | QYECVAEIGEGAYGKVFKARDLKNGGRFVALKRVRVQT |
| *Eudyptula novaehollandiae* | QYECVAEIGEGAYGKVFKARDLKNGGRFVALKRVRVQT |
| *Eurypyga helias* | QYECVAEIGEGAYGKVFKARDLKNGGRFVALKRVRVQT |
| *Falco cherrug* | QYECVAEIGEGAYGKVFKARDLKNGGRFVALKRVRVQT |
| *Falco naumanni* | QYECVAEIGEGAYGKVFKARDLKNGGRFVALKRVRVQT |
| *Falco peregrinus* | QYECVAEIGEGAYGKVFKARDLKNGGRFVALKRVRVQT |
| *Falco rusticolus* | QYECVAEIGEGAYGKVFKARDLKNGGRFVALKRVRVQT |
| *Falco tinnunculus* | QYECVAEIGEGAYGKVFKARDLKNGGRFVALKRVRVQT |
| *Ficedula albicollis* | QYECVAEIGEGAYGKVFKARDLKNGGRFVALKRVRVQT |
| *Gallus gallus* | QYECVAEIGEGAYGKVFKARDLKNGGRFVALKRVRVQT |
| *Gavia stellata* | QYECVAEIGEGAYGKVFKARDLKNGGRFVALKRVRVQT |
| *Geospiza fortis* | QYECVAEIGEGAYGKVFKARDLKNGGRFVALKRVRVQT |
| *Haliaeetus albicilla* | QYECVAEIGEGAYGKVFKARDLKNGGRFVALKRVRVQT |
| *Haliaeetus leucocephalus* | QYECVAEIGEGAYGKVFKARDLKNGGRFVALKRVRVQT |
| *Hirundo rustica* | QYECVAEIGEGAYGKVFKARDLKNGGRFVALKRVRVQT |
| *Hypotaenidia okinawae* | QYECVAEIGEGAYGKVFKARDLKNGGRFVALKRVRVQT |
| *Junco hyemalis* | QYECVAEIGEGAYGKVFKARDLKNGGRFVALKRVRVQT |
| *Lagopus leucura* | QYECVAEIGEGAYGKVFKARDLKNGGRFVALKRVRVQT |
| *Lepidothrix coronata* | QYEYVAEIGEGAYGKVFKARDLKNGGRFVALKRVRVQT |
| *Leptosomus discolor* | QYECVAEIGEGAYGKVFKARDLKNGGRFVALKRVRVQT |
| *Lonchura striata domestica* | RYECVAEIGEGAYGKVFKARDLKNGGRFVALKRVRMQT |
| *Malurus cyaneus samueli* | QYECVAEIGEGAYGKVFKARDLKNGGRFVALKRVRVQT |
| *Manacus vitellinus* | QYEYVAEIGEGAYGKVFKARDLKNGGRFVALKRVRVQT |
| *Megadyptes antipodes antipodes* | QYECVAEIGEGAYGKVFKARDLKNGGRFVALKRVRVQT |
| *Meleagris gallopavo* 1 | QYECVAEIGEGAYGKVFKARDLKNGGRFVALKRVRVQT |
| *Meleagris gallopavo* 2 | QYECVAEIGEGAYGKVFKARDLKNGGRFVALKRVRVQT |
| *Melopsittacus undulatus* | QYECVAEIGEGAYGKVFKARDLKNGGRFVALKRVRVQT |
| *Melospiza melodia maxima* | QYECVAEIGEGAYGKVFKARDLKNGGRFVALKRVRVQT |
| *Molothrus ater* | QYECVAEIGEGAYGKVFKARDLKNGGRFVALKRVRVQT |
| *Merops nubicus* | QYECVAEIGEGAYGKVFKARDLKNGGRFVALKRVRVQT |
| *Mesitornis unicolor* | QYECVAEIGEGAYGKVFKARDLKNGGRFVALKRVRVQT |
| *Motacilla alba alba* | QYECVAEIGEGAYGKVFKARDLKNGGRFVALKRVRVQT |
| *Neopelma chrysocephalum* | QYEYVAEIGEGAYGKVFKARDLKNGGRFVALKRVRVQT |
| *Nestor notabilis* | QYECVAEIGEGAYGKVFKARDLKNGGRFVALKRVRVQT |
| *Nipponia nippon* | QYECVAEIGEGAYGKVFKARDLKNGGRFVALKRVRVQT |
| *Nothoprocta perdicaria* | QYECVAEIGEGAYGKVFKARDLKNGGRFVALKRVRVQT |
| *Numida meleagris* | QYECVAEIGEGAYGKVFKARDLKNGGRFVALKRVRVQT |
| *Onychostruthus taczanowskii* | QYECVAEIGEGAYGKVFKARDLKNGGRFVALKRVRVQT |
| *Opisthocomus hoazin* | QYECVAEIGEGAYGKVFKARDLKNGGRFVALKRVRVQT |
| *Otus sunia* | QYECVAEIGEGAYGKVFKARDLKNGGRFVALKRVRVQT |
| *Oxyura jamaicensis* | QYECVAEIGEGAYGKVFKARDLKNGGRFVALKRVRVQT |
| *Parus major* | QYECVAEIGEGAYGKVFKARDLKNGGRFVALKRVRVQT |
| *Passer montanus* | QYECVAEIGEGAYGKVFKARDLKNGGRFVALKRVRVQT |
| *Patagioenas fasciata monilis* | QYECVAEIGEGAYGKVFKARDLKNGGRFVALKRVRVQT |
| *Pavo cristatus* | QYECVAEIGEGAYGKVFKARDLKNGGRFVALKRVRVQT |
| *Pelecanus crispus* | QYECVAEIGEGAYGKVFKARDLKNGGRFVALKRVRVQT |
| *Phaethon lepturus* | QYECVAEIGEGAYGKVFKARDLKNGGRFVALKRVRVQT |
| *Phalacrocorax carbo* | QYECVAEIGEGAYGKVFKARDLKNGGRFVALKRVRVQT |
| *Phasianus colchicus* | QYECVAEIGEGAYGKVFKARDLKNGGRFVALKRVRVQT |
| *Pipra filicauda* | QYEYVAEIGEGAYGKVFKARDLKNGGRFVALKRVRVQT |
| *Pseudopodoces humilis* | QYECVAEIGEGAYGKVFKARDLKNGGRFVALKRVRVQT |
| *Pterocles gutturalis* | QYECVAEIGEGAYGKVFKARDLKNGGRFVALKRVRVQT |
| *Pygoscelis adeliae* | QYECVAEIGEGAYGKVFKARDLKNGGRFVALKRVRVQT |
| *Pygoscelis antarcticus* | QYECVAEIGEGAYGKVFKARDLKNGGRFVALKRVRVQT |
| *Pygoscelis papua* | QYECVAEIGEGAYGKVFKARDLKNGGRFVALKRVRVQT |
| *Pyrgilauda ruficollis* | QYECVAEIGEGAYGKVFKARDLKNGGRFVALKRVRVQT |
| *Serinus canaria* | QYECVAEIGEGAYGKVFKARDLKNGGRFVALKRVRVQT |
| *Spheniscus demersus* | QYECVAEIGEGAYGKVFKARDLKNGGRFVALKRVRVQT |
| *Spheniscus humboldti* | QYECVAEIGEGAYGKVFKARDLKNGGRFVALKRVRVQT |
| *Spheniscus magellanicus* | QYECVAEIGEGAYGKVFKARDLKNGGRFVALKRVRVQT |
| *Spheniscus mendiculus* | QYECVAEIGEGAYGKVFKARDLKNGGRFVALKRVRVQT |
| *Strigops habroptila* | QYECVAEIGEGAYGKVFKARDLKNGGRFVALKRVRVQT |
| *Strix occidentalis caurina* | QYECVAEIGEGAYGKVFKARDLKNGGRFVALKRVRVQT |
| *Struthio camelus australis* | QYECVAEIGEGAYGKVFKARDL---------------- |
| *Sturnus vulgaris* | QYECVAEIGEGAYGKVFKARDLKNGGRFVALKRVRVQT |
| *Taeniopygia guttata* | RYECVAEIGEGAYGKVFKARDLKNGGRFVALKRVRMQT |
| *Tauraco erythrolophus* | QYECVAEIGEGAYGKVFKARDLKNGGRFVALKRVRVQT |
| *Tinamus guttatus* | QYECVAEIGEGAYGKVFKARDLKNGGRFVALKRVRVQT |
| *Turdus rufiventris* | QYECVAEIGEGAYGKVFKARDLKNGGRFVALKRVRVQT |
| *Turnix velox* | QYECVAEIGEGAYGKVFKARDLKNGGRFVALKRVRVQT |
| *Tyto alba* | QYECVAEIGEGAYGKVFKARDLKNGGRFVALKRVRVQT |
| *Zonotrichia albicollis* | QYECVAEIGEGAYGKVFKARDLKNGGRFVALKRVRVQT |
| *Zosterops borbonicus* | QYECVAEIGEGAYGKVFKARDLKNGGRFVALKRVRVQT |
| *Zosterops lateralis melanops* | QYECVAEIGEGAYGKVFKARDLKNGGRFVALKRVRVQT |

**reptile**

| | |
|---|---|
| *Alligator mississippiensis* | QYECVAEIGEGAYGKVFKARDLKNGGRFVALKRVRVQT |
| *Alligator sinensis* | QYECVAEIGEGAYGKVFKARDLKNGGRFVALKRVRVQT |
| *Anolis carolinensis* | QYECVAEIGEGAYGKVFKARDLKNGGRFVALKRVRVQT |
| *Chelonia mydas* | QYECVAEIGEGAYGKVFKARDLKNGGRFVALKRVRVQT |
| *Chelonoidis abingdonii* | QYECVAEIGEGAYGKVFKARDLKNGGRFVALKRVRVQT |
| *Chelydra serpentina* | QYECVAEIGEGAYGKVFKARDLKNGGRFVALKRVRVQT |
| *Chrysemys picta bellii* | QYECVAEIGEGAYGKVFKARDLKHGGRFVALKRVRVQT |
| *Crocodylus porosus* | QYECVAEIGEGAYGKVFKARDLKNGGRFVALKRVRVQT |
| *Crotalus adamanteus* | QYECVAEIGEGAYGKVFKARDLKNGGRFVALKRVRVQT |
| *Crotalus tigris* | QYECVAEIGEGAYGKVFKARDLKNGGRFVALKRVRVQT |
| *Dermochelys coriacea* | QYECVAEIGEGAYGKVFKARDLKNGGRFVALKRVRVQT |
| *Gavialis gangeticus* | QYECVAEIGEGAYGKVFKARDLKNGGRFVALKRVRVQT |
| *Gekko japonicus* | QYECVAEIGEGAYGKVFKARDLKNGGRFVALKRVRVQT |
| *Gopherus agassizii* | QYECVAEIGEGAYGKVFKARDLKNGGRFVALKRVRVQT |
| *Gopherus evgoodei* | QYECVAEIGEGAYGKVFKARDLKNGGRFVALKRVRVQT |
| *Hypsiglena* sp. JMG-2014 | QYEPVAEIGIGAYGTVFKARDLQSG-KFVALKNVRVQN |
| *Lacerta agilis* | QYECVAEIGEGAYGKVFKARDLKNGGRFVALKRVRVQT |
| *Laticauda laticaudata* | QYECVAEIGEGAYGKVFKARDLKNGGRFVALKRVRVQT |
| *Mauremys mutica* | QYECVAEIGEGAYGKVFKARDLKNGGRFVALKRVRVQT |
| *Mauremys reevesii* | QYECVAEIGEGAYGKVFKARDLKNGGRFVALKRVRVQT |
| *Naja naja* | QYECVAEIGEGAYGKVFKARDLKNGGRFVALKRVRVQT |
| *Notechis scutatus* | QYECVAEIGEGAYGKVFKARDLKNGGRFVALKRVRVQT |
| *Ophiophagus hannah* | QYECVAEIGEGAYGKVFKARDLKNGGRFVALKRVRVQT |
| *Pantherophis guttatus* | QYECVAEIGEGAYGKVFKARDLKNGGRFVALKRVRVQT |
| *Pelodiscus sinensis* | QYECVAEIGEGAYGKVFKARDLKNGGRFVALKRVRVQT |
| *Pelusios castaneus* | QYECVAEIGEGAYGKVFKARDLKNGGRFVALKRVRVQT |
| *Podarcis muralis* | QYECVAEIGEGAYGKVFKARDLKNGGRFVALKRVRVQT |
| *Pogona vitticeps* | QYECVAEIGEGAYGKVFKARDLKNGGRFVALKRVRVQT |
| *Protobothrops mucrosquamatus* | QYECVAEIGEGAYGKVFKARDLKNGGRFVALKRVRVQT |
| *Pseudonaja textilis* | QYECVAEIGEGAYGKVFKARDLKNGGRFVALKRVRVQT |
| *Python bivittatus* | QYECVAEIGEGAYGKVFKARDLKNGGRFVALKRVRVQT |
| *Salvator merianae* | QYECVAEIGEGAYGKVFKARDLKNGGRFVALKRVRVQT |
| *Sceloporus undulatus* | QYECVAEIGEGAYGKVFKARDLKNGGRFVALKRVRVQT |
| *Sphenodon punctatus* | QYECVAEIGEGAYGKVFKARDLKNGGRFVALKRVRVQT |
| *Terrapene carolina triunguis* | QYECVAEIGEGAYGKVFKARDLKHGGRFVALKRVRVQT |
| *Thamnophis elegans* | QYECVAEIGEGAYGKVFKARDLKNGGRFVALKRVRVQT |
| *Thamnophis sirtalis* | QYECVAEIGEGAYGKVFKARDLKNGGRFVALKRVRVQT |
| *Trachemys scripta elegans* | QYECVAEIGEGAYGKVFKARDLKHGGRFVALKRVRVQT |
| *Varanus komodoensis* | QYECVAEIGEGAYGKVFKARDLKNGGRFVALKRVRVQT |
| *Zootoca vivipara* | QYECVAEIGEGAYGKVFKARDLKNGGRFVALKRVRVQT |

**amphibian**

| | |
|---|---|
| *Ambystoma andersoni* | QYECVAEIGQGAYGKVFKARDLKNGGRFVALKQVRVLT |
| *Ambystoma maculatum* | QYECVAEIGQGAYGKVFKARDLKNGGRFVALKQVRVQT |
| *Ambystoma mexicanum* | QYECVAEIGQGAYGKVFKARDLKNGGRFVALKQVRVQT |
| *Ambystoma velasci* | QYECVAEIGQGAYGKVFKARDLKNGGRFVALKQVRVQT |
| *Bufo bufo* | QYECVAEIGEGAYGKVFKARDLKNGGRFVALKRVRVQT |
| *Bufo gargarizans* | QYECVAEIGEGAYGKVFKARDLKNGGRFVALKRVRVQT |
| *Engystomops pustulosus* | QYECVAEIGEGAYGKVFKARDLKNGGRFVALKRVRVQT |
| *Geotrypetes seraphini* | QYECVAEIGEGAYGKVFKARDLKNGGRFVALKRMRVQT |
| *Hymenochirus boettgeri* | QYECVAEIGEGAYGKVFKARDLKNGGRFVALKRVRVQT |
| *Leptobrachium leishanense* | QYECVAEIGEGAYGKVFKARDLKNGGRFVALKRVRVQT |
| *Microcaecilia unicolor* | QYECVAEIGEGAYGKVFKARDLKNGGRFVALKRVRVQT |
| *Nanorana parkeri* | QYECVAEIGEGAYGKVYKARDLKNGGRFVALKRVRVQT |
| *Rana temporaria* | QYECVAEIGEGAYGKVYKARDLKNGGRFVALKRVRVQT |
| *Rhinatrema bivittatum* | QYECVAEIGEGAYGKVFKARDLKNGGRFVALKRVRVQT |
| *Xenopus laevis* 1 | QYECVAEIGEGAYGKVFKARDLKNGGRFVALKRVRVQT |
| *Xenopus laevis* 2 | QYECVAEIGEGAYGKVFKARDLKNGGRFVALKRVRVQT |
| *Xenopus tropicalis* | QYECVAEIGEGAYGKVFKARDLKNGGRFVALKRVRVQT |

**Figure S5e**

**Cdk4**

| | |
|---|---|
| Homo sapiens | RYEPVAEIGVGAYG--TVYKARD-PHSGHFVAL-KSVRVPN |

**Cdk6**

**bony fish**

| | |
|---|---|
| *Acanthochromis polyacanthus* | QYERVAEIGEGAYG--KVYKARDLKNGGRFVAL-KRVRVQT |
| *Acanthopagrus latus* | QYEPVAEIGEGAYG--KVYKARDLKNGGRFVAL-KRVRVQT |
| *Acipenser ruthenus* 1 | QYEPVAEIGEGAYG--KVYKARDLKNGGRFVAL-KRVRVQT |
| *Acipenser ruthenus* 2 | QYEPVAEIGEGAYG--KVYKARDLKNGGRFVAL-KRVRVQT |
| *Albula glossodonta* | QYEPVAEIGEGAYG--KVYKARDLKNGGRFVAL-KRVRVQT |
| *Alosa alosa* | SYEPVAEIGEGAYG--KVYKARDLNNGGRFVAL-KRVRVQT |
| *Alosa sapidissima* | SYEPVAEIGEGAYG--KVYKARDLNNGGRFVAL-KRVRVQT |
| *Ameiurus melas* | RYEPVAEIGEGAYG--KVYKARDLKNG-RFVAL-KRVRVQT |
| *Amphilophus citrinellus* | QYEPVAEIGEGAYG--KVYKARDLKNGGRFVAL-KRVRVQT |
| *Amphiprion ocellaris* | QYEPVAEIGEGAYG--KVYKARDLKNGGRFVAL-KRVRVQT |
| *Amphiprion percula* | QYEPVAEIGEGAYG--KVYKARDLKNGGRFVAL-KRVRVQT |
| *Anabarilius grahami* | QYEPVAEIGEGAYG--KVYKARDLKNGGRFVAL-KRVRVQT |
| *Anabas testudineus* | QYEPVAEIGEGAYG--KVYKARDLKNGGRFVAL-KRVRVQT |
| *Anarrhichthys ocellatus* | QYEPVAEIGEGAYG--KVYKARDLKNGGRFVAL-KRVRVQT |
| *Anguilla anguilla* | QYVPVAEIGEGAYG--KVYKARDLKNGGRFVAL-KRVRVQT |
| *Archocentrus centrarchus* | QYEPVAEIGEGAYG--KVYKARDLKNGGRFVAL-KRVRVQT |
| *Astatotilapia calliptera* | QYEPVAEIGEGAYG--KVYKARDLKNGGRFVAL-KRVRVQT |
| *Astyanax mexicanus* | QYEPVAEIGEGAYG--KVYKARDLKNGGRFVAL-KRVRVQT |
| *Austrofundulus limnaeus* | QYEPVAEIGEGAYG--KVYKARDLKNGGRFVAL-KRVRVQT |
| *Bagarius yarrelli* | QYEPVAEIGEGAYG--KVYKARDLKNNGRFVAL-KRVRVQT |
| *Betta splendens* | QYEPVAEIGEGAYG--KVYKARDLKNGGRFVAL-KRVRVQT |
| *Caranx melampygus* | QYEPVAEIGEGAYG--KVYKARDLKNGGRFVAL-KRVRVQT |
| *Carassius auratus* 1 | QYEPVAEIGEGAYG--KVYKARDLKNGGRFVAL-KRVRVQT |
| *Carassius auratus* 2 | QYEPVAEIGEGAYG--KVYKARDLKNGGRFVAL-KRVRVQT |
| *Channa argus* | QYEPVAEIGEGAYG--KVYKARDLKNGGRFVAL-KRVRVQT |
| *Chanos chanos* | QYEPVAEIGEGAYG--KVYKARDLKNGGRFVAL-KRVRVQT |
| *Cheilinus undulatus* | QYEPVAEIGEGAYG--KVYKARDLKNGGRFVAL-KRVRVQT |
| *Chelmon rostratus* | QYEAVAEIGEGAYG--KVYKARDLKNGGRFVAL-KRVRVQT |
| *Clarias magur* | QYEPVAEIGEGAYG--KVYKARDLKNG-RFVAL-KRVRVQT |
| *Clupea harengus* | SYEPVAEIGEGAYG--KVYKARDLNNGGRFVAL-KRVRVQT |
| *Collichthys lucidus* | QYEPVAEIGEGAYG--KVYKARDLKNGGRFVAL-KRVRVQT |
| *Colossoma macropomum* | QYEPVAEIGEGAYG--KVYKARDLKNGGRFVAL-KRVRVQT |
| *Coregonus clupeaformis* 1 | RYEPVAEIGEGAYG--KVYKARDLKNGGRFVAL-KRVRVQT |
| *Coregonus clupeaformis* 2 | QYEPVAEIGEGAYG--KVYKARDLKNEGRFVAL-KRVRVQT |
| *Coregonus lavaretus* | RYEPVAEIGEGAYG--KVYKARDLKNGGRFVAL-KRVRVQT |
| *Coryphaenoides rupestris* | DYEPVAEIGEGAYG--KVYKARDLKNGGRFVAL-KRVRVQT |
| *Cottoperca gobio* | QYEPVAEIGEGAYG--KVYKARDLKNGGRFVAL-KRVRVQT |
| *Cyclopterus lumpus* | QYEPVAEIGEGAYG--KVYKARDLKNGGRFVAL-KRVRVQT |
| *Cynoglossus semilaevis* | QYEQVAEIGEGAYG--KVYKARDLKNGGRFVAL-KRVRVLT |
| *Cyprinodon tularosa* | QYEPVAEIGEGAYG--KVYKARDLKNGGRFVAL-KRVRVQT |
| *Cyprinodon variegatus* | QYEPVAEIGEGAYG--KVYKARDLKNGGRFVAL-KRVRVQT |
| *Cyprinus carpio* 1 | QYEPVAEIGEGAYG--KVYKARDLKNGGRFVAL-KRVRVQT |
| *Cyprinus carpio* 2 | QYEPVAEIGEGAYG--KVYKARDLKNGGRFVAL-KRVRVQT |
| *Danio rerio* | QYEPVAEIGEGAYG--KVYKARDLKNGGRFVAL-KRVRVQT |
| *Danionella translucida* | QYEPVAEIGEGAYG--KVYKARDLKNGGRFVAL-KRVRVQT |
| *Denticeps clupeoides* | LYEPVAEIGEGAYG--KVYKARDLKNGGRFVAL-KRVRVQT |
| *Dicentrarchus labrax* | QYEPVAEIGEGAYG--KVYKARDLKNGGRFVAL-KRVRVQT |
| *Echeneis naucrates* | QYEAVAEIGEGAYG--KVYKARDLKNGGRFVAL-KRVRVQT |
| *Electrophorus electricus* | QYEPVAEIGEGAYG--KVYKARDLKNGGRFVAL-KRVRVQT |
| *Epinephelus coioides* | QYEPVAEIGEGAYG--KVYKARDLKNGGRFVAL-KRVRVQT |
| *Epinephelus lanceolatus* | QYEPVAEIGEGAYG--KVYKARDLKNGGRFVAL-KRVRVQT |
| *Erpetoichthys calabaricus* | QYEPVAEIGEGAYG--KVYKARDLNNGGRFVAL-KRVRVLT |
| *Esox lucius* | LYEPVAEIGEGAYG--KVYKARDLKNGGRFVAL-KKVRVQT |
| *Etheostoma cragini* | QYEPVAEIGEGAYG--KVYKARDLNNGGRFVAL-KRVRVQT |
| *Etheostoma spectabile* | QYEPVAEIGEGAYG--KVYKARDLNNGGRFVAL-KRVRVQT |
| *Fundulus heteroclitus* | QYEPVAEIGEGAYG--KVYKARDLKNGGRFVAL-KRVRVQT |
| *Gadus morhua* | DYEPVAEIGEGAYG--KVYKARDLKNGGRFVAL-KRVRVQT |
| *Gambusia affinis* | QYEPVAEIGEGAYG--KVYKARDLKNGGRFVAL-KRVRVQT |
| *Gasterosteus aculeatus aculeatus* | QYEPVAEIGEGAYG--KVYKARDLKNGGRFVAL-KRVRVQT |
| *Girardinichthys multiradiatus* | QYEPVAEIGEGAYG--KVYKARDLKNGGRFVAL-KRVRVQT |
| *Gouania willdenowi* | QYEPVAEIGEGAYG--KVYKARDLKNGGRFVAL-KKVRVQT |
| *Gymnodraco acuticeps* | QYEPVAEIGEGAYG--KXYKARDMKNGGRFVAL-KRVRVQT |
| *Haplochromis burtoni* | QYEPVAEIGEGAYG--KVYKARDLKNGGRFVAL-KRVRVQT |
| *Hippocampus comes* | QYEPVAEIGEGAYG--KVYKARDLKNGGRFVAL-KRVRVQT |
| *Hippoglossus hippoglossus* | RYEPVAEIGEGAYG--KVYKARDLKNGGRFVAL-KRVRVQT |
| *Hippoglossus stenolepis* | RYEPVAEIGEGAYG--KVYKARDLKNGGRFVAL-KRVRVQT |
| *Hucho hucho* 1 | ATSPVAEIGEGAYG--KVYKARDLNNAGRFVAL-KRVRVQT |
| *Hucho hucho* 2 | QYEPVAEIGEGAYG--KVYKARDLKNEGRFVAL-KRVRVQT |
| *Iconisemion striatum* | QYEAVAEIGEGAYG--KVYKARDLKNGGRFVAL-KRVRVQT |
| *Ictalurus punctatus* | RYEPVAEIGEGAYG--KVYKARDLKNG-RFVAL-KRVRVQT |
| *Kryptolebias marmoratus* | QYEPVAEIGEGAYG--KVYKARDLKNGGRFVAL-KRVRVQT |
| *Labeo rohita* | QYEPVAEIGEGAYG--KVYKARDLKNGGRFVAL-KRVRVQT |
| *Labrus bergylta* | RYEPVAEIGEGAYG--KVYKARDLKNGGRFVAL-KRVRVQT |
| *Larimichthys crocea* | QYEPVAEIGEGAYG--KVYKARDLKNGGRFVAL-KRVRVQT |
| *Lates calcarifer* | QYEPVAEIGEGAYG--KVYKARDLKNGGRFVAL-KRVRVQT |
| *Latimeria chalumnae* | LYEPVAEIGEGAYG--KVYKARDLKNGGRFVAL-KKVRVQT |
| *Lepisosteus oculatus* | QYEPVAEIGEGAYG--KVYKARDLKNGGRFVAL-KRVRVQT |
| *Liparis tanakae* | QYEPVAEIGEGAYG--KVYKARDLKNGGRFVAL-KRVRVQT |
| *Lucifuga dentata* | QYEPVAEIGEGAYG--KVYKARDLKNGGRFVAL-KRVRVQT |
| *Mastacembelus armatus* | QYEPVAEIGEGAYG--KVYKARDLKNGGRFVAL-KRVRVQT |
| *Maylandia zebra* | QYEPVAEIGEGAYG--KVYKARDLKNGGRFVAL-KRVRVQT |
| *Megalobrama amblycephala* | QYEPVAEIGEGAYG--KVYKARDLKNGGRFVAL-KRVRVQT |
| *Megalops atlanticus* | QYEPVAEIGEGAYG--KVYKARDLKNGGRFVAL-KRVRVQT |
| *Megalops cyprinoides* | QYEPVAEIGEGAYG--KVYKARDLKNGGRFVAL-KRVRVQT |
| *Melanotaenia boesemani* | LYETVAEIGEGAYG--KVYKARDLKNGGRFVAL-KRVRVQT |
| *Micropterus dolomieu* | QYEPVAEIGEGAYG--KVYKARDLKNGGRFVAL-KRVRVQT |
| *Micropterus salmoides* | QYEPVAEIGEGAYG--KVYKARDLKNGGRFVAL-KRVRVQT |
| *Mola mola* | QYEHVAEIGEGAYG--KVYKARDLKNGGRFVAL-KRVRVQT |
| *Monopterus albus* | QYEPVAEIGEGAYG--KVYKARDLKNGGRFVAL-KRVRVQS |
| *Morone saxatilis* | QYEPVAEIGEGAYG--KVYKARDLKNGGRFVAL-KRVRVQT |
| *Mugil cephalus* | QYEPVAEIGEGAYG--KVYKARDLKNGGRFVAL-KRVRVQT |
| *Myripristis murdjan* | QYEPVAEIGEGAYG--KVYKARDLKNGGRFVAL-KRVRVQT |
| *Nematolebias whitei* | QYEPVAEIGEGAYG--KVYKARDLKNGGRFVAL-KRVRVQT |
| *Neolamprologus brichardi* | QYEPVAEIGEGAYG--KVYKARDLKNGGRFVAL-KRVRVQT |
| *Nibea albiflora* | QYEPVAEIGEGAYG--KVYKARDLKNGGRFVAL-KRVRVQT |
| *Nothobranchius furzeri* | QYEAVAEIGEGAYG--KVYKARDLKNGGRFVAL-KRVRVQT |
| *Nothobranchius kadleci* | QYEAVAEIGEGAYG--KVYKARDLKNGGRFVAL-KRVRVQT |
| *Nothobranchius korthausae* | QYEAVAEIGEGAYG--KVYKARDLKNGGRFVAL-KRVRVQT |
| *Nothobranchius kuhntae* | QYEAVAEIGEGAYG--KVYKARDLKNGGRFVAL-KRVRVQT |
| *Nothobranchius pienaari* | QYEAVAEIGEGAYG--KVYKARDLKNGGRFVAL-KRVRVQT |
| *Nothobranchius rachovii* | QYEAVAEIGEGAYG--KVYKARDLKNGGRFVAL-KRVRVQT |
| *Notolabrus celidotus* | QYEPVAEIGEGAYG--KVYKARDLKNGGRFVAL-KRVRVQT |
| *Notothenia coriiceps* | QYEPVAEIGEGAYG--KVYKARDLKNGGRFVAL-KRVRVQT |
| *Oncorhynchus gorbuscha* 1 | RYEPVAEIGEGAYG--KVYKARDLKNAGRFVAL-KRVRVQT |
| *Oncorhynchus gorbuscha* 2 | QYEPVAEIGEGAYG--KVYKARDLKNEGRFVAL-KRVRVQT |
| *Oncorhynchus keta* 1 | RYEPVAEIGEGAYG--KVYKARDLKNAGRFVAL-KRVRVQT |
| *Oncorhynchus keta* 2 | QYEPVAEIGEGAYG--KVYKARDLKNEGRFVAL-KRVRVQT |
| *Oncorhynchus kisutch* 1 | QYEPVAEIGEGAYG--KVYKARDLKNEGRFVAL-KRVRVQT |
| *Oncorhynchus kisutch* 2 | RYEPVAEIGEGAYG--KVYKARDLKNAGRFVAL-KRVRVQT |
| *Oncorhynchus mykiss* 1 | RYEPVAEIGEGAYG--KVYKARDLKNAGRFVAL-KRVRVQT |
| *Oncorhynchus mykiss* 2 | QYEPVAEIGEGAYG--KVYKARDLKNEGRFVAL-KRVRVQT |
| *Oncorhynchus nerka* 1 | RYEPVAEIGEGAYG--KVYKARDLKNAGRFVAL-KRVRVQT |
| *Oncorhynchus nerka* 2 | QYEPVAEIGEGAYG--KVYKARDLKNEGRFVAL-KRVRVQT |
| *Oncorhynchus tshawytscha* 1 | RYEPVAEIGEGAYG--KVYKARDLKNAGRFVAL-KRVRVQT |
| *Oncorhynchus tshawytscha* 2 | QYEPVAEIGEGAYG--KVYKARDLKNEGRFVAL-KRVRVQT |
| *Onychostoma macrolepis* | QYEPVAEIGEGAYG--KVYKARDLKNGGRFVAL-KRVRVQT |
| *Oreochromis aureus* | QYEPVAEIGEGAYG--KVYKARDLKNGGRFVAL-KRVRVQT |
| *Oreochromis niloticus* | QYEPVAEIGEGAYG--KVYKARDLKNGGRFVAL-KRVRVQT |
| *Oryzias javanicus* | QYEPVAEIGEGAYG--KVYKARDLKNEGRFVAL-KRVRVQT |
| *Oryzias latipes* | QYEPVAEIGEGAYG--KVYKARDLKNEGRFVAL-KRVRVQT |
| *Oryzias melastigma* | QYEPVAEIGEGAYG--KVYKARDLKNEGRFVAL-KRVRVQT |
| *Oryzias sinensis* | QYEPVAEIGEGAYG--KVYKARDLKNEGRFVAL-KRVRVQT |
| *Pangasianodon hypophthalmus* | QYEPVAEIGEGAYG--KVYKARDLKNG-RFVAL-KRVRVQT |
| *Paralichthys olivaceus* | QYEPVAEIGEGAYG--KVYKARDLKNGGRFVAL-KRVRVQT |
| *Parambassis ranga* | QYEPVAEIGEGAYG--KVYKARDLKNGGRFVAL-KRVRVQT |
| *Paramormyrops kingsleyae* | QYEPVAEIGEGAYG--KVYKARDLKNG-RFVAL-KRVRVQT |
| *Perca flavescens* | QYEPVAEIGEGAYG--KVYKARDLKNGGRFVAL-KRVRVQT |
| *Perca fluviatilis* | QYEPVAEIGEGAYG--KVYKARDLKNGGRFVAL-KRVRVQT |
| *Pimephales promelas* | QYEPVAEIGEGAYG--KVYKARDLKNGGRFVAL-KRVRVQT |
| *Plectropomus leopardus* | QYEPVAEIGEGAYG--KVYKARDLKNGGRFVAL-KRVRVQT |
| *Poecilia formosa* | QYEPVAEIGEGAYG--KVYKARDLKNGGRFVAL-KRVRVQT |
| *Poecilia latipinna* | QYEPVAEIGEGAYG--KVYKARDLKNGGRFVAL-KRVRVQT |
| *Poecilia mexicana* | QYEPVAEIGEGAYG--KVYKARDLKNGGRFVAL-KRVRVQT |
| *Poecilia reticulata* | QYEPVAEIGEGAYG--KVYKARDLKNGGRFVAL-KRVRVQT |
| *Poeciliopsis prolifica* | QYEPVAEIGEGAYG--KVYKARDLKNGGRFVAL-KRVRVQT |
| *Polyodon spathula* 1 | QYEPVAEIGEGAYG--KVYKARDLKNGGRFVAL-KRVRVQT |
| *Polyodon spathula* 2 | QYEPVAEIGEGAYG--KVYKARDLKNGGRFVAL-KRVRVQT |
| *Polypterus senegalus* | QYEPVAEIGEGAYG--KVYKARDLNNGGRFVAL-KRVRVLT |
| *Protopterus annectens* | QYEYVAEIGEGAYG--KVFKARDLKNGGRFVAL-KRVRVQT |
| *Pseudochaenichthys georgianus* | QYEPVAEIGEGAYG--KVYKARDLKNGGRFVAL-KRVRVQT |
| *Pundamilia nyererei* | QYEPVAEIGEGAYG--KVYKARDLKNGGRFVAL-KRVRVQT |
| *Pungitius pungitius* | QYEPVAEIGEGAYG--KVYKARDLKNGGRFVAL-KRVRVQT |
| *Puntigrus tetrazona* | QYEPVAEIGEGAYG--KVYKARDLKNGGRFVAL-KRVRVQT |
| *Pygocentrus nattereri* | QYEPVAEIGEGAYG--KVYKARDLKNGGRFVAL-KRVRVQT |
| *Salarias fasciatus* | QYEPVAEIGEGAYG--KVYKARDLKNGGRFVAL-KRVRVQT |
| *Salmo salar* 1 | RYEPVAEIGEGAYG--KVYKARDLNNAGRFVAL-KRVRVQT |
| *Salmo salar* 2 | QYEPVAEIGEGAYG--KVYKARDLKNEGRFVAL-KRVRVQT |
| *Salmo trutta* 1 | QYEPVAEIGEGAYG--KVYKARDLKNEGRFVAL-KRVRVQT |
| *Salmo trutta* 2 | RYEPVAEIGEGAYG--KVYKARDLNNAGRFVAL-KRVRVQT |
| *Salvelinus alpinus* 1 | QYEPVAEIGEGAYG--KVYKARDLKNEGRFVAL-KRVRVQT |
| *Salvelinus alpinus* 2 | RYEPVAEIGEGAYG--KVYKARDLKNAGRFVAL-KRVRVQT |
| *Salvelinus namaycush* 1 | QYEPVAEIGEGAYG--KVYKARDLKNEGRFVAL-KRVRVQT |
| *Salvelinus namaycush* 2 | RYEPVAEIGEGAYG--KVYKARDLKNAGRFVAL-KRVRVQT |
| *Sander lucioperca* | QYEPVAEIGEGAYG--KVYKARDLKNGGRFVAL-KRVRVQT |
| *Scatophagus argus* | QYEPVAEIGEGAYG--KVYKARDLKNGGRFVAL-KRVRVQT |
| *Scleropages formosus* | QYEPVAEIGEGAYG--KVYKARDLKNG-RFVAL-KRVRVQT |
| *Scophthalmus maximus* | RYEPVAEIGEGAYG--KVYKARDLKNGGRFVAL-KRVRVQT |
| *Sebastes umbrosus* | QYEPVAEIGEGAYG--KVYKARDLKNGGRFVAL-KRVRVQT |
| *Seriola dumerili* | QYEPVAEIGEGAYG--KVYKARDLKNGGRFVAL-KRVRVQT |
| *Seriola lalandi dorsalis* | QYEPVAEIGEGAYG--KVYKARDLKNGGRFVAL-KRVRVQT |
| *Silurus meridionalis* | QYEPVAEIGEGAYG--KVYKARDVKNG-RFVAL-KRVRVQT |
| *Simochromis diagramma* | QYEPVAEIGEGAYG--KVYKARDLKNGGRFVAL-KRVRVQT |
| *Siniperca chuatsi* | QYEPVAEIGEGAYG--KVYKARDLKNGGRFVAL-KRVRVQT |
| *Sinocyclocheilus anshuiensis* 1 | QYEPVAEIGEGAYG--KVYKARDLKNGGRFVAL-KRVRVQT |
| *Sinocyclocheilus anshuiensis* 2 | QYEPVAEIGEGAYG--KVYKARDLKNGGRFVAL-KRVRVQT |
| *Sinocyclocheilus grahami* 1 | QYEPVAEIGEGAYG--KVYKARDLKNGGRFVAL-KRVRVQT |
| *Sinocyclocheilus grahami* 2 | QYEPVAEIGEGAYG--KVYKARDLKNGGRFVAL-KRVRVQT |
| *Sinocyclocheilus rhinocerous* 1 | QYEPVAEIGEGAYG--KVYKARDLKNGGRFVAL-KRVRVQT |
| *Sinocyclocheilus rhinocerous* 2 | QYEPVAEIGEGAYG--KVYKARDLKNGGRFVAL-KRVRVQT |
| *Solea senegalensis* | QYEQVAEIGEGAYG--KVYKARDLKNGGRFVAL-KRVRVQT |
| *Sparus aurata* | QYEPVAEIGEGAYG--KVYKARDLKNGGRFVAL-KRVRVQT |
| *Sphaeramia orbicularis* | QYEPVAEIGEGAYG--KVYKARDLKNGGRFVAL-KRVRVQT |
| *Stegastes partitus* | QYEPVAEIGEGAYG--KVYKARDLKNGGRFVAL-KRVRVQT |
| *Syngnathus acus* | QYEPVAEIGEGAYG--KVYKARDLKNGGRFVAL-KRVRVQT |
| *Tachysurus fulvidraco* | QYEPVAEIGEGAYG--KVYKARDLKNG-RFVAL-KRVRVQT |
| *Takifugu flavidus* | RYEAVAEIGEGAYG--KVYKARDLKNEGRFVAL-KKVRVQT |
| *Takifugu rubripes* | RYEAVAEIGEGAYG--KVYKARDLKNEGRFVAL-KKVRVQT |
| *Tetraodon nigroviridis* | QYEQVAEIGEGAYG--KVYKARDLKNGGRFVAL-KKVRVQT |
| *Thalassophryne amazonica* | QYEPVAEIGEGAYG--KVYKARDLKNGGRFVAL-KRVRVQT |
| *Thunnus albacares* | QYEPVAEIGEGAYG--KVYKARDLKNGGRFVAL-KRVRVQT |
| *Thunnus maccoyii* | QYEPVAEIGEGAYG--KVYKARDLKNGGRFVAL-KRVRVQT |
| *Toxotes jaculatrix* | QYEPVAEIGEGAYG--KVYKARDLKNGGRFVAL-KRVRVQT |
| *Trematomus bernacchii* | QYEPVAEIGEGAYG--KVYKARDLKNGGRFVAL-KRVRVQT |
| *Triplophysa tibetana* | QYEPVAEIGEGAYG--KVYKARDLKNGGRFVAL-KRVRVQT |
| *Xiphias gladius* | QYEPVAEIGEGAYG--KVYKARDLKNGGRFVAL-KRVRVQT |
| *Xiphophorus couchianus* | QYEPVAEIGEGAYG--KVYKARDLKNGGRFVAL-KRVRVQT |
| *Xiphophorus hellerii* | QYEPVAEIGEGAYG--KVYKARDLKNGGRFVAL-KRVRVQT |
| *Xiphophorus maculatus* | QYEPVAEIGEGAYG--KVYKARDLKNGGRFVAL-KRVRVQT |

**cartilaginous fish**

| | |
|---|---|
| *Amblyraja radiata* | RYEPVAEIGEGAYG--KVYKARDRNNEGRFVAL-KKVRVQT |
| *Callorhinchus milii* 1 | RYEPVAEIGEGAYG--KVYKARDRNNEGRFVAL-KKVRVQT |
| *Callorhinchus milii* 2 | QYESVAELGAGVYGAATTFKARDLRDQGRFVALLKGLRLQT |
| *Callorhinchus milii* 3 | QYEAVAELGAGVYGAATALKARDLRDEGRFVALLKGLRLQT |
| *Carcharodon carcharias* | RYEPVAEIGEGAYG--KVYKARDRDNEGRFVAL-KKVRVQT |
| *Chiloscyllium plagiosum* | RYEPVAEIGEGAYG--KVYKARDRNNEGRFVAL-KKVRVQT |
| *Hydrolagus affinis* 1 | RYEPVAEIGEGAYG--KVYKARDRNNEGRFVAL-KKVRVQT |
| *Hydrolagus affinis* 2 | QYELLAEVGKGAYG--KVYKARDLKKNGMFVAL-KRIDIPR |
| *Rhincodon typus* | RYEPVAEIGEGAYG--KVYKARDRKNEGRFVAL-KKVRVQT |
| *Scyliorhinus canicula* | RYEPVAEIGEGAYG--KVYKARDRKNEGRFVAL-KKVRVQT |
| *Stegostoma fasciatum* | RYEPVAEIGEGAYG--KVYKARDRNNEGRFVAL-KKVRVQT |

**cyclostome**

| | |
|---|---|
| *Eptatretus burgeri* 1 | QYERVAEIGVGAYG--TVYKARDVQNGGRFVAL-KKVRVHT |
| *Eptatretus burgeri* 2 | RYKRVAAIGHGAQG--TVFKSQDLVN-GQFVAT-KKMRLPT |
| *Eptatretus burgeri* 3 | RYKNISTIGCGAHG--TVFKSQDLVN-GQIVAV-KKIRMPT |
| *Entosphenus tridentatus* | QYEPIAEIGVGAYG--TVYKARDLQNEGRFVAL-KKVRVQT |
| *Lethenteron camtschaticum* | QYEPIAEIGVGAYG--TVYKARDLQNEGRFVAL-KKVRVQT |
| *Lethenteron reissneri* | QYEPIAEIGVGAYG--TVYKARDLQNEGRFVAL-KKVRVQT |
| *Petromyzon marinus* | QYEPIAEIGVGAYG--TVYKARDLQNEGRFVAL-KKVRVQT |

**Figure S5f**

**Cdk4**

| | |
|---|---|
| *Homo sapiens* | RYEPVAEIGVGAYGTVYKARD-PHSGHFVALKSVRVPN |

**Cdk21**

**bony fish**

| | |
|---|---|
| *Acanthochromis polyacanthus* | SYELLAEVGEGCYGKVYKAREVGERQRLLAVKKFNIRG |
| *Acanthopagrus latus* | CYELLAEVGKGSYGKVYKAREAGGKRRLLAVKKFNVHV |
| *Acipenser ruthenus* | DYEILAEIGEGAYGKVYKARHLKNKLDFVAIKKIKLEK |
| *Albula glossodonta* | KYEILVEVGEGAYGKVYKARETRGKQRLVALKKLNIPN |
| *Alosa alosa* | NYDYLAEVGEGAYGKVYKAREIAGKQRLVAVKKLNLPG |
| *Alosa sapidissima* | NYDYLAEVGEGAYGKVYKAREIAGKQRLVAVKKLNLPG |
| *Ameiurus melas* | NYEILAEIGEGAYGKVYKARETGEKQRLVAVKRLNISG |
| *Amphilophus citrinellus* | CYELLAEVGEGYYGKVYKAREVGEKQRLLAVKKFTFRG |
| *Amphiprion ocellaris* | SYELLAEVGEGCYGKVYKAREVGERQRLLAVKKFNIRG |
| *Amphiprion percula* | SYELLAEVGEGCYGKVYKAREVGERQRLLAVKKFNIRG |
| *Anabarilius grahami* | DYEILAEIGQGSYGKVYKARERREQQRLIAVKRLNIPE |
| *Anabas testudineus* | CYELLAEVGQGSYGKVYKAREVGDEQRLLAVKKFIVRG |
| *Anarrhichthys ocellatus* | CYELLAEVGEGSYGKVYKAREVGGKQRLLAVKKFNVRE |
| *Anguilla anguilla* 1 | DYEILVEIGEGAYGKVYKARETKDKQRLVALKKLNIPN |
| *Anguilla anguilla* 2 | DYEILVEIGEGAYGKVYKARETKDKQRLVALKKLNIPN |
| *Anguilla anguilla* 3 | DYEILVEIGEGAYGKVYKARETKDKQRLVALKKLNIPN |
| *Archocentrus centrarchus* | CYELLAEVGEGHYGKVYKAREVGEKQRLLAVKKFTFRG |
| *Astatotilapia calliptera* | CYELLAEVGQGYYGKVYKAREVGEKQRLLAVKKFTIHG |
| *Austrofundulus limnaeus* | SYELLAEVGEGSFGKVYKAREVGGTQRLLGVKKLNCRW |
| *Bagarius yarrelli* | NYEILAEIGQGAYGKVYKARELSEKQRLIAVKRLNISA |
| *Betta splendens* | GYELLAEVGEGTYGKVYRAREVGGNQRLLAVKKFNIHS |
| *Boleophthalmus pectinirostris* | CYELLAEVGEGAYGKVYKAREVGGQQRLLAVKKLNIHP |
| *Caranx melampygus* | RFELLSEVGQGSYGKVFKAREVGGEERLLAVKKFNIHG |
| *Carassius auratus* 1 | DYEILAEIGQGAYGKVYKARERRGQQRFIAVKRLNIPE |
| *Carassius auratus* 2 | DYEMLAEIGQGAYGKVYKARERRGLQRFVAVKRLHIPE |
| *Carassius auratus* 3 | DYEMLAEIGQGAYGKVYKARERRGLQRFVAVKRLHIPE |
| *Channa argus* | CYELLAKVGEGSYGTVYKAREMGEKQRILAVKKLNIRG |
| *Chanos chanos* | NYEILAEIGEGAYGKVYKAREVRERQRLVAVKKLNIAA |
| *Cheilinus undulatus* | CYELLTEVGEGSYGKVYKAREVGGKQRLLAVKKFNIRG |
| *Chelmon rostratus* | CYELLTKLGEGAYGAVYKARETGEKQRLLAVKKFNVRG |
| *Clarias magur* | HYEILAEIAEGAYGKVYKARELGEKQRLVAVKRLNISG |
| *Clupea harengus* | DYEYLAEIGEGAYGKVFKAREIGEKQRLVAVKKLNLPG |
| *Collichthys lucidus* | CYELLAEVGQGSYGKVYKAREAAGQQRLLAVKKFNIRA |
| *Colossoma macropomum* | DYEILAELGEGAYGKVYKAREIGGLQRLVAVKKLNVPG |
| *Coregonus clupeaformis* 1 | NYELLAEIGEGAYGKVYKAREVRDQQRLVAVKKLNIPG |
| *Coregonus clupeaformis* 2 | NYEILAEIGEGTYGKVYKAREVRDQQRLVAVKKLKIHG |
| *Coryphaenoides rupestris* | SYEILVEIGHGAYGKVFKARELREQQRLVAVKKLSVPR |
| *Cottoperca gobio* | CYDLLAEVGEGSYGKVYKAREVGGKQRLLAVKKFNIRG |
| *Cyclopterus lumpus* | CYELLAEVGEGSYGKVYKAREVGEKQRLLAVKKFNIRG |
| *Cynoglossus semilaevis* | SYELLSEVGEGSHGKVYKAREVGANHRYLAVKKFDFRG |
| *Cyprinodon tularosa* | RYELLAEVGEGSFGKVYKAREVGEKRRLLAVKKFNFSM |
| *Cyprinodon variegatus* | RYELLAEVGEGSFGKVYKAREVGEKRRLLAVKKFNFSM |
| *Cyprinus carpio* 1 | DYEILAEIGQGAYGRVYKARERRGQQRFIAVKRLNIPE |
| *Cyprinus carpio* 2 | DYEILAEIGQGAYGKVYKARERRGQQRFIAVKRLNIPE |
| *Danio rerio* | DYEILAEIGQGAYGKVYKAREKREQQRLIAVKRLNIPE |
| *Danionella translucida* | EYEILAEIGQGAYGTVYKGREKRGQQRLIALKRMHSTE |
| *Denticeps clupeoides* | DYEILAEIGEGSYGQVFKARETGQQQRLVAVKKMKLPA |
| *Dicentrarchus labrax* | CYELLAEVGEGSYGKVYKAREAGGKQRLLAVKKFNVCA |
| *Echeneis naucrates* | SYELLAEVGEGSYGKVFKARQVGGDRRLLAVKKFNIRD |
| *Electrophorus electricus* | DYEILAEIGKGAYGKVYKAREVRDRRRLLAVKKLNLPG |
| *Epinephelus coioides* | CYELLAEVGEGSYGKVYKAREAGGTQRLLAVKKFNMLE |
| *Epinephelus lanceolatus* | CYELLAEVGEGSYGKVYKAREAGGTQRLLAVKKFNMLE |
| *Esox lucius* | NYEILAEIGEGAYGKVYKGREVNGQQRLVAVKRLNIPR |
| *Etheostoma cragini* | CYELLAEVGVGSYGRVYKAREVGEKRRLLAVKKFNFRG |
| *Etheostoma spectabile* | CYELLAEVGVGSYGKVYKAREVGEKRRLLAVKKFNFRG |
| *Fundulus heteroclitus* | RYELLTEVGEGSFGRVYKAREVGGKQRLLAVKKFHFSC |
| *Gadus morhua* | SYELLAEIGEGAYGKVFKAREISAEQRLVAIKKLNFSG |
| *Gambusia affinis* | HYELLAEVGEGSFGKVYKAREVGEKQRLLAVKKFNFRW |
| *Gasterosteus aculeatus aculeatus* | CYELLAEVGEGSYGKVYKAREVGDERRLLAVKKFNICG |
| *Girardinichthys multiradiatus* 1 | HYELLAEVGEGSFGKVYKAREIGEKRRLLAVKKFNFRW |
| *Girardinichthys multiradiatus* 2 | HYELLAEVGEGSFGKVYKAREIGEKRRLLAVKKFNFRW |
| *Gouania willdenowi* | RYQLLAEVGKGSYGSVYKARELGGNQRLLAVKKFNIHG |
| *Gymnodraco acuticeps* | CYELLAEVGEGAYGKVYKARELGEKQRFLAVKKFNIRG |
| *Haplochromis burtoni* | CYELLAEVGQGYYGKVYKAREVGEKQRLLAVKKFTIHG |
| *Hippocampus comes* | FYELLTEIGQGSYGKVYKAREVGEGGRLLAVKKFHFCS |
| *Hippoglossus hippoglossus* | CYELLAAVGKGSFGTVYKAREVGDEPRFLAVKKFKIYG |
| *Hippoglossus stenolepis* | CYELLAAVGKGSFGTVYKAREVGDEPRFLAVKKFKIYG |
| *Hucho hucho* | NYEILAEIGEGAYGKVYKAREVRDQQRLVAVKKLNIPG |
| *Hypomesus transpacificus* | NYEILVEIGEGAYGKVYKAREVRDQQRLVAVKKLNIPK |
| *Ictalurus punctatus* | NYEILAEIGEGAYGKVYKARETGEKQRLVAVKRLNIGA |
| *Kryptolebias marmoratus* | SYELLAEVGQGSFGRVFRAREVGGAQRLLAVKKFNCRW |
| *Labeo rohita* | DYEILAEIGQGAYGKVYKARERRGQQRLVAVKKLNIPE |
| *Labrus bergylta* | CYELLAEVGEGSYGTVYKAREVGGKQRLLAVKKFNIRG |
| *Larimichthys crocea* | CYELLAEVGVGSYGKVYKAREAAGKQRLLAVKKFNIRG |
| *Lates calcarifer* | CYELLSEVGEGSYGKVYKAREVGGTQRLLAVKKFNIRG |
| *Latimeria chalumnae* | NYELQAVIGEGAYGKVYKARDL--DGRFVALKKMNLFS |
| *Lepisosteus oculatus* | EYERLAEVGEGAYGKVYKARETQGQRRLVALKKLSVAG |
| *Liparis tanakae* | RYELLVEVGQGSYGKVYKARQLGGQQRLLAVKKFNIRG |
| *Lucifuga dentata* | NYDILVQIGEGSYGKVYKAREVGGQQRLLAVKKFNITG |
| *Mastacembelus armatus* | RYELLAEVGEGSYGKVYKAREVGEKHRLLAVKKFNIRG |
| *Maylandia zebra* | CYELLAEVGQGYYGKVYKAREVGEKQRLLAVKKFTIHG |
| *Megalobrama amblycephala* | DYEILAEIGQGSYGKVYKARERREQQRLIAVKRLNVPE |
| *Megalops atlanticus* | DYEILVDIGEGAYGKVYKARETKGQQRLVALKKLNIPN |
| *Megalops cyprinoides* | DYEILVDIGEGAYGKVFKARETKGQQRLVALKKLNIPN |
| *Melanotaenia boesemani* | SYELLTEVGEGSFGKVYKAREVGEKQRLLAVKKFHLSC |
| *Micropterus dolomieu* | CYELLAEVGEGSYGKVYKAREVGEKQRILAVKKFNIHG |
| *Micropterus salmoides* | CYELLAEVGEGSYGKVYKAREVGEKQRILAVKKFNIHG |
| *Mola mola* | TYELLTKLGEGAYGQVYKGREVGGKQRLLAVKKFNIPG |
| *Monopterus albus* | SYELLVEIGKGSYGKVYKAREMGEKQRLLAVKKFNTRV |
| *Morone saxatilis* | CYELLAEVGEGSYGKVYKAREAGGKQRLLAVKKFNVRA |
| *Mugil cephalus* | PYELLVEVGQGSYGKVYKAREVGGKQRLLAVKKYNVRG |
| *Myripristis murdjan* | SYEILAQVGEGTYGKVYKARELGERQRLLAVKKLNIPG |
| *Nematolebias whitei* | SYEVLAEVGEGSFGKVYKAREVGGTQRLLGVKKVNCRW |
| *Neogobius melanostomus* | CYELLAEIGEGSYGKVYKAREVGGKQRLLAVKKINIRP |
| *Neolamprologus brichardi* | SYELLAEVGQGYYGKVYKAREVGEKQRLLAVKKFTIHG |
| *Nibea albiflora* | CYELLAEVGEGSYGKVYKAREAGGKQRLLAVKKFIIRG |
| *Nothobranchius furzeri* | SYELLAEVGEGSFGKVYKARELGEKQRLLAVKKFNFQI |
| *Nothobranchius kuhntae* | SYELLAEVGEGSFGKVYKARELGEKQRLLAVKKFNFQI |
| *Notolabrus celidotus* | CYELLAEVGEGTYGTVYKAREVGGTQRLLAVKKFNFHG |
| *Notothenia coriiceps* | CYELLAEVGEGSYGKVYKARELGEKQRFLAVKKFNVRR |
| *Oncorhynchus gorbuscha* | NYEILAEIGEGAYGKVYKAREVRDQQRLVAVKKLNIPG |
| *Oncorhynchus keta* | NYEILAEIGEGAYGKVYKAREVRDQQRLVAVKKLNIPG |
| *Oncorhynchus kisutch* | NYEILAEIGEGAYGKVYKARAVRDQQRLVAVKKLNIPG |
| *Oncorhynchus mykiss* | NYEILAEIGEGAYGKVYKAREVRDQQRLVAVKKLNIPG |
| *Oncorhynchus nerka* | NYEILAEIGEGAYGKVYKAREVRDQQRLVAVKKLNIPG |
| *Oncorhynchus tshawytscha* | NYEILAEIGEGAYGKVYKAREVRDQQRLVAVKKLNIPG |
| *Onychostoma macrolepis* | DYEILAEIGQGAYGKVYKARERRGQQRFVAVKRLHIPE |
| *Oreochromis aureus* | CYELLAEVGQGYYGKVYKAREVGEKQRLLAVKKFTIHG |
| *Oreochromis niloticus* | CYELLAEVGQGYYGKVYKAREVGEKQRLLAVKKFTIHG |
| *Oryzias javanicus* | SYELLAEVGEGSFGTVYKAREIGGKQRLLAVKKLNFRV |
| *Oryzias latipes* | RYELLAEVGEGSFGRVYKAREIGEKQRLLAVKKLNFSL |
| *Oryzias melastigma* | SYELLAEVGEGSFGRVYKAREIGGKQRLLAVKKLNFRL |
| *Oryzias sinensis* | RYELLAEVGEGSFGRVYKAREIGEKQRLLAVKKLNFSL |
| *Pangasianodon hypophthalmus* | NYEILAEIGEGAYGKVYKAREMGDKQRLVAVKRLNISG |
| *Paralichthys olivaceus* | SYQLLAAVGKGSFGTVYKARELGDKQRFLAVKKFTIHG |
| *Parambassis ranga* | SYELLAEVGQGAYARVYKAREVGDTRRLLAVKKFIIRA |
| *Paramormyrops kingsleyae* | DYEILVEIGEGAYGKVYKARAIREPQRLVAVKKLNIPV |
| *Perca flavescens* | CYELLAEVGVGSYGKVYKAREVGEKRRLLAVKKFNIRG |
| *Perca fluviatilis* | CYELLAEVGVGSYGKVYKAREVGEKRRLLAVKKFNIRG |
| *Periophthalmus magnuspinnatus* | CYELLAEIGEGSYGKVYKAREVGGQQRLLAVKKLNIRA |
| *Pimephales promelas* | DYEFLAEIGQGSYGKVYKARERRDKQRLIAVKRLNLPE |
| *Plectropomus leopardus* | CFELLVEIGEGSYGKVYKAREVGEKQRLLAVKKFNIRG |
| *Poecilia formosa* | HYELLAEVGEGSFGKVYKAREVGEKQRLLAVKKFNFRW |
| *Poecilia latipinna* | HYELLAEVGEGSFGKVYKAREVGEKQRLLAVKKFNFRW |
| *Poecilia mexicana* | HYELLAEVGEGSFGKVYKAREVGEKQRLLAVKKFNFRW |
| *Poecilia reticulata* | HYELLAEVGEGSFGKVYKAREVGEKQRLLAVKKFNFRW |
| *Poeciliopsis prolifica* | -----AEVGGGSFGKVYKAREVGEKQRLLAVKKFNFRW |
| *Protopterus annectens* | GYEPLAEIGEGAYGKVFKARDLKNEGRFVALKKVNVCS |
| *Pseudochaenichthys georgianus* | CYELLAEVGEGSYGKVYKARELGEKQRFLAVKKFNIRG |
| *Pundamilia nyererei* | CYELLAEVGQGYYGKVYKAREVGEKQRLLAVKKFTIHG |
| *Pungitius pungitius* | CYELLAEVGEGSYGKVYKAREVGEKRRLLAVKKFNIRG |
| *Puntigrus tetrazona* | DYEILAEVGQGAYGKVYKARERRGKQRFIAVKRLNIPE |
| *Pygocentrus nattereri* | DYEILAEIGEGAYGKVYKAREIGGLQRLVAMKKINVPG |
| *Salarias fasciatus* | RFQLLAEVGKGSYGKVFKAREVGGEQRLLAVKKFNIRG |
| *Salmo salar* | NYEILAEIGEGAYGKVYKAREVRDQQRLVAVKKLNIPG |
| *Salmo trutta* | NYEILAEIGEGAYGKVYKAREVRNQQRLVAVKKLNIPG |
| *Salvelinus alpinus* | NYEILAEIGEGAYGKVYKAREVRDQQRLVAVKKLNIPG |
| *Salvelinus namaycush* | NYEILAEIGEGAYGKVYKAREVRDQQRLVAVKKLNIPG |
| *Sander lucioperca* | CYELLAEVGVGSYGKVYKAREVGEKRRLLAVKKFNIRG |
| *Scatophagus argus* | CYELLAEVGEGSYGKVFKAREAGQKQRLLAVKKFKIRS |
| *Scleropages formosus* 1 | CYEVLAAVGEGASGRVFKARETGGCQRRVALKKVRVLE |
| *Scleropages formosus* 2 | HYEVLAAVGEGASGRVFKARETGGCQRRVALEKVRVLE |
| *Scleropages formosus* 3 | RYEVLAAVGEGASGRVFKARETGGCQCLVALKKVRVLK |
| *Scleropages formosus* 4 | RYEVLAAVGEGASGRVFKARETGGCQRLVALKKVRVLE |
| *Scleropages formosus* 5 | RYEVLAAVGEGASGRVFKARETGGCQRLVALKKVRVLE |
| *Scleropages formosus* 6 | RYEVLAAVGEGASGRVFKARETGGCQRRVALEKVRVLE |
| *Scleropages formosus* 7 | RYEVLAAVGEGASRRVFKARETGGCQRLVALKKVRVLE |
| *Scophthalmus maximus* | RYELLAEVGQGSHGRVYRARRLGGDQRLLAVKKFNVPA |
| *Sebastes umbrosus* | SYELLAAVGEGSYGKVFKAREVEGRQRLLAVKKFNIRG |
| *Seriola dumerili* | CYELLSEVGEGSYSKVYKAREVGEKQRLLAVKKFNIHG |
| *Seriola lalandi dorsalis* | CYELLSEVGEGSYSKVYKAREVGEKQRLLAVKKFNIHG |
| *Silurus meridionalis* | NYEILAEIGAGAYGTVYKARDLVDKQRLVAVKKLKISA |
| *Simochromis diagramma* | CYELLAEVGQGYYGKVYKAREVGEKQRLLAVKKFTIHG |
| *Siniperca chuatsi* | CYELLAEVGEGSYGKVYKAREVGEKQRLLAVKKFNIRG |
| *Sinocyclocheilus anshuiensis* 1 | DYEILAEIGQGAYGKVYKARERRGQQRFIAVKRLNIPE |
| *Sinocyclocheilus anshuiensis* 2 | DYEILAEIGQGAYGKVYKARERRGQQRFVAVKRLNIPE |
| *Sinocyclocheilus grahami* 1 | DYEILAEIGQGAYGKVYKARERRGQQRFIAVKRLNIPE |
| *Sinocyclocheilus grahami* 2 | DYEILAEIGQGAYGKVFKARERRGQQRFIAVKRLNIPE |
| *Sinocyclocheilus rhinocerous* 1 | DYEIWAEIGQGAYGKVYKARERRGQQRFIAVKRLNIPE |
| *Sinocyclocheilus rhinocerous* 2 | DYEILTEIGQGAYGKVYKARERRGQQRFIAVKRLNIPE |
| *Solea senegalensis* | CFELLSEVGEGSYAKVYKAREVGEEQRLLAVKKFNFHA |
| *Sparus aurata* | CYELLAEVGKGSYGKVYKAREAGGKRRLLAVKKFNVHV |
| *Sphaeramia orbicularis* | TYDLLAEVGHGSYGTVYKARQVGGEQRLLAVKKLNVRA |
| *Stegastes partitus* | DYELLAEVGVGCYGKVYKAKEVGDRQRLLAVKKFSIRG |
| *Syngnathus acus* | LYELLAEIGQGSFGKVYKARAVGEEGRLLAVKKFHFCS |
| *Tachysurus fulvidraco* | NYEILAEIGQGAYGKVYKARELGEKQRLVAVKKLNIST |

| | |
|---|---|
| *Takifugu rubripes* | CYELLAEIGEGSYGKVYKAREEGGKQRLLAVKKFNFRG |
| *Tetraodon nigroviridis* | -YELLAEVGEGTYGKVYKAREEGGEQRLLAVKKLHFRG |
| *Thalassophryne amazonica* | RYEILAKVGEGSYSQVFKARETGEQQRLLAVKKFTIQG |
| *Thunnus albacares* | CYELLAEVGEGSYGKVYKAREVGEKQRLLAVKKFNIRG |
| *Thunnus maccoyii* | CYELLAEVGEGSYGKVYKAREVGEKQRLLAVKKFNIRG |
| *Toxotes jaculatrix* | RYELLAEVGQGSYGKVYKAREVGGKERLLAVKKFNIRG |
| *Trematomus bernacchii* | CYELLAEVGEGSYGKVYKARELGEKQRFLAVKKFNIRG |
| *Triplophysa tibetana* 1 | DYEILAEIGQGAYGKVYKAREVRDRQRLVAVKRLNISE |
| *Triplophysa tibetana* 2 | DYEILAEIGQGAYGKVYKAREVRDRQRLVAVKRLNISE |
| *Xiphias gladius* | RYELLSGVGEGSYGTVYKAREVGGEQRLLAVKKFNLHR |
| *Xiphophorus couchianus* | HYELLAEVGEGSFGKVYKAREVGEKQRLLAVKKLNFRW |
| *Xiphophorus hellerii* | HYELLAEVGEGSFGKVYKAREVGEKQRLLAVKKLNFRW |
| *Xiphophorus maculatus* | HYELLAEVGEGSFGKVYKAREVGEKQRLLAVKKLNFRW |
| **cartilaginous fish** | |
| *Amblyraja radiata* | RYELLAEIGEGAYGKVYKARDLENNGKCVALKRIVVPR |
| *Callorhinchus milii* | QYELLAEIGKGAYGKVYKARDLKNNGKFVALKRIEIPR |
| *Carcharodon carcharias* | RYELMAEIGEGAYGKVYKARDLENNGRFVALKRIEIPR |
| *Chiloscyllium plagiosum* | RYELLAEIGEGAYGKVYKARDLENNGRFVALKRIEIPR |
| *Rhincodon typus* | RYELLAEIGEGAYGKVYKARDLENNGRFVALKRIEIPR |
| *Scyliorhinus canicula* | RYELIAEIGEGAYGKVYKARDLENNGRFVALKRIEIPR |
| *Stegostoma fasciatum* | RYELLAEIGEGAYGKVYKARDLENNGRFVALKRIEIPR |

### Figure S5g

**Figure S5** Cdk4/6/21 amino acid alignment around the residue corresponding to R24/R31 in vertebrates. Cdk4 in (a) mammals, (b) non-mammalian/non-fish vertebrates, and (c) fish; Cdk6 in (d) mammals, (e) non-mammalian/non-fish vertebrates, and (f) fish; Cdk21 in (g) fish. The residue corresponding to Cdk4/6 R24/31 is in green if the strong positive charge is conserved and in magenta, if not.
