## Supplementary Tables S1-S11 and Figures S1-S6 for "Evolution of the *Cdk4/6*–*Cdkn2* system in invertebrates": Figure S6.pdf

**(a)**

| Phyla | Number of species in which Cdk4/6 was detected **[1]** | Number of species in which both Cdkn2 and Cdk4/6 were detected **[2]** | **[2]** / **[1]** ( *y* -axis) | ratio of the number of species with R or K at Cdk4/6 R24/31 ( *x*-axis ) |
|---|---|---|---|---|
| Urochordata | 11 | 0 | 0.000 | 0.818 |
| Echinodermata | 15 | 15 | 1.000 | 1.000 |
| Arthropoda | 898 | 0 | 0.000 | 0.963 |
| Nematoda | 66 | 0 | 0.000 | 0.379 |
| Rotifera | 7 | 0 | 0.000 | 0.714 |
| Annelida | 7 | 4 | 0.571 | 1.000 |
| Mollusca | 55 | 16 | 0.291 | 1.000 |
| Platyhelminthes | 40 | 0 | 0.000 | 0.025 |
| Cnidaria | 56 | 0 | 0.000 | 0.929 |

**(b)**

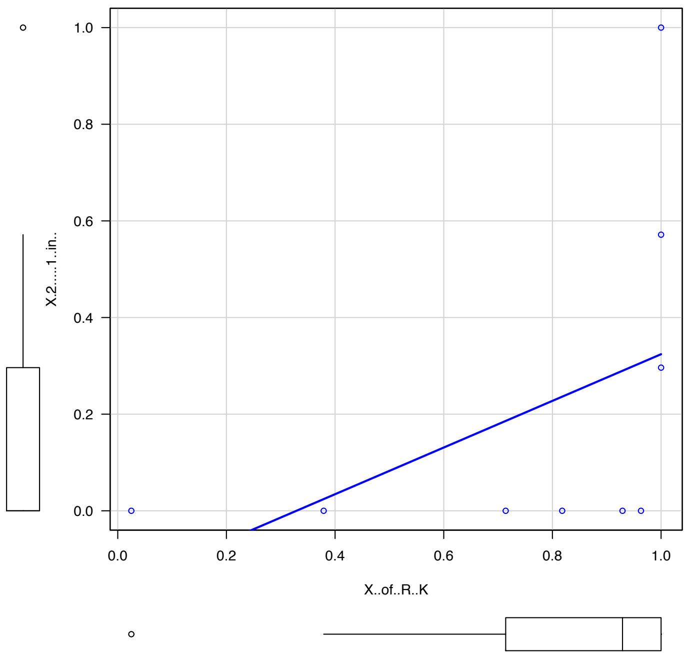

**Figure S6** Determination of the Spearman's rank correlation coefficient. (a) The data used and (b) the scatter plot.
